# Biosynthetic allene and alkyne formation by enzymatic prenyl demethylation

**DOI:** 10.64898/2026.05.26.727785

**Authors:** Mengting Liu, Masao Ohashi, Wenyu Han, Qingyang Zhou, Kendall N. Houk, Yi Tang

## Abstract

Allenes and alkynes are versatile functional groups in total synthesis, medicinal chemistry, and bioorthogonal conjugation. The biosynthetic logic of how Nature installs allene or alkyne in natural products, especially that of allenes, is not well understood. Here we uncovered allenes and alkynes can be formed enzymatically through oxidative C(*sp*^2^)-demethylation of the common five-carbon prenyl group. Two fungal cytochrome P450 monooxygenases, PpnB and NseB, from the penipratynolene and sinuxylamide biosynthetic pathways, respectively, were shown to catalyze oxidative removal of a C(*sp*^2^)-methyl group in *O*-prenyl-L-tyrosine to afford *O*-homoallenyl-L-tyrosine and *O*-but-2-ynyl-L-tyrosine, respectively. Combining density functional theory calculations, heterologous expression, biotransformation and enzymatic assays with isotopically labeled substrates, a mechanism involving selective C–C bond cleavage followed by product-determining hydrogen atom abstraction is presented. An additional P450 enzyme from the penipratynolene pathway, PpnD, acts as an oxidative isomerase that converts the four-carbon terminal allene into a terminal alkyne. This unprecedented enzymatic editing strategy to install allene and alkyne expands the catalytic repertoire of P450 enzymes.

**Graphical abstract:** 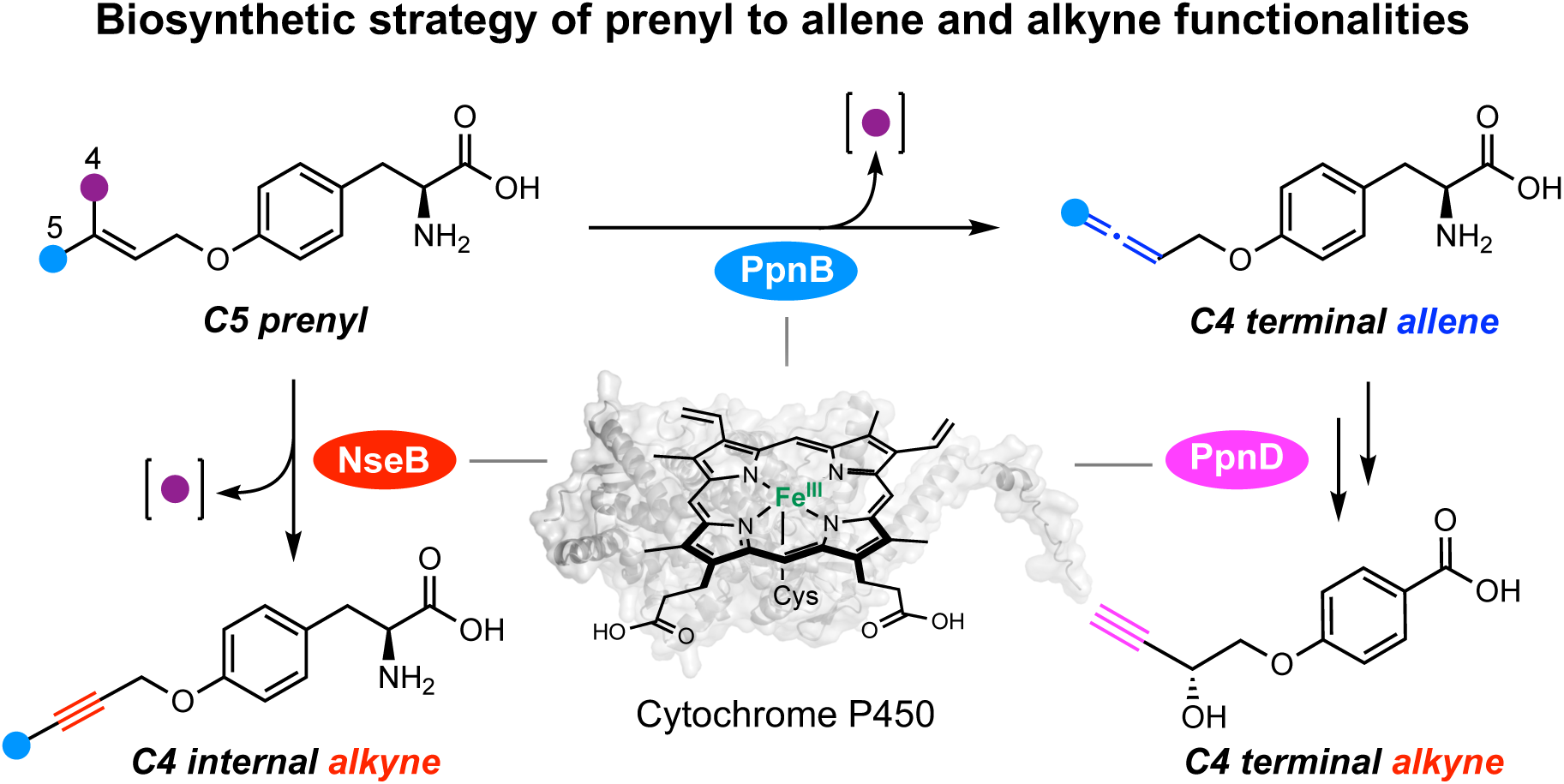

## Main

Allenes and alkynes are versatile functional groups useful in synthetic applications and bioorthogonal chemistry^1–12^. In particular, terminal and strained alkynes can react with azides via the 1,3-dipolar cycloaddition widely known as Click Chemistry^8,9,13,14^. Click reactions using allenes have also been developed, such as bisphosphine–copper-catalyzed phenoxydiazaborinine formation (CuPDF) using terminal allenes^10,11^. Installation of alkynes, and to a lesser extent, allenes, into small molecules have therefore become a mainstay in drug discovery and chemical biology^15–18^. While organic synthesis remains the primary method to introduce these functionalities, enzymatic installations that are compatible with cellular conditions remain rare^19,20^. More than 150 natural products containing allene moieties have been reported^21,22^, exemplified by the terpenoid ginamallene^23^; the cyclic peptide pseudoxylallemycin B that contains *O-*homoallenyl*-*L*-*Tyr (**1**)^24^; and the benzyl alcohol derived terricollene C^25^ (**3**). Over 2000 natural products bearing alkyne functionality are known across various kingdoms of life, including sinuxylamide B^26^ (**2**) and terriolyne^25^ (**4**) (**Fig. 1a** and **Supplementary Figs. 1 and 2**)^27^. These allene and alkyne functional groups are important for the biological activities of the natural products^15,21,22,27^. Despite such prevalence of alkynes in natural products, only a few alkyne-forming biosynthetic enzymes have been identified and characterized (**Supplementary Fig. 3**)^20,28–33^. The sole example of an allene-forming enzyme is the algal violaxanthin de-epoxidase-like enzyme VDL1 that catalyzes the conversion of violaxanthin to neoxanthin via 1,2-elimination (**Supplementary Fig. 3**)^34^. Therefore, new allene- and alkyne-forming enzymes can be discovered from uncharacterized biosynthetic pathways, such as those leading to **1** and **2**.

**Figure 1.**
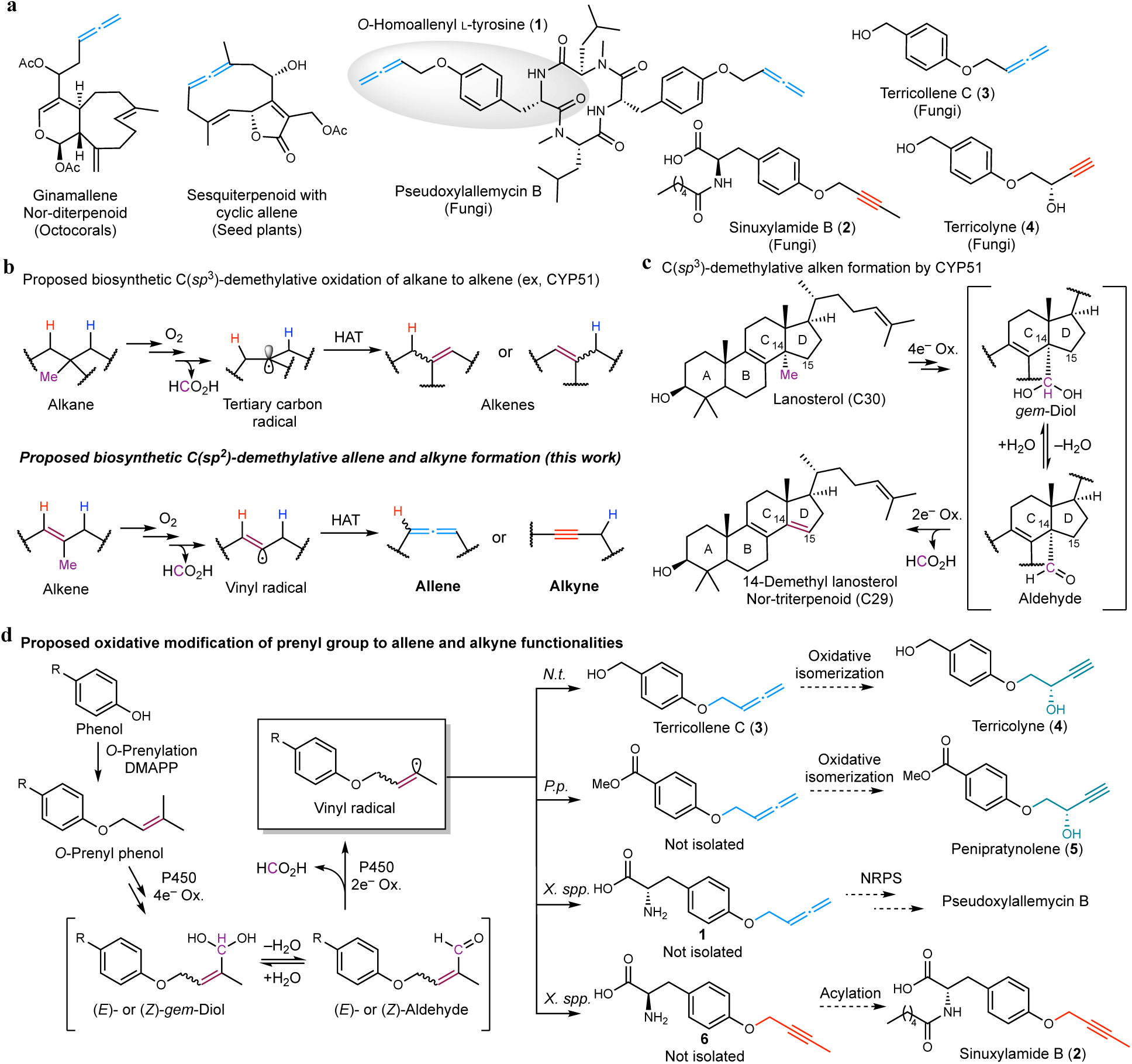
Discovery of allene and alkyne-forming enzymes from natural product biosynthesis. **a**, Structures of selected allene and alkyne-containing natural products of which biosynthesis are uncharacterized. **b**, Proposed allene and alkyne formation via oxidative C(*sp*^2^)-demethylation (this work). **c**, Lanosterol 14-α-demethylase CYP51 catalyzes oxidative C(*sp*^3^)-demethylation of lanosterol to form 14-demethyl lanosterol. **d**, Proposed biosynthetic formation of allene and alkyne from prenyl group in the phenol-containing natural products such as terricollene C (**3**), penipratynolene (**5**), pseudoxyallemycin B, and sinuxylamide B (**2**). By following CYP51-catalyzed demethylative alkene formation, the reaction can be initiated by the oxygenation of the terminal (*E*)- or (*Z*)-methyl group in the prenyl moiety to the *gem*-diol or the aldehyde, which can be oxidatively transformed into allene and alkyne. The producing organisms for the corresponding compounds are shown above the arrow; *N.t.* (*Neurospora terricola*), *P.p.* (*Penicillium polonicum*), and *X.*spp (*Xylaria* species). The proposed oxidation of allene to α-hydroxy alkyne for the formation of **4** and **5** are shown.

**Figure 2.**
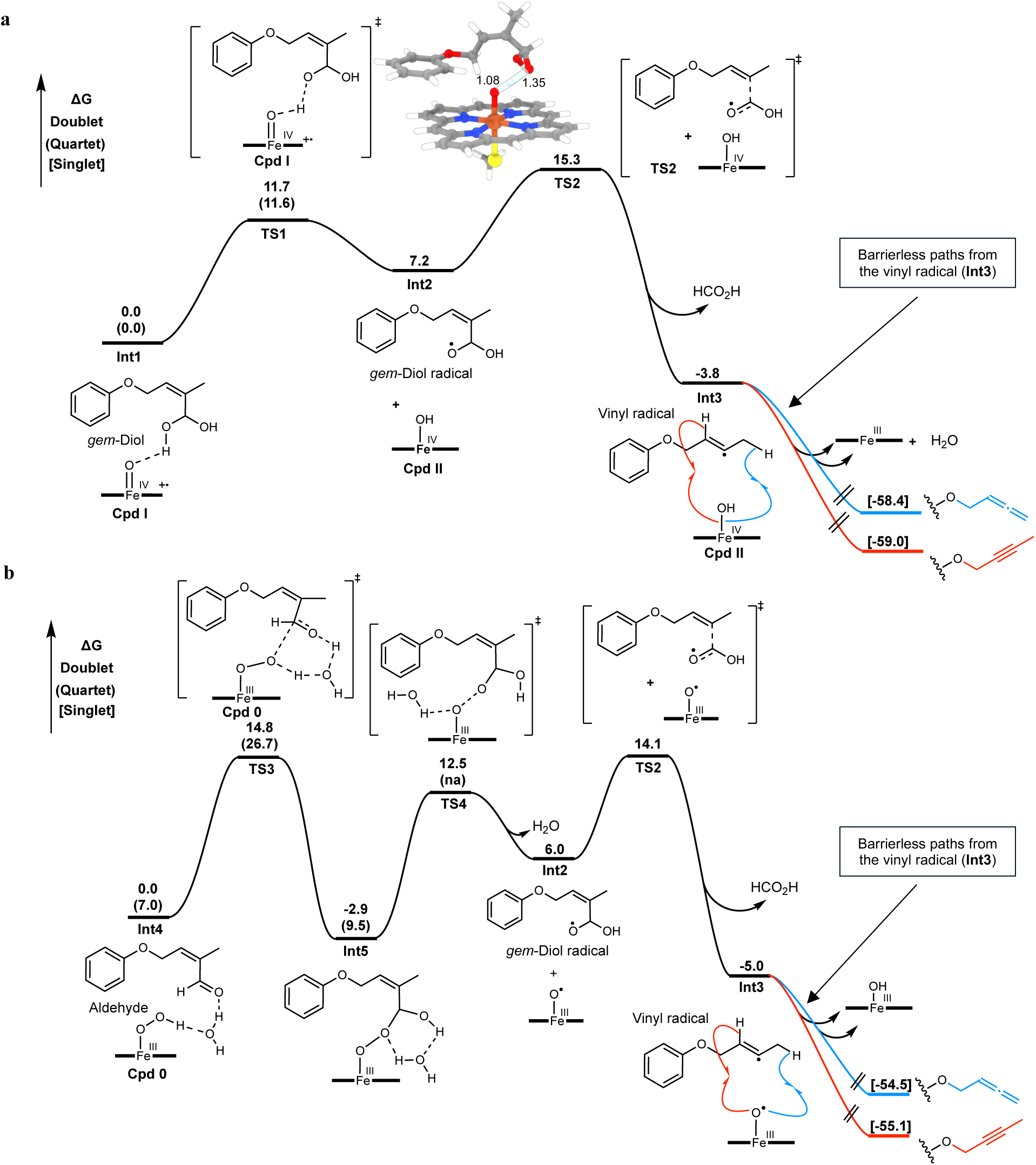
Theoretical validation of allene and alkyne formation from prenyl group. **a**, Potential mechanism initiated by Cpd I with the (*Z*)-*gem*-diol. **b**, Potential mechanism initiated by Cpd 0. The energies were calculated by B3LYP-D3(BJ)/def2-TZVP/SMD(Et_2_O)//B3LYP-D3(BJ)/def2-SVP/IEFPCM(Et_2_O) at 298 K, 1 atm and are given in kcal/mol. Bond distances are labeled in angstrom. For all iron-containing species, possible spin states with free energy are reported.

Although not reported in chemical or biological synthesis, one plausible route to allene and alkyne functionalities could initiate from oxidative C–C bond cleavage of substituted alkenes (**Fig. 1b**). This is a challenging transformation as unstrained C–C bonds are among the least reactive bonds in organic chemistry due to the high bond dissociation energy (BDE) (∼90 kcal/mol), weak polarization and poor interactions with metal catalysts^35–37^. In primary and secondary metabolisms, enzymatic C–C bond cleavage is deployed in the biosynthesis of small molecules, such as terpenoids and oxylipins^38^. One prominent example is lanosterol-14-α-demethylase (CYP51) that converts triterpenoid lanosterol to 14-demethyl-lanosterol, via C–C bond fragmentation of a proposed C_14_ *gem*-diol/aldehyde intermediate (**Fig. 1c** and **Extended Data Fig. 1**)^38–44^. Different mechanisms of C–C bond cleavage with either homolytic or heterolytic cleavage pathway by the heme center of CYP51 have been proposed^42–45^. The heterolytic cleavage pathway involving Baeyer–Villiger (BV)-type intermediates has been extensively discussed with the experimental detection of a putative BV intermediate during CYP51 catalysis^42,45^. On the other hand, theoretical studies proposed the mechanisms with homolytic cleavage pathways^43,44^, including (i) hydrogen abstraction from the C_14_ *gem*-diol by Cpd I (high valent iron oxo species) to generate a *gem*-diol radical; or (ii) nucleophilic attack of Cpd 0 (Fe(III)–peroxo species) on the C_14_ aldehyde to form a Criegee intermediate. In both homolytic cleavage proposals, subsequent C–C bond cleavage yields the same C_14_ tertiary carbon radical and formic acid, followed by a second hydrogen atom transfer (HAT) step to the heme to form the C_14_–C_15_ double bond in 14-demethyl lanosterol. The overall transformation is oxidation of an alkane to an alkene via excision of a C(*sp*^3^)-methyl group (**Fig. 1b**).

Inspired by the CYP51-catalyzed demethylative oxidation, we propose that an analogous oxidative demethylation of a C(*sp*^2^)-methyl substituted alkene could generate an allene or an alkyne (**Fig. 1b**). In nature, the five-carbon (C5) prenyl (dimethylallyl) group is the most abundant methyl-substituted olefin. The thermodynamically activated prenyl precursor, dimethylallyl diphosphate (DMAPP), can be transferred to different nucleophiles in a prenylation reaction^46^. Four-electron oxidation of the terminal methyl group can first yield the aldehyde or *gem-*diol intermediate (**Fig. 1d**). The subsequent oxidative homolytic C–C bond cleavage can generate a four-carbon (C4) vinyl radical intermediate, which can be followed by an allyl HAT to afford a terminal allene, as present in **3**^25^ and **1**^24^; or a vinyl HAT to arrive at an internal alkyne as present in **2** (**Fig. 1d**)^26^. Subsequent oxidative isomerization of C4 terminal allene could also account for the biosynthesis of related compounds with a α-hydroxy alkyne, such as **4** and penipratynolene (**5**) (**Fig. 1d**). In this work, we examined the biosynthetic pathways of **5** and **2** isolated from fungi, and discovered a fungal P450 enzyme in each pathway that catalyzes the oxidative C(*sp*^2^)-demethylation of *O*-prenyl-L-Tyr (**7**) to give *O-*homoallenyl-L-Tyr (**1**) and *O*-but-2-ynyl-L-Tyr (**6**), respectively. In the biosynthetic pathway of **5**, a second P450 enzyme catalyzes the oxidative isomerization of a C4 terminal allene to a terminal alkyne. Because prenylation is a prevalent modification of both small molecules and macromolecules, our findings have the potential for *in situ* generation of allene and alkyne containing biomolecules for chemical biology applications.

## Results

### Validation of the Proposed Mechanism by DFT Calculations

The proposed P450-catalyzed oxidative C(*sp*^2^)-methyl excision from a prenyl group to form allene or alkyne involves the generation of a vinyl radical (**Fig. 1b**). This radical is less stable than the tertiary carbon radical proposed in the CYP51 reaction (C–H BDEs are 111 and 96-98 kcal/mol, respectively)^37^. To assess the energetics of the vinyl radical intermediate, density functional theory (DFT) calculations were performed using heme-bound iron as the oxidant. The terminal aldehyde or the hydrated form (*gem*-diol) of *O*-prenylated phenol was used as model substrates (**Fig. 2**). The reaction pathway for Cpd I-dependent mechanism starting from (*Z*)-*gem*-diol is shown in Fig. 2a, while Cpd 0-dependent mechanism starting from the aldehyde is shown in Fig. 2b. Our DFT calculations indicate that C–C bond cleavage could proceed via the same *gem*-diol radical intermediate (**Int2**). In the Cpd I-dependent mechanism, binding of the *gem*-diol substrate to Cpd I forms **Int1** is followed by hydrogen atom abstraction from one hydroxyl group by Cpd I to give the *gem*-diol radical **Int2** via **TS-1** (Δ*G*^‡^ = 11.7 kcal/mol) and Cpd II. On the other hand, in the Cpd 0-dependent mechanism, the reaction initiates with a water-assisted direct addition of the peroxide oxygen from Cpd 0 to the heme-bound aldehyde (**Int4**) via **TS3** (Δ*G*^‡^ = 14.8 kcal/mol) to form a Criegee-like intermediate (**Int5**). This adduct can subsequently undergo homolysis of the peroxide O–O bond via **TS4** (Δ*G*^‡^ = 15.4 kcal/mol) to yield the *gem*-diol radical **Int2** and a ferric-oxo radical (Fe(III)-O•).

In both mechanisms, C–C bond fragmentation from **Int2** via **TS2** (Δ*G*^‡^ = 8.1 kcal/mol) can occur readily to form the vinyl radical **Int3** with release of formic acid (Δ*G°= –*11.0 kcal/mol). Although vinyl radicals are generally considered unstable, the thermodynamic driving force here arises from the release of formic acid, a stable, closed-shell molecule, which provides a favorable enthalpic contribution, combined with the entropic gain of this fragmentation step. Previous QM/MM calculations of the CYP51 mechanism predicted formation of the tertiary carbon radical is substantially more exothermic (-34.4 kcal/mol for the Cpd I–dependent mechanism and -44.0 kcal/mol for the Cpd 0–dependent mechanism), which is expected for the more stable tertiary radical^43,44^. Our calculation indicates that generation of the vinyl radical remains both kinetically and thermodynamically feasible. The fate of **Int3** is determined by the regioselectivity of the second HAT step to Cpd II (Cpd I mechanism) or Fe(III)-O• (Cpd 0 mechanism): abstraction from C(*sp^2^*)-H to form alkyne or from terminal C(*sp^3^*)-H to form allene. Both pathways are calculated to be barrierless on the potential energy surface (**Fig. 2** and **Supplementary Fig. 4**), therefore indicating that HAT regioselectivity should be determined by the binding conformation of **Int3** with respect to the heme-iron center in the P450 active site. The rate-determining step of the overall reaction is predicted to be generation of **TS2** (Δ*G*^‡^ = 15.3 kcal/mol) or **TS4** (Δ*G*^‡^ = 15.4 kcal/mol), depending on the oxidizing heme species. Previously computed rate-determining barriers for the CYP51 reaction are 3.9 kcal/mol for a Cpd I-dependent mechanism and 17.3 kcal/mol for a Cpd 0-dependent mechanism^43,44^. Notably, although **TS2** and **TS4** are predicted to have similar reaction barriers, the actual rate is also controlled by the relative stability of *gem*-diol versus aldehyde. As the aldehyde substrate is stabilized by conjugation with the adjacent C=C double bond, the hydration energy is calculated to be +4.4 kcal/mol (**Supplementary Fig. 27**). Therefore, the Cpd 0–dependent route with aldehyde would be overall more favored, but this preference could also be readily overturned by the actual enzyme environment, where the *gem*-diol form might benefit from additional hydrogen bonds with surrounding residues. Overall, calculations indicate that both pathways are plausible, and support the feasibility of P450-catalyzed C(*sp*^2^)-demethylative formation of allenes and alkynes via the vinyl radical intermediate **Int3**.

### Identification of Biosynthetic Gene Cluster for Allene Formation

Based on the mechanistic hypothesis, we reasoned that a multifunctional P450 enzyme and a prenyltransferase (PT) should be involved in formation of a C4 allene starting from a phenolic substrate (**Fig. 1d**). Terricollene C (**3**) and terricolyne (**4**) were co-isolated from the endolichenic fungus *Neurospora terricola*^25^, and the α-hydroxy alkyne in **4** can be formed via oxidation of the allene in **3** (**Fig. 1a, d**). However, due to unavailability of *N. terricola* and genome information, we instead targeted 4-hydroxybenzoate methyl ester analog of terricolyne, penipratynolene (**5**) isolated from *Penicillium polonicum*^47–49^. Although the allene analog of **5** was not isolated, we propose that allene formation should precede the formation of **5** (**Fig. 1d**). Using clustering of genes encoding PT and P450 as a criterion, the *ppn* BGC from *P. polonicum* was identified as the top candidate. The BGC encodes an ABBA-type PT (PpnA), two P450 enzymes (PpnB and PpnD), an aromatic L-amino acid ammonia-lyase (AAL, PpnC)^32^, a methyltransferase (MT, PpnE), a NAD(P)H-dependent oxidoreductase PpnF, and a berberine bridge enzyme (BBE)-like enzyme PpnG (**Fig. 3a** and **Supplementary Table 1**). The P450 enzymes PpnB and PpnD do not show high sequence similarity to known P450s.

**Figure 3.**
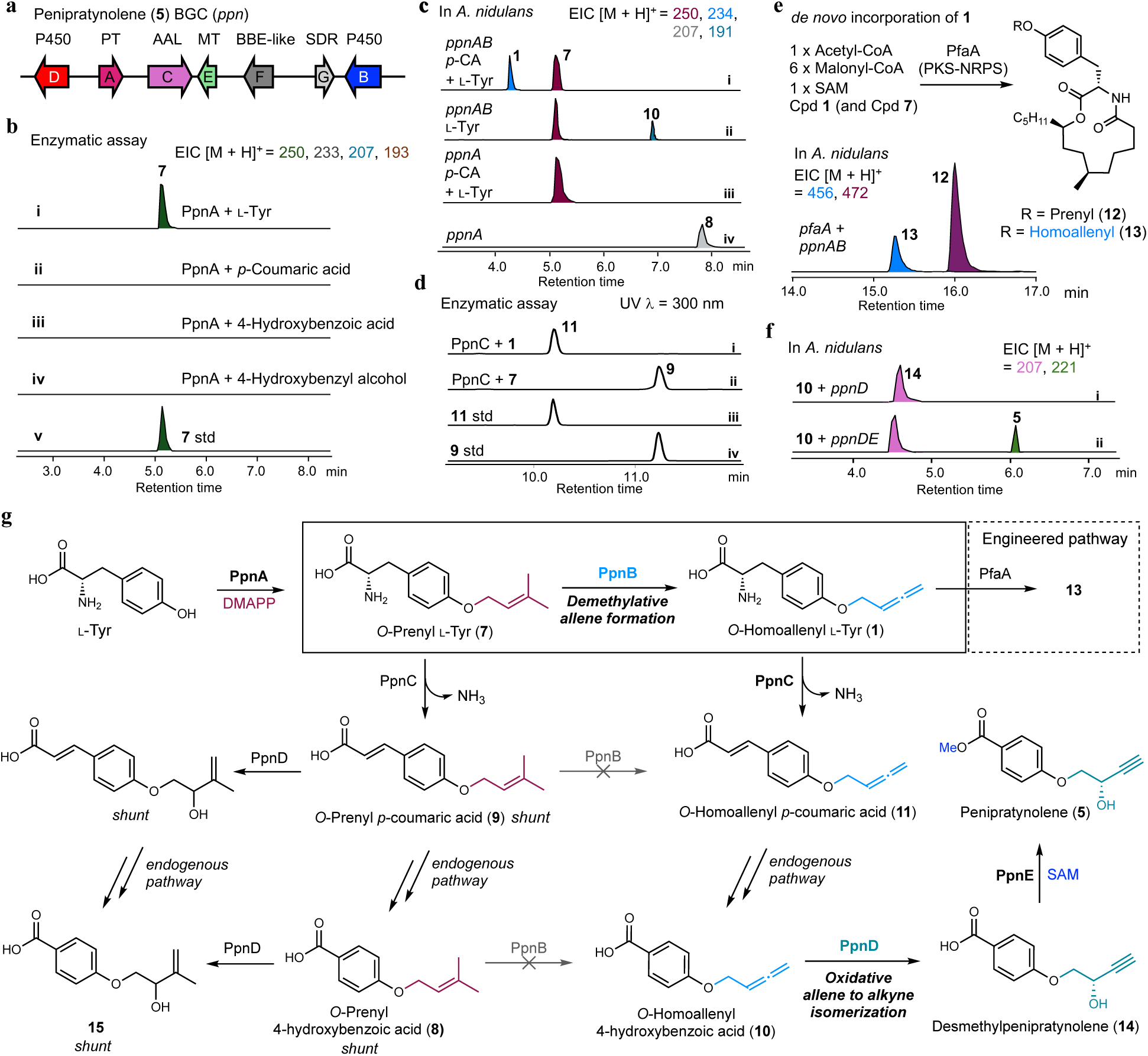
Biosynthesis of penipratynolene (5). **a**, Identification of the *ppn* BGC from *P. polonicum*. **b**, Enzymatic reaction of PpnA with potential substrates. Selected-ion chromatography traces presented on the same scale are shown and the colors of the traces show the MS of corresponding prenylated compounds (i-v). **c**, LC/MS analysis of metabolites produced by *A. nidulans* expressing *ppnA* and *ppnAB* (i-iv). Compound **1** was detected from the transformant expressing *ppnAB* only when _L_-Tyr and *p*-coumaric acid were supplemented to the production media (trace i). **d**, LC/MS analysis (λ = 300 nm) of enzymatic reaction of PpnC with **1** (trace i) or **7** (trace ii) as the substrate. Authentic standards of **9** and **11** are synthetically prepared. **e**, Engineered biosynthesis of the allene-substituted *N*-demethylmelearoride A (**12**) in *A. nidulans*. Previous study showed that PKS-NRPS PfaA can construct *N*- demethylmelearoride A (**12**) using **7** as an amino acid building block^53^. We show that PfaA can use **1** as the amino acid building block to biosynthesize **13**. LC/MS analysis of metabolites produced by the heterologous host *A. nidulans* expressing *ppnAB* coexpressed with *pfaA* is shown. Selected ion chromatography traces presented on the same scale are shown. **f**, LC/MS analysis of biotransformation from **10** to **14** (trace i) by PpnD and from **10** to **5** by PpnD and PpnE (trace ii). Selected-ion chromatography traces are shown and the colors of the traces show the MS of corresponding compounds. **g**, Biosynthetic pathway of L-Tyr to **5**.

Heterologous expression of *ppnA* in *Aspergillus nidulans* ΔEMΔST led to the formation of *O*-prenyl-4-hydroxybenzoic acid (**8**) (**Fig. 3c** and **Supplementary Figs. 5-6**), which was structurally confirmed by NMR (**Supplementary Table 9 and Supplementary Figs. 47-48**). Recombinant PpnA purified from *Escherichia coli* BL21(DE3) (**Supplementary Figs. 7-8**), however, could not catalyze formation of **8** in the presence of 4-hydroxybenzoic acid (4-HBA), DMAPP and Mg^2+^ (**Fig. 3b**). This led us to search for the *bona fide* substrate of PpnA, which may span the entire catabolic pathway of L-Tyr, including L-Tyr, *p*-coumaric acid (*p*-CA), 4-hydroxybenzyl alcohol and 4-hydroxybenzylaldehyde. Biochemical assay showed that PpnA can only prenylate L-Tyr to *O*-prenyl-L-Tyr (**7**) (**Fig. 3b**). This is consistent with feeding studies in which (^13^C_9_,^15^N)-L-Tyr, but not 4-HBA-*d_4_*, can be incorporated into **8** (**Extended Data Figs. 2, 3 and Supplementary Fig. 14**). PpnA displays weak sequence identity (∼30%) to L-Tyr *O*-prenyltransferase SirD^50,51^. Recombinant PpnA catalyzed prenylation of various L-Tyr analogs and D-Tyr (**Supplementary Fig. 9**), and could use either GPP or FPP as the prenyl donor (**Supplementary Fig. 10)**.

These results indicate that **7** is converted to **8** through the L-Tyr catabolic pathway in *A. nidulan*s. This was confirmed by directly feeding **7** to *A. nidulans,* which generated *O*-prenyl-*p*-coumaric acid (*p*-CA) (**9**) and **8** (**Supplementary Fig. 11**). To accumulate **7** in *A. nidulans* for biosynthetic studies, two approaches were used. First, media supplementation of L-Tyr and excess amount of *p*-CA, a known negative feedback regulator of tyrosine ammonia lyase (TAL)^52^, led to detectable levels of **7** from *A. nidulans* expressing *ppnA* (**Fig. 3c**). In parallel, gene-knockout of two putative TALs in *A. nidulans* host led to accumulation of **7** without feeding of L-Tyr and *p*-CA (**Supplementary Fig. 12**).

### PpnB is an Allene-Forming Cytochrome P450 Enzyme

With enzymatic formation of *O*-prenyl-L-Tyr (**7**) established, the next biosynthetic step is hypothesized to be the P450-catalyzed C(*sp*^2^)-demethylation of **7** to *O-*homoallenyl*-*L*-*Tyr (**1**) via a *gem*-diol or an aldehyde intermediate as shown in Figure 1d. Coexpression of *ppnB* with *ppnA* in *A. nidulans* supplemented with both L-Tyr and *p*-CA led to the appearance of a compound with MW of 233, the same as that of **1** (**Fig. 3c**). NMR spectra of the purified compound confirmed the product to be **1**, including characteristic chemical shift of the *sp*-hybridized allenic carbon (209 ppm) in ^13^C NMR (**Supplementary Table 4 and Supplementary Figs. 28-32**). Repeating the coexpression in *A. nidulans* without supplementation of *p*-CA resulted in appearance of a new compound with MW of 190, which was isolated and confirmed to be *O*-homoallenyl-4-hydroxybenzoic acid (**10)** (**Supplementary Table 11 and Supplementary Figs. 6, and 51-52**). To further pinpoint the timing of PpnB, **7** and other downstream catabolic products such as *O*-prenyl-*p*-coumaric acid (**9**), *O*-prenyl-4-hydroxybenzoic acid (**8**) were fed to *A. nidulans* expressing only *ppnB*. While no conversion was observed with **8** and **9**, **7** was clearly converted into **1** (**Extended Data Fig. 4**), supporting that PpnB is solely responsible for catalyzing the unprecedented C(*sp*^2^)-demethylative allene formation from **7** to **1**.

The formation of **10** upon coexpression of *ppnA* and *ppnB* suggests that endogenous TAL or AAL can catalyze the deamination of **1** to *O*-homoallenyl *p*-coumaric acid (**11**), which can then be further modified into **10** (**Supplementary Fig. 6, 13, and 14**). The role of the coclustered putative AAL PpnC was therefore examined. Recombinant PpnC does not catalyze the deamination of L-Phe and L-Tyr, but can catalyze the conversion **7** to **9**, as well as **1** to **11** (**Fig. 3d** and **Supplementary Fig. 15 and 16**). Hence, while it is not essential, PpnC is a dedicated amino acid ammonia lyase for *O*-alkylated L-Tyr in the biosynthetic pathway of **5**.

Our results demonstrate PpnB together with PpnA can transform L*-*Tyr to **1** (**Fig. 3g**). To demonstrate *de novo* synthesis and incorporation of **1** into natural products, we targeted *N*-demethylmelearoride A (**12**), which is a biosynthetic intermediate of the sterol-C_4_-methyl oxidase inhibitor PF1163A^53^ (**Supplementary Fig. 17**). The polyketide-nonribosomal peptide synthetase (PKS-NRPS) PfaA activates and condenses **7** with the polyketide precursor, and catalyzes macrolactonization to give **12** (**Fig. 3e**). When *ppnA* and *ppnB* were coexpressed with *pfaA* in *A. nidulans*, the transformant produced the expected analog homoallenyl-*N-*demethylmeleaoride (**13**) together with **12** (**Fig. 3e**). The presence of the allene function group in **13** was confirmed by NMR (**Supplementary Tables 13** and **Supplementary Figs. 55-59**).

### PpnD Catalyzes Oxidative Isomerization of Allene to Alkyne

Biosynthesis of the allene-containing **10** by the *ppn* enzymes supported the biosynthetic hypothesis that the homoallenyl moiety is the precursor to the α-hydroxy alkyne moiety in penipratynolene (**5**). The second P450 enzyme encoded in the pathway, PpnD, was assayed for the oxidative isomerization activity. When **10** was supplemented to *A. nidulans* expressing *ppnD*, a new compound with MW matching that of desmethylpenipratynolene (**14**) was readily formed. Isolation and structural characterization confirmed the presence of the α-hydroxy alkyne (**Fig. 3f, Supplementary Table 14 and Supplementary Figs. 60-61**). The absolute stereochemistry of **14** was assigned based on matching optical rotation value to that reported for **5**^49^, thereby supporting that PpnD can catalyze the oxidative isomerization of the terminal allene **10** into the terminal alkyne **14**. Coexpression of *ppnD* with *ppnA* (but without *ppnB*) in *A. nidulans* resulted in formation of a new product with MW of 222 (**Supplementary Fig. 6**). NMR characterization revealed the compound to be the 3-methylbut-3-en-2-ol ether of 4-hydroxybenzoate (**15**) (**Fig. 3g**, **Supplementary Table 15** and **Supplementary Figs. 62-66**). Direct feeding of **8** to the *A. nidulans* expressing PpnD similarly led to the formation of **15**, suggesting that PpnD catalyzes oxidative modification of benzoate substrates (**Supplementary Fig. 18**).

Based on the activities of PpnD towards **10** and **8**, two mechanism differing in the initial step can be proposed: (i) HAT from a terminal allenic hydrogen; or (ii) direct attack of allene by Cpd I which is similar to first step of P450-catalyzed epoxidation (**Extended Data Fig. 5)**. Using heme-bound iron as the oxidant with *O*-homoallenyl phenol as a model substrate, DFT calculations show that initiation with HAT is kinetically more favorable over the direct attack by Cpd I (**Extended Data Fig 5b, c**). While generation of an allene radical by natural enzyme is unprecedented, the calculated BDE of the terminal allenic C(*sp*^2^)– H bond in **10** (88.7 kcal/mol) is similar to that of allylic C(*sp*^3^)–H bond in **8** (88.2 kcal/mol)^54,55^, suggesting the HAT step is energetically feasible. To form **14** from **10** via the HAT pathway, a high valent iron-oxo species Cpd I in PpnD abstracts the terminal allenic hydrogens in **10** to generate the allene radical. Delocalization of the radical leads to formation of the more stable propargyl radical, which can undergo stereoselective OH rebound from Cpd II to complete the isomerization (**Extended Data Fig. 5a, b**). Analogously if **8** is the substrate, hydrogen abstraction from the terminal methyl group followed by delayed rebound can give the *exo* olefin **15** (**Supplementary Fig. 18**). PpnD is therefore the first example of an oxidative “allene-alkyne isomerase” that modifies allene to the alkyne.

To complete the biosynthetic pathway, coexpressing the MT-encoding gene *ppnE* with *ppnD* in *A. nidulans* with feeding of **10** resulted in formation of the target compound **5** (**Fig. 3f, 3g**). While the functions of PpnF and PpnG are not assigned in the pathway from L-Tyr to **5** (**Fig. 3g**), these two enzymes might be involved in the conversion of **11** to **10**, which can also be catalyzed by endogenous catabolic enzymes.

### Genome-guided Discovery of the Internal Alkyne forming P450 Enzyme

Identification and activity reconstitution of PpnB complete one half of our initial mechanistic proposal, in which a prenyl group can undergo oxidative demethylation to give a terminal allene (**Fig. 1b, 1d and Fig. 2**). On the other hand, if the second HAT step occurs at the vinyl position instead of the allyl position in the vinyl radical intermediate, an internal alkyne could be formed (**Fig. 1b, 1d**). To search for a homologous P450 enzyme with such activity, the biosynthesis of sinuxylamide B (**2**) isolated from *Xylaria* spp. was explored^26^. Biosynthetically, **2** is derived from *N-*hexanoylation of *O*-but-2-ynyl-L-Tyr (**6**) (**Fig. 1d)**. The BGC of **2** therefore should encode a minimum of three enzymes: homologs of PpnA, PpnB and a *N-*acyltransferase. The top 100 homologs of PpnB were compiled from BLASTP analysis and subjected to phylogenetic analysis. Among the three clades (I, II, and III) emerged from the analysis (**Fig. 4a**), PpnB belongs in Clade I, which also includes enzyme AarB and Xte1B encoded in homologous BGCs from *Apiospora arundinis* and *Xylaria telfairii*, respectively (**Fig. 4a**). While Clade III is distantly related to Clade I, members of Clade II are evolutionarily closely related to those in Clade I. Genome context analysis showed that nearly all PpnB homologs in Clade II are encoded in BGCs that also encode a PpnA homolog and a putative acyltransferase (**Fig. 4a**). These include a second BGC from *X. telfairii* (*xte2*) in which Xte2B is the P450 homolog with 65% and 70% amino acid sequence identity to PpnB and Xte1B, respectively; as well as the *nse* BGC from *Nemania serpens* of which NseB is 89% identical to Xte2B.

**Figure 4.**
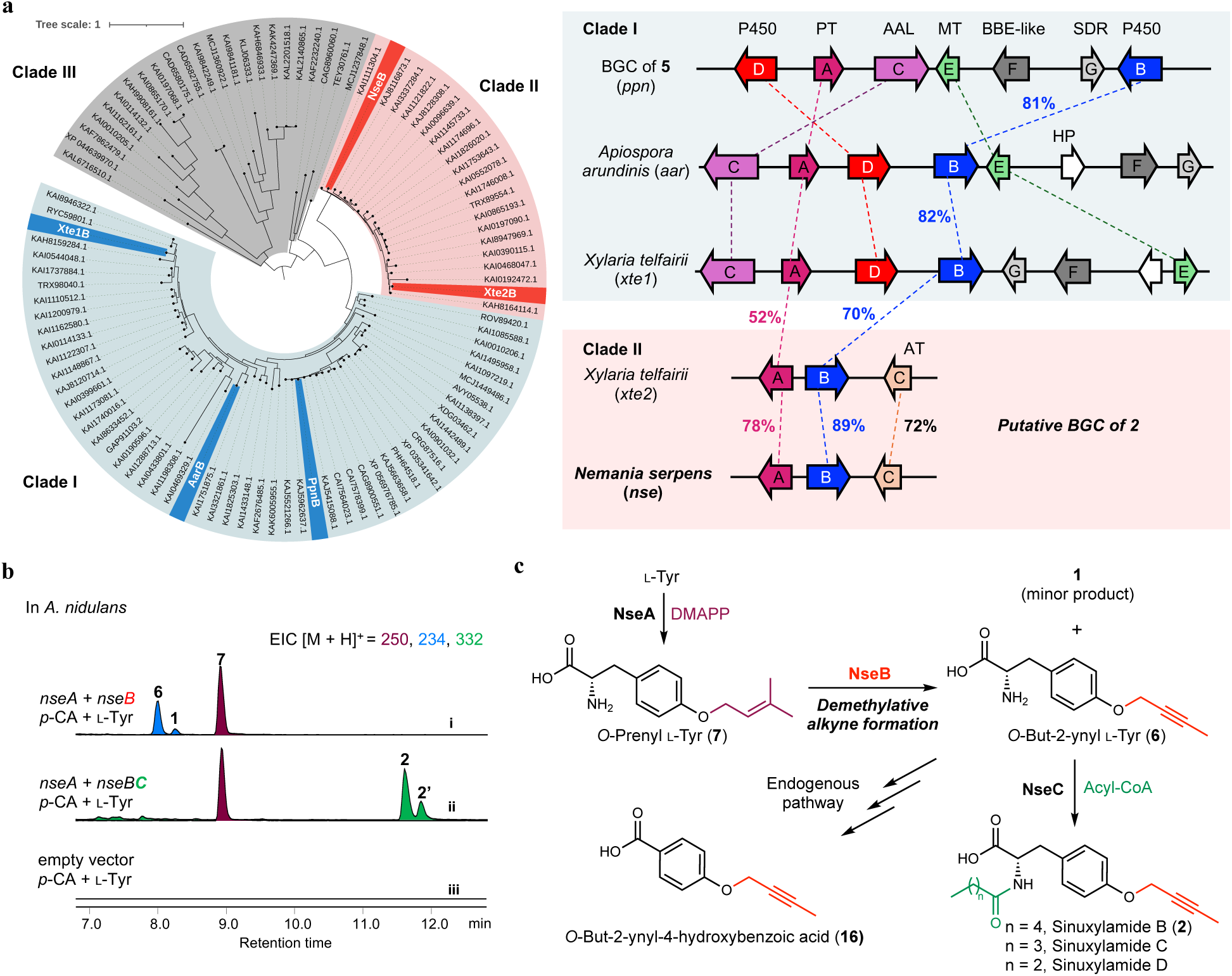
Characterizing the biosynthetic pathway of sinuxylamide B (2). **a**, Phylogenetic analysis of PpnB homologs. The phylogenetic tree is based on 100 sequences obtained by BLASTP using PpnB as the query sequence, and constructed by the maximum likelihood method. The tree is classified into three clades (I, II, and III). Clade I and Clade II are evolutionally closely related as by sharing a late ancestral protein. Two BGCs homologous to *ppn* BGC that contain a PpnB homolog in Clade I are shown: *aar* BGC from *Apiospora arundinis* and *xte1* BGC from *Xylaria telfairii*. The enzymes in Clade II reside in BGCs with a homolog of PpnA and a putative *N-*acyltransferase, which are mainly from *Xylaria* spp and *Nemania* spp (e.g., *nse* BGC from *Nemania serpens*). The numeric values represent amino acid sequence identities (%) between two corresponding enzymes. **b**, LC/MS analysis of metabolites produced by heterologous host *A. nidulans* expressing *nseAB* (trace i) and *nseABC* (trace ii). L-Tyr and *p*-coumaric acid were supplemented to the production media. **2’** is proposed to be the putative *N*-hexanoylated **1**. **c**, Biosynthetic pathway of **2** from L-Tyr.

When the *nseABC* cassette was heterologously expressed in *A. nidulans*, a major product with MW of 331, which matches that of sinuxylamide B (**2**), was detected and isolated (**Fig. 4b**). Both ^1^H and ^13^C NMR spectra, as well as optical rotation, matched with the published data of **2**^26^ (**Supplementary Table 5** and **Supplementary Figs. 33-34**). Two minor products with MWs of 317, and 303 were also detected, which match to those of sinuxylamide C with a *N-*pentanoyl chain and sinuxylamide D with a *N-*butanoyl chain, respectively (**Supplementary Fig. 19**). Coexpression of *nseA* with *nseB* in *A. nidulans* led to formation of three products: the internal alkyne *O*-but-2-ynyl-L-Tyr (**6**) as a major product, *O-*homoallenyl*-*L*-*Tyr (**1**) as a minor product, and *O*-but-2-ynyl-4-hydroxybenzoic acid (**16**) as a catabolite of **6 (Fig. 4b and Supplementary Fig. 19)**. The identity of **16** was verified through NMR analysis (**Supplementary Table 16 and Supplementary Figs. 67-68**). Direct feeding of *O*-prenyl-L-Tyr (**7**) to *A. nidulans* expressing *nseB* also resulted in the formation of **6** and **1** in the same ratio (4:1), while feeding *O*-prenyl-4-hydroxybenzoic acid (**8**) did not lead to formation of **16** (**Extended Data Fig. 4**). These results support that NseB is solely responsible for catalyzing the C(*sp*^2^)-demethylative alkyne formation from **7** to **6**.

In contrast to PpnB which exclusively produces the allene **1**, NseB can generate both **1** and **6** from **7**. This suggests a more flexible active site environment with respect to the second HAT step, in which Cpd II or Fe(III)-O• species can access both the vinyl and the allylic hydrogens (**Fig. 1b and Fig. 2**). These results also confirmed the role of NseC in catalyzing the *N-*acyltransfer step in biosynthesis of **2** (**Fig. 4c**). A putative *N*-hexanoylated **1** (labeled as **2’**) was also detected from the *A. nidulans* expressing *nseABC* (**Fig. 4b**), suggesting that NseC could acylate **1** in addition to **6**. Acylation of the α-amino group in **6** prevents deamination and subsequent catabolic steps that form **16**.

### Mechanistic Investigation of PpnB and NseB using Deuterated Substrates

Based on our proposed mechanism for the transformation of the C5 prenyl group in **7** into the C4 homoallenyl in **1** or the C4 but-2-ynyl group in **6** (**Figs. 1d and 2**), one of the terminal C(*sp^2^*)-methyl groups in the prenyl precursor must be oxidatively excised during the transformation. To determine which of the two methyl groups (*cis* C_4_ vs. *trans* C_5_) is removed and to gain more mechanistic insight, we prepared a series of deuterated **7**, including 4,4,4-[*d*_3_]-**7**, 5,5,5-[*d*_3_]-**7**, 2-[*d*_1_]-**7**, and 1,1-[*d*_2_]-**7**. These were used as substrates for enzymatic reactions with microsomal fractions containing PpnB or NseB prepared from *A. nidulans* transformants. The enzymatic products were analyzed by LC/QTOF to determine the regioselectivity of methyl excision and hydrogen abstraction. When 4,4,4-[*d*_3_]-**7** was used as the substrate for PpnB reaction, no mass increase in the product **1** was observed. In contrast, 5,5,5-[*d*_3_]-**7** led to +2 Da increase in mass of **1** (**Fig. 5a**). The same pattern of mass change for **1** was also observed in the biotransformation of the methyl ester forms of 4,4,4-[*d*_3_]-**7** and 5,5,5-[*d*_3_]-**7** by *A. nidulans* expressing *ppnB* (**Extended Data Fig. 6**). Consistent observations were seen with NseB: no mass change in **6** was observed with 4,4,4-[*d*_3_]-**7** as substrate, while +3 Da mass increase in **6** was detected with 5,5,5-[*d*_3_]-**7** as substrate (**Extended Data Fig. 7**). These results unambiguously establish that the *cis* C_4_ methyl group is selectively removed by both PpnB and NseB.

**Figure 5.**
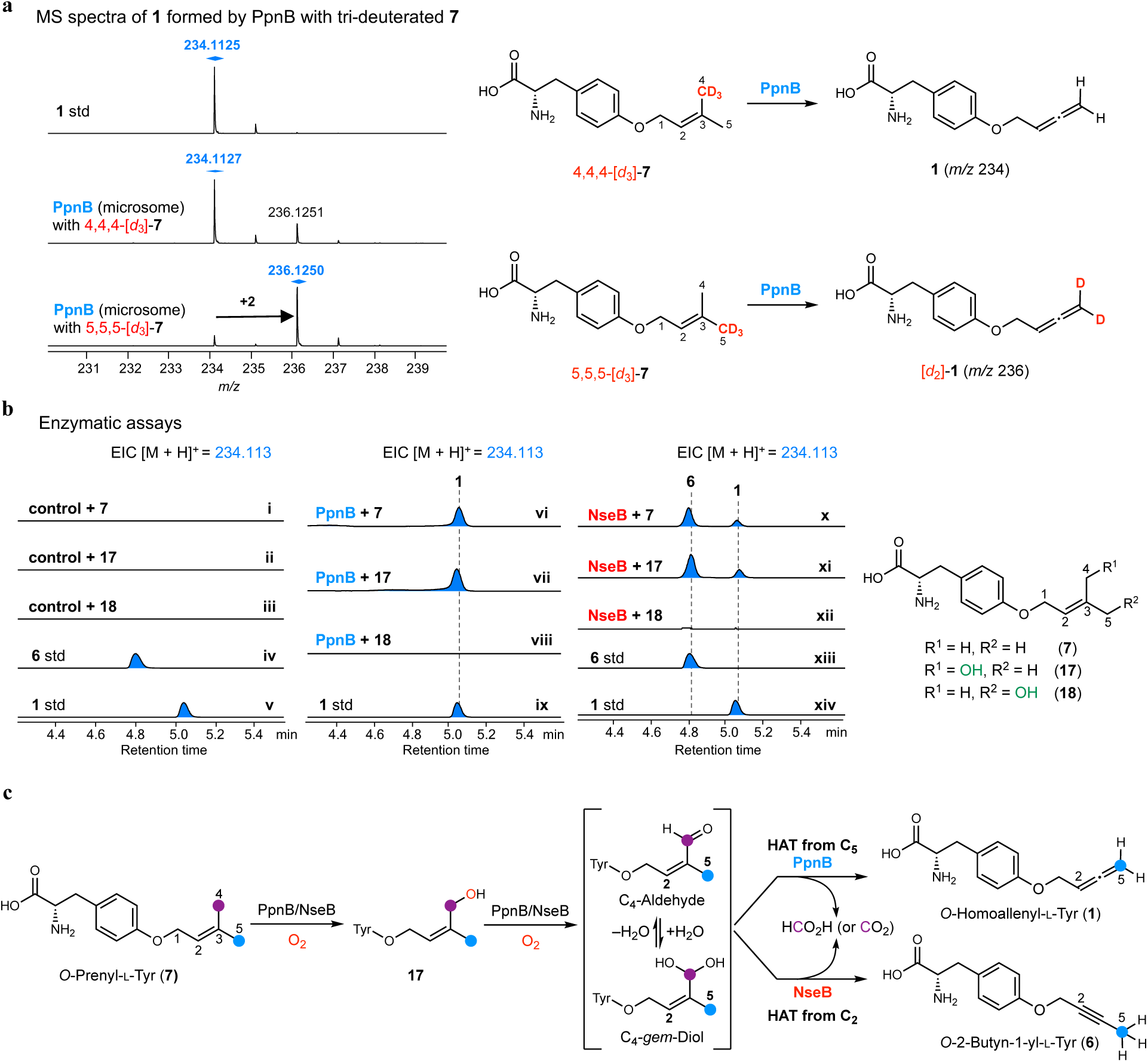
Mechanistic study and the proposed mechanism of the PpnB and NseB-catalyzed allenylation and alkynation. **a**, LC/QTOF MS analysis of PpnB-catalyzed reaction with deuterated **7** such as 4,4,4-[*d*_3_]-**7** and 5,5,5-[*d*_3_]-**7**. Microsomal fraction containing PpnB was prepared from *A. nidulans* expressing PpnB. Enzymatic reactions were performed in 100 µL of 50 mM Tris-HCl buffer containing PpnB (microsome), 100 µM substrate, and 2 mM NADPH at 28°C for 16 h. Clear +2 Da mass shift of **1** was only observed when 5,5,5-[*d*_3_]-**7** was used for the enzymatic assays. **b**, LC/QTOF analysis of PpnB and NseB-catalyzed reactions with the putative substrates such as **7** (trace vi for PpnB and trace x for NseB), **17** (trace vii for PpnB and trace xi for NseB), **18** (trace viii for PpnB and trace xii for NseB). The same reaction conditions described in **a** were used. Control experiments were performed with microsome prepared from *A. nidulans* harboring empty vectors. **c**, Proposed mechanism of the oxidative conversion of C5 prenyl group to C4 allene and C4 alkyne catalyzed by PpnB and NseB, respectively.

The detected mass shifts in **1** (+2 Da) and **6** (+ 3Da) with 5,5,5-[*d*_3_]-**7** as substrate are consistent with the proposed product-determining second HAT step by PpnB and NseB, respectively. In the PpnB reaction, the *trans* C_5_ methyl group becomes the terminal allene carbon in **1** and loses one hydrogen upon HAT, resulting in the +2 Da shift. In the NseB reaction, all hydrogens on the *trans* C_5_ methyl group are retained in **6** to give the +3 Da shift. To confirm the second HAT step for NseB takes places at vinylogous C_2_, 2-[*d*_1_]-**7** was used in the enzyme assay. As expected, no mass increase in **6** was detected in the presence of NseB, indicating the loss of the deuterium label; while a +1 Da increase in the mass of **1** was detected in the presence of PpnB (**Supplementary Fig. 20**). Finally, both PpnB and NseB reactions gave a +2 Da increase in the mass of products when 1,1-[*d*_2_]-**7** was used as the substrate (**Supplementary Fig. 21**). Collectively, these results are consistent with our proposed mechanism shown in Fig. 2.

The mechanistic proposal in Fig. 2 involved iterative hydroxylation of the to-be-excised methyl group to the *gem*-diol or the aldehyde intermediate, which can then undergo fragmentation reaction in a Cpd 0- or Cpd I-dependent fashion. To test if hydroxylated **7** is indeed an intermediate of PpnB and NseB, **17** and **18**, which are C_4_-hydoxyl-**7** and C_5_-hydroxyl-**7**, respectively, were chemically synthesized and used as substrates for the enzymatic reactions. Consistent with the deuterium labeling results, only **17** was converted into **1** by PpnB, while **18** was not modified (**Fig. 5b**). The same selectivity towards **17** but not **18** was also observed in NseB-catalyzed formation of **6** (major) and **1** (minor) (**Fig. 5b**). Searching for putative intermediates during the enzymatic reaction from **7** to **1** led to the identification of **17** based on the retention time of the synthetic standard (**Supplementary Fig. 22**). Attempts to synthesize C_4_-aldehyde-**7** for enzymatic rection was not successful, as chemical oxidation of the methyl ester of **17** led to the formation of C_5_-aldehyde-**7** methyl ester due to the rapid *Z* to *E* isomerization as previously documented^26^. While we cannot yet conclusively pinpoint whether the C_3_–C_4_ fragmentation occurs in either C_4_-aldehyde (or C_4_-*gem*-diol) or C_4_-carboxylate form of **7**, our results strongly support that the reaction is initiated by the regioselective hydroxylation at C_4_ (**Fig. 5c**).

## Discussion

Allene and alkyne are among the most useful functional groups in complex organic compounds for structural diversification, semisynthesis and bioconjugation^1–12^. However, in clear contrast to the development of synthetic methods for allene and alkyne formation, the biosynthesis of these functionalities is underexplored (alkyne)^28^ or is nearly nonexistent (allene)^34,56^. In this study, we discovered three fungal P450 enzymes (PpnB, PpnD, and NseB) from the biosynthesis of penipratynolene (**5**) and sinuxylamide B (**2**) that can form allene or alkyne functionality via unprecedented oxidative mechanisms. In biosynthesis of **5**, PpnB catalyzes the demethylative terminal allene formation of *O*-prenyl-L-Tyr (**7**) to *O*-homoallenyl-L-Tyr (**1**); whereas NseB, a homolog of PpnB, catalyzes the demethylative internal alkyne formation of **7** to *O*-but-2-ynyl L-Tyr (**6**). To the best of our knowledge, no demethylase had been shown to remove a methyl group directly from a *sp*^2^-hybridized carbon prior to this work, making PpnB and NseB the first examples of demethylative “allenylase” and “alkynase”. Moreover, such demethylative allene and alkyne-forming reactions have not been documented in synthetic chemistry. This discovery highlights the use of PpnB and NseB by Nature to achieve precise C–C bond scission, representing a novel strategy for functional group installation.

PpnB- and NseB-catalyzed oxidative demethylation likely proceed through a mechanistically analogous pathway as that proposed for CYP51^42–44^. While an analogous heterolytic C_3_–C_4_ cleavage pathway via a BV intermediate could also be considered for the Cpd 0-dependent mechanism (**Supplementary Fig. 23**), one proposed mechanism involving a homolytic C_3_–C_4_ cleavage (**Fig. 2**) could begin with hydrogen abstraction from the C_4_-*gem*-diol compound (**Fig. 5c**) by Cpd I species, or attack of the C_4_-aldehyde by Fe(III)-peroxo species (**Fig. 5c**), results in C_3_–C_4_ homolytic fragmentation to generate a C_3_-vinyl radical intermediate. Regioselectivity of the second HAT step dictates product outcome: PpnB abstracts a hydrogen from the terminal C_5_-methyl group to yield the terminal allene **1**, whereas NseB abstracts C_2_-vinylic hydrogen to form the internal alkyne **6**. The bifurcation of the vinyl radical as a shared intermediate to allene and alkyne products is consistent with our DFT calculations (**Fig. 2**), and is reflected in formation of both **6** (major) and **1** (minor) in the NseB reaction (**Figs. 4b and 5b**). The hydroxylation and desaturation reactions catalyzed by PpnB and NseB indicate versatile active sites in PpnB and NseB that can switch between distinct modes of heme-based chemistry depending on substrate. While purification of active membrane-anchored eukaryotic P450s remains a persistent challenge, further biochemical and biophysical characterization of PpnB and NseB will be essential to fully elucidate these unusual and mechanistically intriguing transformations.

Because the position of second HAT dictates product outcome (allene vs alkyne), it is reasonable to propose that the active sites of each enzyme control the positions of the heme iron relative to the prenyl group in **7**: in PpnB, the heme iron is positioned between C_4_ and C_5_ of **7**, whereas in NseB, it is positioned between C_4_ and C_2_. A comparison of the AlphaFold3-predicted structures of PpnB and NseB shows that residues lining both substrate-binding and heme-binding pockets are nearly identical. The only notable differences are two residues (V127/I126 and L380/F379 in PpnB/NseB) found in the heme-binding pocket. Introducing the corresponding double mutation in NseB (I126V/F379L) increases the ratio of allene **1** relative to alkyne **6** by ∼2-fold (**Supplementary Fig. 25**), whereas the reciprocal double mutant in PpnB (V127I/L380F) does not result in **6** formation. This suggests that product selectivity is primarily dictated by distal (second- and third-shell) residues that result in subtle differences in active site environment. Indeed, a growing body of work showed that such residues can tune catalytic efficiency, reaction selectivity, and substrate scope by modulating conformational dynamics, reshaping the active-site cavity, and altering pocket volume^57–60^. For example, a directed evolution campaign converted a P450 into a ketone synthase for selective oxidation of internal alkenes by targeting distal positions^61^; while distal beneficial mutations in P450 BM3 improved activity and broadened substrate scope^58,62^.

The discovery of allene-forming enzyme PpnB suggests a plausible biosynthetic route to xenicin-type norditerpenoids bearing terminal allenes, such as ginamallene, isolated from octocoral *Acalycigorgia* spp^23^ (**Fig. 6**). Our findings suggest a pathway in which the terminal allene in ginamallene can be formed from a diterpene precursor via oxidative one-carbon deletion. Recent work from Burkhardt et al. identified a diterpene synthase XsTC-1 from *Xenia* spp. that catalyzes the cyclization of geranylgeranyl pyrophosphate (GGPP) to xeniaphyllene^63^, a proposed precursor to a large group of xenicane diterpenoids, including 9-deacetoxy-14,15-deepoxyxeniculin^64^. Because ginamallene and 9-deacetoxy-14, 15-deepoxyxeniculin have been co-isolated from octocoral *Acalycigorgia* spp^23^, one can proposed that 9-deacetoxy-14, 15-deepoxyxeniculin can undergo oxidative demethylation to form ginamallene (**Fig. 6**). Although no *Acalycigorgia* genomes are currently available (as of January 2026), a targeted search for P450s that are colocalized^63,65,66^ or co-expressed with a homolog of XsTC-1 could be one strategy to identify the new allene-forming enzymes once genomic resources emerge.

**Figure 6.**
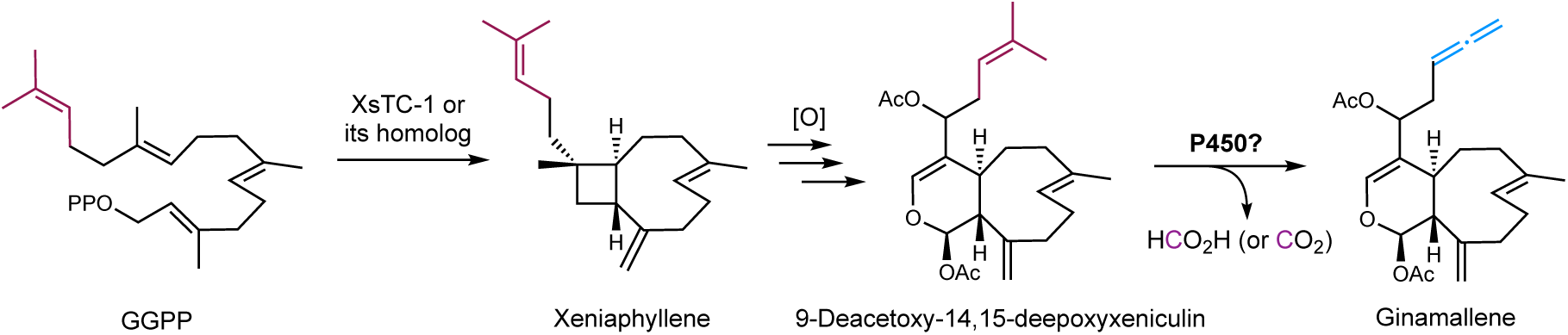
Proposed biosynthetic pathway of ginamallene, featuring oxidative editing of C20-diterpenoid scaffold into C19 nor-diterpenoid with the terminal allene formation. Because ginamallene and 9-deacetoxy-14,15-deepoxyxxeniculin were co-isolated from *Acalycigorgia inermis*, we propose that 9-deacetoxy-14,15-deepoxyxxeniculin is a biosynthetic intermediate en route to ginamallene. XsTC-1 in *Xenia* spp. produces xeniaphyllene (a putative xenicane precursor), which could be converted to 9-deacetoxy-14,15-deepoxyxxeniculin via oxidative cyclobutene cleavage and tailoring oxidations, and then to ginamallene via a P450-catalyzed oxidative demethylation—potentially by an allene-forming P450 unique to *Acalycigorgia* spp.

In summary, we uncovered three unique P450 enzymes (PpnB, NseB, and PpnD) that generate terminal allene, internal alkyne, and terminal alkyne functionalities originated from a common C5 isoprenoid unit. These enzymes expand the catalytic repertoire of P450s and provide a foundation for developing biocatalysts to install allene and alkyne groups, with broad applications in synthetic and medicinal chemistry.

## Online Methods

### Metabolite analysis and compound isolation from *A. nidulans* transformants

For small scale metabolite analysis in *A. nidulans*, transformants containing the desired plasmids were selected from CD sorbitol agar (2% glucose as carbon source) appropriately supplemented with riboflavin, uracil, and/or pyridoxine. CD-ST agar was inoculated with spores and incubated at 28 °C for four to five days. The agar was then collected and extracted with acetone for 30 min with sonication. After centrifugation, the supernatant (200 µL) was concentrated and re-suspended in methanol (100 µL). The LC/MS analyses were performed on an Agilent LC/MSD iQ (AgilentTM InfinityLab Poroshell 120 Aq-C18, 2.7 μm, 100 Å, 2.1 × 100 mm) using positive and negative mode electrospray ionization with a linear gradient of 1-99% CH_3_CN-H_2_O supplemented with 0.1% (v/v) formic acid in 13.25 min followed by 99% CH_3_CN for 3 min with a flow rate of 0.6 mL/min. The data were collected and analyzed by OpenLab CDS 2.4 (Agilent). Some of LC-MS analyses were performed on a Shimadzu 2020 EV LC-MS with a reverse-phase column (Phenomenex Kinetex, C18, 1.7 μm, 100 Å, 2.1 × 100 mm) using positive and negative-mode electrospray ionization with a linear gradient of 5-95% acetonitrile-H_2_O (containing 0.1% formic acid) in 15 min followed by 95% acetonitrile for 3 min with a flow rate of 0.3 ml/min.

For large-scale analysis, *A. nidulans* transformants were inoculated to plates each of which contains 50 mL CD-ST agar, and were placed in a 28 °C incubator for 3–4 days. After 4 days, the solid agar cultures were cut into small pieces and were extracted extensively with acetone. The residual was loaded on a reverse-phase CombiFlash^®^ system and was subjected to flash chromatography with a gradient of water (A) and methanol (B) (100:0 to 0:100, v/v) for initial separation. Metabolites of interest, tracked by analytical HPLC and LC/MS, were purified from the corresponding fractions by reverse-phase semipreparative HPLC with a COSMOSIL column with flow rate of 4 mL/min of solvents A (H_2_O with 0.1% formic acid) and B (MeCN). NMR spectra were obtained with a Bruker AV500 spectrometer with a 5 mm dual cryoprobe at the UCLA Molecular Instrumentation Center. (^1^HNMR 500 MHz, ^13^CNMR 125 MHz). High resolution mass spectra were also recorded on a Agilent 6545 Quadrupole Time of Flight high resolution mass spectrometer (UCLA Molecular Instrumentation Center). The mass and NMR spectra were analyzed by MassHunter 10.0 (Agilent) and MestReNova-9.0.1 (Mestrelab Research), respectively.

For isolation of **2**, an *A. nidulans* transformant expressing *nseA* and *nseBC* was grown on 8 L of solid CD media supplying with 2 mM L-Tyr, and 2 mM *p*-coumaric acid for 4 days at 28 °C and then extracted with acetone. The crude extracts were subjected to fractionation using a CombiFlash flash chromatography system (Teledyne). Fractions determined by LC-MS to contain target compounds were pooled. After solvent evaporation, the residue was further separated by semi-preparative RP HPLC using a Shimadzu UFLC system and a COSMOSIL column (Nacalai Tesque Inc., 5C18-AR-II, 20ID x 250mm, flow rate of 4 mL/min of solvents A (0.1% formic acid in water) and B (acetonitrile), with an isocratic concentration of 50% acetonitrile (MeCN)–50% water, to afford **2**. For isolation of **5**, *penicillium polonicum*^18^ was cultured in 4 L PDB liquid. The culture was then extracted by EtOAc. Similarly, after CombiFlash chromatography, fractions containing the target compound were combined and purified by HPLC with a semi-preparative reverse-phase column (5C18-AR-II, 10ID × 250 mm) with water (A) and acetonitrile (B) with 0.1% formic acid using a gradient of 0–5 minutes 20% B; 5–40 minutes 20–80% B. For isolation of **8**, an *A. nidulans* transformant expressing PpnA was grown on 4 L of solid CD media for 5 days at 28 °C and then extracted with acetone. For isolation of **7**, an *A. nidulans* transformant expressing PpnA was grown on 4 L of solid CD media supplying 2 mM L-Tyr, and 2 mM *p*-coumaric acid for 4 days at 28 °C and then extracted with acetone. Purification was also done similarly by fractionation using a CombiFlash system followed by semi-preparative RP HPLC (5C18-AR-II, 10ID x 250mm, flow rate of 4 mL/min), with an isocratic concentration MeOH–H_2_O (60:40, v/v) to yield **7**. For isolation of **10**, an *A. nidulans* transformant expressing PpnAB was grown on 4 L of solid CD media for 3 days at 28 °C and then extracted with acetone. Fractions containing the target compound from the CombiFlash were combined and purified by HPLC with a semi-preparative reverse-phase column (5C18-AR-II, 10ID × 250 mm) with water (A) and acetonitrile (B) with 0.1% formic acid using an isocratic concentration of 40% B (0–40 minutes). To isolate **13**, an *A. nidulans* transformant coexpressing PpnAB and PfaA was cultivated on 8 L of solid CD medium supplemented with 2 mM L-Tyr and 2 mM *p*-coumaric acid for 4 days at 28 °C, followed by extraction with acetone. After CombiFlash chromatography, the extract was then purified by HPLC using MeOH as the organic phase under a gradient of 80% B from 0–15 min and 80–100% B from 15–25 min. For the isolation of **14**, compound **5** underwent basic hydrolysis, followed by pH adjustment and extraction with EtOAc. The resulting organic extract was subsequently purified by HPLC using a gradient of 20% B from 0–5 min and 20–80% B from 5–20 min. Compound **14** was also obtained by feeding 1 mM compound **10** to *A. nidulans* strains overexpressing PpnD, cultivated on 4 L agar medium for 5 days. For isolation of **15**, an *A. nidulans* transformant expressing PpnAD was grown on 4 L of solid CD media for 4 days at 28 °C and then extracted with acetone. After CombiFlash chromatography, fractions containing the target compound were combined and further purified by HPLC on a semi-preparative reverse-phase column (5C18-AR-II, 10ID × 250 mm) using water (A) and acetonitrile (B), each containing 0.1% formic acid, with a gradient of 20% B from 0–5 min and 20–80% B from 5–40 min. To isolate **16**, an *A. nidulans* transformant expressing PpnA and NseB was grown on 8 L of solid CD medium for 4 days at 28 °C and extracted with acetone. Followed by CombiFlash chromatography, the crude extract was purified by HPLC using MeOH as the organic phase with a gradient of 35% (0–20 min) to 50% (20–26 min).

### Proteins expression and purification

The intron free ORFs encoding PpnA, and PpnC were amplified by PCR using cDNA which was reverse transcribed from mRNA extracted from the corresponding *A. nidulans* transformant as a template, and ligated to linear expression vector pET28a via Gibson Assembly (New England Biolabs, NEB), following the manufacturer’s protocol. The plasmids were then transformed into *E. coli* BL21(DE3) individually and grown overnight in 5 mL of LB medium with 50 µg/mL kanamycin at 37 °C. The overnight cultures were used as seed cultures for 1 L fresh LB media containing 50 µg/mL kanamycin and incubated at 37 °C until the OD_600_ reached 0.8. The cultures were cooled on ice, before addition of 0.1 mM isopropyl-β-D-thiogalactopyranoside (IPTG, GoldBio, USA) to induce protein expression. The expression was performed at 16 °C for 20 h, 220 rpm. *E. coli* cells were harvested by centrifugation at 5200 x *g* for 15 min and resuspended in 30 mL A10 buffer (50 mM sodium phosphate buffer, 150 mM NaCl, 10 mM imidazole, pH 8.0) containing 1 tablet of PierceTM protease inhibitor (Thermo Scientific). The cell suspension was lysed on ice by sonication and the lysate was centrifuged at 17,000 x *g* for 30 min at 4°C to remove the insoluble cellular debris. Recombinant hexa-His-tagged proteins were purified at 4°C from corresponding soluble fractions by affinity chromatography with Ni-NTA agarose resin (GE Healthcare). Briefly, recombinant proteins on the resin was initially washed with wash buffer A1 (50 mM sodium phosphate, 150 mM NaCl, 10 mM imidazole, pH 8.0) until no protein was detected in the eluent using the Bradford reagent. Then, the same procedure was repeated with wash buffer A2 (50 mM sodium phosphate, 150 mM NaCl, 20 mM imidazole, pH 8.0). The target protein was eluted by elution buffer A (50 mM sodium phosphate, 150 mM NaCl, 250 mM imidazole, pH 8.0). The purified proteins were concentrated and exchanged into storage buffer (50 mM sodium phosphate, 200 mM NaCl, 10% glycerol, pH 8.0) with Amicon Ultra-30K concentrator (Merck Millipore). SDS-PAGE was performed to check the protein purity and Bradford Protein Assay (Bio-Rad) was used to calculate protein concentration with bovine serum albumin (BSA, Sigma) as standard. The proteins were aliquoted and stored at −80 °C until used in in vitro assays. The plasmids used for protein purification are listed in **Supplementary Table 2**. See **Supplementary Figs. 7** for SDS–PAGE analysis.

### Enzymatic assays for PpnA and PpnC

To assay the activities of PpnA, reactions were performed in 100 µL of 50 mM sodium phosphate buffer (pH 8.0) containing 1 mM of substrate, 6 mM DMAPP/FPP/GPP, 5 mM MgCl_2_, and 10 µM of PpnA at 28 °C for overnight. The reaction mixture in the absence of protein was prepared as the negative control. Enzyme reactions were quenched by adding 100 µL acetonitrile and centrifuged at 17,000 x *g* for 5 min. LC-MS analyses were performed on a Shimadzu 2020 EV LC-MS with a reverse-phase column (Phenomenex Kinetex, C18, 1.7 μm, 100 Å, 2.1 × 100 mm) using positive and negative-mode electrospray ionization with a linear gradient of 5-95% acetonitrile-H_2_O (containing 0.1% formic acid) in 15 min followed by 95% acetonitrile for 3 min with a flow rate of 0.3 ml/min.

To assay the activities of PpnC, a typical reaction contains 20 µM enzyme, 0.5 mM **1** and **7**, in 100 µL of 50 mM sodium phosphate buffer (pH 8.0). After incubation at 28 °C for overnight, the reaction was then quenched with 100 µL MeCN and centrifuged at 17,000 x *g* for 5 min. The supernatant was subjected to Shimadzu 2020 EV LC-MS analysis with the same conditions as mentioned above.

### Biotransformation assays for PpnB/NseB

To feed compounds to the *A. nidulans* transformants overexpressing *PpnB/NseB*, spores of the transformants were inoculated onto 25 mL of solid CD-ST medium (20 g L⁻¹ soluble starch, 20 g L⁻¹ tryptone, 6 g L⁻¹ nitrate salts, 0.52 g L⁻¹ KCl, 0.52 g L⁻¹ MgSO₄·7H₂O, 1.52 g L⁻¹ KH₂PO₄, and 1 mL L⁻¹ trace elements) and incubated at 28 °C. After three days, the substrate **7**, **7**-Me, **17**-Me, **18**-Me, **8**, and **9**, were added to a final concentration of ∼500 μM, along with supplementation of 2 mM L-Tyr and 2 mM *p*-coumaric acid. The cultures were then incubated for an additional three days, extracted with acetone, and analyzed by LC–MS as mentioned above.

### Microsomal assays for PpnB/NseB

The *A. nidulans* transformants expressing PpnB or NseB, or empty vectors, were cultured in 500 mL Erlenmeyer flasks containing 100 mL CD-ST liquid medium at 28°C, 220 rpm for 4 days. The liquid culture was vacuum filtered through cheesecloth on a Büchner funnel to collect the mycelia. The mycelia were successively washed with 1 L of ultrapure water (Milli-Q water) and 400 mL of Buffer A (50 mM Tris-HCl, 1 mM EDTA, 0.6 M sorbitol, 20% glycerol, pH 7.5). The mycelia can be freeze-dried and/or directly stored at -80°C until used for microsome extraction.

To extract microsomes, 4.0 g of the mycelia was added into a cold-chilled unglazed mortar (pre-chilled by pouring liquid nitrogen). Grinding with a pestle was continued until the mycelia became a white powder, with occasional additions of liquid nitrogen to prevent melting. The resulting powder was suspended in 20 mL of Buffer B (50 mM Tris-HCl, 1 mM EDTA, 0.6 M sorbitol, 20% glycerol, 2.0 mM β-mercaptoethanol, Pierce™ protease inhibitor cocktail, pH 7.5) or Homogenization Buffer II from Microsome isolation kit (Abcam). The suspension was centrifuged at 4,300 × g at 4°C for 20 min to remove cellular debris. The supernatant was transferred into microcentrifuge tubes in 1 mL aliquots and centrifuged at 10,000 × g at 4°C for 15 min. The resultant supernatant was transferred to new microcentrifuge tubes (1 mL per tube) and centrifuged at 21,000 × g at 4°C for at least 2 hours to pellet the microsomal fraction. After removing the supernatant, 800 µL of Buffer C (50 mM Tris-HCl, 50 mM NaCl, 20% glycerol, pH 7.5) was added to the first tube to resuspend the pale brown pellet. This mixture was sequentially transferred to the remaining tubes to resuspend and pool all microsomal fractions.

For the in vitro enzymatic assay, a 100 µL reaction mixture was prepared containing the PpnB or NseB or control microsomal fraction, 100 µM substrate (e.g., compounds **7**, **17**, or **18**), and 2 mM NADPH. The reaction was incubated at 22°C for 16 h and subsequently quenched by the addition of an equal volume (100 µL) of methanol. The quenched mixture was centrifuged at 17,000 × g for 5 min, and the resulting supernatant was subjected to LC–QTOF (quadrupole time-of-flight) analysis with an Agilent 6545 QTOF equipped with a reverse-phase column (Agilent Poroshell, 120 EC-C18; 2.7 μm, 3.0 × 50 mm) using positive-mode electrospray ionization with 1% acetonitrile in H_2_O (containing 0.1% formic acid) for the first 2 min, then a linear gradient of 1–95% for 9 min and finally 95% acetonitrile for 3 min with a flow rate of 0.6 ml min^−1^.

### Biotransformation assays for PpnD or PpnDE

To feed compounds to the heterologously *A. nidulans* strains heterologously overexpressing *PpnD* or *PpnDE*, spores of the transformed strain were inoculated onto 25 ml of solid CD-ST medium and incubated at 28 °C. After three days, substrate **10** was added to a final concentration of ∼500 μM. The cultures were then incubated for an additional three days, extracted with methanol, and analyzed by LC–MS as mentioned above.

### Phylogenetic tree analysis of allene- and alkyne-forming P450s

Homologs of PpnB were identified by BLASTP searches against the NCBI non-redundant (nr) protein database using PpnB as the query sequence. Retrieved homologous P450 sequences were aligned by multiple sequence alignment (MSA) using ClustalW with default parameters. A phylogenetic tree was inferred using the maximum-likelihood (ML) method implemented in the IQ-TREE web server (http://iqtree.cibiv.univie.ac.at/) with default settings. Branch support was assessed using 1,000 ultrafast bootstrap replicates. The resulting tree was visualized and annotated using iTOL (https://itol.embl.de/).

## Computational Methods

All the calculations were carried out using Gaussian 16 program^65^. The geometries were optimized using B3LYP-D3(BJ)^68–72^ functional with def2-SVP^71^ basis set with IEFPCM solvent model^72^ to describe the diethyl ether environment. Transition structures have also been verified by intrinsic reaction coordinate (IRC)^73^ calculations. No geometrical restraints are applied for optimizations. Single point energies were calculated using B3LYP-D3(BJ) functional, def2-TZVP basis set and SMD solvent model^74^. Quasiharmonic^75^ and concentration corrections to enthalpy and entropy were made using Paton’s GoodVibes software^76^. For the hydronium ion formed in the reaction, proton solvation energy that reported by Kelly et al. was used^79^, while the thermodynamic correction of a free proton in gas phase was calculated using Fermi-Dirac distribution^80^.

**Supplementary Information** is available in the online version of the paper.

## Acknowledgement

This work was supported by grant from NIH (R35GM118056) to Y.T., and NSF (CHE-2153972) to K.N.H. This work used computational resources provided by Expanse at SDSC through allocation CHE-040014 from the Advanced Cyberinfrastructure Coordination Ecosystem: Services & Support (ACCESS) program.

## Author contributions

M.L., M.O., K.N.H., and Y.T. developed the hypothesis and conceived the idea for the study. M.L., M.O., K.N.H., and Y.T. designed the experiments. M.L. performed all *in vivo*, *in vitro* experiments as well as compound synthesis, isolation and characterization. M.L. and M.O. performed bioinformatic analysis and identified the biosynthetic gene clusters for penipratynolene and sinuxylamides. M.O. and W.H. synthesized the compounds. Q.Z. performed all computational studies. All authors analyzed and discussed the results. M.L., M.O., K.N.H., and Y.T. prepared the main text of the manuscript. All authors participated in the preparation of methods section and supplementary information.

## Data availability

The data that support the findings of this study are available within the paper and its Supplementary Information, or are available from the corresponding authors upon request.

---

**Extended Data Fig. 1.**
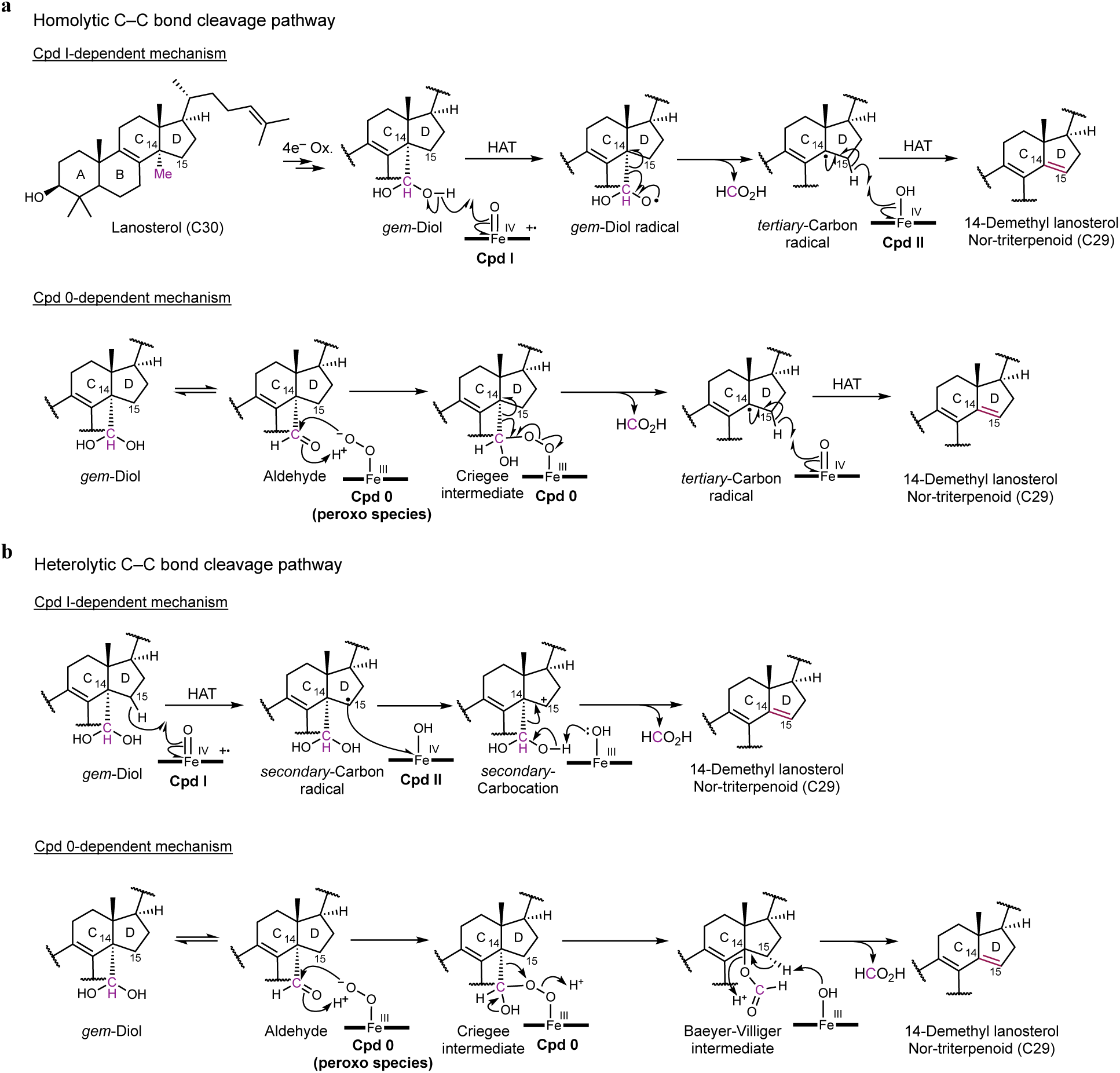
Proposed mechanisms of CYP51-catalyzed C(*sp^3^*)-demethylation of lanosterol. **a.** Proposed homolytic C–C bond cleavage pathway with proposed Cpd I-dependent mechanism starting from the *gem*-diol and proposed Cpd 0-dependent mechanism starting from the aldehyde^43,44^. **b.** Proposed heterolytic C–C bond cleavage pathway with proposed Cpd I-dependent mechanism (carbocation-mediated mechanism) starting from the *gem*-diol and proposed Cpd 0-dependent mechanism via a Baeyer-Villiger (BV) intermediate starting from the aldehyde^42,45^.

**Extended Data Fig. 2.**
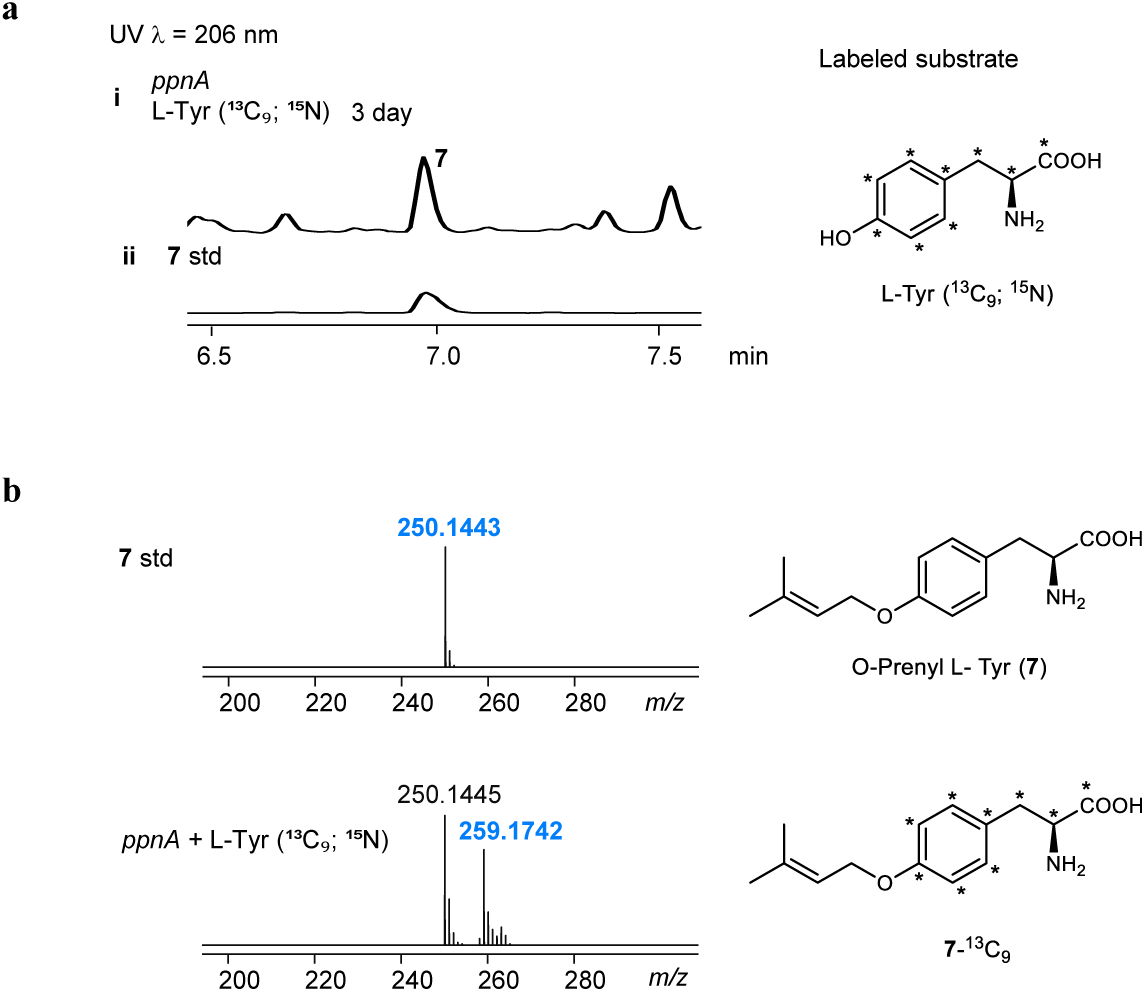
LC-MS trace of the metabolites extracted from *A. nidulans* expressing *ppnA* supplemented with labeled L-Tyr (^13^C_9_; ^15^N). **a.** Compound **7** can be detected in *A. nidulans* expressing *ppnA* after 3 days, supplemented with 0.5 mM L-Tyr (^13^C_9_; ^15^N). **b.** MS analysis of **7** produced by PpnA with the isotope-labeled L-Tyr. Incorporation of ^13^C-labeled L-Tyr into compound **7** was confirmed by a mass shift from ([M + H]^+^ = 250) (unlabeled standard) to **7**-^13^C_9_ ([M + H]^+^ = 259), consistent with full ^13^C_9_ incorporation. The ^15^N label was not incorporated, likely due to deamination by endogenous transaminases.

**Extended Data Fig. 3.**
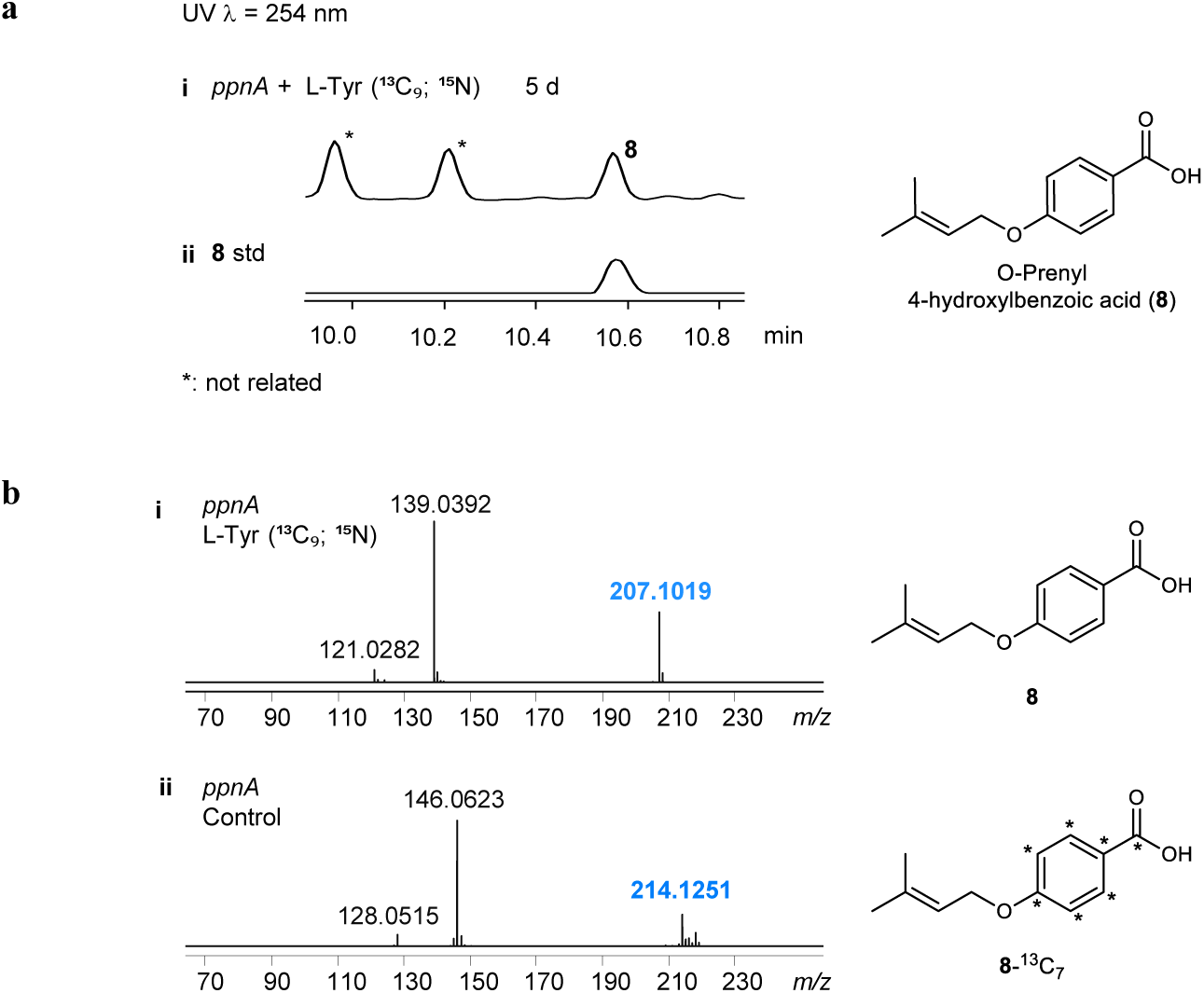
LC-MS analysis of 8 produced by *A. nidulans* expressing *ppnA*. **a.** Compound **8** accumulated in *A. nidulans* expressing *ppnA* after 5 days, supplemented with 0.5 mM L-Tyr (^13^C_9_; ^15^N). **b.** MS analysis of **8**. The mass shift from **8** ([M + H]^+^ = 207.1016 as calculated mass) (unlabeled standard) to **8-**^13^C7 ([M + H]^+^ = 214.1251 as calculated mass), consistent with ^13^C_7_ incorporation.

**Extended Data Fig. 4.**
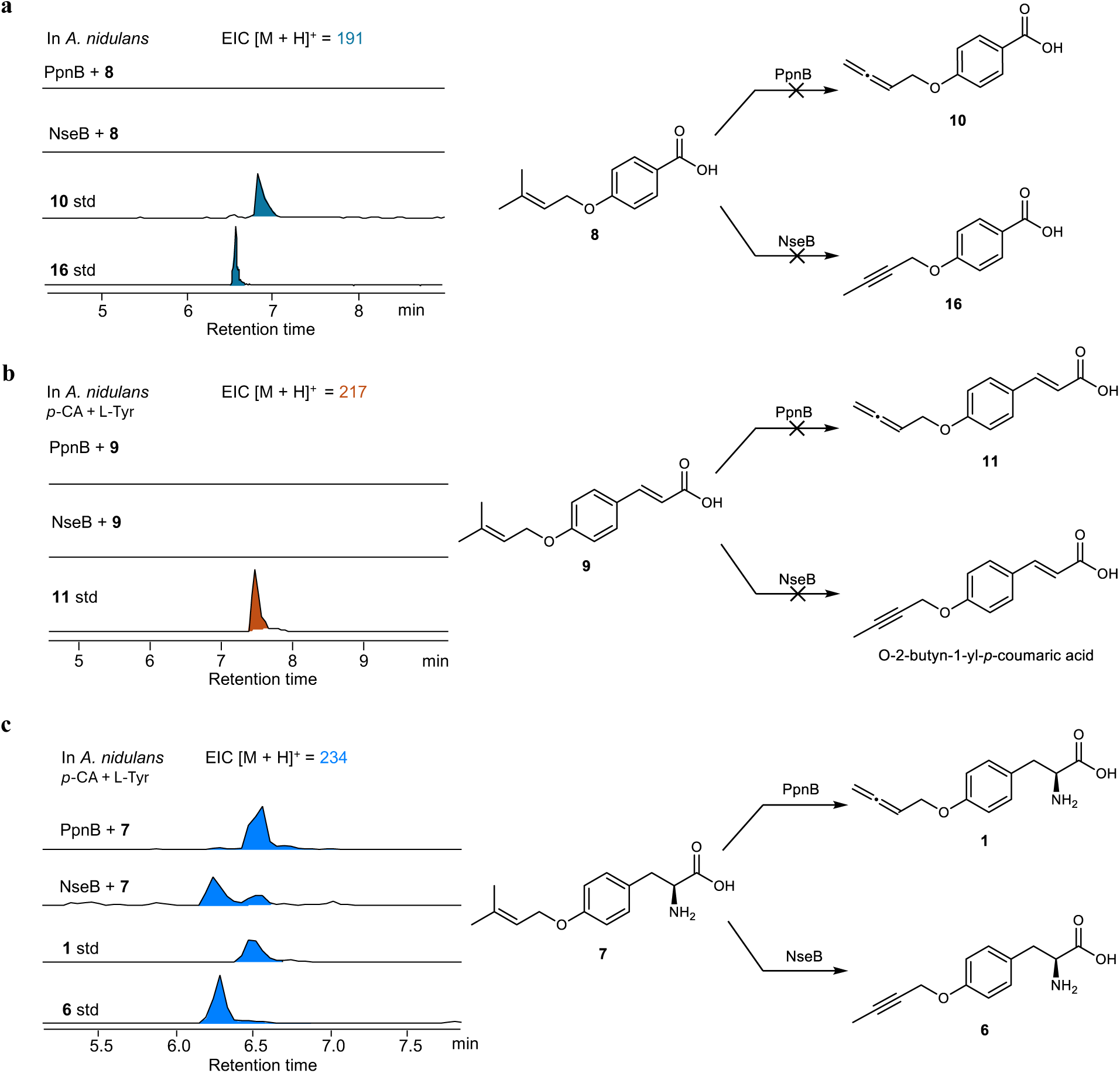
Native substrate of PpnB/NseB. **a.** LC/MS analysis of PpnB and NseB biotransformation in *A. nidulans* with *O*-prenyl-4-hydroxybenzoic acid (**8**). Feeding of **8** to *A. nidulans* expressing PpnB/NseB did not yield corresponding allene (**10**)/internal alkyne product (**16**). **b**. LC/MS analysis of PpnB and NseB biotransformation in *A. nidulans* with *O*-prenyl-*p*-coumaric acid (**9**). Similarly, corresponding allene (**11**)/internal alkyne product was not observed when **9** was fed to *A. nidulans* expressing PpnB/NseB. **c**. LC/MS analysis of PpnB and NseB biotransformation in *A. nidulans* with *O*-prenyl-L-Tyr (**7**). Feeding of the native substrate **7** led to the conversion into **1** and **6**, confirming that PpnB/NseB specifically catalyzes the transformation of **7** as its natural substrate in the pathway. In panels **b** and **c**, L-Tyr and *p*-coumaric acid (*p*-CA) were co-fed to suppress endogenous metabolism of the substrate. Together, these results confirm that **7** is the native substrate for PpnB and NseB, rather than **8** or **9**.

**Extended Data Fig. 5.**
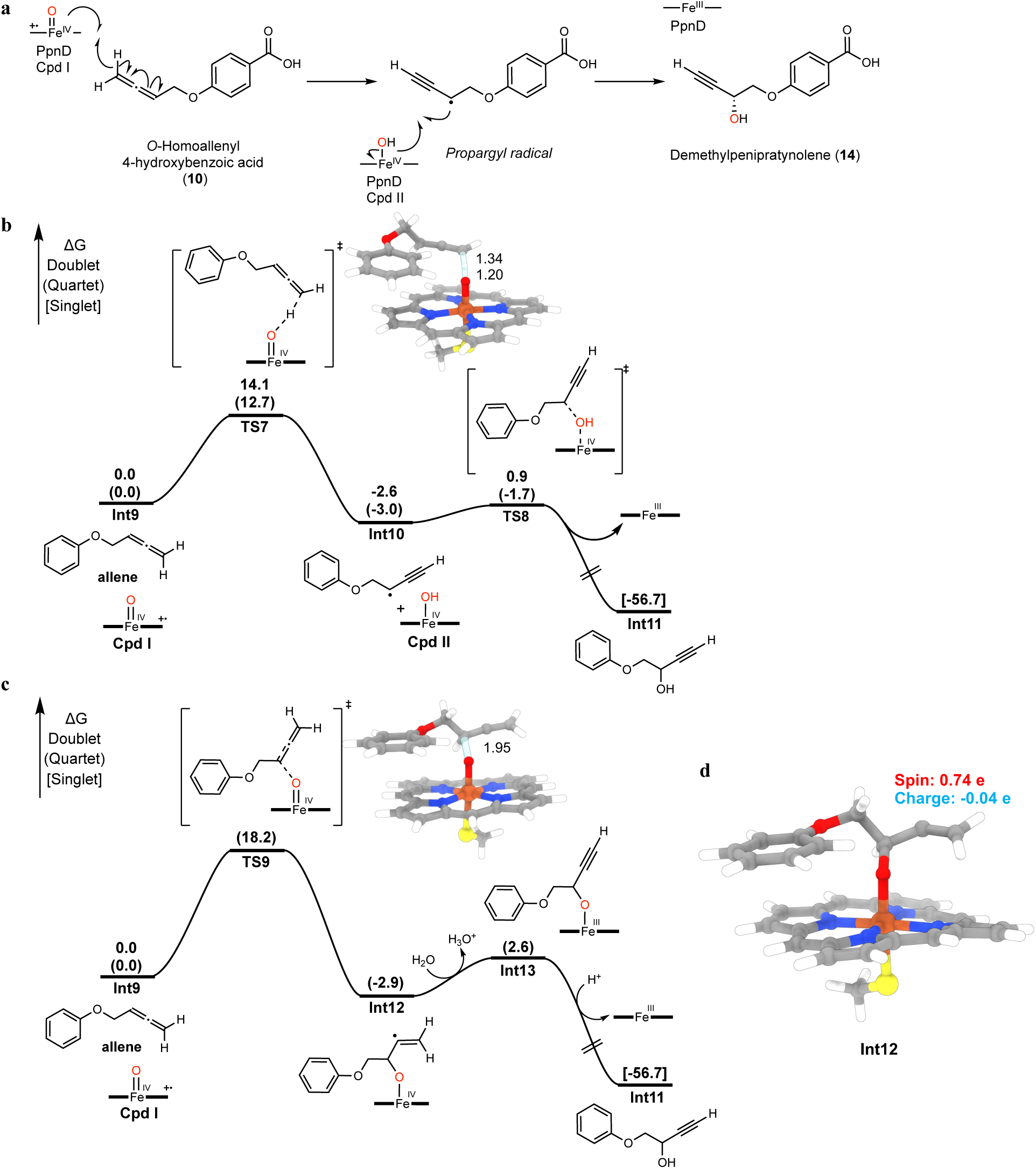
Proposed radical mediated allene to alkyne isomerization by PpnD. **a**, Proposed mechanism of PpnD-catalyzed oxidative isomerization supported by DFT calculations with a model compound. PpnD is proposed to catalyze the abstraction of a hydrogen atom from the terminal allenic carbon of *O*-homoallenyl 4-hydroxybenzoic acid (**10**), forming a propargyl radical intermediate via delocalization, which subsequently undergoes hydroxyl rebound to yield demethylprenipratynolene (**14**). Computed energy profile for allene to alkyne isomerization mechanism using the model substrate is shown in **b** for the isomerization mechanism-initiated HAT and **c** for the isomerization mechanism initiated by Cpd I direct attack. **d**, 3D structure of **Int12** and Hirshfeld spin & charge population. The energies were calculated by B3LYP-D3(BJ)/def2-TZVP/SMD(Et_2_O)//B3LYP-D3(BJ)/def2-SVP/IEFPCM(Et_2_O) at 298 K, 1 atm and are given in kcal/mol. Bond distances are labeled in angstrom.

**Extended Data Fig. 6.**
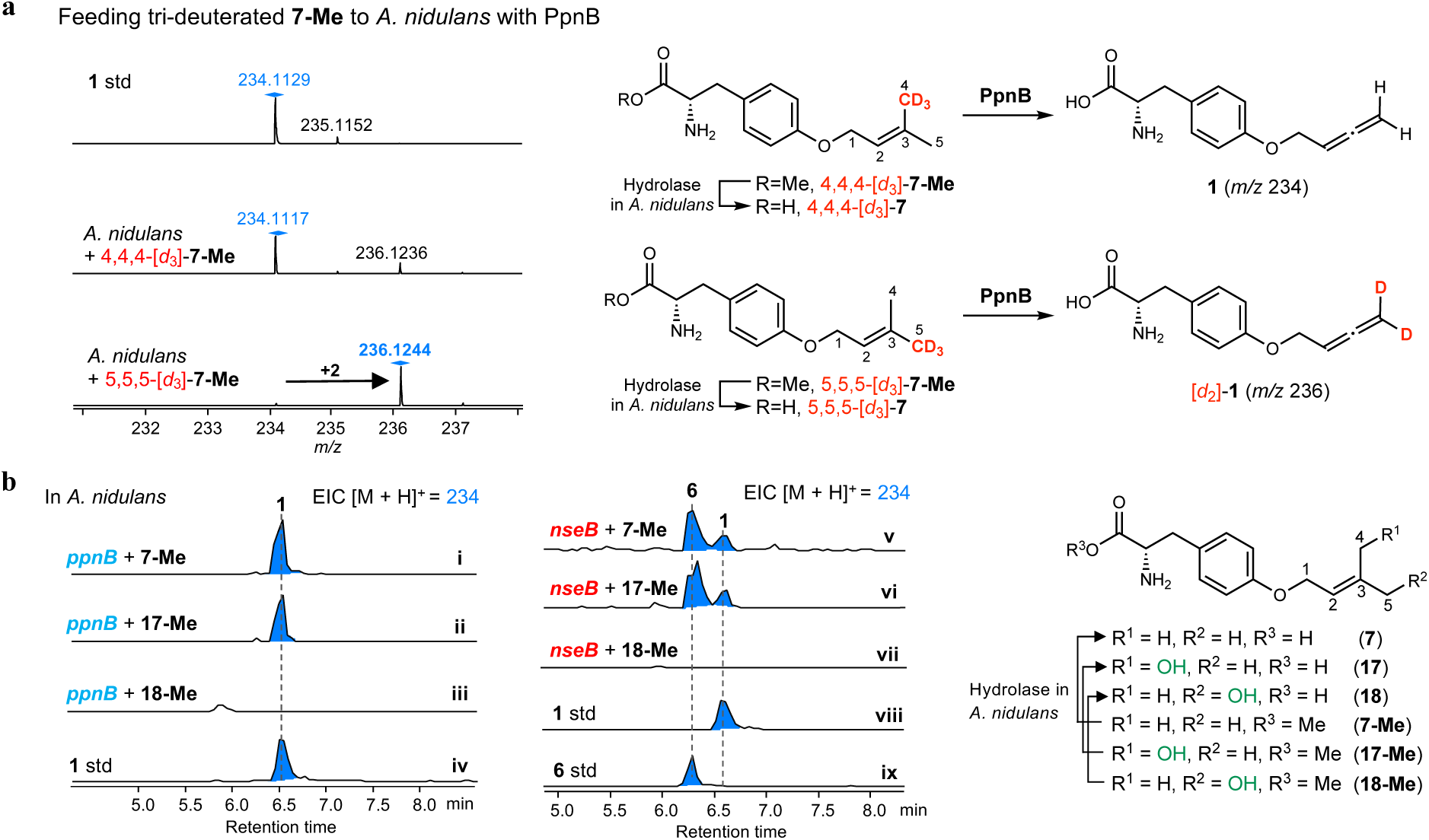
Mechanistic study and the proposed mechanism of the PpnB and NseB-catalyzed allenylation and alkynation. **a**, LC/QTOF analysis of PpnB biotransformation in *A. nidulans* with the methyl ester of deuterated **7** such as 4,4,4-[*d*_3_]-**7-Me** and 5,5,5-[*d*_3_]-**7-Me**, which are hydrolyzed by endogenous hydrolases in *A. nidulans* to be 4,4,4-[*d*_3_]-**7** and 5,5,5-[*d*_3_]-**7**, respectively. +2 Da mass shift of **1** was only observed when 5,5,5-[*d*_3_]-**7-Me** was supplied for the biotransformation. **b**, LC/MS analysis of PpnB and NseB biotransformation in *A. nidulans* with the methyl ester of the putative substrates such as **7** (trace i for PpnB and trace v for NseB), **17** (trace ii for PpnB and trace vi for NseB), **18** (trace iii for PpnB and trace vii for NseB).

**Extended Data Fig. 7.**
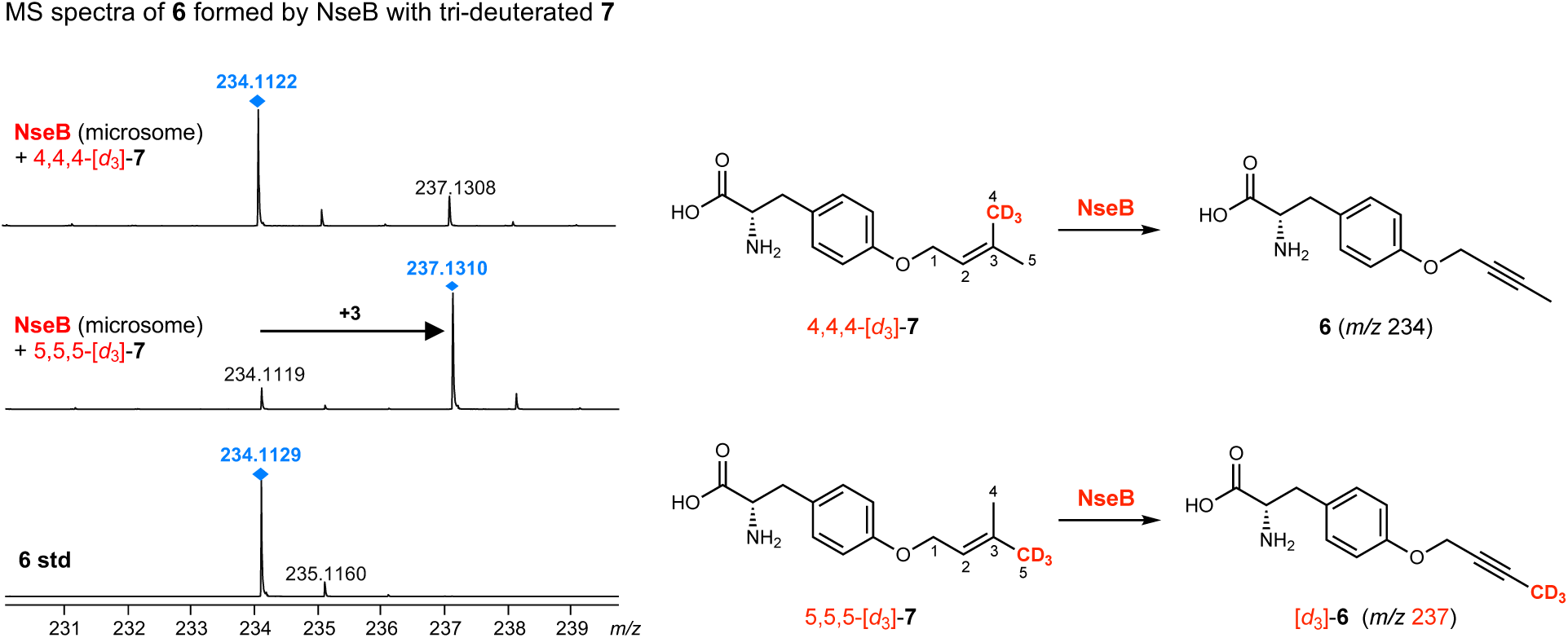
LC–QTOF analysis of NseB-containing microsomes incubated with 4,4,4-[*d*₃]-7 or 5,5,5-[*d*₃]-7. +3 Da mass shift was observed for product **6** only when 5,5,5-[*d*₃]-**7** was supplied in the microsomal assay, supporting retention of the *trans*-C_5_-methyl group during the NseB-catalyzed transformation.

