## supplemental Files for "Biosynthetic allene and alkyne formation by enzymatic prenyl demethylation"

### Table of Contents


**Experimental Procedures**
1. Strains and culture conditions **S5**
2. General molecular biology techniques **S5**
3. Plasmid construction for heterologous expression in *A. nidulans* **S5**
4. Preparation of protoplast of *A. nidulans* and transformations **S5**
5. General procedure for the synthesis of substrates **S6**
6. Amino acid sequences used in this study **S16**

**Supplementary Tables**
**Table 1**. Bioinformatic analysis of *ppn* gene cluster **S18**
**Table 2.** Plasmids used in this study **S19**
**Table 3.** Primers used in this study **S20**
**Table 4.** Spectroscopic data of compound *O*-homoallenyl-L-Tyr (**1**) **S22**
**Table 5.** Spectroscopic data of compound sinuxylamide (**2**) **S23**
**Table 6.** Spectroscopic data of compound penipratynolene (**5**) **S24**
**Table 7.** Spectroscopic data of compound *O*-2-butyn-1-yl L-Tyr (**6**) **S25**
**Table 8.** Spectroscopic data of compound *O*-prenyl L-Tyr (**7**) **S26**
**Table 9.** Spectroscopic data of compound *O*-prenyl 4-hydroxybenzoic acid (**8**) **S27**
**Table 10.** Spectroscopic data of compound *O*-prenyl *p*-coumaric acid **(9)** **S28**
**Table 11.** Spectroscopic data of compound *O*-homoallenyl 4-hydroxybenzoic acid (**10**) **S29**
**Table 12.** Spectroscopic data of compound *O*-homoallenyl *p*-coumatic acid **(11)** **S30**
**Table 13.** Spectroscopic data of compound **13** **S31**
**Table 14.** Spectroscopic data of compound demethylpenipratynolene (**14**) **S32**
**Table 15.** Spectroscopic data of compound *shunt* **15** **S33**
**Table 16.** Spectroscopic data of compound *O*-but-2-ynyl 4-hydroxybenzoic acid (**16**) **S34**
**Table 17.** Spectroscopic data of compound **17**-Me **S35**
**Table 18.** Spectroscopic data of compound **18**-Me **S36**
**Table 19.** Spectroscopic data of compound 4,4,4-[$d_3$]-**7-Me** **S37**
**Table 20.** Spectroscopic data of compound 5,5,5-[$d_3$]-**7-Me** **S38**
**Table 21.** Spectroscopic data of compound **2’-[$d_1$]-7** **S39**
**Table 22.** Spectroscopic data of compound **1’,1’-[$d_2$]-7** **S40**
**Table 23.** Spectroscopic data of compound **19** **S41**
**Table 24.** Spectroscopic data of compound **20** **S42**
**Table 25.** Computed energy components (in Hartree) for all intermediates and transition states **S43**

**Supplementary Figures**
**Figure 1.** Structures of representative allene and alkyne containing natural products in Nature **S44**
**Figure 2.** Allenic ether or alkyne ether containing natural products from fungi **S45**
**Figure 3.** Reported enzymes which catalyze the formation of alkyne or allene **S46**
**Figure 4.** Computational relaxed scan of the directly attack from Int4 to allene and alkyne product **S47**
**Figure 5.** Plasmids used for heterologous expression in *A. nidulans* heterologous host **S48**
**Figure 6.** Heterologous expression of *ppn* cluster **S49**
**Figure 7.** SDS-PAGE gels of purified proteins used in this study **S50**
**Figure 8.** Annotation of PpnA **S51**
**Figure 9.** Substrate specificity of PpnA toward L-tyrosine analogs **S52**
**Figure 10.** Substrate specificity of PpnA toward different prenyl donors **S53**
**Figure 11.** LC-MS analysis of metabolites produced by *A. nidulans* feeding **7**-Me **S54**
**Figure 12.** Diagnostic PCR for *A. nidulans* A1145 Δ*TAL1*Δ*TAL2* mutant **S55**

**Figure 13.** LC-MS analysis of metabolites produced by *A. nidulans* feeding **1**-Me S56
**Figure 14.** LC-MS analysis of metabolites produced by *A. nidulans* expressing *ppnAB* or *ppnAD* feeding with labeled L-Tyr ($^{13}C_9$; $^{15}N$) and 4-hydroxybenzoic acid-$d_4$ (4-HBA-$d_4$) S57
**Figure 15.** Pathways for 4-hydroxybenzoic acid (4-HBA) formation in plants, bacteria, and fungi S58
**Figure 16.** In vitro reaction of PpnC S59
**Figure 17.** Reported native pathway for melearolide A and PF1163A S60
**Figure 18.** PpnD catalyzes oxidative modification of O-prenyl-4-hydroxybenzoic acid S61
**Figure 19.** Overexpression of NseABC leads to production of internal alkyne-containing products S62
**Figure 20.** LC–QTOF analysis of microsomes containing PpnB or NseB incubated with 2-[$d_1$]-**7** S63
**Figure 21.** LC–QTOF analysis of microsomes containing PpnB or NseB incubated with 1,1-[$d_2$]-**7** S64
**Figure 22.** Identification of intermediate **17** from incubation of **7** with PpnB-containing microsomes S65
**Figure 23.** Alternative proposed mechanism of allene and alkyne formation via Baeyer-Villiger intermediate. S66
**Figure 24.** Compounds **15** and **16** are not the intermediate toward **1** or **6** S67
**Figure 25.** Comparison of residues forming substrate binding pocket between PpnB and NseB S68
**Figure 26.** Computed energy profile for the potential cationic allene formation mechanism S70
**Figure 27.** The energy difference between gem-diol and aldehyde + $H_2O$ S71
**Figure 28.** $^{1}H$ NMR spectrum of *O*-homoallenyltyrosine **(1)** in DMSO-$d_6$ S72
**Figure 29.** $^{13}C$ NMR spectrum of *O*-homoallenyltyrosine **(1)** in DMSO-$d_6$ S72
**Figure 30.** HSQC spectrum of *O*-homoallenyltyrosine **(1)** in DMSO-$d_6$ S73
**Figure 31.** HMBC spectrum of *O*-homoallenyltyrosine **(1)** in DMSO-$d_6$ S73
**Figure 32.** $^{1}H$-$^{1}H$ COSY spectrum of *O*-homoallenyltyrosine **(1)** in DMSO-$d_6$ S74
**Figure 33.** $^{1}H$ NMR spectrum of Sinuxylamide B **(2)** in $CD_3OD$ S74
**Figure 34.** $^{13}C$ NMR spectrum of Sinuxylamide B **(2)** in $CD_3OD$ S75
**Figure 35.** $^{1}H$ NMR spectrum of penipratynolene (**5**) in acetone-$d_6$ S75
**Figure 36.** $^{13}C$ NMR spectrum of penipratynolene (**5**) in acetone-$d_6$ S76
**Figure 37.** $^{1}H$ NMR spectrum of *O*-2-butyn-1-yl-L-tyrosine (**6**) in DMSO-$d_6$ S76
**Figure 38.** $^{13}C$ NMR spectrum of *O*-2-butyn-1-yl-L-tyrosine (**6**) in DMSO-$d_6$ S77
**Figure 39.** HSQC spectrum of *O*-2-butyn-1-yl-L-tyrosine (**6**) in DMSO-$d_6$ S77
**Figure 40.** HMBC spectrum of *O*-2-butyn-1-yl-L-tyrosine (**6**) in DMSO-$d_6$ S78
**Figure 41.** $^{1}H$-$^{1}H$ COSY spectrum of *O*-2-butyn-1-yl-L-tyrosine (**6**) in DMSO-$d_6$ S78
**Figure 42.** $^{1}H$ NMR spectrum of *O*-prenyl-L-tyrosine (**7**) in DMSO-$d_6$ S79
**Figure 43.** $^{13}C$ NMR spectrum of *O*-prenyl-L-tyrosine (**7**) in DMSO-$d_6$ S79
**Figure 44.** HSQC spectrum of *O*-prenyl-L-tyrosine (**7**) in DMSO-$d_6$ S80
**Figure 45.** HMBC spectrum of *O*-prenyl-L-tyrosine (**7**) in DMSO-$d_6$ S80
**Figure 46.** $^{1}H$-$^{1}H$ COSY spectrum of *O*-prenyl-L-tyrosine (**7**) in DMSO-$d_6$ S81
**Figure 47.** $^{1}H$ NMR spectrum of *O*-prenyl-benzoic acid (**8**) in $CDCl_3$ S81
**Figure 48.** $^{13}C$ NMR spectrum of *O*-prenyl-benzoic acid (**8**) in $CDCl_3$ S82
**Figure 49.** $^{1}H$ NMR spectrum of *O*-prenyl-*trans*-cinnamic acid (**9**) in $CDCl_3$ S82
**Figure 50.** $^{13}C$ NMR spectrum of *O*-prenyl-*trans*-cinnamic acid (**9**) in $CDCl_3$ S83
**Figure 51.** $^{1}H$ NMR spectrum of *O*-homoallenylbenzoic acid (**10**) in $CDCl_3$ S83
**Figure 52.** $^{13}C$ NMR spectrum of *O*-homoallenylbenzoic acid (**10**) in $CDCl_3$ S84
**Figure 53.** $^{1}H$ NMR spectrum of *O*-homoallenyl-*trans*-cinnamic acid (**11**) in $CDCl_3$ S84
**Figure 54.** $^{13}C$ NMR spectrum of *O*-homoallenyl-*trans*-cinnamic acid (**11**) in $CDCl_3$ S85
**Figure 55**. $^{1}H$ NMR spectrum of compound **13** in $CDCl_3$ S85
**Figure 56.** $^{13}C$ NMR spectrum of compound **13** in $CDCl_3$ S86
**Figure 57.** HSQC spectrum of compound **13** in $CDCl_3$ S86
**Figure 58.** HMBC spectrum of compound **13** in $CDCl_3$ S87
**Figure 59.** $^{1}H$-$^{1}H$ COSY spectrum of compound **13** in $CDCl_3$ S87
**Figure 60.** $^{1}H$ NMR spectrum of demethylpenipratynolene (**14**) in acetone-$d_6$ S88
**Figure 61.** $^{13}C$ NMR spectrum of demethylpenipratynolene (**14**) in acetone-$d_6$ S88

**Figure 62.** $^{1}$H NMR spectrum of compound **15** in $CDCl_3$ **S89**
**Figure 63.** $^{13}$C NMR spectrum of compound **15** in $CDCl_3$ **S89**
**Figure 64.** HSQC spectrum of compound **15** in $CDCl_3$ **S90**
**Figure 65.** HMBC spectrum of compound **15** in $CDCl_3$ **S90**
**Figure 66.** $^{1}$H-$^{1}$H COSY spectrum of compound **15** in $CDCl_3$ **S91**
**Figure 67.** $^{1}$H NMR spectrum of *O*-2-butyn-1-ylbenzoic acid (**16**) in DMSO-$d_6$ **S91**
**Figure 68.** $^{13}$C NMR spectrum of *O*-2-butyn-1-ylbenzoic acid (**16**) in DMSO-$d_6$ **S92**
**Figure 69.** $^{1}$H NMR spectrum of compound **17**-Me in $D_2O$ **S92**
**Figure 70.** $^{13}$C NMR spectrum of compound **17**-Me in $D_2O$ **S93**
**Figure 71.** HSQC spectrum of compound **17**-Me in $D_2O$ **S93**
**Figure 72.** HMBC spectrum of compound **17**-Me in $D_2O$ **S94**
**Figure 73.** $^{1}$H-$^{1}$H COSY spectrum of compound **17**-Me in $D_2O$ **S94**
**Figure 74.** NOESY spectrum of compound **17**-Me in $D_2O$ **S95**
**Figure 75.** $^{1}$H NMR spectrum of compound **18**-Me in DMSO-$d_6$ **S95**
**Figure 76.** $^{13}$C NMR spectrum of compound **18**-Me in DMSO-$d_6$ **S96**
**Figure 77.** HSQC spectrum of compound **18**-Me in DMSO-$d_6$ **S96**
**Figure 78.** HMBC spectrum of compound **18**-Me in DMSO-$d_6$ **S97**
**Figure 79.** $^{1}$H-$^{1}$H COSY spectrum of compound **18**-Me in DMSO-$d_6$ **S97**
**Figure 80**. $^{1}$H NMR spectrum of compound 4’,4’,4’-[$^{2}H_3$]-**7-Me** in DMSO-$d_6$ **S98**
**Figure 81.** $^{13}$C NMR spectrum of compound 4’,4’,4’-[$^{2}H_3$]-**7-Me** in DMSO-$d_6$ **S98**
**Figure 82**. $^{1}$H NMR spectrum of compound 5’,5’,5’-[$^{2}H_3$]-**7-Me** in DMSO-$d_6$ **S99**
**Figure 83.** $^{13}$C NMR spectrum of compound 5’,5’,5’-[$^{2}H_3$]-**7-Me** in DMSO-$d_6$ **S99**
**Figure 84**. $^{1}$H NMR spectrum of compound **19-Me** in $D_2O$ **S100**
**Figure 85.** $^{13}$C NMR spectrum of compound **19-Me** in $D_2O$ **S100**
**Figure 86.** HSQC spectrum of compound **19-Me** in $D_2O$ **S101**
**Figure 87.** HMBC spectrum of compound **19-Me** in $D_2O$ **S101**
**Figure 88.** $^{1}$H-$^{1}$H COSY spectrum of compound **19-Me** in $D_2O$ **S102**
**Figure 89.** NOESY NMR spectrum of compound **19-Me** in $D_2O$ **S102**
**Figure 90**. $^{1}$H NMR spectrum of compound **20** in $CD_3OD$ **S103**
**Figure 91.** $^{13}$C NMR spectrum of compound **20** in $CD_3OD$ **S103**
**Figure 92.** HSQC spectrum of compound **20** in $CD_3OD$ **S104**
**Figure 93.** HMBC spectrum of compound **20** in $CD_3OD$ **S104**
**Figure 94.** $^{1}$H-$^{1}$H COSY spectrum of compound **20** in $CD_3OD$ **S105**
**Figure 95.** NOESY spectrum of compound **20** in $CD_3OD$ **S105**
**Figure 96**. $^{1}$H NMR spectrum of compound **2’-[*d*$_1$]-7** in DMSO-$d_6$ **S106**
**Figure 97.** $^{13}$C NMR spectrum of compound **2’-[*d*$_1$]-7** in DMSO-$d_6$ **S107**
**Figure 98.** HSQC spectrum of compound **2’-[*d*$_1$]-7** in DMSO-$d_6$ **S107**
**Figure 99.** HMBC spectrum of compound **2’-[*d*$_1$]-7** in DMSO-$d_6$ **S108**
**Figure 100.** $^{1}$H-$^{1}$H COSY spectrum of compound **2’-[*d*$_1$]-7** in DMSO-$d_6$ **S108**
**Figure 101**. $^{1}$H NMR spectrum of compound **1’,1’-[*d*$_2$]-7** in DMSO-$d_6$ **S109**
**Figure 102.** $^{13}$C NMR spectrum of compound **1’,1’-[*d*$_2$]-7** in DMSO-$d_6$ **S109**
**Figure 103.** HSQC spectrum of compound **1’,1’-[*d*$_2$]-7** in DMSO-$d_6$ **S110**
**Figure 104.** HMBC spectrum of compound **1’,1’-[*d*$_2$]-7** in DMSO-$d_6$ **S110**
**Figure 105.** $^{1}$H-$^{1}$H COSY spectrum of compound **1’,1’-[*d*$_2$]-7** in DMSO-$d_6$ **S111**

**Supplementary Computational Data** **S111**

### Experimental Procedures

#### 1. Strains and culture conditions

*Penicillium polonicum*[1] was maintained in liquid PDB medium (PDA medium without agar) at 28 °C for isolation of genomic DNA. Genomic DNA of *Nemania serpens* was kindly provided by Prof. Michio Sato. *Aspergillus nidulans* A1145 ΔEMΔST host was previously developed in our lab[2]. The *A. nidulans* strain was grown at 28 °C in CD media (1 L: 10 g Glucose, 50 mL 20 × Nitrate salts, 1 mL Trace elements, pH 6.5, and 20 g/L Agar for solid cultivation) for sporulation or in CD-ST media (1L: 20 g Starch, 20 g Casamino acids, 50 mL 20 × Nitrate salts, 1 mL Trace elements, pH 6.5)[2] for heterologous expression of gene clusters, compound production and RNA extraction. *Escherichia coli* strains were cultivated either on lysogeny broth (LB) agar plates or in LB liquid medium. Growth media were supplied with antibiotics as required at the following concentrations: kanamycin (50 µg $mL^{-1}$), and ampicillin (100 µg $mL^{-1}$). *Saccharomyces cerevisiae* strain BJ5464-NpgA (*MATα ura3-52 his3-Δ200 leu2-Δ1 trp1 pep4::HIS3 prb1 Δ1.6R can1 GAL*) was used as the yeast host for *in vivo* homologous recombination to construct the *A. nidulans* plasmids.

#### 2. General molecular biology techniques

*E. coli* TOP10 cells were used for cloning, following standard recombinant DNA techniques. *E. coli* BL21(DE3) (Novagen) was used as the *E. coli* host for protein expression. DNA restriction enzymes were used as recommended by the manufacturer (New England Biolabs, NEB). Genomic DNA from all fungal strains was prepared using LETS isolation buffer (10 mM Tris-HCI, pH 8.0, 20 mM EDTA, 0.5% SDS, 0.1 M LiCl). PCR was performed using Q5 High-Fidelity DNA Polymerase (NEB). The gene-specific primers are listed in Supplementary Table 2. PCR products were confirmed by DNA sequencing. For isolation of RNA from *A. nidulans* transformants, the strains were grown on CD-ST liquid for 3 days at 28 °C. The RNA extraction steps were performed using RiboPure™ Yeast RNA Isolation Kit (Ambion) following the manufacturer's instructions. Residual genomic DNA in the extracts was digested by DNase I (2 U/mL) (Invitrogen) at 37 °C for 4 hours. SuperScript III First-Strand Synthesis System (Invitrogen) was used for cDNA synthesis with Oligo-dT primers following directions from the user manual.

#### 3. Plasmid construction for heterologous expression in *A. nidulans*

For heterologous expression in *A. nidulans*, three plasmid vectors, pYTU, pYTP, and pYTR containing auxotrophic markers for uracil (*pyrG*), pyridoxine (*pyroA*), and riboflavin (*riboB*), respectively, were used to construct plasmids for *A. nidulans* heterologous expression. Genes in the *ppn* and *nse* cluster were amplified with PCR from the genomic DNA of *A. flavus*. The *gpdA* promoters from *Penicillium oxalicum (*constitutive *POgpdA), A. niger* (constitutive *gpdA*, *glaA* induced by starch) and *Penicillium expansum (*constitutive *PEgpdA)* were amplified by PCR. pYTP and pYTR were digested with PacI/PspXI. pYTU was digested with PacI/NotI. The amplified gene fragments and the corresponding vectors were co-transformed into *S. cerevisiae* strain BJ5464-NpgA for homologous recombination. The yeast plasmids were extracted using Zymoprep™ Yeast Plasmid Miniprep I (Zymo Inc. USA), and then electrically transformed into *E. coli* TOP10 to isolate single plasmids. The plasmids were extracted from *E. coli* using the Zyppy™ Plasmid Miniprep Kit (Zymo Research) and confirmed with sequencing by Laragen and Primordium Lab.

#### 4. Preparation of protoplast of *A. nidulans* and transformation

The transformation of *Aspergillus nidulans* A1145 ΔEMΔST[2], spores were inoculated into 50 mL liquid CD media in a 125-mL flask and germinated at 30 °C shaking at 250 rpm for ~9 h. The 20 X Nitrate salts solution was prepared by dissolving 120 g $NaNO_3$, 10.4 g KCl, 10.4 g $MgSO_4$•$7H_2O$, and 30.4 g $KH_2PO_4$ in 1 L double distilled water. The trace elements solution (100 mL) contained 2.20 g $ZnSO_4$•$7H_2O$, 1.10 g $H_3BO_3$, 0.50 g $MnCl_2$•$4H_2O$, 0.16 g $FeSO_4$•$7H_2O$, 0.16 g $CoCl_2$•$5H_2O$, 0.16 g $CuSO_4$•$5H_2O$, and 0.11 g $(NH_4)_6Mo_7O_{24}$•$4H_2O$. The dropout components for selection for the three expression vectors were uracil/uridine, pyridoxine and riboflavin. *A. nidulans* A1145 ΔEM was initially grown on CD agar plates containing 10 mM uridine, 5 mM uracil, 0.5 µg/mL pyridoxine HCl and 2.5 µg/mL riboflavin at 37°C for 5 days. The germinated spores were harvested by centrifugation at 3,500 rpm for 10 min, and washed with Osmotic buffer (10 mL, 1.2 M $MgSO_4$, 10 mM sodium phosphate buffer, pH 5.8). The mycelia were then mixed with Osmotic buffer (10 mL, 30 mg lysing enzymes from *Trichoderma*, 20 mg Yatalase) in a 125-mL flask. Protoplasts were prepared by incubating the mixture overnight at 30 °C with gentle shaking at 80 rpm. Cells were collected in a 30-mL Corex tube and overlaid gently by 10 mL of Trapping buffer (0.6 M sorbitol, 0.1 M Tris HCI, pH 7.0). Centrifugation at 3,500 rpm for 15 min at 4 °C layered the protoplasts at the interface of the two buffers. The protoplasts were then pipetted to a sterile 15-mL falcon tube and washed with STC buffer (10 mL, 1.2 M sorbitol, 10 mM $CaCl_2$, 10 mM Tris-HCI pH 7.5). The protoplasts were resuspended in STC buffer (1 mL).

For each transformation, 3 µL of each plasmid (>100 ng/µL) was added to 60 µL of the *A. nidulans* A1145 ΔSTΔEM protoplast suspension prepared as above, and the mixture was incubated for 1 h on ice. 600 µL PEG solution (60% PEG, 50 mM of $CaCl_2$, and 50 mM of Tris-HCl, pH 7.5) was added to the protoplast mixture, followed by additional incubation at room temperature for 20 min. The mixture was spread on the CD sorbitol plate (CD solid medium with 1.2 M sorbitol and the appropriate supplements: 10 mM of uridine, 5 mM of uracil, 0.5 µg/mL of pyridoxine HCl, and/or 2.5 µg/mL of riboflavin according to the markers in the transformed plasmids) and incubated at 37 °C for 3-4 days.

**5. General procedure for the synthesis of substrates[3–5]**

**5.1 Chemical synthesis of 17-Me and 17.**

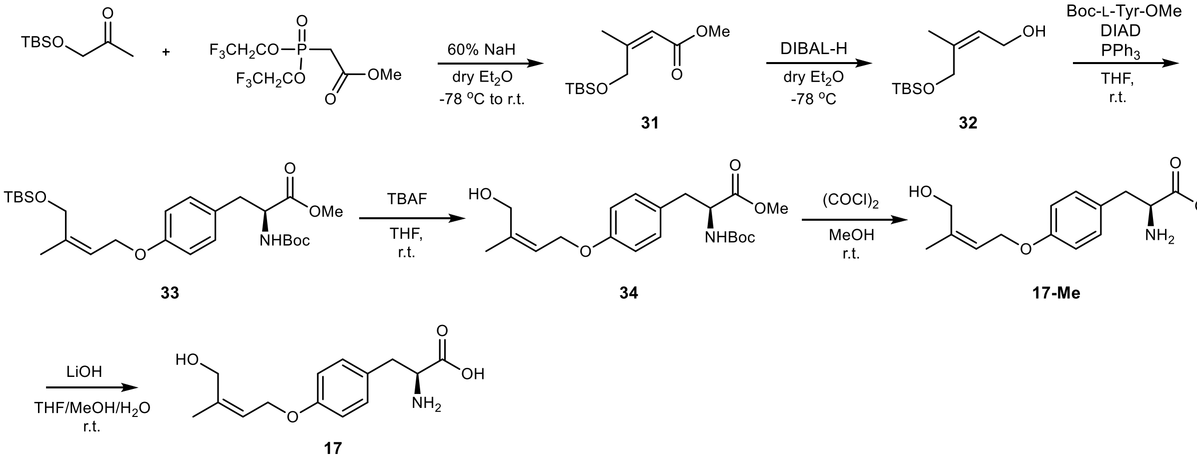


Under an argon atmosphere, to a solution of sodium hydride (NaH, 60 % dispersion in mineral oil, 0.96 g, 24 mmol) in 30 mL of dry diethylether ($Et_2O$) was added triethyl bis(2,2,2-trifluoroethyl) (methoxycarbonylmethyl)phosphonate (4.4 mL, 20 mmol) dropwise at 0 ˚C. After stirring for 1 hour at 0 ˚C, the reaction mixture was cooled down to -78 ˚C. 1-(tert-butyldimethylsilyloxy)-2-propanone (5.2 mL, 24 mmol) was then added to the reaction mixture dropwise. The reaction was allowed stirring at -78 ˚C to r.t. overnight. The mixture was quenched by 50 mL of saturated $NH_4Cl$ aqueous solution. The mixture was extracted with 3 x 50 mL of diethyl ether, and the combined organic layers were washed with brine, dried using anhydrous $Na_2SO_4$. The organic layer was filtered to remove $Na_2SO_4$, and carefully concentrated under reduced pressure to give 1.6 g of crude product **31**. The product was used for next reaction without further purification.

Under an argon atmosphere, to a solution of the crude (1.6 g) in 30 mL of dry $Et_2O$ was added diisobutylaluminium hydride (1 M DIBAL-H in hexane, 16.4 mL, 16.4 mmol) dropwise at -78 ˚C. After addition, the reaction kept stirring at the same temperature until no starting material could be detected by TLC. The reaction mixture was quenched by a few drops of water and 4M NaOH. The resultant solid was removed by filtration and washed with $Et_2O$. The filtrate was carefully concentrated under reduced pressure to give 0.92 g of the corresponding alcohol **32**. The product was used for next reaction without further purification.

A mixture of alcohol (0.92 g), *N*-(*tert*-butoxycarbonyl)-L-tyrosine methyl ester (1.26 g, 4.3 mmol), triphenylphosphine (1.35 g, 5.15 mmol), DIAD (1.10 mL, 5.15 mmol) and 30 mL of THF were stirred for 24 h at room temperature under an argon atmosphere. The reaction mixture was concentrated under reduced pressure. The residue was purified by reverse-phase column using Combi-Flash system with a linear gradient of 5-95% acetonitrile: water supplemented with 0.1% formic acid to give 1.24 g of the corresponding compound **33** as yellow oil (13% in 3 steps).

To a solution of the compound (1.24 g, 2.5 mmol) in 20 mL of THF was added 1M TBAF in THF (3.0 mL, 3.0 mmol) dropwise at r.t. After stirring the reaction mixture at r.t. overnight, 30 mL of water was added to the reaction mixtute. The mixture was extracted by 3 × 30 mL of EtOAc, and the combined organic layer was washed with 50 mL of brine, and dried by anhydrous $Na_2SO_4$. After the filtration to remove $Na_2SO_4$, the organic layer was concentrated under reduced pressure to give 0.73 g of the compound **34**. The product was used for next reaction without further purification. $^1H$ NMR (500 MHz, DMSO-$d_6$) δ 7.25 (d, *J* = 8.1 Hz, 1H), 7.12 (dd, *J* = 9.0, 3.0 Hz, 2H), 6.82 (d, *J* = 8.1 Hz, 2H), 5.42 (t, *J* = 6.5 Hz, 1H), 4.55 (d, *J* = 6.4 Hz, 2H), 4.10 (ddd, *J* = 9.8, 8.0, 5.1 Hz, 1H), 3.99 (s, 2H), 3.60 (s, 3H), 2.90 (dd, *J* = 13.8, 5.2 Hz, 1H), 2.77 (dd, *J* = 13.8, 9.9 Hz, 1H), 1.75 (d, *J* = 1.5 Hz, 3H), 1.33 (s, 9H). $^{13}C$ NMR (126 MHz, DMSO) δ 21.0, 28.1, 35.6, 51.7, 55.5, 59.8, 63.6, 78.3, 114.3, 121.2, 129.3, 130.1, 140.3, 155.4, 157.0, 172.7.

Compound **34** (0.73 g, 1.82 mmol) was dissolved in MeOH (15 mL) and allowed to stir at room temperature for 5 min. oxalyl chloride (0.50 mL, 5.80 mmol) was then added to the solution *via* micropipette directly into the reaction solvent mixture dropwise. The reaction mixture was allowed to stir at r.t. for 4 h. Upon complete consumption of the *N*-Boc-protected compound, water (10 mL) was added to the flask slowly. The crude material was subsequently extracted with 3 x 30 mL of EtOAc and washed with 30 mL of water and 30 mL of brine. The organic layer was dried over anhydrous $Na_2SO_4$, filtered, and concentrated under the reduced pressure. The residue was purified by reverse-phase column using Combi-Flash system with a gradient of 5-20% (acetonitrile : water) supplemented with 0.1% formic acid to give 0.33 g of the corresponding compound **17-Me** as yellow oil (47% in 2 steps). **17**-Me further undergo hydrolysis to give **17** following the method for chemical synthesis of **7-Me** and **7**. $^1H$ NMR (500 MHz, $D_2O$) δ 7.14 (d, *J* = 8.6 Hz, 2H), 6.88 (d, *J* = 8.6 Hz, 2H), 5.57 (dt, *J* = 7.0, 4.3 Hz, 1H), 4.55 (d, *J* = 6.9 Hz, 2H), 4.11 (s, 2H), 3.40 (dd, *J* = 7.4, 5.4 Hz, 1H), 2.89 (dd, *J* = 13.7, 5.4 Hz, 1H), 2.72 (dd, *J* = 13.6, 7.4 Hz, 1H), 1.78 (m, 3H). $^{13}C$ NMR (126 MHz, $D_2O$) δ 20.4, 39.8, 57.3, 59.9, 64.1, 115.0, 115.0, 122.0, 130.5, 130.5, 131.1, 140.8, 156.2, 182.3. HRMS (ESI, M+H$^+$) calculated for $C_{14}H_{20}O_4^+$ 266.1387; found 266.1387.

**5.2 Chemical synthesis of 18-Me and 18.**

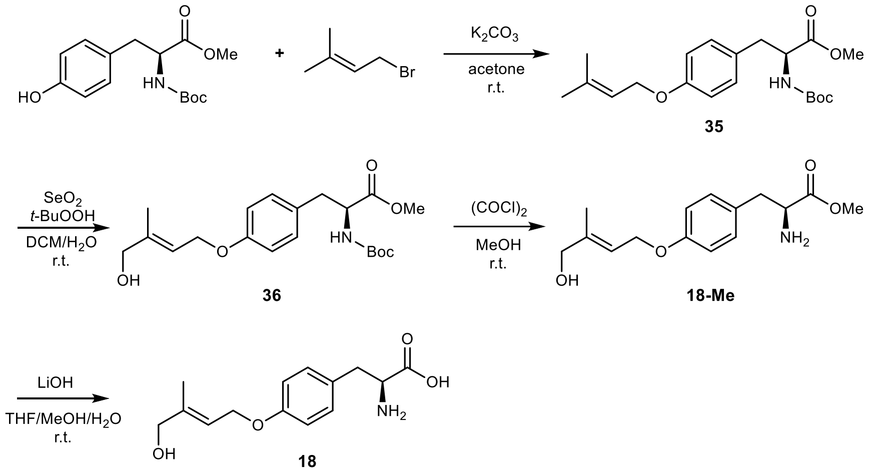


A suspension of Boc-L-Tyr-OMe (0.89 g, 3.0 mmol, 1.0 equiv) and $K_2CO_3$ (0.50 g, 3.6 mmol, 1.2 equiv) in 20 mL of acetone was stirred at room temperature, and 3,3-dimethylallyl bromide (0.42 mL, 3.6 mmol, 1.2 equiv) was added dropwise. The mixture was stirred at r.t. for 18 h, and then poured into 150 mL of saturated $NaHCO_3$ aqueous solution. The mixture was extracted by 3 x 30 mL of EtOAc , and the combined organic layer was washed with brine, dried using anhydrous $Na_2SO_4$, and concentrated under reduced pressure to give 1.01 g of the compound **35** as colorless oil. $^1$H NMR (500 MHz, DMSO-$d_6$) δ 7.25 (d, $J$ = 8.0 Hz, 1H), 7.12 (d, $J$ = 8.2 Hz, 2H), 6.83 (d, $J$ = 8.5 Hz, 2H), 5.45 – 5.36 (m, 1H), 4.47 (d, $J$ = 6.6 Hz, 2H), 4.10 (ddd, $J$ = 9.9, 8.1, 5.1 Hz, 1H), 2.90 (dd, $J$ = 13.8, 5.2 Hz, 1H), 2.77 (dd, $J$ = 13.8, 9.9 Hz, 1H), 1.73 (s, 3H), 1.69 (s, 3H), 1.33 (s, 9H). $^{13}$C NMR (126 MHz, DMSO) δ 18.0, 25.4, 28.1, 51.7, 55.5, 64.2, 78.2, 114.3, 120.1, 129.3, 130.0, 136.8, 155.4, 157.1, 172.7.

1.01 g of **35** was dissolved in DCM (18 mL), followed by the addition of 70% $t$-BuOOH aqueous solution (2.0 mL, 9.0 mmol) and $SeO_2$ (66 mg, 0.60 mmol). The mixture was stirred for 3 days at r.t. The reaction mixture was poured into 300 mL of water. The mixture was extracted with 3 x 30 mL of EtOAc, and the combined organic layer was washed with brine, dried using anhydrous $Na_2SO_4$, and concentrated under reduced pressure to give crude product containing alcohol **36** and aldehyde **48**. The crude mixture was further purified by reverse-phase column using Combi-Flash instrument with a gradient of 5-95% (acetonitrile : water) supplemented with 0.1% formic acid to give 0.21 g of **36** as pale yellow oil (20% yield) and 0.38 g of aldehyde **48** as colorless oil. The NMR data of synthetic **36:** $^1$H NMR (500 MHz, $CDCl_3$) δ 7.02 (d, $J$ = 8.2 Hz, 2H), 6.87 – 6.77 (m, 2H), 5.75 (tq, $J$ = 6.5, 1.5 Hz, 1H), 4.56 (d, $J$ = 6.5 Hz, 2H), 4.53 (m, 1H), 4.08 (s, 2H), 3.71 (s, 3H), 3.08 – 2.94 (m, 2H), 1.75 (d, $J$ = 1.2 Hz, 3H), 1.41 (s, 9H). $^{13}$C NMR (126 MHz, $CDCl_3$) δ 14.1, 28.4, 37.6, 52.4, 54.7, 64.4, 67.9, 80.4, 114.9, 120.4, 128.1, 130.4, 139.8, 155.5, 157.8, 172.6. The NMR data of synthetic **48:** $^1$H NMR (500 MHz, $CDCl_3$) δ 9.45 (s, 1H), 7.04 (d, $J$ = 8.3 Hz, 2H), 6.82 (d, $J$ = 8.6 Hz, 2H), 6.66 (tq, $J$ = 5.4, 1.4 Hz, 1H), 5.04 (d, $J$ = 8.4 Hz, 1H), 4.83 (dq, $J$ = 5.5, 1.2 Hz, 2H), 4.52 (q, $J$ = 6.6 Hz, 1H), 3.69 (s, 3H), 3.10 – 2.92 (m, 2H), 1.81 (d, $J$ = 1.3 Hz, 3H), 1.39 (s, 9H). $^{13}$C NMR (126 MHz, $CDCl_3$) δ 9.7, 28.4, 37.5, 52.3, 54.6, 64.8, 80.1, 114.7, 129.0, 130.6, 139.8, 147.9, 157.2, 172.5, 194.2.

Compound **36** (0.21 g, 0.52 mmol) was dissolved in MeOH (10 mL) and allowed to stir at room temperature for 5 min. oxalyl chloride (0.14 mL, 1.60 mmol) was then added to the solution *via* micropipette directly into the reaction solvent mixture dropwise. The reaction mixture was allowed to stir at r.t. for 4 h. Upon complete consumption of the $N$-Boc-protected compound, water (10 mL) was added to the flask slowly. The crude material was subsequently extracted with 3 x 30 mL of EtOAc and washed with 30 mL of water and 30 mL of brine. The organic layer was dried over anhydrous $Na_2SO_4$, filtered, and concentrated under the reduced pressure. The residue was purified by reverse-phase column using Combi-Flash system with a gradient of 5-20% (acetonitrile : water) supplemented with 0.1% formic acid to give 0.10 g of the corresponding compound **18-Me** as yellow oil (69% yield). **18**-Me further undergo hydrolysis to give **18** following the method for chemical synthesis of **7-Me** and **7**. $^1$H NMR (500 MHz, $D_2O$) δ 7.05 (d, $J$ = 8.5 Hz, 2H), 6.87 – 6.77 (m, 2H), 5.54 (dp, $J$ = 6.7, 1.7 Hz, 1H), 4.50 (d, $J$ = 6.7 Hz, 2H), 3.87 (s, 2H), 3.30 (m, 1H), 2.78 (dd, $J$ = 13.6, 5.5 Hz, 1H), 2.63 (dd, $J$ = 13.7, 7.3 Hz, 1H), 1.58 (s, 3H). $^{13}$C NMR (126 MHz, $D_2O$) δ 13.1, 25.0, 39.8, 48.8, 57.3, 64.7, 66.4, 67.8, 115.0, 115.0, 119.1, 130.5, 130.5, 131.1, 141.1, 156.3, 182.4. HRMS (ESI, M+H$^+$) calculated for $C_{14}H_{20}O_4^+$ 266.1387; found 266.1401.

#### 5.3 Chemical synthesis of 4’,4’,4’-[*d*3]-7-Me and -7 and 5’,5’,5’-[*d*3]-7-Me and -7.

#### 5.3.1 Synthesis of (*E*)-37 and (*Z*)-37.

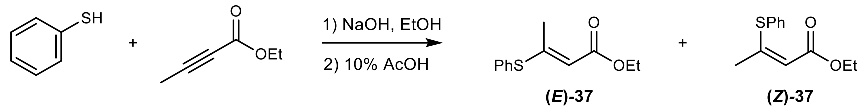


To a stirring suspension of NaOH (1.2 g, 30 mmol) in 60 mL of EtOH was added PhSH (3.5 mL, 34.2 mmol) and ethyl 2-butynoate (3 mL, 25.7 mmol). After stirring at r.t. for 4 hours, the reaction was acidified with 15 mL of 10% AcOH. The organic layer was then washed with 60 mL of water and 60 mL of brine, dried over anhydrous $Na_2SO_4$, filtered and concentrated under reduced pressure. The residue was purified by normal-phase silica gel chromatography using Combi-Flash instrument with a gradient of 10% ($Et_2O$ : hexane) to give 2.94 g **(*E*)-37** (51%) and 0.92 g **(*Z*)-37** (16%) as yellow oil.

#### 5.3.2 Synthesis of 5’,5’,5’-[*d*3]-7-Me and -7.

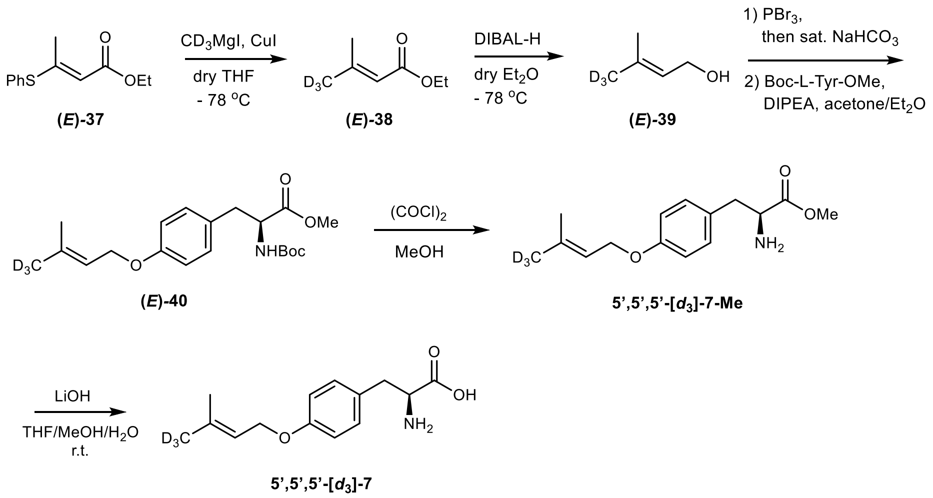

#### 5.3.3 Synthesis of 4’,4’,4’-[$d_3$]-7-Me and -7.

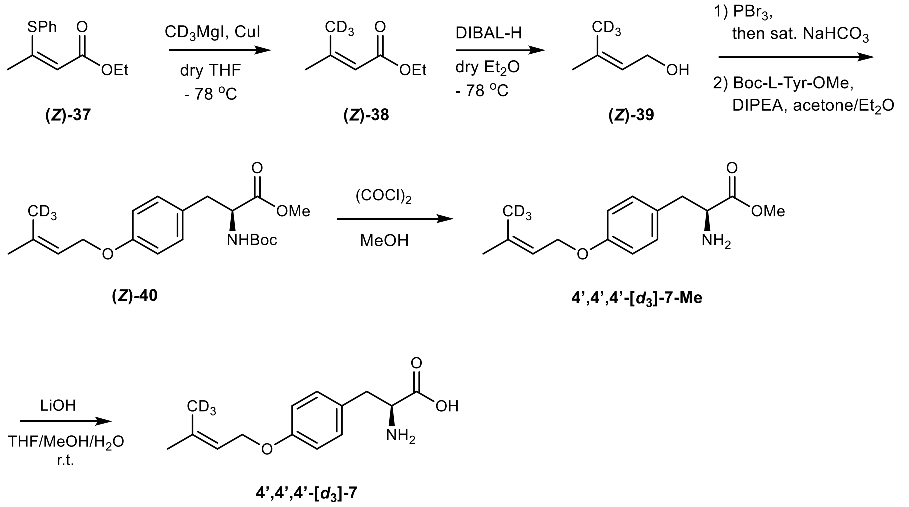


Compound **(*E*)-38** and **(*Z*)-38**. Under an argon atmosphere, to a stirring suspension of CuI (3 g, 16 mmol) in 30 mL dry THF at -78 °C was added $CD_3MgI$ (1 M solution in $Et_2O$, 20 mL, 20 mmol). After the mixture was stirred at -78 °C for 15 minutes, a solution of **(*E*)-37** (888 mg, 4 mmol) in 12 mL of dry THF was added dropwise and stirred at -78 °C for 4 hours. The mixture was then stirred at 0 °C and 50 mL of saturated $NH_4Cl$ was added slowly. The aqueous layer was extracted with $Et_2O$ twice, the organic layers were combined and washed with 30 mL of water and 30 mL of brine, dried over anhydrous $Na_2SO_4$, filtered and concentrated under reduced pressure carefully. The residue was purified by normal-phase silica gel chromatography using Combi-Flash instrument with a gradient of 2.5% ($Et_2O$ : pentane) to give 192 mg of **(*E*)-38** (37%) as white oil. **(*Z*)-38** (222 mg, 42%, white oil) was acquired through the same procedure.

Compound **(*E*)-39** and **(*Z*)-39**. Under an argon atmosphere, to a stirring solution of **(*E*)-38** (475 mg, 3.6 mmol) in 2.5 mL of dry $Et_2O$ was added DIBAL-H (1 M DIBAL-H in hexane, 9 mL, 9 mmol) dropwise at - 78 °C. After stirring for 1 hour, the mixture was then stirred at 0 °C and a few drops of water and 4 M NaOH was added. The resultant solid was removed by filtration and washed twice with 5 mL of cold $Et_2O$, then dried over anhydrous $Na_2SO_4$ at 0 °C. The product solution containing **(*E*)-39** was filtered and used for next reaction without further purification.

Compound **(*E*)-40** and **(*Z*)-40**. $PBr_3$ (266 µL, 2.8 mmol) in 2 mL of dry $Et_2O$ was added dropwise into the resulted product solution upon stirring at 0 °C for 2 hours. The reaction was quenched at 0 °C by adding 15 mL of saturated $NaHCO_3$. The aqueous layer was extracted with 2 x 5 mL of $Et_2O$ and the organic layers were combined. Boc-L-Tyr-OMe (1.26 g, 4.3 mmol) in 20 mL of 1:1 acetone/$Et_2O$ and DIPEA (1.4 mL, 8 mmol) were added into the resulted organic layer and stirred for 16 hours. The reaction was concentrated under reduced pressure and the residue was purified by reverse-phase column using Combi-Flash instrument with a gradient of 30-95% (acetonitrile : water) to give 588 mg of **(*E*)-40** (45%) as yellow oil. **(*Z*)-40** (563 mg, 43%) was acquired through the same procedure.

Compound 4’,4’,4’-[$d_3$]-**7-Me** and 5’,5’,5’-[$d_3$]-**7-Me**. **(*E*)-40** (363 mg, 1 mmol) was dissolved in MeOH (10 mL) and allowed to stir at room temperature for 5 min. Oxalyl chloride (170 µL, 2 mmol) was then added to the solution into the reaction mixture dropwise. The reaction mixture was allowed to stir at r.t. for 4 h. Upon complete consumption of **(*E*)-40**, 10 mL of water was added slowly. The crude material was subsequently extracted with 3 x 10 mL of EtOAc, combined and washed with 30 mL of water and 30 mL of brine. The organic layer was dried over anhydrous $Na_2SO_4$, filtered, and concentrated under the reduced pressure. The residue was purified by reverse-phase column using Combi-Flash system with a gradient of 10-50% (acetonitrile : water) to give 134 mg of 5',5',5'-[$d_3$]-**7-Me** (51%) as yellow oil. 4',4',4'-[$d_3$]-**7-Me** (130 mg, 50%) was acquired through the same procedure. 4',4',4'-[$d_3$]-**7-Me** and 5',5',5'-[$d_3$]-**7-Me** further undergo hydrolysis to give 4',4',4'-[$d_3$]-**7** and 5',5',5'-[$d_3$]-**7** following the method for chemical synthesis of **7-Me** and **7**. 4',4',4'-[$d_3$]-**7**: $^1$H NMR (500 MHz, $CD_3OD$) δ 7.17 (d, $J$ = 8.1 Hz, 2H), 6.85 (d, $J$ = 8.1 Hz, 2H), 5.44 (t, $J$ = 6.7 Hz, 1H), 4.49 (d, $J$ = 6.7 Hz, 2H), 3.55 (dd, $J$ = 8.5, 4.6 Hz, 1H), 3.12 (dd, $J$ = 14.1, 4.6 Hz, 1H), 2.82 (dd, $J$ = 14.0, 8.3 Hz, 1H), 1.77 (s, 3 H). $^{13}$C NMR (126 MHz, $CD_3OD$) δ 18.1, 25.8, 39.9, 58.4, 65.8, 115.9, 115.9, 121.4, 130.6, 131.4, 131.4, 138.3, 159.2, 178.2. HRMS (ESI, M+H$^+$) calculated for $C_{14}H_{17}D_3NO_3^+$ 253.1626; found 253.1624. 5',5',5'-[$d_3$]-**7**: $^1$H NMR (500 MHz, $CD_3OD$) δ 7.20 (d, $J$ = 8.1 Hz, 2H), 6.89 (d, $J$ = 8.1 Hz, 2H), 5.44 (t, $J$ = 6.7 Hz, 1H), 4.52 (d, $J$ = 6.6 Hz, 2H), 3.72 (dd, $J$ = 8.8, 4.3 Hz, 1H), 3.24 (dd, $J$ = 14.6, 4.4 Hz, 1H), 2.94 (dd, $J$ = 14.6, 8.7 Hz, 1H), 1.74 (s, 3H). $^{13}$C NMR (126 MHz, MeOD) δ 18.1, 25.8, 37.5, 57.7, 65.8, 116.2, 116.2, 121.3, 129.0, 131.4, 131.4, 138.5, 159.6, 173.9. HRMS (ESI, M+H$^+$) calculated for $C_{14}H_{17}D_3NO_3^+$ 253.1626; found 253.1620.

#### 5.4 Chemical synthesis of compound 1.

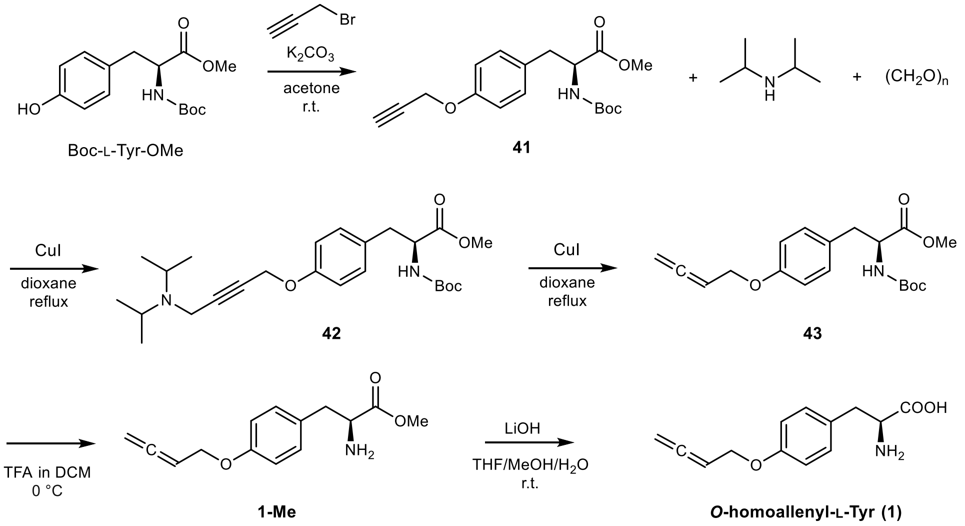


A suspension of Boc-L-Tyr-OMe (2.95 g, 10.0 mmol, 1.0 equiv) and $K_2CO_3$ (1.66 g, 12.0 mmol, 1.2 equiv) in 30 mL of acetone was stirred at room temperature, and propargyl bromide (0.91 mL, 12.0 mmol, 1.2 equiv) was added dropwise. The mixture was stirred at r.t. for 18 h, and then poured into 150 mL of saturated $NaHCO_3$ aqueous solution. The mixture was extracted by 3 x 50 mL of EtOAc, and the combined organic layer was washed with brined, dried using anhydrous $Na_2SO_4$, and concentrated under reduced pressure to give 2.1 g of crude **41**.

0.70 g of the crude residue was dissolved in 30 mL of 1,4-dioxane, and CuI (84mg, 0.44 mmol, 0.2 equiv), diisopropylamine (0.62 mL, 4.4 mmol, 2.0 equiv), and paraformaldehyde (0.17 g, 5.5 mmol, 2.5 equiv) were added. The mixture was heated to reflux and stirred overnight. After cooling to r.t., the mixture was filtered through Celite to remove precipitates and washed with EtOAc. The combined filtrates were concentrated under reduced pressure to afford crude compound **43**. The crude compound **43** was further purified by reverse-phase column using Combi-Flash instrument with a gradient of 5-95% (acetonitrile : water) supplemented with 0.1% formic acid to give 0.21 g of **43** as brown oil. $^1$H NMR (500 MHz, $CDCl_3$) δ 7.02 (d, $J$ = 8.1 Hz, 2H), 6.84 (d, $J$ = 8.2 Hz, 2H), 5.38 (p, $J$ = 6.7 Hz, 1H), 4.95 (d, $J$ = 8.3 Hz, 1H), 4.86 (dd, $J$ = 6.2, 3.1 Hz, 2H), 4.54 (dd, $J$ = 6.5, 3.0 Hz, 2H), 3.70 (s, 3H),

3.02 (qd, $J$ = 14.0, 5.9 Hz, 2H), 1.42 (s, 9H). $^{13}C$ NMR (126 MHz, $CDCl_3$) δ 28.4, 37.6, 52.3, 54.6, 66.0, 80.0, 87.2, 115.1, 128.4, 130.4, 155.2, 157.6, 172.6, 209.6.

0.21 g of **43** was dissolved in $CH_2Cl_2$ (5 mL), and trifluoroacetic acid (TFA, 5 mL) was added dropwise at 0 °C. The mixture was stirred at rt for 0.5 h and then concentrated under reduced pressure to give the deprotected intermediate to give 0.20 g of **1-Me** as brown solid with quantitative yield. $^1H$ NMR (500 MHz, DMSO-$d_6$) δ 7.16 – 7.06 (m, 2H), 6.96 – 6.86 (m, 2H), 5.48 (p, $J$ = 6.7 Hz, 1H), 4.97 (dt, $J$ = 6.7, 2.6 Hz, 2H), 4.54 (dt, $J$ = 6.7, 2.6 Hz, 2H), 4.25 (t, $J$ = 6.6 Hz, 1H), 3.67 (s, 3H), 3.03 (qd, $J$ = 14.2, 6.6 Hz, 2H); $^{13}C$ NMR (126 MHz, DMSO) δ 35.2, 52.6, 53.5, 65.1, 76.9, 86.9, 114.9, 126.5, 130.5, 157.3, 169.5, 208.7.

0.20 g of **1-Me** was dissolved in THF/MeOH/$H_2O$ (4:2:3 (v/v/v), 9.0 mL), LiOH (23 mg, 0.97 mmol, 1.2 equiv) was added, and the mixture was stirred at rt for 3 h. After completion (LCMS monitoring), the reaction mixture was directly purified by reverse-phase column using Combi-Flash instrument with a gradient of 5-30% (acetonitrile : water) supplemented with 0.1% formic acid to give 0.11 g of **1** (58% yield) as yellow solid.

#### 5.5 Chemical synthesis of compound 6.

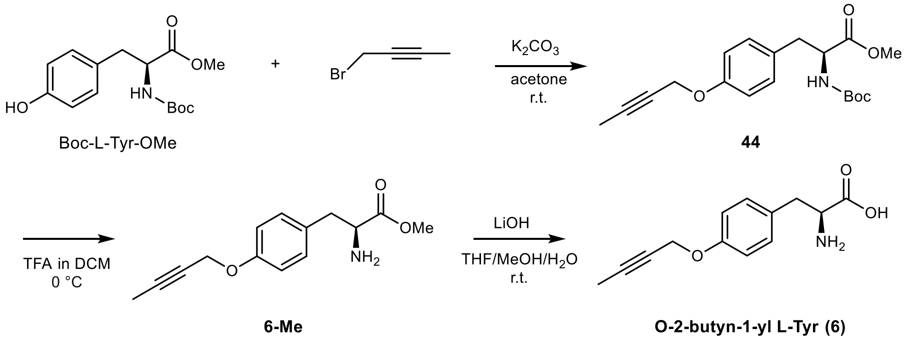


1-bromo-2-butyne (0.32 mL, 3.6 mmol) was added to a suspension of Boc-L-Tyr-OMe (0.88 g, 3.0 mmol) and $K_2CO_3$ (0.50 g, 3.6 mmol) in 20 mL of acetone. The mixture was stirred at r.t. for 18 h, and then poured into 100 mL of saturated $NaHCO_3$ aqueous solution. The mixture was extracted by 3 x 30 mL of EtOAc , and the combined organic layer was washed with brined, dried using anhydrous $Na_2SO_4$, and concentrated under reduced pressure to give 0.90 g of crude **44**. The 0.20 g of crude **44** was dissolved in 5 mL of $CH_2Cl_2$, and 5 mL of trifluoroacetic acid was added dropwise at 0 °C. The reaction mixture was stirred at r.t. until deprotection was complete. The solvent and excess TFA were removed under reduced pressure to give crude **6-Me**. The crude **6-Me** was dissolved in THF/MeOH/$H_2O$ (4:2:3 (v/v/v), 9.0 mL), LiOH (24 mg, 1.0 mmol) was added, and the mixture was stirred at rt for 3 h. After completion (LCMS monitoring), the reaction mixture was directly purified by reverse-phase column using Combi-Flash instrument with a gradient of 5-30% (acetonitrile : water) supplemented with 0.1% formic acid to give 0.10 g of **6** as yellow solid.

#### 5.6 Chemical synthesis of O-prenylbenzoic acid (8).

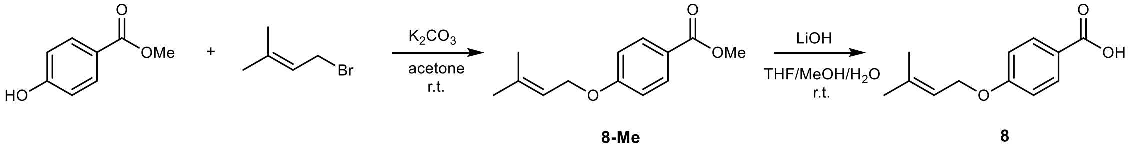

3,3-Dimethylallyl bromide (0.14 mL, 1.2 mmol, 1.2 equiv) was added to a suspension of 4-hydroxybenzoic acid methyl ester (0.15 g, 1.0 mmol, 1.0 equiv) in 15 mL of acetone. The mixture was stirred at r.t. for 18 h, and then poured into 100 mL of saturated $NaHCO_3$ aqueous solution. The mixture was extracted by 3 x 30 mL of EtOAc, and the combined organic layer was washed with brined, dried using anhydrous $Na_2SO_4$, and concentrated under reduced pressure to give 201 mg of crude **8-Me**. The crude **8-Me** was dissolved in THF/MeOH/$H_2O$ (4:2:3 (v/v/v), 9.0 mL), LiOH (48 mg, 2.0 mmol) was added, and the mixture was stirred at rt for 1 h. After completion (LCMS monitoring), the pH of the reaction mixture was adjusted to ~ 3, and extracted with 3 x 30 mL of EtOAc. The combined organic layer was washed with brined, dried using anhydrous $Na_2SO_4$, and concentrated under reduced pressure to give crude **6**, which was subsequently purified by reverse-phase column using Combi-Flash instrument with a gradient of 5-95% (acetonitrile : water) supplemented with 0.1% formic acid to give 0.15g of **6** as white solid (73% yield, 2 steps).

#### 5.7 Chemical synthesis of O-prenyl-*p*-coumaric acid (9).

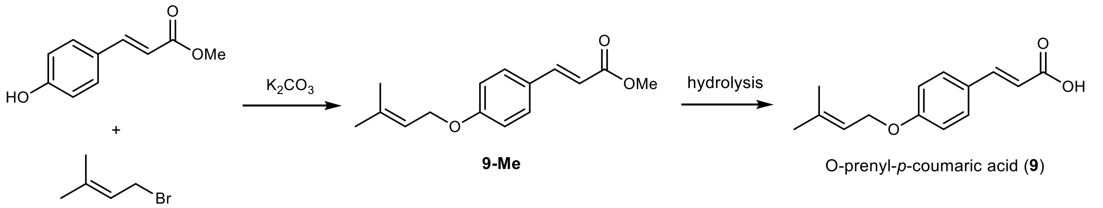


Compound **9** was synthesized following the same procedure used for compound **8**, except that a *p*-coumaric acid ester was employed as the substrate in place of the 4-hydroxybenzoic acid ester. The NMR data of synthetic **9-Me:** $^1$H NMR (500 MHz, $CDCl_3$) δ 7.64 (d, $J$ = 15.9 Hz, 1H), 7.49 – 7.40 (m, 2H), 6.90 (d, $J$ = 8.8 Hz, 2H), 6.30 (d, $J$ = 15.9 Hz, 1H), 5.47 (tp, $J$ = 6.6, 1.6 Hz, 1H), 4.52 (d, $J$ = 6.8 Hz, 2H), 3.78 (s, 3H), 1.84 – 1.78 (m, 3H), 1.74 (d, $J$ = 1.3 Hz, 3H). $^{13}$C NMR (126 MHz, $CDCl_3$) δ 18.3, 25.9, 51.6, 65.0, 115.1, 115.2, 119.3, 127.1, 129.8, 138.7, 144.7, 160.8, 167.9.

#### 5.8 Chemical synthesis of O-homoallenyl-4-hydroxybenzoic acid (10).

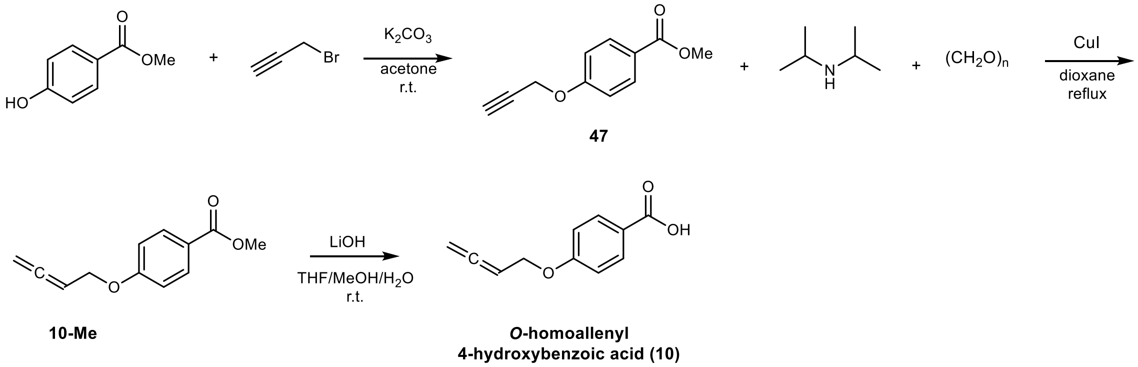


A suspension of 4-hydroxybenzoic acid ester (0.61 g, 4.0 mmol, 1.0 equiv) and $K_2CO_3$ (0.53 g, 4.8 mmol, 1.2 equiv) in 30 mL of acetone was stirred at room temperature, and propargyl bromide (0.52 mL, 4.8 mmol, 1.2 equiv) was added dropwise. The mixture was stirred at r.t. for 18 h, and then poured into 150 mL of saturated $NaHCO_3$ aqueous solution. The mixture was extracted by 3 x 50 mL of EtOAc, and the combined organic layer was washed with brined, dried using anhydrous $Na_2SO_4$, and concentrated under reduced pressure to give 0.70g of crude **10-Me**.

The crude residue **47** was dissolved in 30 mL of 1,4-dioxane, and CuI (0.15 g, 0.80 mmol, 0.2 equiv) diisopropylamine (1.1 mL, 8.0 mmol, 2.0 equiv), and paraformaldehyde (0.31 g, 10.0 mmol, 2.5 equiv) were added. The mixture was heated to reflux and stirred overnight. After cooling to r.t., the mixture was filtered through Celite to remove precipitates and washed with EtOAc. The combined filtrates were concentrated under reduced pressure to afford crude mixture. The mixture was further purified by reverse-phase column using Combi-Flash instrument with a gradient of 5-95% (acetonitrile : water) supplemented with 0.1% formic acid to give 0.68 g of **10-Me** (83% yield, 2 steps) as brown oil. The NMR data of synthetic **10-Me** matched the reported values[5]. $^1$H NMR (500 MHz, $CDCl_3$) δ 7.92 (d, $J$ = 8.9 Hz, 1H), 6.85 (d, $J$ = 8.9 Hz, 1H), 5.33 (p, $J$ = 6.7 Hz, 0H), 4.83 (dt, $J$ = 6.8, 2.5 Hz, 1H), 4.52 (dt, $J$ = 6.6, 2.6 Hz, 1H), 3.81 (s, 1H). $^{13}$C NMR (126 MHz, $CDCl_3$) δ 51.5, 65.6, 76.6, 86.5, 114.2, 122.5, 131.3, 161.9, 166.4, 209.3.

To a solution of **10-Me** (0.68 g, 3.3 mmol) in THF/MeOH/$H_2O$ (4:2:3 (v/v/v), 18 mL), added LiOH (0.16 g, 6.6 mmol). The reation mixture was allowed to stir at room temperature. After reaction completion, adjust the pH to 3 and extracted with 3 x 30 mL of EtOAc. The extracts were washed with water and saturated brine, and then dried over anhydrous $MgSO_4$ to give 0.63 g of **10** with quantitative yield.

#### 5.9 Chemical synthesis of O-homoallenyl-*p*-coumaric acid (11).

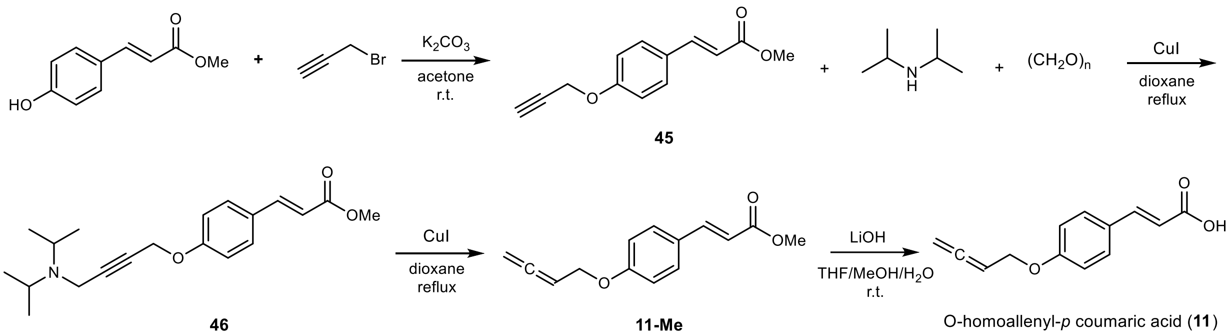


The synthesis of **11** followed the same procedure as for compound **10**, except that a *p*-coumaric acid ester was used in place of the 4-hydroxybenzoic acid ester. The NMR data of synthetic **11-Me** matched the reported values[5]. **11-Me:** $^1$H NMR (500 MHz, $CDCl_3$) δ 7.64 (d, $J$ = 15.9 Hz, 1H), 7.46 (d, $J$ = 8.4 Hz, 2H), 6.91 (d, $J$ = 8.3 Hz, 2H), 6.31 (d, $J$ = 15.9 Hz, 1H), 5.38 (p, $J$ = 6.7 Hz, 1H), 4.87 (dd, $J$ = 6.3, 2.9 Hz, 2H), 4.59 (dd, $J$ = 6.5, 3.0 Hz, 2H), 3.79 (s, 3H). $^{13}$C NMR (126 MHz, $CDCl_3$) δ 51.7, 66.0, 86.9, 115.3, 115.5, 127.4, 129.8, 144.6, 160.3, 167.9, 209.7.

#### 5.10 Chemical synthesis of O-2-butyn-1-ylbenzoic acid (16).

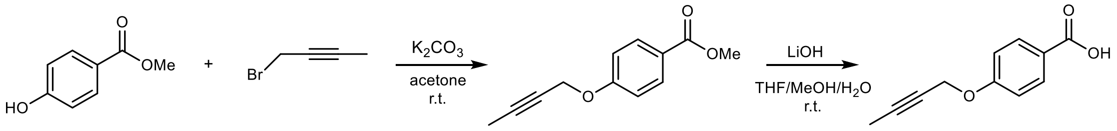


Compound **16** was synthesized following the same procedure used for compound **8**, except that a 1-bromo-2-butyne (0.36 mL, 4.06 mmol) was employed as the substrate in place of the prenyl bromide. The NMR data of synthetic **12:** $^1$H NMR (500 MHz, DMSO-$d_6$) δ 7.89 (d, $J$ = 9.0 Hz, 2H), 7.04 (d, $J$ = 9.0 Hz, 2H), 4.82 (s, 1H), 1.83 (d, $J$ = 2.3 Hz, 3H). $^{13}$C NMR (126 MHz, DMSO-$d_6$) δ 3.1, 56.1, 74.3, 84.0, 114.6, 123.4, 131.2, 160.9, 166.9.

**5.11 Chemical synthesis of 19-Me.**

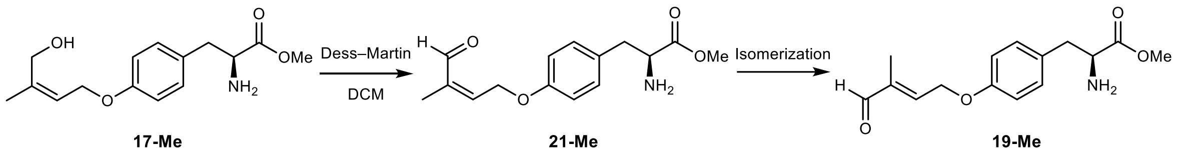


To a stirred solution of **17-Me** (20 mg, 0.072 mmol, 1.0 eq.) in DCM was added Dess-Martin periodinane (36 mg, 0.086 mmol, 1.2 eq.) at 0°C. The reaction mixture was stirred at 0°C for 0.5 hour, then filtered through Celite and the precipitate was further washed with DCM. The filtrate was successively washed with a saturated solution of $NaHCO_3$ and brine. The organic layer was dried over $Na_2SO_4$, filtered, and concentrated under reduced pressure. The crude residue was further purified by reverse-phase column using HPLC system with a linear gradient of 5-50% acetonitrile: water supplemented with 0.1% formic acid to afford 5.0 mg of **19-Me** as a yellow oil (25% yield). We were unable to obtain the corresponding $C_4$-aldehyde (**21-Me**) likely due to the spontaneous *E*/*Z* isomerization during the reaction and purification.

**5.12 Chemical synthesis of 20.**

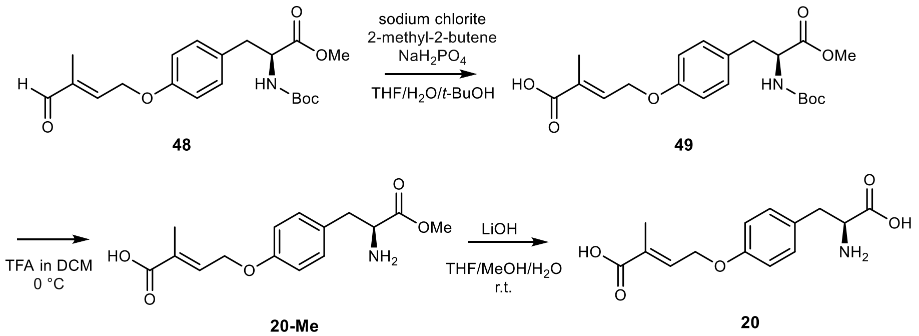


To a stirred solution of the **48** (0.39 g, 1.0 mmol) in tetrahydrofuran/water/*tert*-butanol (22.5 mL, 4:4:1) were added 2-methyl-2-butene (1.60 mL, 15.0 mmol) at 0 ºC, followed by the sequential addition of sodium dihydrogen phosphate monohydrate (0.72 g, 6.0 mmol) and sodium chlorite (80% purity, 0.34 g, 3.0 mmol). The resulting reaction mixture was stirred at r.t. overnight. The reaction mixture was diluted with brine and extracted with 3 x 30 mL of EtOAc. The combined organic layers were dried over $Na_2SO_4$, filtered, and concentrated reduced pressure to give 0.40 g of the crude compound **49**, which was directly used in the next step without further purification.

To a stirred solution of **49** (70 mg, 0.18 mmol) in 10 mL of DCM were added 3 mL of TFA at 0 ºC. The resulting reaction mixture was stirred at r.t. until all substrate is deprotected (checked by LC/MS). The reaction mixture was concentrated under reduced pressure to afford **20-Me**, which was subsequently subjected to hydrolysis with lithium hydroxide. The resultant residue was purified by reverse-phase column using Combi-Flash instrument with a gradient of 5-95% (acetonitrile : water) supplemented with 0.1% formic acid to give 25 mg of **20** as pale yellow oil.

**5.13 Chemical synthesis of 7-Me and 7.**

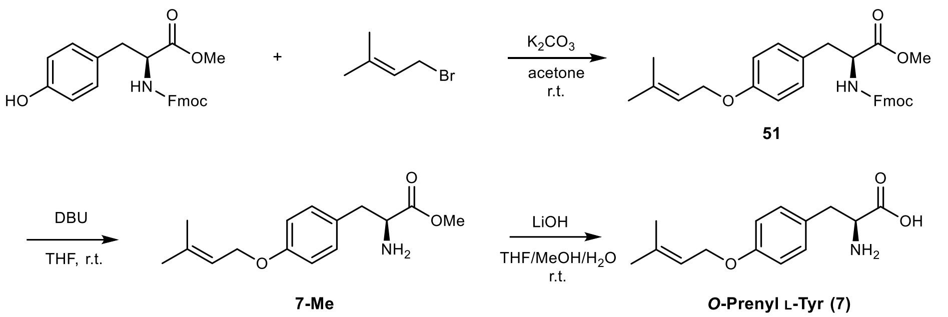


Compound **51** (120 mg, 0.20 mmol) was synthesized as same method as **35**, except that Fmoc-L-Tyr-OMe was employed as the substrate instead of Boc-L-Tyr-OMe. The crude **51** was dissolved in THF, followed by dropwise addition of DBU (45 μL, 0.30 mmol, 1.5 equiv) at room temperature. The reaction mixture was stirred for 2 h. After completion, the reaction was quenched with saturated aqueous $NH_4Cl$ (5 mL) and extracted with EtOAc (3 × 10 mL). The combined organic layers were dried over anhydrous $Na_2SO_4$, filtered, and concentrated under reduced pressure. The crude residue was purified by silica gel column chromatography (hexanes/EtOAc 3:1) to yield methyl ester **7** as a white solid (85 mg). Further hydrolysis of **7-Me** was carried out by dissolving the purified ester in THF/MeOH/$H_2O$ (1:1), followed by addition of LiOH (2.0 equiv). The mixture was stirred at room temperature for 2 h. Upon completion, the reaction was lyophilized to yield **7** as a white solid.

#### 5.14 Chemical synthesis of 1',1'-[$d_2$]-7 and 2'-[$d_1$]-7.

Regarding the synthesis of 1',1'-[$d_2$]-**7**, the same synthetic scheme from 32 to 17 in **Chemical synthesis of 17-Me and 17** without TBAF step was applied with 1,1-[$d_2$]-prenol as the starting material. Similarly, regarding the synthesis of 2'-[$d_1$]-**7**, the same synthetic scheme was used with 2-[$d_2$]-prenol as the starting material. 1,1-[$d_2$]-prenol[6] and 2-[$d_2$]-prenol[7] were synthesized by the reported procedures.

### 6. Sequence information

PpnA amino acid sequence

MAQTLLDESPEFQFGRYQFASNGLSTSSADLEPREHDRRITRYPYYPVELSIWQRVNSELDSFESVHHRFW
WSRHTGKALAVLLYNAQYPADLQYWNLKFFAEAVAPHLGVAPEILGSDTPIWPSFMTDDGTPVELSWDW
GTKDAPPMVRYSVEPIGLHAGTSVDPGNLTAGPAFQERLTRSLPTMRLEWFHHFKDFFNIPNAKEGEFHE
DTRDHNSSIFYGFDCSETEITPKVYFFPKLRAKASGQSNLDVLFQAMRTAPHVTDRNFEAGDIFHAFCSSV
GSKSLEHEMLAIDLIDPLQSRLKIYFRSRETTFQSVINIMTLEGRIRNPKLYEGLVDLHRLWTALFGVYAVD
QPLREVEHRTSGILYNFEFRLGEALPVAKIYLPVRHYCTSDEAVIRALNDYFQGQQKGKYMPDYVRAMST
LFTPKSMRENSGVQTYVGCAIRPDGTLRVVSYFKPQVPVHLFESEVYL

PpnB amino acid sequence

MLESIQQALFPVGRSLIVDYIATGLTLFLVYLIRTRYFHGLNKIPGPFLASVTSLWKWDTVRREKMPFVNT
QLHERYGPLVRIGPNHISASSAESIQVVHRSKSGFTKSGIYGILQPSYEGTDLHNVFSTQDAEYHAALKRT
MGSLYTTTAVLGLEFHLDDCTKLFISKMNEIIGTKASAPVDLAAWLQYYAFDSLGAVNFSEMLGFMESGT
DVDGICHLDHDQMMYFAVWGQITTIERAWAKVKALATGTRKENPLFNFALELVQKREANPTDSTDMLN
LFLALHKSVPEKFTIRDVIAAAYINVVTAHDVVAITLRAVVYYLAKNPAMQKKLQQEIDDAETAGKLSNP

AKYTEITPLSYVLAVTNEALRIHPSTGLILERRCPTGGVTLHGQYIPEGTIIGVNCWVVNRDKGIFGHDAH
EFRPERWIDSDPKDVSRMRTNMFTFGAGARNCIGKNLAMMQLTKIIVELYRNFDIVLENPDKDWTVSGG
WLTRQTEMDMVLTKRK

PpnC amino acid sequence

MNHSDIVLREWECLETLICGKNSAKVVLTGRTLSIPDVIAVARYNVLSDIDLSAIKAMEMSLSILEKRLHSG
DVIYGVNTGFGGSADVRTKKHVELQRALIRELHYGILPTGERDHRPLQLEGNLGQHRHDLSLESGLESTY
LPWSWARASILIRINSLISGCSAVRPVIVQRMQDLLKHDIIPMIPLRGSISSSGDLSPLSYICGAIQGKSTLRV
LSRDNKHVYADSAFERAGLEPVVIQAKEGLAVTNGTTVSAAVAALALHDTHSLALLAQVLTAMSVEALA
GTSESFHPFFSEVRPHPGQMESARNILNFLSGSRLTKVNDGAHSSLRQDRYSIRTAPQWLGPILEDLSLAHQ
QVFIECNSATDNPLVTPEGEFIHGGNFQAKSVTSAMEKARQGIQGIGRMLFSQCTEIINPATSRGLPPNLVA
EDPGISLIFKGFDLNIASLAAELGFLASPVNHVQTAEMGNQSLNSLALISARYTHTANEVLSQLMAAHLIA
VCQALDLRAMHAQFLEGYHPEFVQLVDAHYAEETLPSDGIPKIPNPIEVSGNLTNGLADPQLKPVEPKTV
ETLSELLWAQLLAAFDTTTSMDAEQRFPVMAKSLRSVLLDHVGFNTAADFTPRLQSFTQALSASLHDAW
CAHRDAYLVHGDATGLLGKASRAMYVFLRRSLGVPLLATRNLATPTVGGMNGESGTHGGVEAPTVGSY
TGAVYRALRDGTLAKVAVDILRDSVKAEVEVGE

PpnD amino acid sequence

MDESLILVALCALCLAFQLLRTRYHRGLNAIPGPFSASFCNLWKILAVYNNDMPRRNISVHKKYGPVVRIG
PKHVSFSSPEALHIIHGSRQAYPKSDFYNPAAAPFEGSPLLNLFSVRDVSYHSSLKKVIGGLYTKAAVLDLE
SKIDTCVEMFTNQLRKRTQDDGPTNVDMSLWVHLFAFDCLGELNVSKKFGYLDTGRDFNGFIEGSDKVL
IKTGLFGQAPFLQVIRKLIDTRWGAEKLNPVLKYTTAVVRQRLEKPTETPDMLNSFLALRKAQPEKLSIRD
ITGSIYINLMAGHDVLAVTLRTILYYVARSPRVEDKLRGELATIITHYRPTDAIPYTETSKLPYLGAVINESLR
IHGNLGLINERVTPPEGARIDGYHIPGGTIVGINPWVIHRNTEIFGEDVETFRPERWLDGPEDSIQEMKRNL
FSFGAGPRMCIGKNIAMMQIYKFITQFYRHFTFELASPEKDWHVIGNWVTKQTEMDMLVTQAKPRQI

NseB amino acid sequence

MKYPTTGQLQQVHLGIGPKGYEPVASYQGDKQLYTQEHEILQASILGFCPEHLWHHGSNKASCPRPILVT
AKHQEQLEQLHNALVTAIVDIVKRWWTDLDARFPERMPLTRDEEDLLRWLEHQHSHNGVPYEARLGSW
RPDFLVGDYSGGPSTETYRLTEINARFCFNGFMHQAYGQEGLSDLGAGRNGLIHATDSSKILDGLLSLFNP
DRPLHLLKGEEPGIDIHMFIDFVYRHIGIKPRLITPADLRLIPDPQKKDGSKLCCLVKDQQNASLINESRLLV
TSKGEVVEEVHQVGLELHQHELFGLSREMLREISLRCFNDMRTILLVHDKRMLGIIKQEMPTLVARKVLT
HDQGEALERGISDSFIPGSSELNELIQTLTDSPELRKEYLLKPIRGGKGAGIIFGDEVGPDEWLSTLERLRNP
HFVSGNTMYVVQRRIWPRLYEVILNSSGDRGNYPLIGTYHTTNGQLLGLGTWRSSPDRICAVSHGGGWIC
SVLDEYAESSE

**Supplementary Tables**

**Supplementary Table 1.** Comparative BLASTP analysis of the *ppn* gene cluster with homologous cluster from *Penicillium polonicum*, *Nemania serpens*, and *Xylaria telfairii*.

| *Penicillium polonicum (ppn)* | *Xylaria telfairii* (% identity) *(xte1)* | *Xylaria telfairii* (% identity) *(xte2)* | *Nemania serpens* (% identity) *(nse)* | Homolog in Swiss-Prot database (% identity) |
|---|---|---|---|---|
| PpnA (PT) OQD61507.1 | Xte1A (51%) KAI0446169.1 | Xte2A (68%) KAI0441401.1 | NseA (PT) (66%) KAI1163401.1 | 4-O-dimethylallyl-L-tyrosine synthase, M1VV66.1 (34%) |
| PpnB (P450) OQD61637.1 | Xte1B (79%) KAI0446171.1 | Xte2B (68%) KAI0441400.1 | NseB (P450) (69%) KAI1163402.1 | Cytochrome P450 monooxygenase, D7PI20.2 (GsfF, 33%) |
| PpnC (TAL) OQD61545.1 | Xte1C (65%) KAI0446168.1 | Xte2C (AT) KAI0441399.1 | NseC (AT) (72%) KAI1163403.1 | Phenylalanine ammonia-lyase, A0AAN4PAE6.1 (LenB, 40%) |
| PpnD (P450) OQD61524.1 | Xte1D (70%) KAI0446170.1 | N/A | N/A | Cytochrome P450 monooxygenase, C8V0D4.1 (CicH, 33%) |
| PpnE (MT) OQD61540.1 | Xte1E (60%) KAI0446175.1 | N/A | N/A | Methyltransferase, A0A6F8RNE2.1, (GrgD, 28%) |
| PpnF (BBE) OQD61464.1 | Xte1F (65%) KAI0446173.1 | N/A | N/A | VAO-type flavoprotein oxidase VAO615, G2QDQ9.1, (46%) |
| PpnG (SDR) OQD61477.1 | Xte1G (65%) KAI0446172.1 | N/A | N/A | Ketoreductase CTB6, A0A2G5ICG8.1, (37%) |

**Supplementary Table 2.** Plasmids used in this study

| Plasmids | Vector | Genes |
|---|---|---|
| pML 9001 | pYTU | *ppnA* (dimethylallyltryptophan synthase) |
| pML 9002 | pYTP | *ppnB* (cytochrome P450) |
| pML 9003 | pYTU | *ppnC* (phenylalanine ammonia lyase) |
| pML 9004 | pYTR | *ppnD* (cytochrome P450)-*ppnE* (methyltransferase) |
| pML 9005 | pYTU | *ppnA* (dimethylallyltryptophan synthase)-*ppnB* (cytochrome P450)-*ppnC* (phenylalanine ammonia lyase) |
| pML 9006 | pYTR | *ppnD* (cytochrome P450) |
| pML 9007 | pYTP | *ppnF* (BBE-like)-*ppnG* (SDR) |
| pML 9008 | pYTP | *ppnA* (dimethylallyltryptophan synthase) |
| pML 9009 | pYTU | *nseA* (dimethylallyltryptophan synthase) |
| pML 9010 | pYTU | *nseA* (dimethylallyltryptophan synthase)-*nseB* (cytochrome P450) |
| pML 9011 | pYTR | *nseB* (cytochrome P450) |
| pML 9012 | pYTU | *nseC* (acyltransferase) |
| pML 9013 | pYTU | *ppnA* (dimethylallyltryptophan synthase)-*pfaA* (PKS-NRPS) |

**Supplementary Table 3.** Primers used in this study

| Primers | Sequence (5'-3') |
|---|---|
| pML 9001 F1 | TAACCTCGCGGGTGTTCTTGACGATGGCATCCTGCACTCCGGTGAATTGATTTGGGTGAC |
| pML 9001 R1 | AATTCGGGGCTTTCATCAAGTAAAGTCTGAGCCATTGTTTAGATGTGTCTATGTGGCGGG |
| pML 9001 F2 | ACCATTACCCCGCCACATAGACACATCTAAACAATGGCTCAGACTTTACTTGATGAAAGC |
| pML 9001 R2 | ACAGTGGAGGACATACCCGTAATTTTCTGATTTAAATCACATTCCTTCAGCTCTATGCCC |
| pML9002 F1 | TTAGTAACCTCGCGGGTGTTCTTGACGATGGCATCCTACTCCGGTGAATTGATTTGGGTG |
| pML9002 R1 | ACGGGGAAGAGAGCTTGTTGTATCGATTCAAGCATTGTTTAGATGTGTCTATGTGGCGGG |
| pML9002 F2 | ACCATTACCCCGCCACATAGACACATCTAAACAATGCTTGAATCGATACAACAAGCTCTC |
| pML9002 F3 | TGGACATGGTTTTGACGAAGAGGAAGTAACCACTTAACGTTACTGAAATCATCAAACAGC |
| pML9002 R2 | AGCTGTTTGATGATTTCAGTAACGTTAAGTGGTTACTTCCTCTTCGTCAAAACCATGTCC |
| pML9002 R3 | ATGAGACCCAACAACCATGATACCAGGGGAAGAAGGATTACCTCTAAACAAGTGTACCTG |
| pML9003 F1 | AGTAACCTCGCGGGTGTTCTTGACGATGGCATCCTGCACTCCGGTGAATTGATTTGGGTG |
| pML9003 R1 | TCCCACTCCCTCAGAACAATATCCGAGTGATTCATTGTTTAGATGTGTCTATGTGGCGGG |
| pML9003 R2 | ACACAGTGGAGGACATACCCGTAATTTTCTGATTTAAATGGCGCTATGTACGCATTCGAC |
| pML9003 F2 | CCATTACCCCGCCACATAGACACATCTAAACAATGAATCACTCGGATATTGTTCTGAGGG |
| pML9004 F1 | TACCCCGCCACATAGACACATCTAAACAATGGATGAGTCTTTAATATTGGTTGCACTTTG |
| pML9004 F2 | CAAGCCCCCCCTACCTTCCAGCTCGCTCGAAACAGATTTGATTAAAGGTGCCGAACGAGC |
| pML9004 F3 | CCCTTCTCTGAACAATAAACCCCACAGAAGGCATTTATGTCAGAAGTCAAGAATGCCCTC |
| pML9004 R2 | AGGATATTCCTCGAGGGCATTCTTGACTTCTGACATAAATGCCTTCTGTGGGGTTTATTG |
| pML9004 R3 | GCTAAAGGGTATCATCGAAAGGGAGTCATCCAATTTTGCCGTGGATATTCTTAGAGACTC |
| pML9004 R1 | TTATATCATTTATAGCTCGTTCGGCACCTTTAATCAAATCTGTTTCGAGCGAGCTGGAAG |
| pML9005 F1 | CCTCGCGGGTGTTCTTGACGATGGCATCCTGCACTCCGGTGAATTGATTTGGGTG |
| pML9005 F2 | ACCATTACCCCGCCACATAGACACATCTAAACAATGAATCACTCGGATATTGTTCTGAGG |
| pML9005 F3 | CCGACCACGACCACCCTGTCGAATGCGTACATAGCGCCGATTCGTCCAGGGCTTCCCAAG |
| pML9005 F4 | ACATGATCTAACAACTTCTAGTAAACCGCAATCATGGCTCAGACTTTACTTGATGAAAGC |
| pML9005 F5 | TTTCTATTAGGGCATAGAGCTGAAGGAATGTGTTTGCTCCAGGAATACATGTGAGCTTAC |
| pML9005 F6 | AGTGCATACAGAACACTTCAAACAATCGCAAAAATGCTTGAATCGATACAACAAGCTCTC |
| pML9005 R1 | TCCCACTCCCTCAGAACAATATCCGAGTGATTCATTGTTTAGATGTGTCTATGTGGCGGG |
| pML9005 R2 | CCAATATTCCAACCTTGGGAAGCCCTGGACGAATCGGCGCTATGTACGCATTCGACAGG |
| pML9005 R3 | GGGGCTTTCATCAAGTAAAGTCTGAGCCATGATTGCGGTTTACTAGAAGTTGTTAGATCA |
| pML9005 R4 | AGAATCAGTAAGCTCACATGTATTCCTGGAGCAAACACATTCCTTCAGCTCTATGCCCT |
| pML9005 R5 | GAAGAGAGCTTGTTGTATCGATTCAAGCATTTTTGCGATTGTTTGAAGTGTTCTGTATGC |
| pML9005 R6 | AACACAGTGGAGGACATACCCGTAATTTTCTGATTTGACCCCGTGGTTCGGGTACCATAC |
| pML9006 F1 | TACCCCGCCACATAGACACATCTAAACAATGGATGAGTCTTTAATATTGGTTGCACTTTG |
| pML9006 R1 | CTAAAGGGTATCATCGAAAGGGAGTCATCCAATTTAAATCTGTTTCGAGCGAGCTGGAAG |
| pML9007 F1 | TCCCTTCTCTGAACAATAAACCCCACAGAAGGCATTTATGTCACCCTACTGGCAGAATCG |
| pML9007 F2 | AGTAAGTAAGCAAGAGCGTTACGTATACCCTCTGGCATTTGATTCGTCCAGGGCTTCCCA |
| pML9007 F3 | TATACATGATCTAACAACTTCTAGTAAACCGCAATCATGTGTTCCGCAGACCTTCTCATC |
| pML9007 R1 | CCCAATATTCCAACCTTGGGAAGCCCTGGACGAATCAAATGCCAGAGGGTATACGTAACG |
| pML9007 R2 | CGGTGATGAGAAGGTCTGCGGAACACATGATTGCGGTTTACTAGAAGTTGTTAGATCATG |
| pML9007 R3 | GAGTGATGAGACCCAACAACCATGATACCAGGGGATTTCAGCAGTGTCTCTCGTGGTCTT |
| pML9008 F1 | AGTAACCTCGCGGGTGTTCTTGACGATGGCATCCTACTCCGGTGAATTGATTTGGGTG |
| pML9008 F2 | ACCATTACCCCGCCACATAGACACATCTAAACAATGGCTCAGACTTTACTTGATGAAAGC |
| pML9008 F3 | ACTTGTTCGAATCTGAGGTATATCTCTGACCACTTAACGTTACTGAAATCATCAAACAGC |
| pML9008 R1 | TGTTTAGATGTGTCTATGTGGCGGG |
| pML9008 R2 | GTTTGATGATTTCAGTAACGTTAAGTGGTCAGAGATATACCTCAGATTCGAACAAGTGTA |
| pML9008 R3 | ATGAGACCCAACAACCATGATACCAGGGGAAGAAGGATTACCTCTAAACAAGTGTACCTG |
| pML9009 F1 | CCTGAGCTTCATCCCCAGCATCATTACACCTCAGCAATGTTCTCTTCGTTCGGTGACTTG |
| pML9009 R1 | CAACACAGTGGAGGACATACCCGTAATTTCTGGCTGAGGTAGTTCCTGATTGTTTCG |
| pML9010 F1 | CCTGAGCTTCATCCCCAGCATCATTACACCTCAGCAATGTTCTCTTCGTTCGGTGACTTG |
| pML9010 F2 | GGGATGAATGAATCGAAACAATCAGGAACTACCTCAGCGATTCGTCCAGGGCTTCCCAAG |
| pML9010 F3 | CTTATACATGATCTAACAACTTCTAGTAAACCGCAATCATGGAGTCCACACAGCACATCG |

**Supplementary Table 3 (continued).** Primers used in this study.

| Primers | Sequence (5'-3') |
|---|---|
| pML9010 R1 | CCAATATTCCAACCTTGGGAAGCCCTGGACGAATCGCTGAGGTAGTTCCTGATTGTTTCG |
| pML9010 R2 | GATTGCGGTTTACTAGAAGTTGTTAGATCATGTA |
| pML9010 R3 | CAACACAGTGGAGGACATACCCGTAATTTTCTGGGCGCAATCCGTGCCGCAATGTC |
| pML9011 F1 | GACTAACCATTACCCCGCCACATAGACACATCTAAACAATGGAGTCCACACAGCACATC |
| pML9011 F2 | GCAAGAGGGGAGCCTTGCTTGGAGCATAACCACTTAACGTTACTGAAATCATCAAACAGC |
| pML9011 R1 | CGTCAAGCTGTTTGATGATTTCAGTAACGTTAAGTGGTTATGCTCCAAGCAAGGCTCCC |
| pML9011 R2 | AAAGGGTATCATCGAAAGGGAGTCATCCAAAGAAGGATTACCTCTAAACAAGTGTACCTG |
| pML9012 F1 | CCTGAGCTTCATCCCCAGCATCATTACACCTCAGCAATGGTTAGCCTGGGCCTCGCTAAG |
| pML9012 F2 | CAGGGGACAAGCAGGCTGCTAAATTGTGACCACTTAACGTTACTGAAATCATCAAACAGC |
| pML9012 R1 | CTGTTTGATGATTTCAGTAACGTTAAGTGGTCACAATTTAGCAGCCTGCTTGTCCCCTGC |
| pML9012 R2 | ACAGTGGAGGACATACCCGTAATTTTCTGAAGAAGGATTACCTCTAAACAAGTGTACCTG |
| pML9013 F1 | GTAACCTCGCGGGTGTTCTTGACGATGGCATCCTGCACTCCGGTGAATTGATTTGGGTG |
| pML9013 F2 | AACCATTACCCCGCCACATAGACACATCTAAACAATGGCTCAGACTTTACATGGAGAAAG |
| pML9013 F3 | AACCTTCACCCGACAATGCAGTATCTGCATTTTTTGCTCCAGGAATACATGTGAGCTTAC |
| pML9013 F4 | CAAGTGCATACAGAACACTTCAAACAATCGCAAAAATGTCGCAGCGACAGCCTCTAGC |
| pML9013 F5 | GGACGAGATTACGCCCGACAG |
| pML9013 F6 | CAGGTCTACTCGAAGCGCTGAAGGAATAGCCACTTAACGTTACTGAAATCATCAAACAGC |
| pML9013 R1 | CGGGGCTTTCTCCATGTAAAGTCTGAGCCATTGTTTAGATGTGTCTATGTGGCGGG |
| pML9013 R2 | TAGAATCAGTAAGCTCACATGTATTCCTGGAGCAAAAAATGCAGATACTGCATTGTCGGG |
| pML9013 R3 | AACGATAGCTAGAGGCTGTCGCTGCGACATTTTTGCGATTGTTTGAAGTGTTCTGTATGC |
| pML9013 R4 | GGCTATCGAGTCCGTAGTCAAGTAAAC |
| pML9013 R5 | TGTTTGATGATTTCAGTAACGTTAAGTGGCTATTCCTTCAGCGCTTCGAGTAGACCTGCC |
| pML9013 R6 | TGGAGGACATACCCGTAATTTTCTGATTTAAGAAGGATTACCTCTAAACAAGTGTACCTG |

**Supplementary Table 4.** Spectroscopic data of compound *O*-homoallenyl-L-Tyr (**1**)

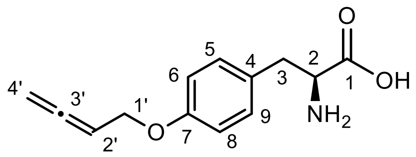


O-homoallenyl-L-Tyr (**1**)

| | *O*-homoallenyl-L-Tyr **(1)** in DMSO-$d_6$ | |
|---|---|---|
| Position | $\delta_H$ ($J$ in Hz) | $\delta_C$, type |
| 1 | | 175.4, C |
| 2 | 3.56, m | 55.8, CH |
| 3 | 2.70, m; 2.78, m | 36.6, $CH_2$ |
| 4 | | 130.2, C |
| 5 | 7.17, d (8.0) | 130.4, CH |
| 6 | 6.85, d (7.2) | 114.7, CH |
| 7 | | 156.7, C |
| 8 | 6.85, t (7.2) | 114.7, CH |
| 9 | 7.17, d (8.0) | 130.4, CH |
| 1' | 4.52, d (6.6) | 65.1, $CH_2$ |
| 2' | 5.47, m | 87.1, CH |
| 3' | | 208.7, C |
| 4' | 4.97, d (6.6) | 76.9, $CH_2$ |

NMR spectrum (500 MHz) for $^1$H, NMR spectrum (125 MHz) for $^{13}$C, DMSO-$d_6$, "m" means overlapped or multiple with other signals. Chemical shifts are reported in ppm.

HRMS (ESI, M+H$^+$) calculated for $C_{13}H_{16}NO_3^+$ 234.1125; found 234.1130.

$[\alpha]_D^{24.1}$ – 4° ($c$ 0.1, MeOH/$H_2O$, 70%, v/v).

**Supplementary Table 5.** Spectroscopic data of compound sinuxylamide B **(2)**

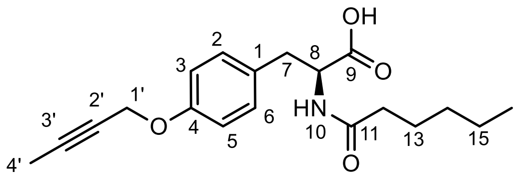


Sinuxylamide B (**2**)

| Position | Sinuxylamide B **(2)** in $CD_3OD$ $\delta_H$ (*J* in Hz) | $\delta_C$, type | Reported Sinuxylamide B **(2)** in $CD_3OD$[8] $\delta_C$, type |
|---|---|---|---|
| 1 | | 165.8, C | 174.5, C |
| 2 | 4.61, m | 55.8, CH | 54.3, CH |
| 3 | 2.88, m; 3.16, m | 36.6, $CH_2$ | 36.6, $CH_2$ |
| 4 | | 130.1, C | 129.8, C |
| 5 | 7.13, d (8.1) | 129.9, CH | 129.7, CH |
| 6 | 6.85, d (8.0) | 114.4, CH | 114.3, CH |
| 7 | | 156.9, C | 156.7, C |
| 8 | 6.85, d (8.0) | 114.4, CH | 114.3, CH |
| 9 | 7.13, d (8.1) | 129.9, CH | 129.7, CH |
| 10 | | 174.6, C | 174.4, C |
| 11 | 2.15, t (7.5) | 35.7, $CH_2$ | 35.3, $CH_2$ |
| 12 | 1.49, m | 25.3, $CH_2$ | 25.3, $CH_2$ |
| 13 | 1.18, m | 31.1, $CH_2$ | 31.1, $CH_2$ |
| 14 | 1.28, m | 22.1, $CH_2$ | 22.1, $CH_2$ |
| 15 | 0.87, t (7.3) | 13.0, $CH_3$ | 13.1, $CH_3$ |
| 1’ | 4.62, s | 55.8, $CH_2$ | 55.6, $CH_2$ |
| 2’ | | 74.0, C | 73.9, C |
| 3’ | | 82.6, C | 82.6, C |
| 4’ | 1.81, s | 1.8, $CH_3$ | 1.8, $CH_3$ |

NMR spectrum (500 MHz) for $^1H$, NMR spectrum (125 MHz) for $^{13}C$, $CD_3OD$, “m” means overlapped or multiple with other signals. Chemical shifts are reported in ppm.

HRMS (ESI, M+H$^+$) calculated for $C_{19}H_{26}NO_4^+$ 332.1856; found 332.1861.

$[\alpha]_D^{24.1}$ + 12° (*c* 0.1, MeOH). Compound **2** showed the same positive optical rotation as reported Sinuxylamide B[8].

**Supplementary Table 6.** Spectroscopic data of penipratynolene (**5**)

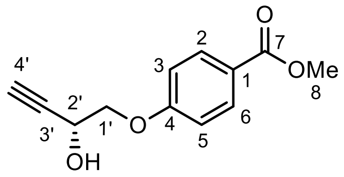


Penipratynolene (**5**)

| Position | penipratynolene (**5**) in acetone-*d*$_6$ $\delta_H$ (*J* in Hz) | $\delta_C$, type | Reported penipratynolene (**5**) in acetone-*d*$_6$[9] $\delta_C$, type |
|---|---|---|---|
| 1 | | 166.9, C | 166.7, C |
| 2 | | 123.7, C | 123.6, C |
| 3 | 7.96, d (8.9) | 132.2, CH | 132.1, CH |
| 4 | 7.07, d (8.9) | 115.3, CH | 115.1, CH |
| 5 | | 163.4, C | 163.2, C |
| 6 | 7.07, d (8.9) | 115.3, CH | 115.1, CH |
| 7 | 7.96, d (8.9) | 132.2, CH | 132.1, CH |
| 1’ | 4.18, m | 72.7, $CH_2$ | 72.6, $CH_2$ |
| 2’ | 4.74, s | 61.2, CH | 61.0, CH |
| 3’ | | 83.5, C | 83.3, C |
| 4’ | 2.98, m | 74.7, CH | 74.6, CH |
| OMe | 3.84 s | 52.0, $CH_3$ | 51.9, $CH_3$ |

NMR spectrum (500 MHz) for $^1$H, NMR spectrum (125 MHz) for $^{13}$C, acetone-*d*$_6$, “m” means overlapped or multiple with other signals. Chemical shifts are reported in ppm.

HRMS (ESI, M+H$^+$) calculated for $C_{12}H_{13}O_4^+$ 221.0808; found 221.0802.

$[\alpha]_D^{24.1}$ – 16° (*c* 0.1, $CDCl_3$). Compound **1** showed the same negative optical rotation as reported penipratynolene[9].

**Supplementary Table 7.** Spectroscopic data of compound *O*-2-butyn-1-yl L-Tyr (**6**)

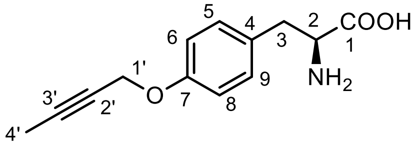


O-2-butyn-1-yl L-Tyr (**6**)

| Position | *O*-2-butyn-1-yl-L-tyrosine (**6**) in DMSO-$d_6$ (500MHz) $\delta_H$ (*J* in Hz) | $\delta_C$, type |
|---|---|---|
| 1 | | 169.2, C |
| 2 | 3.34, m | 55.6, CH |
| 3 | 2.78, m; 3.06, m | 36.1, $CH_2$ |
| 4 | | 129.9, C |
| 5 | 7.17, d (8.7) | 130.3, CH |
| 6 | 6.85, d (8.6) | 114.5, CH |
| 7 | | 156.2, C |
| 8 | 6.85, t (8.6) | 114.5, CH |
| 9 | 7.17, d (8.7) | 130.3, CH |
| 1’ | 4.69, q (2.4) | 55.7, $CH_2$ |
| 2’ | | 74.9, C |
| 3’ | | 83.3, C |
| 4’ | 1.82, t (2.3) | 3.2, $CH_3$ |

NMR spectrum (500 MHz) for $^1$H, NMR spectrum (125 MHz) for $^{13}$C, DMSO-$d_6$, “m” means overlapped or multiple with other signals. Chemical shifts are reported in ppm.

HRMS (ESI, M+H$^+$) calculated for $C_{13}H_{16}NO_3^+$ 234.1125; found 234.1132.

$[\alpha]_D^{24.1}$ – 32 (c 0.1, MeOH/$H_2O$, 70%, v/v)

**Supplementary Table 8.** Spectroscopic data of compound *O*-Prenyl L-Tyr (**7**)

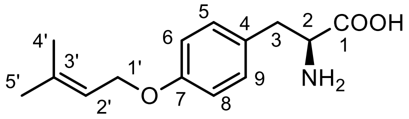


O-Prenyl L-Tyr (**7**)

| Position | *O*-Prenyl L-Tyr (**7**) in DMSO-$d_6$ | |
|---|---|---|
| | $\delta_H$ ($J$ in Hz) | $\delta_H$ ($J$ in Hz) |
| 1 | | 165.5, C |
| 2 | 3.26, m | 55.9, CH |
| 3 | 2.70, m; 2.78, m | 36.5, $CH_2$ |
| 4 | | 130.2, C |
| 5 | 7.17, d (8.0) | 130.3, CH |
| 6 | 6.85, d (7.2) | 114.4, CH |
| 7 | | 157.1, C |
| 8 | 6.85, t (7.2) | 114.4, CH |
| 9 | 7.17, d (8.0) | 130.3, CH |
| 1’ | 4.48 (d, 6.7) | 64.1, $CH_2$ |
| 2’ | 5.41, m | 120.2, CH |
| 3’ | | 136.8, C |
| 4’ | 1.69 (d, 1.4) | 18.0, $CH_3$ |
| 5’ | 1.73 (d, 1.4) | 25.4, $CH_3$ |

NMR spectrum (500 MHz) for $^1$H, NMR spectrum (125 MHz) for $^{13}$C, DMSO-$d_6$, “m” means overlapped or multiple with other signals. Chemical shifts are reported in ppm.

HRMS (ESI, M+H$^+$) calculated for $C_{14}H_{20}NO_3^+$ 250.1438; found 250.1434.

$[\alpha]_D^{24.1}$ – 22 (c 0.1, MeOH/$H_2O$, 70%, v/v)

Compound **3** showed the same positive optical rotation as reported *O*-Prenyl L-Tyr[10].

**Supplementary Table 9.** Spectroscopic data of *O*-prenyl-benzoic acid (**8**)

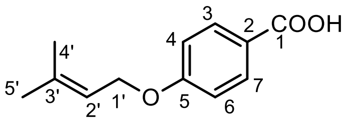


O-prenyl-benzoic acid (**8**)

| | *O*-prenyl-benzoic acid (**8**) in $CDCl_3$ | | Reported *O*-prenyl-benzoic acid (**8**) in $CDCl_3$[11] |
|---|---|---|---|
| Position | $\delta_H$ (*J* in Hz) | $\delta_C$, type | $\delta_H$ (*J* in Hz) |
| 1 | | 172.3, C | |
| 2 | | 121.7, C | |
| 3 | 8.06, d (8.9) | 131.5, CH | 8.09, d (6.8) |
| 4 | 6.95, d (8.9) | 114.5, CH | 6.98, d (6.8) |
| 5 | | 163.5, C | |
| 6 | 6.95, d (8.9) | 114.5, CH | 6.98, d (6.8) |
| 7 | 8.06, d (8.9) | 131.5, CH | 8.09, d (6.8) |
| 1’ | 4.58 (d, 6.3) | 65.2, $CH_2$ | 4.62, d (5.4) |
| 2’ | 5.49, m | 119.1, CH | 5.52, m |
| 3’ | | 139.1, C | |
| 4’ | 1.76, d (1.3) | 18.4, $CH_3$ | 1.79, s |
| 5’ | 1.81, d (1.3) | 26.0, $CH_3$ | 1.84, s |

NMR spectrum (500 MHz) for $^1H$, NMR spectrum (125 MHz) for $^{13}C$, $CDCl_3$, “m” means overlapped or multiple with other signals. Chemical shifts are reported in ppm.

HRMS (ESI, $M+H^+$) calculated for $C_{12}H_{15}O_3^+$ 207.1016; found 207.1006.

**Supplementary Table 10.** Spectroscopic data of *O*-prenyl *p*-coumaric acid (**9**)

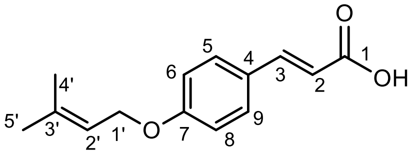

O-prenyl *p*-coumaric acid (**9**)

| Position | *O*-prenyl *p*-coumaric acid (**9**) in $CDCl_3$ $\delta_H$ (*J* in Hz) | $\delta_C$, type | Reported *O*-prenyl *p*-coumaric acid (**9**) in $CDCl_3$[12] $\delta_C$, type |
|---|---|---|---|
| 1 | | 173.1, C | 172.9, C |
| 2 | 6.32 (d, 15.9) | 114.7, CH | 114.2, CH |
| 3 | 7.75 (d, 15.8) | 146.9, CH | 146.8, CH |
| 4 | | 126.8, C | 126.7, C |
| 5 | 7.50, d (8.8) | 130.2, CH | 130.1, CH |
| 6 | 6.93, d (8.8) | 115.2, CH | 115.0, CH |
| 7 | | 161.2, C | 161.1, C |
| 8 | 6.93, d (8.8) | 115.2, CH | 115.0, CH |
| 9 | 7.50, d (8.8) | 130.2, CH | 130.1, CH |
| 1’ | 4.55 (d, 6.7) | 65.1, $CH_2$ | 64.1, $CH_2$ |
| 2’ | 5.49, m | 119.3, CH | 119.2, CH |
| 3’ | | 138.9, C | 138.8, C |
| 4’ | 1.75 (d, 1.4) | 18.4, $CH_3$ | 18.2, $CH_3$ |
| 5’ | 1.81 (d, 1.4) | 26.0, $CH_3$ | 25.8, $CH_3$ |

NMR spectrum (500 MHz) for $^1H$, NMR spectrum (125 MHz) for $^{13}C$, $CD_3CN$, “m” means overlapped or multiple with other signals. Chemical shifts are reported in ppm.

HRMS (ESI, $M+H^+$) calculated for $C_{14}H_{17}O_3^+$ 233.1172; found 233.1159.

**Supplementary Table 11.** Spectroscopic data of compound *O*-homoallenyl 4-hydroxybenzoic acid (**10**)

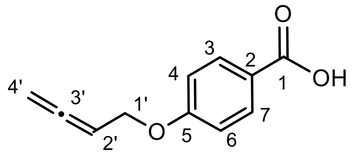


O-homoallenyl
4-hydroxylbenzoic acid (**10**)

| | *O*-homoallenyl 4-hydroxybenzoic acid (**10**) in $CDCl_3$ (500MHz) | |
|---|---|---|
| Position | $\delta_H$ (*J* in Hz) | $\delta_C$, type |
| 1 | | 171.5, C |
| 2 | | 121.9, C |
| 3 | 8.06, d (8.9) | 132.5, CH |
| 4 | 6.96, d (8.9) | 114.7, CH |
| 5 | | 163.0, C |
| 6 | 6.96, d (8.9) | 114.7, CH |
| 7 | 8.06, d (8.9) | 132.5, CH |
| 1’ | 4.64, dt (2.5, 6.8) | 66.1, $CH_2$ |
| 2’ | 5.40, m | 86.7, CH |
| 3’ | | 209.8, C |
| 4’ | 4.90, dt (2.5, 6.6) | 77.1, $CH_2$ |

NMR spectrum (500 MHz) for $^1$H, NMR spectrum (125 MHz) for $^{13}$C, $CDCl_3$, “m” means overlapped or multiple with other signals. Chemical shifts are reported in ppm.

HRMS (ESI, M+H$^+$) calculated for $C_{11}H_{11}O_3^+$ 191.0703; found 191.0696.

**Supplementary Table 12.** Spectroscopic data of *O*-homoallenyl *p*-coumaric acid (**11**)

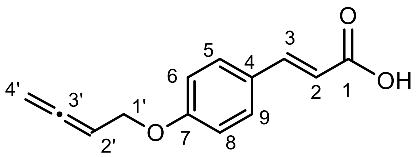


O-homoallenyl *p*-coumaric acid (**11**)

| Position | *O*-homoallenyl *p*-coumaric acid (**11**) in $CDCl_3$ $\delta_H$ (*J* in Hz) | $\delta_C$, type | Reported *O*-homoallenyl *p*-coumaric acid (**11**) in $CDCl_3$[5] $\delta_C$, type |
|---|---|---|---|
| 1 | | 172.8, C | 171.8, C |
| 2 | 6.32 (d, 15.7) | 114.9, CH | 114.7, CH |
| 3 | 7.75 (d, 15.7) | 146.8, CH | 146.7, CH |
| 4 | | 127.1, C | 127.0, C |
| 5 | 7.50, d (8.2) | 130.2, CH | 130.1, CH |
| 6 | 6.93, d (8.3) | 115.4, CH | 115.3, CH |
| 7 | | 160.6, C | 160.5, C |
| 8 | 6.93, d (8.3) | 115.4, CH | 115.3, CH |
| 9 | 7.50, d (8.2) | 130.2, CH | 130.1, CH |
| 1’ | 4.61, m | 66.1, $CH_2$ | 66.2, $CH_2$ |
| 2’ | 5.39, m | 86.8, CH | 86.9, CH |
| 3’ | | 209.7, C | 209.9, C |
| 4’ | 4.89, m | 77.0, $CH_2$ | 77.1, $CH_2$ |

NMR spectrum (500 MHz) for $^1H$, NMR spectrum (125 MHz) for $^{13}C$, $CDCl_3$, “m” means overlapped or multiple with other signals. Chemical shifts are reported in ppm.

HRMS (ESI, $M+H^+$) calculated for $C_{13}H_{13}O_3^+$ 217.0859; found 217.0864.

**Supplementary Table 13.** Spectroscopic data of compound **13**

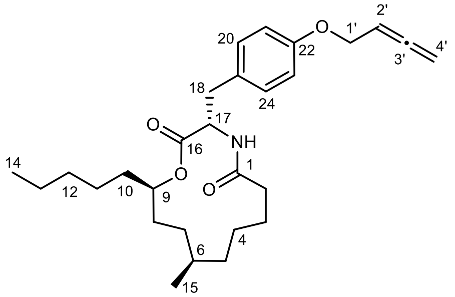


*Unnatural N*-demethylmelearoride A with allene tag (**13**)

| | **13** in $CDCl_3$ | |
|---|---|---|
| Position | $\delta_H$ (*J* in Hz) | $\delta_C$, type |
| 1’ | 4.52, dt (2.5, 6.9) | 66.0, $CH_2$ |
| 2’ | 5.37, m | 87.2, CH |
| 3’ | | 209.6, C |
| 4’ | 4.85, m | 76.6, $CH_2$ |
| 1 | | 173.0, C |
| 2 | 2.12, m; 2.24, m | 37.1, $CH_2$ |
| 3 | 1.48, m; 1.62, m | 25.4, $CH_2$ |
| 4 | 1.33, m | 23.7, $CH_2$ |
| 5 | 1.22, m | 34.0, $CH_2$ |
| 6 | 1.31, m | 29.7, CH |
| 7 | 1.07, m; 1.42, m | 27.4, $CH_2$ |
| 8 | 1.50, m | 28.8, $CH_2$ |
| 9 | 4.80, m | 75.8, CH |
| 10 | 1.38, m; 1.52, m | 33.6, $CH_2$ |
| 11 | 1.08, m | 25.1, $CH_2$ |
| 12 | 1.18, m | 31.7, $CH_2$ |
| 13 | 1.24, m | 22.6, $CH_2$ |
| 14 | 0.85, m | 14.2, $CH_3$ |
| 15 | 0.81, m | 20.6, $CH_3$ |
| 16 | | 172.1, C |
| 17 | 4.80, m | 53.8, CH |
| 18 | 3.04, m | 36.6, $CH_2$ |
| 19 | | 129.1, C |
| 20 | 7.13, d (8.6) | 130.4, CH |
| 21 | 6.82, d (8.6) | 115.0, CH |
| 22 | | 157.4, C |
| 23 | 6.82, d (8.6) | 115.0, CH |
| 24 | 7.13, d (8.6) | 130.4, CH |
| NH | 5.65, d (9.6) | |

NMR spectrum (500 MHz) for $^1H$, NMR spectrum (125 MHz) for $^{13}C$, $CDCl_3$, “m” means overlapped or multiple with other signals. Chemical shifts are reported in ppm.

HRMS (ESI, M+H$^+$) calculated for $C_{28}H_{42}NO_4^+$ 456.3108; found 456.3110.

$[\alpha]_D^{24.1}$ − 38° (*c* 0.1, MeOH).

**Supplementary Table 14.** Spectroscopic data of compound demethyl-**5** (**14**)

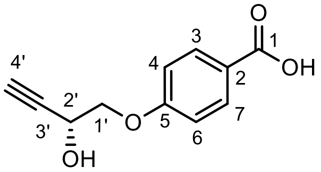


demethyl-**5** (**14**)

| Position | demethyl-**5** (**14**) in acetone-$d_6$ $\delta_H$ ($J$ in Hz) | $\delta_C$, type |
|---|---|---|
| 1 | | 167.6, C |
| 2 | | 124.0, C |
| 3 | 7.99, d (8.9) | 132.5, CH |
| 4 | 7.06, d (8.9) | 115.2, CH |
| 5 | | 163.3, C |
| 6 | 7.06, d (8.9) | 115.2, CH |
| 7 | 7.99, d (8.9) | 132.5, CH |
| 1' | 4.18, m | 72.7, $CH_2$ |
| 2' | 4.74, m | 61.2, CH |
| 3' | | 83.4, C |
| 4' | 2.98, m | 74.8, CH |

NMR spectrum (500 MHz) for $^1$H, NMR spectrum (125 MHz) for $^{13}$C, acetone-$d_6$, "m" means overlapped or multiple with other signals. Chemical shifts are reported in ppm.

HRMS (ESI, M+H$^+$) calculated for $C_{11}H_{11}O_4^+$ 207.0652; found 207.0651.

$[\alpha]_D^{24.1}$ – 10° ($c$ 0.1, MeOH).

**Supplementary Table 15.** Spectroscopic data of compound **15**

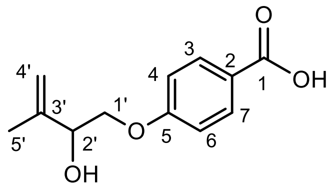


| Position | **15** in $CDCl_3$ $\delta_H$ ($J$ in Hz) | $\delta_C$, type |
|---|---|---|
| 1 | | 170.9, C |
| 2 | | 122.3, C |
| 3 | 8.05, d (8.6) | 132.5, CH |
| 4 | 6.97, d (8.6) | 114.5, CH |
| 5 | | 163.0, C |
| 6 | 6.97, d (8.6) | 114.5, CH |
| 7 | 8.05, d (8.6) | 132.5, CH |
| 1’ | 4.01, m; 4.12, m | 71.4, $CH_2$ |
| 2’ | 4.51, m | 73.7, CH |
| 3’ | | 143.2, C |
| 4’ | 5.04, s; 5.17, s | 113.3, $CH_2$ |
| 5’ | 1.84, s | 19.0, $CH_3$ |

NMR spectrum (500 MHz) for $^1H$, NMR spectrum (125 MHz) for $^{13}C$, $CDCl_3$, “m” means overlapped or multiple with other signals. Chemical shifts are reported in ppm.

HRMS (ESI, $M+H^+$) calculated for $C_{12}H_{15}O_4^+$ 223.0965; found 223.0962.

$[\alpha]_D^{24.1}$ + 44° ($c$ 0.1, MeOH).

**Supplementary Table 16.** Spectroscopic data of *O*-but-2-ynyl benzoic acid (**16**)

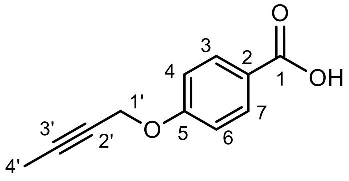

O-but-2-ynyl benzoic acid (**16**)

| | *O*-but-2-ynyl benzoic acid (**16**) in DMSO-$d_6$ | |
|---|---|---|
| Position | $\delta_H$ ($J$ in Hz) | $\delta_C$, type |
| 1 | | 166.9, C |
| 2 | | 123.4, C |
| 3 | 7.89, d (8.9) | 131.2, CH |
| 4 | 7.03, d (8.9) | 114.6, CH |
| 5 | | 160.9, C |
| 6 | 7.03, d (8.9) | 114.6, CH |
| 7 | 7.89, d (8.9) | 131.2, CH |
| 1' | 4.82, m | 56.1, $CH_2$ |
| 2' | | 74.3, C |
| 3' | | 84.0, C |
| 4' | 1.83, t (2.3) | 3.1, $CH_3$ |

NMR spectrum (500 MHz) for $^1$H, NMR spectrum (125 MHz) for $^{13}$C, DMSO-$d_6$, "m" means overlapped or multiple with other signals. Chemical shifts are reported in ppm.

HRMS (ESI, M+H$^+$) calculated for $C_{11}H_{11}O_3^+$ 191.0703; found 191.0691.

**Supplementary Table 17.** Spectroscopic data of compound **17** methyl ester (**17**-Me)

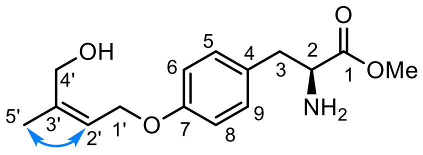


| Position | **17**-Me in $D_2O$ $\delta_H$ ($J$ in Hz) | $\delta_C$, type |
|---|---|---|
| 1 | | 170.0, C |
| 2 | 4.39, m | 54.1, CH |
| 3 | 3.19, m; 3.29, m | 34.7, $CH_2$ |
| 4 | | 126.4, C |
| 5 | 7.22, d (8.7) | 130.6, CH |
| 6 | 7.00, d (8.6) | 115.6, CH |
| 7 | | 157.3, C |
| 8 | 7.00, d (8.6) | 115.6, CH |
| 9 | 7.22, d (8.7) | 130.6, CH |
| 1’ | 4.65, d, 6.6) | 64.1, $CH_2$ |
| 2’ | 5.66, m | 122.0, CH |
| 3’ | | 141.0, C |
| 4’ | 4.18, s | 60.0, $CH_2$ |
| 5’ | 1.84 (d, 1.5) | 20.4, $CH_3$ |
| OMe | 3.83, s | 53.5, $CH_3$ |

NMR spectrum (500 MHz) for $^{1}H$, NMR spectrum (125 MHz) for $^{13}C$, $D_2O$, “m” means overlapped or multiple with other signals. Chemical shifts are reported in ppm.

HRMS (ESI, M+H$^+$) calculated for $C_{15}H_{22}NO_4^+$ 280.1543; found 280.1539.

$[\alpha]_D^{24.1}$ − 12° ($c$ 0.1, MeOH).

**Supplementary Table 18.** Spectroscopic data of compound **18** methyl ester (**18**-Me)

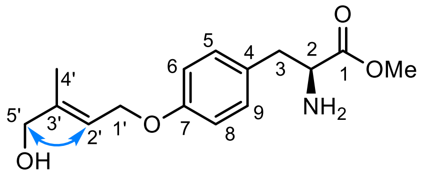


| Position | **18**-Me in DMSO-$d_6$ (500MHz) $\delta_H$ (*J* in Hz) | $\delta_C$, type |
|---|---|---|
| 1 | | 169.5, C |
| 2 | 4.24, m | 53.4, CH |
| 3 | 3.04, m | 35.2, $CH_2$ |
| 4 | | 126.2, C |
| 5 | 7.12, d (8.5) | 130.5, CH |
| 6 | 6.89, d (8.7) | 114.8, CH |
| 7 | | 157.8, C |
| 8 | 6.89, d (8.7) | 114.8, CH |
| 9 | 7.12, d (8.5) | 130.5, CH |
| 1’ | 4.57, d (6.5) | 64.1, $CH_2$ |
| 2’ | 5.64, m | 118.0, CH |
| 3’ | | 140.6, C |
| 4’ | 1.64, s | 13.8, $CH_3$ |
| 5’ | 3.84, s | 65.5, $CH_2$ |
| OMe | 3.68, s | 52.6, $CH_3$ |

NMR spectrum (500 MHz) for $^1$H, NMR spectrum (125 MHz) for $^{13}$C, DMSO-$d_6$, “m” means overlapped or multiple with other signals. Chemical shifts are reported in ppm.

HRMS (ESI, M+H$^+$) calculated for $C_{15}H_{22}NO_4^+$ 280.1543; found 280.1543.

$[\alpha]_D^{24.1}$ − 14° (*c* 0.1, MeOH).

**Supplementary Table 19.** Spectroscopic data of compound 4’,4’,4’-[$d_3$]-**7-Me**

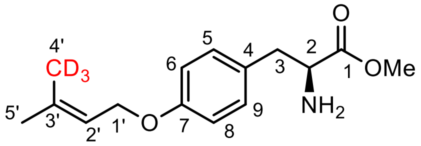


4',4',4'-[$d_3$]-**7-Me**

| Position | 4’,4’,4’-[$d_3$]-**7-Me** in $CD_3CN$ | |
|---|---|---|
| | $\delta_H$ ($J$ in Hz) | $\delta_C$, type |
| 1 | | 165.1, C |
| 2 | 3.81, m | 56.0, CH |
| 3 | 2.90, m; 2.98, m | 39.3, $CH_2$ |
| 4 | | 129.3, C |
| 5 | 7.10, d (8.9) | 131.4, CH |
| 6 | 6.84, d (8.4) | 115.5, CH |
| 7 | | 158.9, C |
| 8 | 6.84, d (8.4) | 115.5, CH |
| 9 | 7.10, d (8.9) | 131.4, CH |
| 1’ | 4.50 (d, 7.2) | 65.5, $CH_2$ |
| 2’ | 5.44, m | 120.9, CH |
| 3’ | | 138.5, C |
| 4’ | | 18.1, $CD_3$ |
| 5’ | 1.77, s | 25.7, $CH_3$ |
| OMe | 3.66, s | 52.7, $CH_3$ |

NMR spectrum (500 MHz) for $^1$H, NMR spectrum (125 MHz) for $^{13}$C, $CD_3CN$, “m” means overlapped or multiple with other signals. Chemical shifts are reported in ppm.

HRMS (ESI, M+H$^+$) calculated for $C_{15}H_{19}D_3NO_3^+$ 267.1783; found 267.1770.

**Supplementary Table 20.** Spectroscopic data of compound 5',5',5'-[$d_3$]-**7-Me**

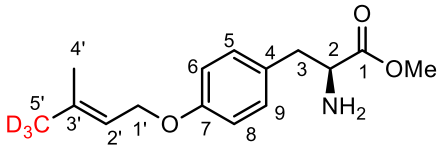


5',5',5'-[$d_3$]-**7-Me**

| Position | 5',5',5'-[$d_3$]-**7-Me** in $CD_3CN$ (500MHz) $\delta_H$ (*J* in Hz) | $\delta_C$, type |
|---|---|---|
| 1 | | 173.5, C |
| 2 | 4.31, m | 56.1, CH |
| 3 | 2.83, m; 3.02, m | 37.4, $CH_2$ |
| 4 | | 129.8, C |
| 5 | 7.09, d (8.6) | 131.2, CH |
| 6 | 6.84, d (8.4) | 115.4, CH |
| 7 | | 158.7, C |
| 8 | 6.84, d (8.4) | 115.4, CH |
| 9 | 7.09, d (8.6) | 131.2, CH |
| 1' | 4.50 (d, 6.7) | 65.4, $CH_2$ |
| 2' | 5.43, m | 120.9, CH |
| 3' | | 138.5, C |
| 4' | 1.72, s | 18.1, $CH_3$ |
| 5' | | 25.7, $CH_3$ |
| OMe | 3.66, s | 52.6, $CH_3$ |

NMR spectrum (500 MHz) for $^1$H, NMR spectrum (125 MHz) for $^{13}$C, $CD_3CN$, "m" means overlapped or multiple with other signals. Chemical shifts are reported in ppm.

HRMS (ESI, M+H$^+$) calculated for $C_{15}H_{19}D_3NO_3^+$ 267.1783; found 267.1780.

**Supplementary Table 21.** Spectroscopic data of compound **2'-[$d_1$]-7**

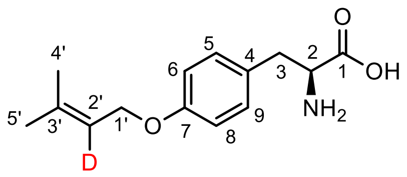


2'-[$d_1$]-**7**

| Position | **2'-[$d_1$]-7** in DMSO-$d_6$ (500MHz) $\delta_H$ ($J$ in Hz) | $\delta_C$, type |
|---|---|---|
| 1 | | 165.7, C |
| 2 | 3.59, d (6.1) | 67.0, CH |
| 3 | 1.76, m | 25.1, $CH_2$ |
| 4 | | 130.1, C |
| 5 | 7.10, d (8.0) | 130.1, CH |
| 6 | 6.80, d (7.7) | 114.1, CH |
| 7 | | 156.6, C |
| 8 | 6.80, t (7.7) | 114.1, CH |
| 9 | 7.10, d (8.0) | 130.1, CH |
| 1' | 4.46, s | 64.1, $CH_2$ |
| 2' | | 119.9, C |
| 3' | | 136.6, C |
| 4' | 1.69, s | 18.0, $CH_3$ |
| 5' | 1.73, s | 25.4, $CH_3$ |

NMR spectrum (500 MHz) for $^1$H, NMR spectrum (125 MHz) for $^{13}$C, DMSO-$d_6$, "m" means overlapped or multiple with other signals. Chemical shifts are reported in ppm.

HRMS (ESI, M+H$^+$) calculated for $C_{14}H_{19}DNO_3^+$ 251.1500; found 251.1496.

**Supplementary Table 22.** Spectroscopic data of compound **1’,1’-[$d_2$]-7**

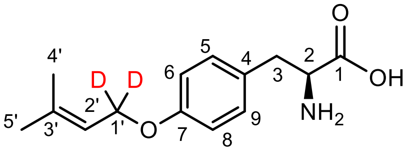


1',1'-[$d_2$]-**7**

| Position | **1’,1’-[$d_2$]-7** in DMSO-$d_6$ (500MHz) | |
|---|---|---|
| | $\delta_H$ ($J$ in Hz) | $\delta_C$, type |
| 1 | | 178.1, C |
| 2 | 3.09, m | 57.9, CH |
| 3 | 2.46, m; 2.94, m | 40.8, $CH_2$ |
| 4 | | 132.3, C |
| 5 | 7.10, d (8.1) | 130.2, CH |
| 6 | 6.80, d (8.1) | 114.2, CH |
| 7 | | 156.6, C |
| 8 | 6.80, d (8.1) | 114.2, CH |
| 9 | 7.10, d (8.1) | 130.2, CH |
| 1’ | | 63.5 |
| 2’ | 5.40, s | 120.1, CHl |
| 3’ | | 136.8, C |
| 4’ | 1.69, s | 18.0, $CH_3$ |
| 5’ | 1.73, s | 25.5, $CH_3$ |

NMR spectrum (500 MHz) for $^1$H, NMR spectrum (125 MHz) for $^{13}$C, DMSO-$d_6$, “m” means overlapped or multiple with other signals. Chemical shifts are reported in ppm.

HRMS (ESI, M+H$^+$) calculated for $C_{14}H_{18}D_2NO_3^+$ 252.1563; found 252.1561.

**Supplementary Table 23.** Spectroscopic data of compound **19-Me**

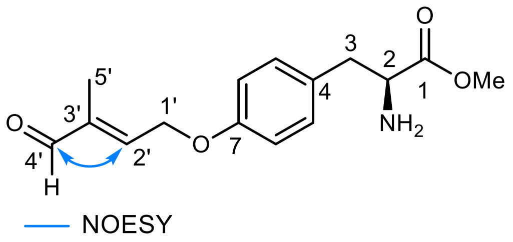


**19-Me**

| Position | **19-Me** in $D_2O$ (500MHz) | |
|---|---|---|
| | $\delta_H$ (*J* in Hz) | $\delta_C$, type |
| 1 | | 170.0, C |
| 2 | 4.40, m | 54.1, CH |
| 3 | 3.21, m; 3.30, m | 34.7, $CH_2$ |
| 4 | | 126.7, C |
| 5 | 7.24, d (8.6) | 130.7, CH |
| 6 | 6.87, d (8.7) | 115.5, CH |
| 7 | | 157.2, C |
| 8 | 6.87, d (8.7) | 115.5, CH |
| 9 | 7.24, d (8.6) | 130.7, CH |
| 1’ | 5.04, m | 65.1, $CH_2$ |
| 2’ | 6.87, td (1.4, 5.5) | 150.3, CH |
| 3’ | | 139.9, C |
| 4’ | 9.38, s | 198.2, CHO |
| 5’ | 1.80, d (1.3) | 8.7, $CH_3$ |
| OMe | 3.83, s | 53.5, $CH_3$ |

NMR spectrum (500 MHz) for $^1H$, NMR spectrum (125 MHz) for $^{13}C$, $D_2O$, “m” means overlapped or multiple with other signals. Chemical shifts are reported in ppm.

HRMS (ESI, $M+H^+$) calculated for $C_{15}H_{20}NO_4^+$ 278.1387; found 278.1392.

**Supplementary Table 24.** Spectroscopic data of compound **20**

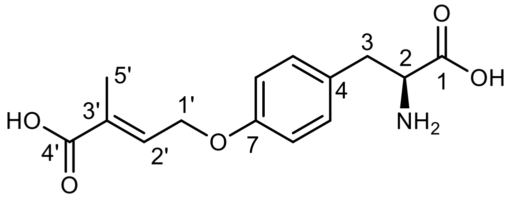


**20**

| Position | **20** in $CD_3OD$ (500MHz) $\delta_H$ (*J* in Hz) | $\delta_C$, type |
|---|---|---|
| 1 | | 176.7, C |
| 2 | 3.19, brs | 58.8, CH |
| 3 | 2.47, m; 2.82, m | 42.0, $CH_2$ |
| 4 | | 132.1, C |
| 5 | 6.93, d (8.5) | 131.5, CH |
| 6 | 6.61, d (8.6) | 115.7, CH |
| 7 | | 158.9, C |
| 8 | 6.61, d (8.6) | 115.7, CH |
| 9 | 6.93, d (8.5) | 131.5, CH |
| 1’ | 4.41, m | 66.3, $CH_2$ |
| 2’ | 6.33, m | 131.9, CH |
| 3’ | | 137.9, C |
| 4’ | | 176.7, C |
| 5’ | 1.65, d (1.3) | 14.4, $CH_3$ |

NMR spectrum (500 MHz) for $^1H$, NMR spectrum (125 MHz) for $^{13}C$, $CD_3OD$, “m” means overlapped or multiple with other signals. Chemical shifts are reported in ppm.

HRMS (ESI, $M+H^+$) calculated for $C_{14}H_{18}NO_5^+$ 280.1179; found 280.1180.

**Supplementary Table 25.** Computed energy components (in Hartree) for all intermediates and transition states calculated at the B3LYP-D3(BJ)/def2-TZVP/SMD($Et_2O$)//B3LYP-D3(BJ)/def2-SVP/IEFPCM($Et_2O$) level of theory at 298 K, 1 atm.

| Structure | E_SPC[a] | E[b] | ZPE | H_SPC[c] | TΔS | TΔqh-S[d] | G(T)_SPC | qh-G(T)_SPC[e] |
|---|---|---|---|---|---|---|---|---|
| **Int1** | -3495.6026 | -3493.2517 | 0.5639 | -3494.9978 | 0.1121 | 0.1053 | -3495.1099 | -3495.1032 |
| **Int1_Q** | -3495.5859 | -3493.2109 | 0.5605 | -3494.9835 | 0.1154 | 0.1085 | -3495.0989 | -3495.0919 |
| **Int10** | -3228.5165 | -3226.4850 | 0.4871 | -3227.9940 | 0.1019 | 0.0953 | -3228.0959 | -3228.0893 |
| **Int10_Q** | -3228.5164 | -3226.4848 | 0.4870 | -3227.9939 | 0.1026 | 0.0960 | -3228.0965 | -3228.0899 |
| **Int11** | -537.7144 | -537.1050 | 0.1756 | -537.5269 | 0.0498 | 0.0482 | -537.5767 | -537.5751 |
| **Int12** | -3228.5212 | -3226.4975 | 0.4907 | -3227.9964 | 0.0995 | 0.0933 | -3228.0959 | -3228.0898 |
| **Int13** | -3228.0693 | -3226.0450 | 0.4780 | -3227.5572 | 0.0985 | 0.0930 | -3227.6557 | -3227.6502 |
| **Int2** | -3495.6102 | -3493.2629 | 0.5660 | -3495.0044 | 0.1098 | 0.1033 | -3495.1143 | -3495.1077 |
| **Int2_Q** | -3495.5866 | -3493.2407 | 0.5634 | -3494.9826 | 0.1123 | 0.1054 | -3495.0950 | -3495.0880 |
| **Int3** | -652.8939 | -652.1516 | 0.2191 | -652.6606 | 0.0565 | 0.0539 | -652.7171 | -652.7145 |
| **Int4** | -463.0313 | -462.5131 | 0.1806 | -462.8393 | 0.0492 | 0.0477 | -462.8885 | -462.8870 |
| **Int5** | -3419.6261 | -3417.3743 | 0.5525 | -3419.0353 | 0.1081 | 0.1009 | -3419.1434 | -3419.1362 |
| **Int5_Q** | -3419.6254 | -3417.3736 | 0.5525 | -3419.0346 | 0.1090 | 0.1016 | -3419.1436 | -3419.1363 |
| **Int6** | -652.9179 | -652.1755 | 0.2200 | -652.6839 | 0.0551 | 0.0530 | -652.7390 | -652.7369 |
| **Int7** | -652.7114 | -651.9626 | 0.2202 | -652.4769 | 0.0555 | 0.0533 | -652.5324 | -652.5302 |
| **Int8** | -3419.6305 | -3417.3806 | 0.5522 | -3419.0401 | 0.1075 | 0.1005 | -3419.1476 | -3419.1406 |
| **Int8_Q** | -3419.6301 | -3417.3804 | 0.5523 | -3419.0397 | 0.1077 | 0.1009 | -3419.1474 | -3419.1406 |
| **Int9** | -3228.5148 | -3226.4853 | 0.4895 | -3227.9903 | 0.1011 | 0.0946 | -3228.0914 | -3228.0850 |
| **Int9_Q** | -3228.5144 | -3226.4850 | 0.4896 | -3227.9900 | 0.1014 | 0.0951 | -3228.0914 | -3228.0851 |
| **Product_allene** | -462.4603 | -461.9409 | 0.1703 | -462.2791 | 0.0475 | 0.0460 | -462.3266 | -462.3251 |
| **Product_yne** | -462.4605 | -461.9405 | 0.1706 | -462.2785 | 0.0498 | 0.0475 | -462.3282 | -462.3260 |
| **TS1** | -3495.5766 | -3493.2306 | 0.5592 | -3494.9787 | 0.1069 | 0.1010 | -3495.0855 | -3495.0796 |
| **TS1_Q** | -3495.5527 | -3493.2081 | 0.5564 | -3494.9566 | 0.1111 | 0.1039 | -3495.0677 | -3495.0605 |
| **TS2** | -3495.5837 | -3493.2353 | 0.5635 | -3494.9806 | 0.1085 | 0.1026 | -3495.0891 | -3495.0832 |
| **TS2_Q** | -3228.5144 | -3226.4820 | 0.4868 | -3227.9928 | 0.1021 | 0.0951 | -3228.0949 | -3228.0879 |
| **TS3** | -652.8781 | -652.1340 | 0.2163 | -652.6475 | 0.0566 | 0.0541 | -652.7041 | -652.7016 |
| **TS4** | -3419.5905 | -3417.3464 | 0.5461 | -3419.0070 | 0.1059 | 0.0990 | -3419.1129 | -3419.1060 |
| **TS4_Q** | -3419.5940 | -3417.3443 | 0.5458 | -3419.0106 | 0.1070 | 0.0999 | -3419.1176 | -3419.1105 |
| **TS5** | -652.6825 | -651.9361 | 0.2183 | -652.4502 | 0.0537 | 0.0522 | -652.5039 | -652.5024 |
| **TS6** | -3419.6047 | -3417.3529 | 0.5447 | -3419.0222 | 0.1071 | 0.0998 | -3419.1293 | -3419.1220 |
| **TS6_Q** | -3419.6057 | -3417.3548 | 0.5456 | -3419.0227 | 0.1059 | 0.0994 | -3419.1286 | -3419.1221 |
| **TS7** | -3228.4869 | -3226.4563 | 0.4829 | -3227.9698 | 0.0987 | 0.0928 | -3228.0684 | -3228.0626 |
| **TS7_Q** | -3228.4891 | -3226.4599 | 0.4832 | -3227.9720 | 0.0983 | 0.0928 | -3228.0703 | -3228.0648 |
| **TS8** | -3228.5128 | -3226.4821 | 0.4877 | -3227.9907 | 0.0990 | 0.0930 | -3228.0897 | -3228.0837 |
| **TS9** | -3228.4845 | -3226.4570 | 0.4876 | -3227.9627 | 0.0993 | 0.0934 | -3228.0620 | -3228.0561 |

[a]Electronic energy at single point level. [b]Electronic energy at optimization level. [c]Enthalpy after single point energy correction. [d]TΔS with quasi-harmonic correction. [e]Gibbs free energy with single point energy and quasi-harmonic corrections.

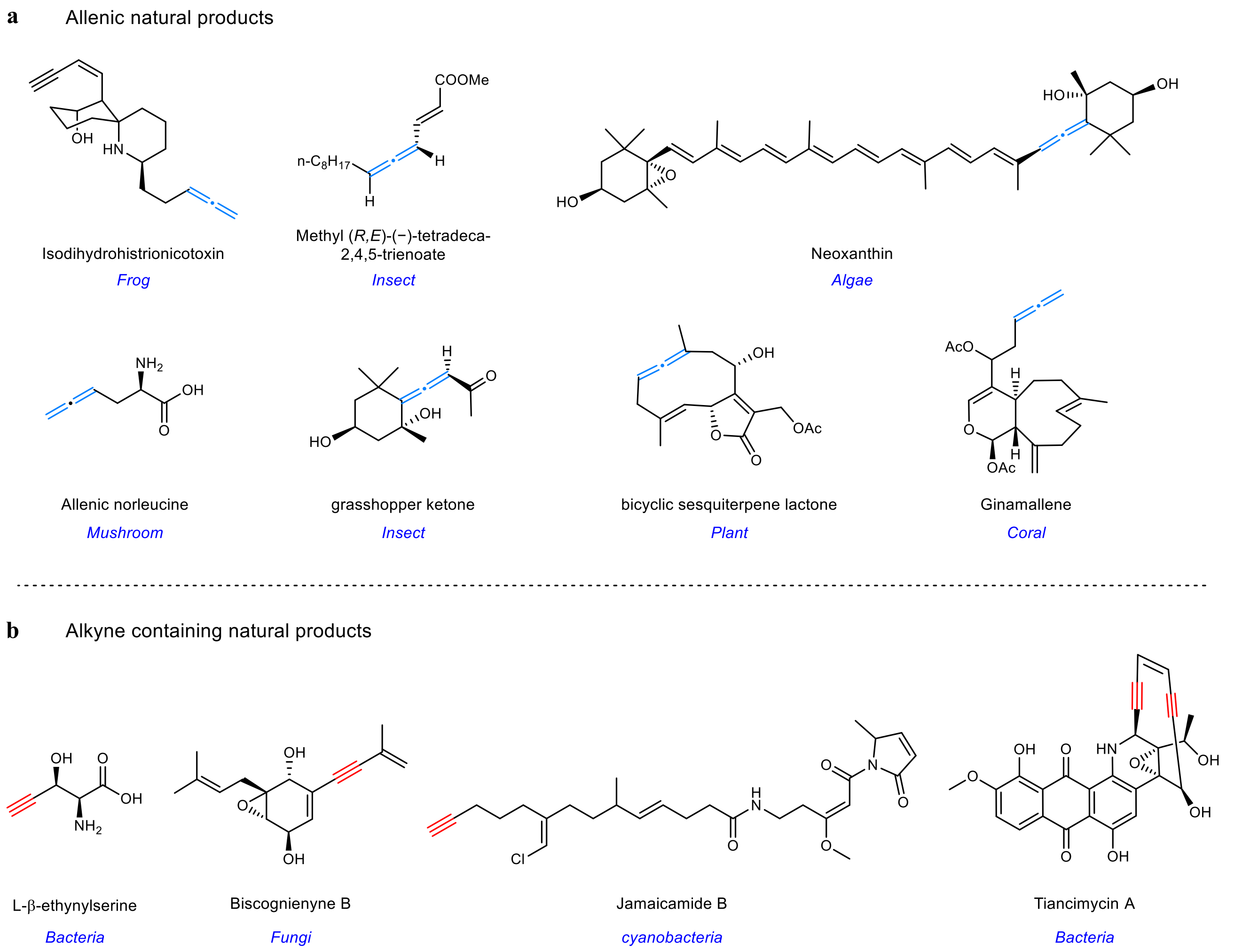


**Supplementary Fig. 1 | Representative allene- and alkyne-containing natural products.**[13–17] **a**, Allene containing natural product, were isolated from diverse origins. Several exhibit notable biological activities: isotetrahydrohistrionicotoxin, from "Poison-dart frog", interacts with ion channels of the nicotin acetylcholin receptor[13]; methyl (R,E)-(−)-tetradeca-2,4,5-trienoate functions as an insect pheromone[13]; allenic norleucine is associated with mushroom toxicity[13]; Grassshopper ketone is secreted as a defensive metabolite by *Romalea microptera*[13]; and bicyclic sesquiterpene lactones from *Vernonia* sp feature endocyclic allenes[13]. **b**, Selected internal or terminal alkyne containing natural products from fungi and bacteria[14-17], whose corresponding alkyne-forming enzymes have been characterized/identified.

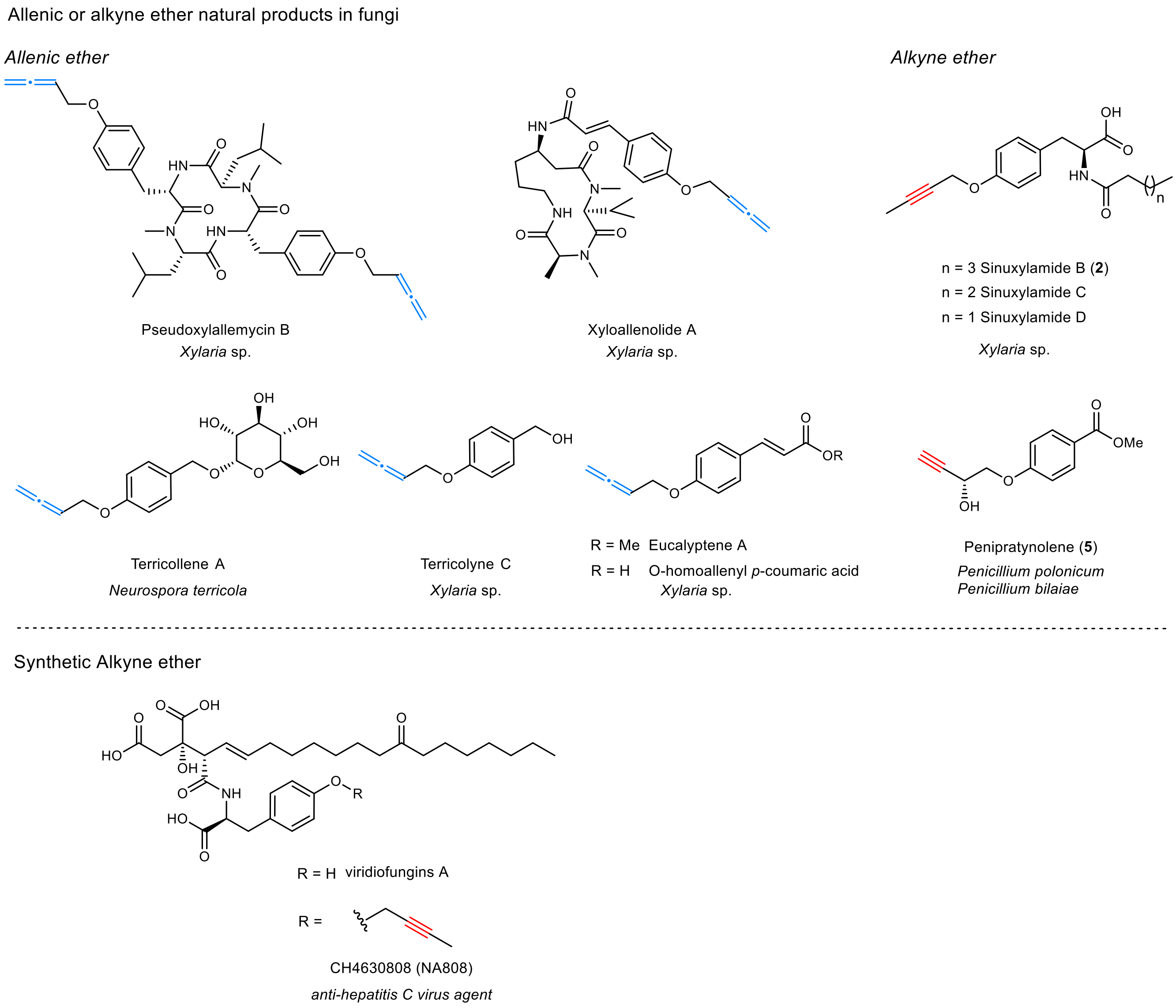


**Supplementary Fig. 2 | Structures of synthetic and natural allenic ether or alkynyl ethers. a**, Examples of natural products featuring aromatic allenic or alkynyl ether motifs, with allene and alkyne groups highlighted in light blue and red, respectively.[1,8,18–20] Most were identified from *Xylaria* sp, with the exceptions of terricollene from *Neurospora terricola* and penipratynolene from *Penicillium polonicum*. **b**, Structure of synthetic alkyne ether CH4630808 (NA808), a novel anti-hepatitis C virus agent.[21]

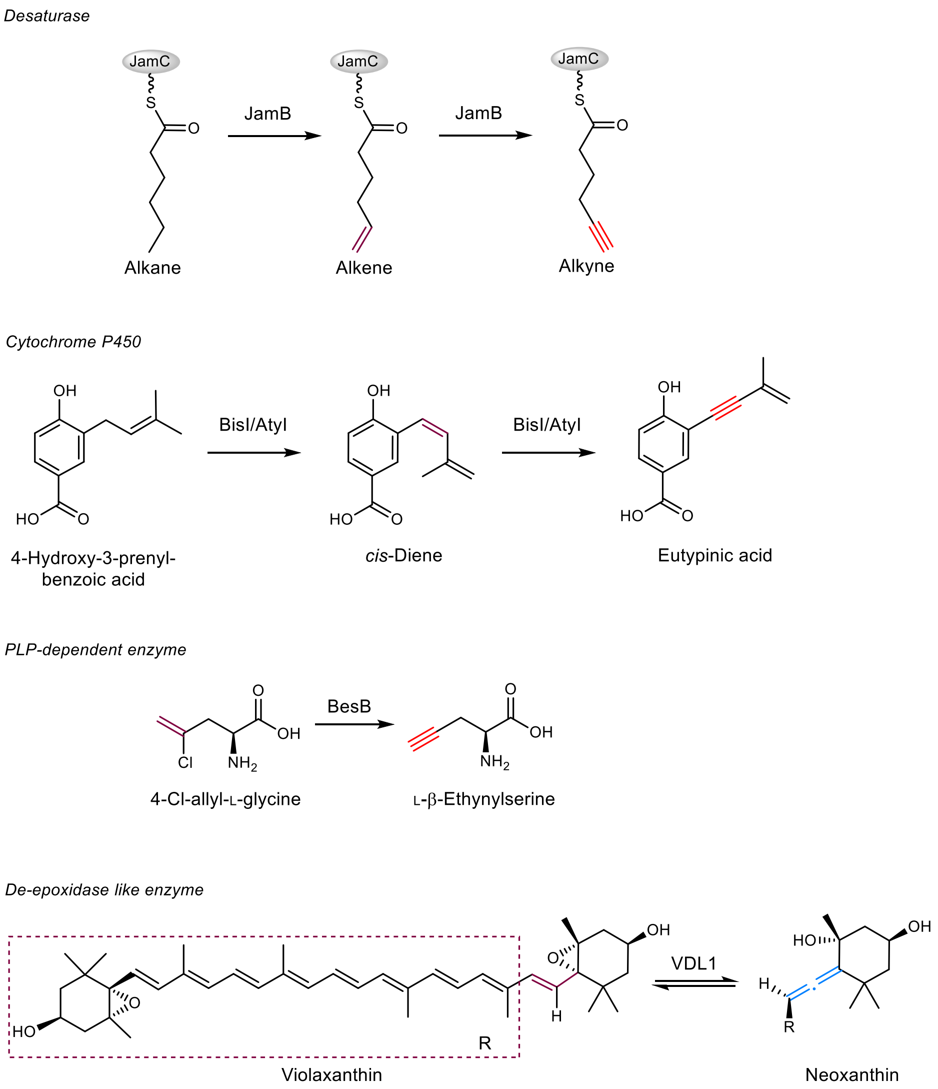


**Supplementary Fig. 3 | Reported enzymes which catalyze the formation of alkyne or allene. a**, For alkynes-forming enzymes, Zhang's group reported a bacterial membrane-bound di-iron-dependent desaturase, JamB[14], which catalyzes the successive oxidation of JamC-tethered saturated fatty acid to generate a terminal alkyne. Both Lin's and Gao's groups independently discovered P450s AtyI[22] and BisI[15] that catalyze the consecutive desaturations of a prenyl group to yield 1,3-enyne moiety. Additionally, Chang's and Ryan's groups reported a PLP-dependent enzyme BesB that catalyzes the formation of L-β-ethynylserine from 4-Cl-allyl-L-glycine by a redox-neutral ionic mechanism via a PLP-tethered allene intermediate[16,23]. **b**, Only known enzyme for formation of allenes is an algal violaxanthin de-epoxidase like enzyme VDL1 that catalyzes the conversion of violaxanthin to neoxanthin with the allene formation through 1,2-elimination[24].

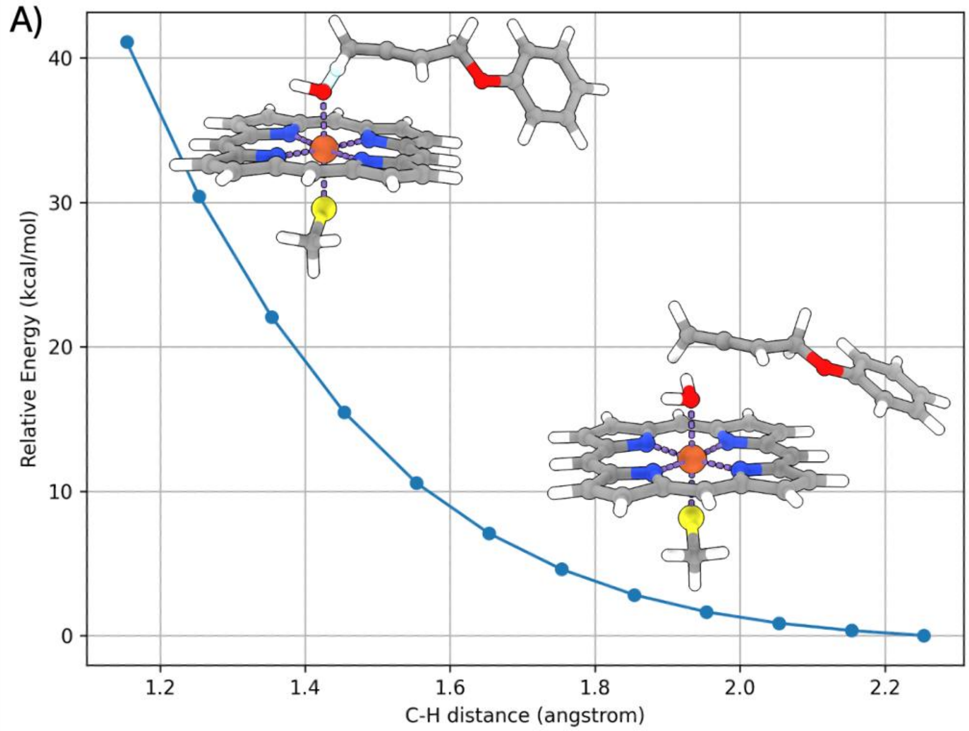


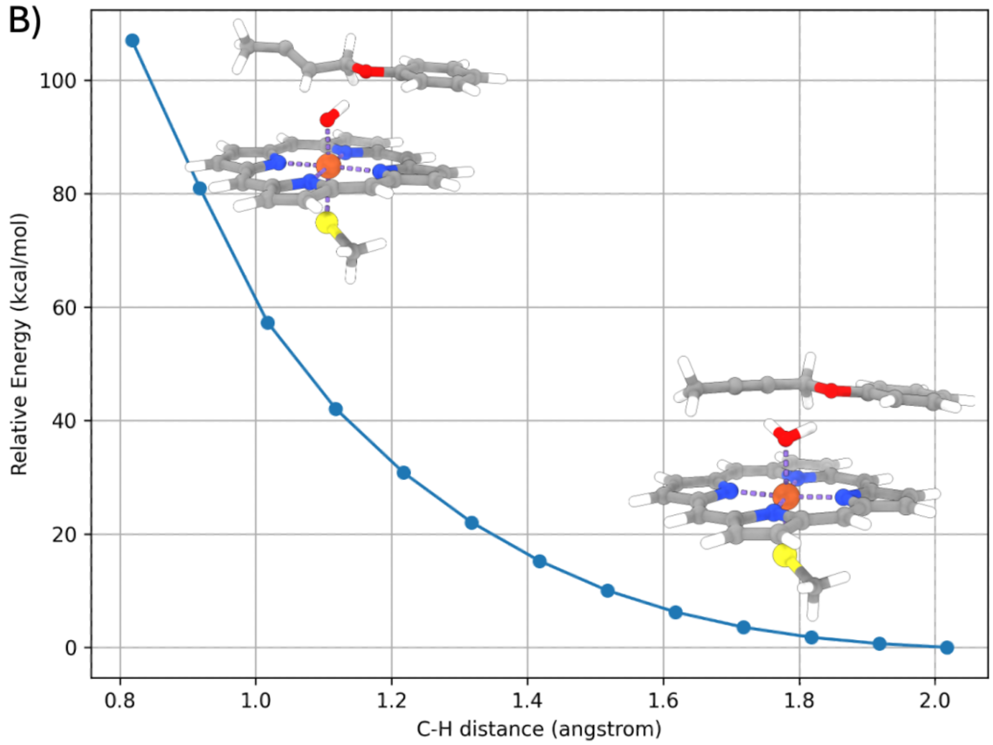


**Supplementary Fig. 4 | Computational relaxed scan of the directly attack from Int4 to allene (A) and alkyne product (B).** The results show a dramatic energy decrease during the HAT process. We tried TS optimization, but no saddle points can be located. Therefore, we believe these processes are barrierless on the potential energy surface, but could have very small barriers on the free energy surface.

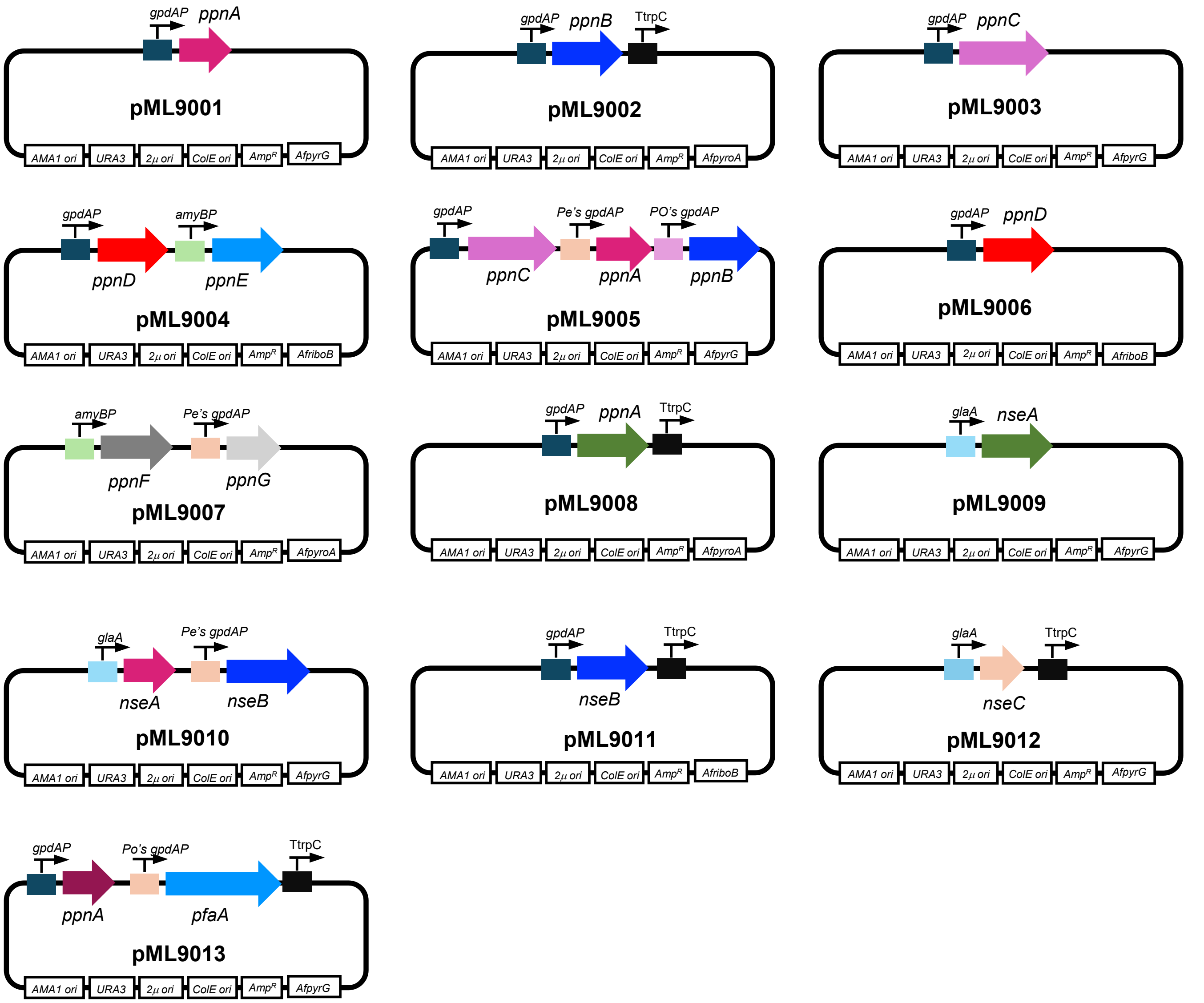


**Supplementary Fig. 5 | Plasmids used for heterologous expression of *ppn* and *nse* genes in *A. nidulans* heterologous host.**

.

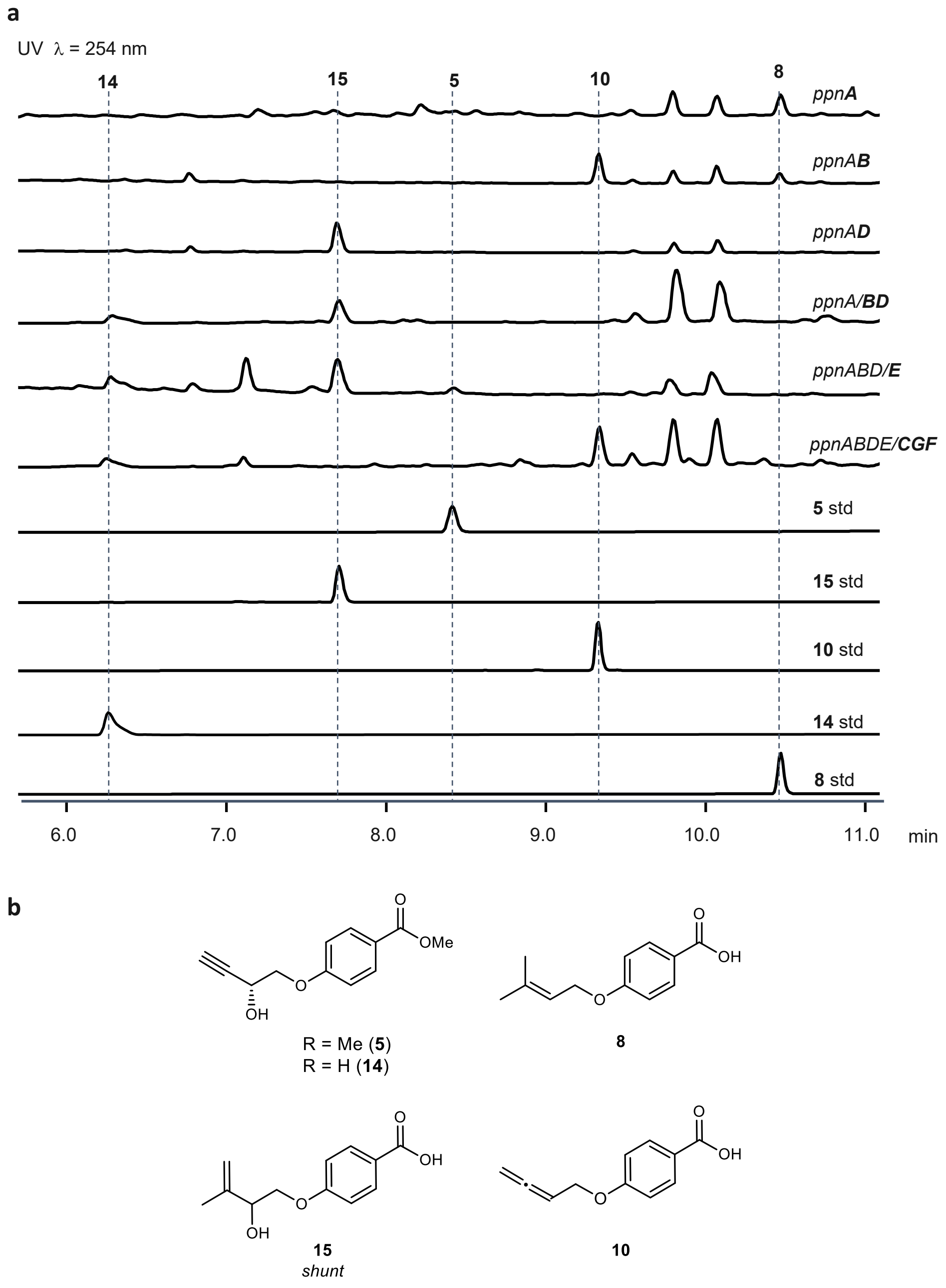


**Supplementary Fig. 6 | Heterologous expression of the *ppn* cluster. a.** Heterologous overexpression of *ppn* genes in *A. nidulans*. Overexpression of *ppnA* led to the formation of **8**, a shunt product. Co-expression of *ppnAB* produced **10**. Expression of *ppnAD* yielded **15**, and expression of *ppnABD* generated **15** as the major product with trace amounts of **14**. Overexpression of *ppnABDE* resulted in trace production of **5**. Introduction of the full cluster (*ppnABDECFG*) did not significantly increase the production of **5**. **b.** Structure of compounds **5**, **8**, **10**, **14** and **15**.

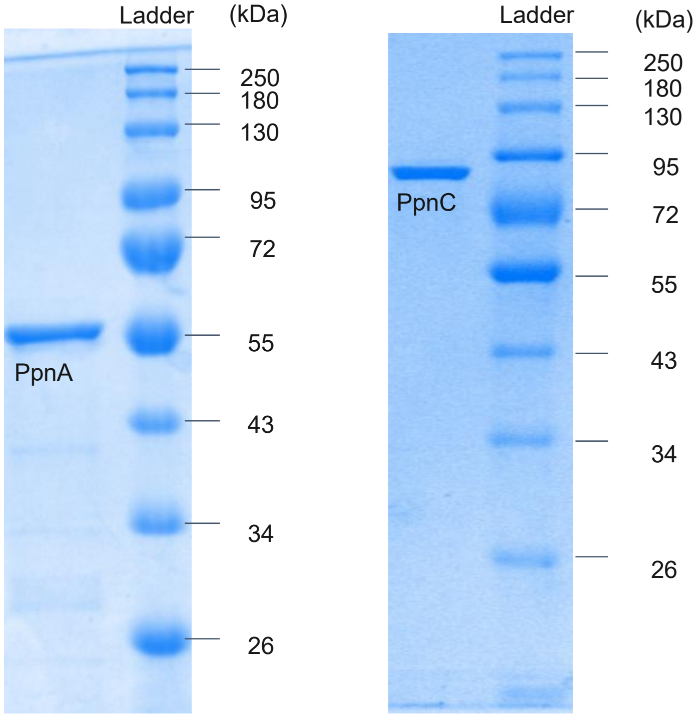


**Supplementary Fig. 7 | SDS-PAGE gels of purified proteins used in this study.** Expected molecular weights of PpnA (prenyltransferase), PpnC (aromatic amino acid ammonia lyase), 54 kDa, and 80 kDa, respectively. These experiments were repeated three times independently and representative results are shown.

**a**

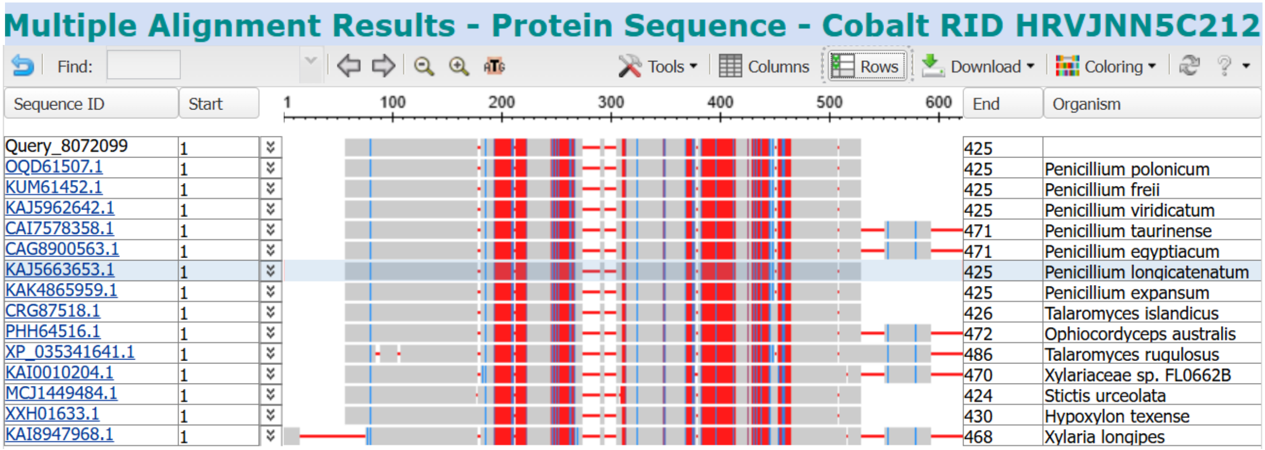


**b** PpnA_425aa

MAQTLLDESPEFQFGRYQFASNGLSTSSADLEPREHDRRITRYPYYPVELSIWQRVNSELDSFESVHHRFWWSRHTGKALAVLLYNAQYPADLQYWNLKFFAE
AVAPHLGVAPEILGSDTPIWPSFMTDDGTPVELSWDWGTKDAPPMVRYSVEPIGLHAGTSVDPGNLTAGPAFQERLTRSLPTMRLEWFHHFKDFFNIPNAKEGE
FHEDTRDHNSSIFYGFDCSETEITPKVYFFPKLRAKASGQSNLDVLFQAMRTAPHVTDRNFEAGDIFHAFCSSVGSKSLEHEMLAIDLIDPLQSRLKIYFRSRETTF
QSVINIMTLEGRIRNPKLYEGLVDLHRLWTALFGVYAVDQPLREVEHRTSGILYNFEFRLGEALPVAKIYLPVRHYCTSDEAVIRALNDYFQGQQKGKYMPDYVRA
MSTLL*

PpnA_471aa

MAQTLLDESPEFQFGRYQFASNGLSTSSADLEPREHDRRITRYPYYPVELSIWQRVNSELDSFESVHHRFWWSRHTGKALAVLLYNAQYPADLQYWNLKFFAE
AVAPHLGVAPEILGSDTPIWPSFMTDDGTPVELSWDWGTKDAPPMVRYSVEPIGLHAGTSVDPGNLTAGPAFQERLTRSLPTMRLEWFHHFKDFFNIPNAKEGE
FHEDTRDHNSSIFYGFDCSETEITPKVYFFPKLRAKASGQSNLDVLFQAMRTAPHVTDRNFEAGDIFHAFCSSVGSKSLEHEMLAIDLIDPLQSRLKIYFRSRETTF
QSVINIMTLEGRIRNPKLYEGLVDLHRLWTALFGVYAVDQPLREVEHRTSGILYNFEFRLGEALPVAKIYLPVRHYCTSDEAVIRALNDYFQGQQKGKYMPDYVRA
MSTL**FTPKSMRENSGVQTYVGCAIRPDGTLRVVSYFKPQVPVHLFESEVYL***

**c**

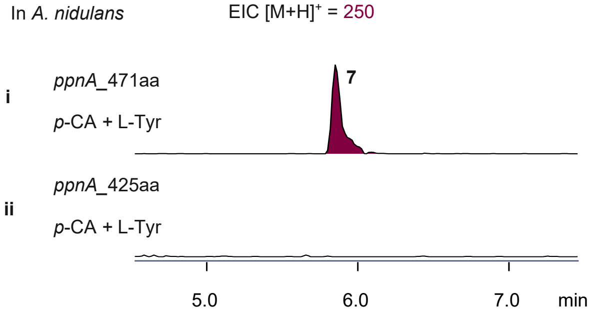


**Supplementary Fig. 8 | Annotation of PpnA. a**. Multiple sequence alignment of PpnA homologs retrieved from the NCBI database shows two alternative annotations for PpnA: a shorter around 425 amino acids (aa) form and a longer form of around 471aa. **b**. Amino-acid sequences of the two annotated PpnA variants. The 425aa form corresponds to a truncated annotation lacking the C-terminal region, whereas the 471aa variant contains the full-length sequence. **c**. LC–MS analysis of *A. nidulans* expressing the two PpnA variants. Only expression of the full-length PpnA_471aa supported in vivo production of compound **7** (EIC *m/z* +250), whereas the truncated PpnA_425aa failed to generate detectable products. These results indicate that the 471 aa variant is the catalytically active form and that the current NCBI gene annotation of PpnA from *P. polonicum* is misannotated.

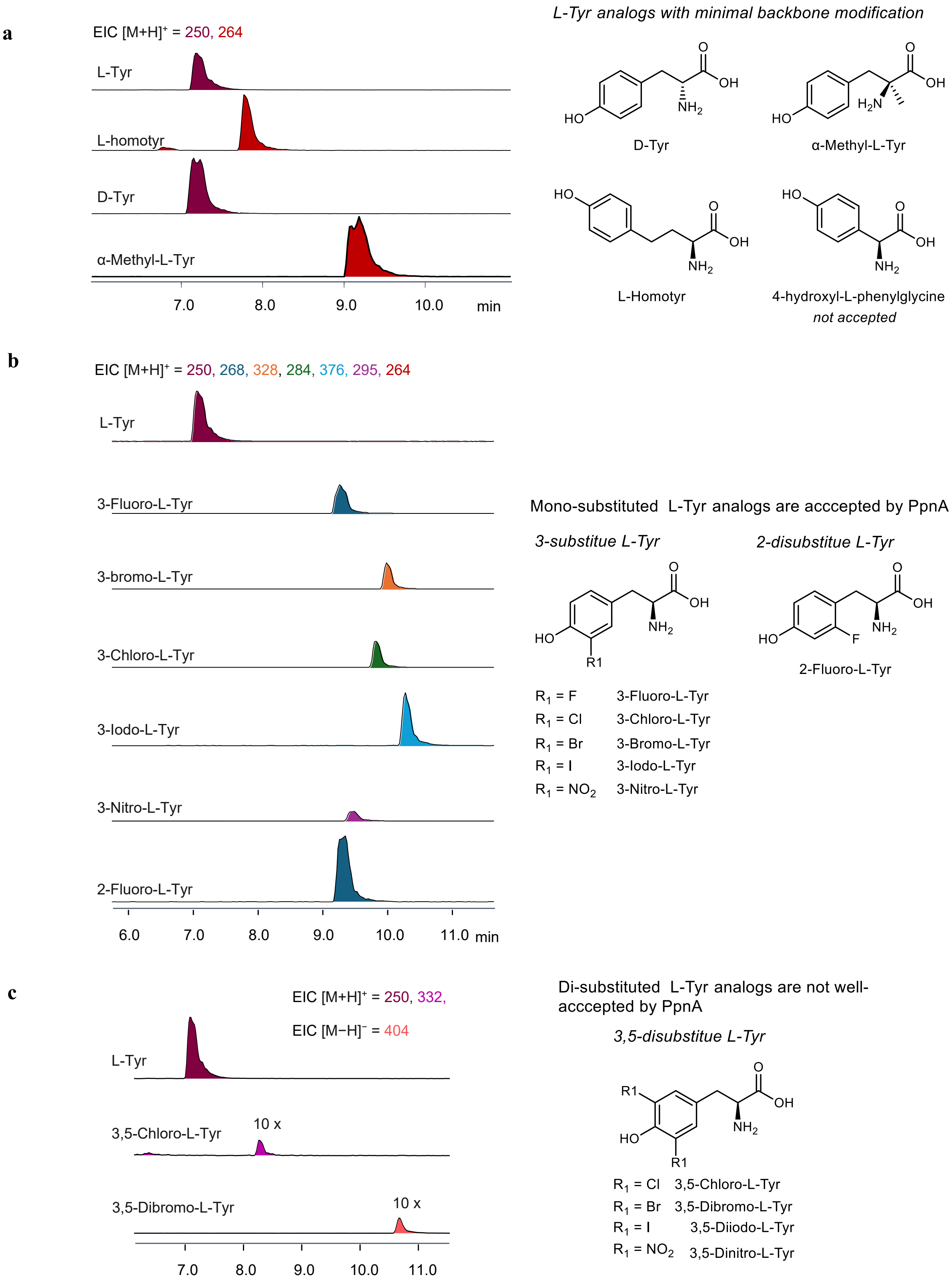


**Supplementary Fig. 9 | Substrate specificity of PpnA toward L-tyrosine analogs. a.** PpnA tolerates L-Tyr analogs with minor backbone modifications, including D-Tyr, α-Me-L-Tyr, and L-homotyrosine, yielding products with [M

+ H]$^+$ = 250 for L-Tyr and D-Tyr; [M + H]$^+$ = 264 for α-Me-L-Tyr and L-homotyrosine. In contrast, 4-hydroxy-L-phenylglycine is not recognized as a substrate. **b.** Substrate scope of PpnA toward mono-substitute L-tyrosine analogs. PpnA activates a range of mono-substituted L-Tyr derivatives, including 3-fluoro-, 3-chloro-, 3-bromo-, 3-iodo-, 3-nitro-, and 2-fluoro-L-Tyr, as shown by LC–MS detection of corresponding prenylated products ([M + H]$^+$ = 250 for L-Tyr; [M + H]$^+$ = 268 for 3-fluoro-, and 2-fluoro-L-Tyr; [M + H]$^+$ = 284 for 3-bromo-L-Tyr; [M + H]$^+$ = 376 for 3-Iodo-L-Tyr; [M + H]$^+$ = 285 for 3-nitro-L-Tyr). **c.** Di-substituted L-Tyr analogs are not efficiently accepted by PpnA. No product formation was observed for 3,5-diiodo-L-Tyr or 3,5-dinitro-L-Tyr, while trace levels of products were detected for 3,5-dichloro- and 3,5-dibromo-L-Tyr ([M + H]$^+$ = 332 for 3,5-dichloro-L-Tyr; [M − H]$^-$ = 404 for 3,5-dibromo-L-Tyr). The y-axis represents ion counts, and all selected ion chromatograms are shown on the same scale. The colors of the traces correspond to the MS spectra of the respective prenylated products.

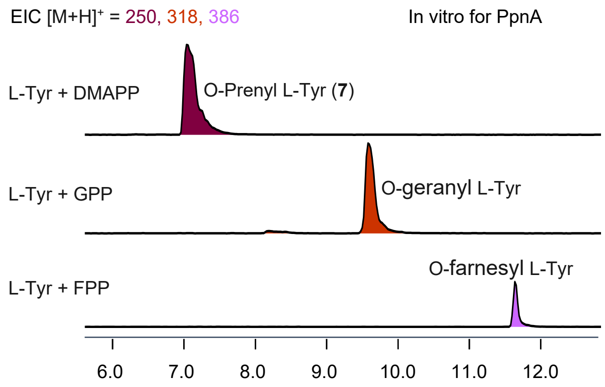


**Supplementary Fig. 10 | Substrate specificity of PpnA toward different prenyl donors.** LC–MS analysis of reactions using L-Tyr with various prenyl donors shows that PpnA accepts DMAPP, GPP, and FPP to generate distinct prenylated products with [M + H]$^+$ = 250, 318, and 386, respectively. The chromatograms (EICs) correspond to the mono-, geranyl-, and farnesyl-tyrosine products, confirming that PpnA can utilize multiple isoprenoid diphosphates as prenyl donors.

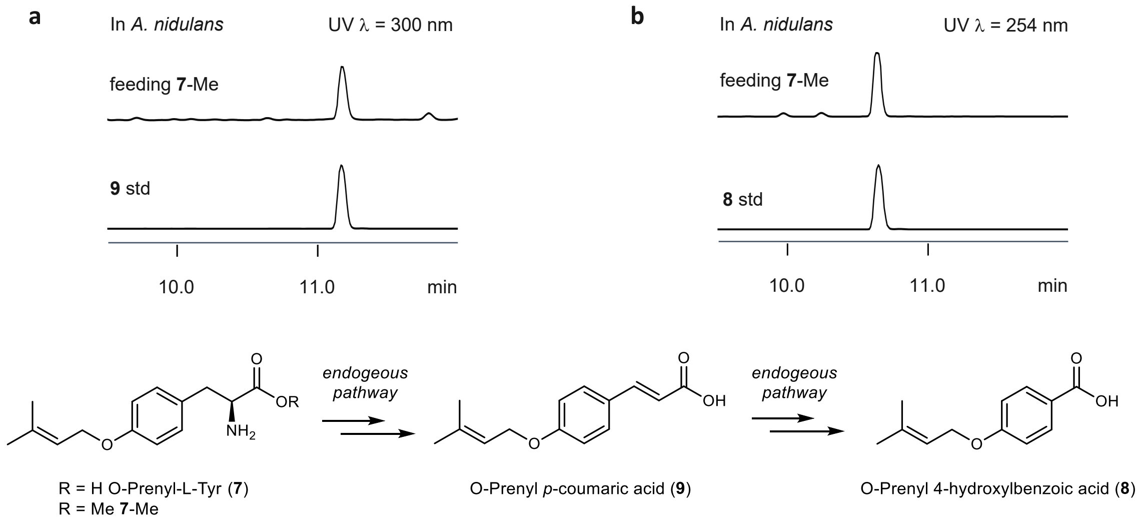


**Supplementary Fig. 11 | LC-MS analysis of the metabolites extracted from *A. nidulans* feeding 7-Me. a.** LC traces of *A. nidulans* extracts after feeding **7**-Me, monitored at UV = 300 nm. The formation of *O*-prenyl-*p*-coumaric acid (**9**) was observed, suggesting conversion of **7**-Me through an endogenous pathway. **b.** LC traces monitored at UV = 254 nm showing formation of *O*-prenyl-4-hydroxybenzoic acid (**8**) via intermediate **9** through an endogenous pathway. These results support that compound **8** is the shunt from **7** through endogenous metabolism.

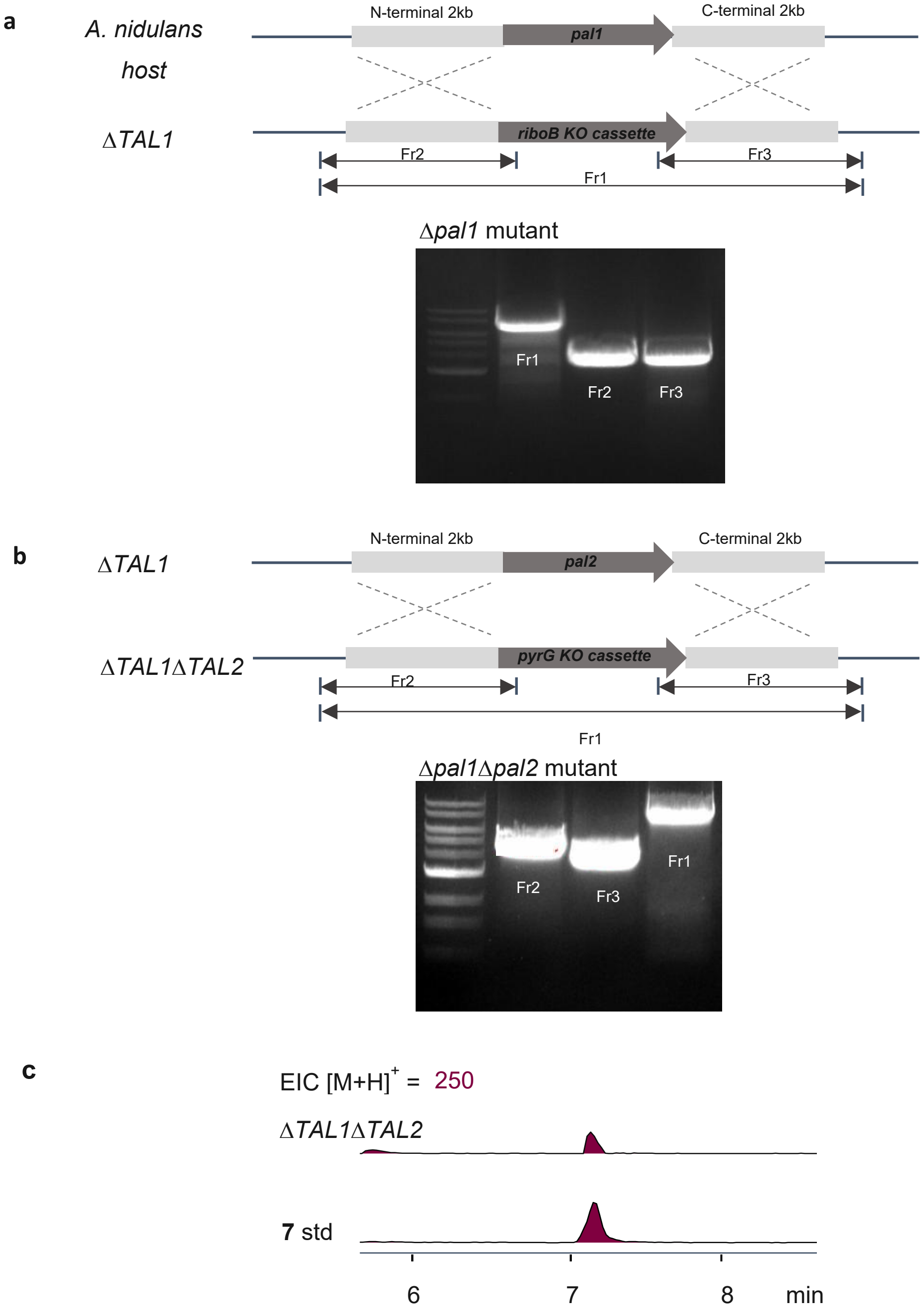


**Supplementary Fig. 12 | Diagnostic PCR for *A. nidulans* A1145 Δ*TAL1*Δ*TAL2* mutant.** Sequence identity of PpnC to *A. nidulans* endogenous tyrosine ammonia-lyases (TALs): TAL1 (XP_661501.2), 41%; TAL2, 41% (XP_663679.1). Recombinant expression and purification of TAL1 and TAL2 from *E. coli* yielded no detectable soluble protein, preventing confirmation of their proposed deamination activity in vitro. Instead, functional analysis proceeded through gene deletion in *A. nidulans*. **a.** PCR analysis confirmed knockout of TAL1 via insertion of a DNA cassette containing the riboB (riboflavin) selection marker. **b.** PCR confirmation of TAL2 knockout via insertion of a DNA cassette containing the *pyrG* (uracil) selection marker. **c.** Overexpression of *ppnA* in the *A. nidulans* A1145 Δ*TAL1*Δ*TAL2* mutant led to accumulation of compound **7** without feeding of L-Tyr. However, **7**

was still slowly converted to *O*-prenyl-4-hydroxybenzoic acid (**8**). This suggests that additional endogenous pathways may contribute to processing of the tyrosine derived moiety in vivo, potentially including transaminase-mediated reaction pathways.

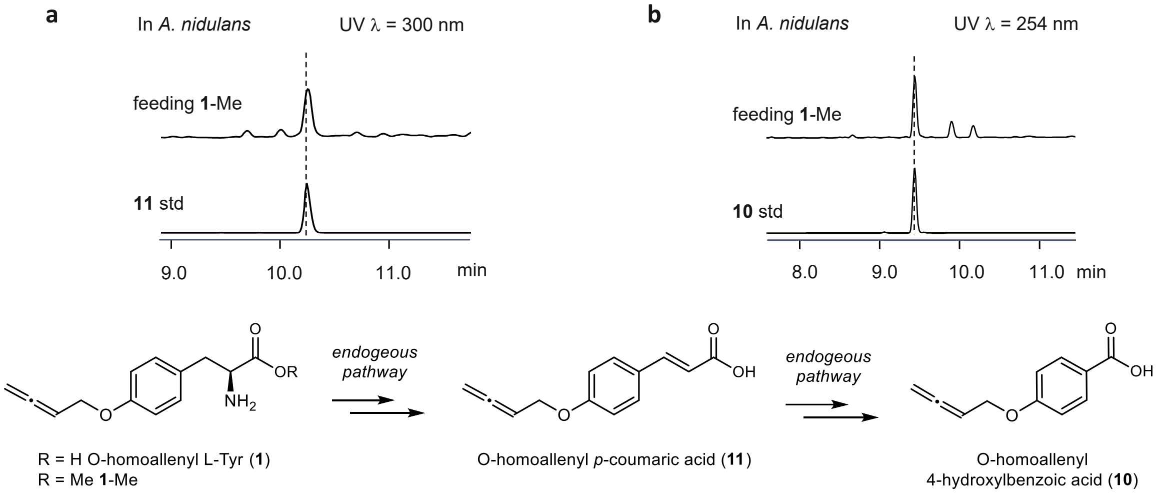


**Supplementary Fig. 13 | LC-MS analysis of metabolites extracted from *A. nidulans* feeding 1-Me. a.** LC traces of *A. nidulans* extracts after feeding **1**-Me, monitored at 300 nm. The formation of *O*-homoallenyl *p*-coumaric acid (**11**) was observed, suggesting conversion of **1**-Me through an endogenous pathway. **b.** LC traces monitored at 254 nm showing formation of *O*-homoallenyl 4-hydroxybenzoic acid (**10**) via intermediate **11** through an endogenous pathway.

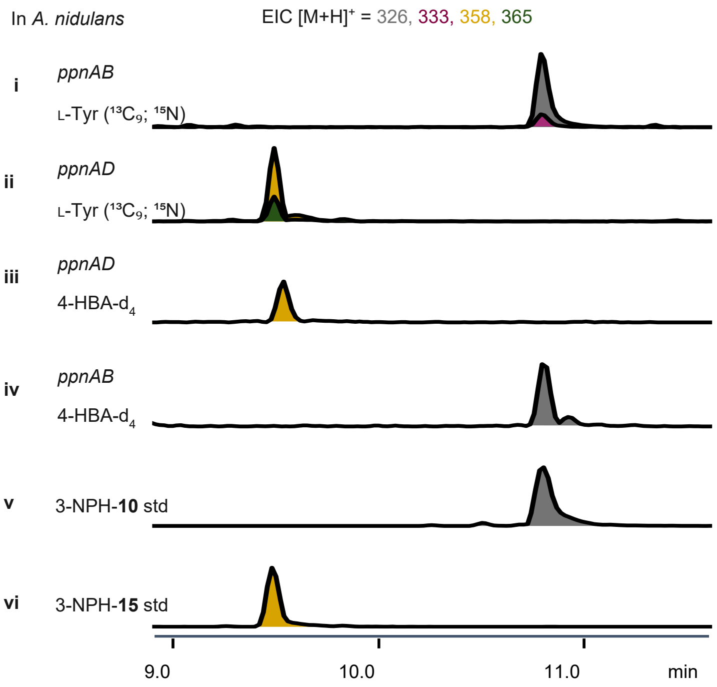


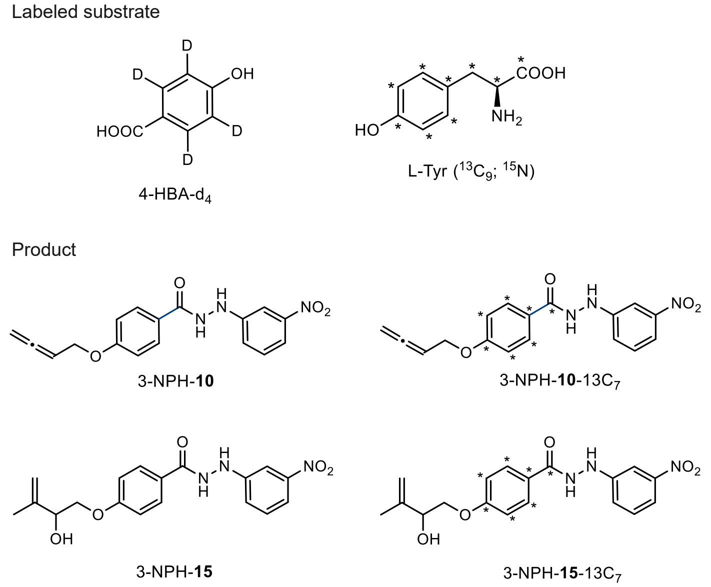


**Supplemenary Fig. 14 | LC-MS analysis of the metabolites extracted from *A. nidulans* expressing *ppnAB* or *ppnAD* feeding with labeled L-Tyr ($^{13}C_9$; $^{15}N$) and 4-hydroxybenzoic acid-$d_4$ (4-HBA-$d_4$) after 5 days.** The crude extracts were derivatized with 3-nitrophenylhydrazine (3-NPH) prior to LC-MS analysis, in order to increase the MS sensitivity. Labeled 3-NPH-**10** and 3-NPH-**15** were detected when feeding 0.5 mM L-Tyr ($^{13}C_9$; $^{15}N$) to *A. nidulans* expressing *ppnAB* and *ppnAD* respectively. Feeding 0.5 mM 4-HBA-$d_4$ did not lead to deuterium incorporation in 3-NPH-**10** or 3-NPH-**15**. Selected ion monitoring of 3-NPH-**10** ($[M + H]^+ = 326$), 3-NPH-**10-**$^{13}$C7 ($[M + H]^+ = 333$), 3-NPH-**15** ($[M + H]^+ = 358$), and 3-NPH-**15-**$^{13}$C7 ($[M + H]^+ = 365$) is shown. Y-axis represents ion counts and the chromatograms are presented on the same scale.

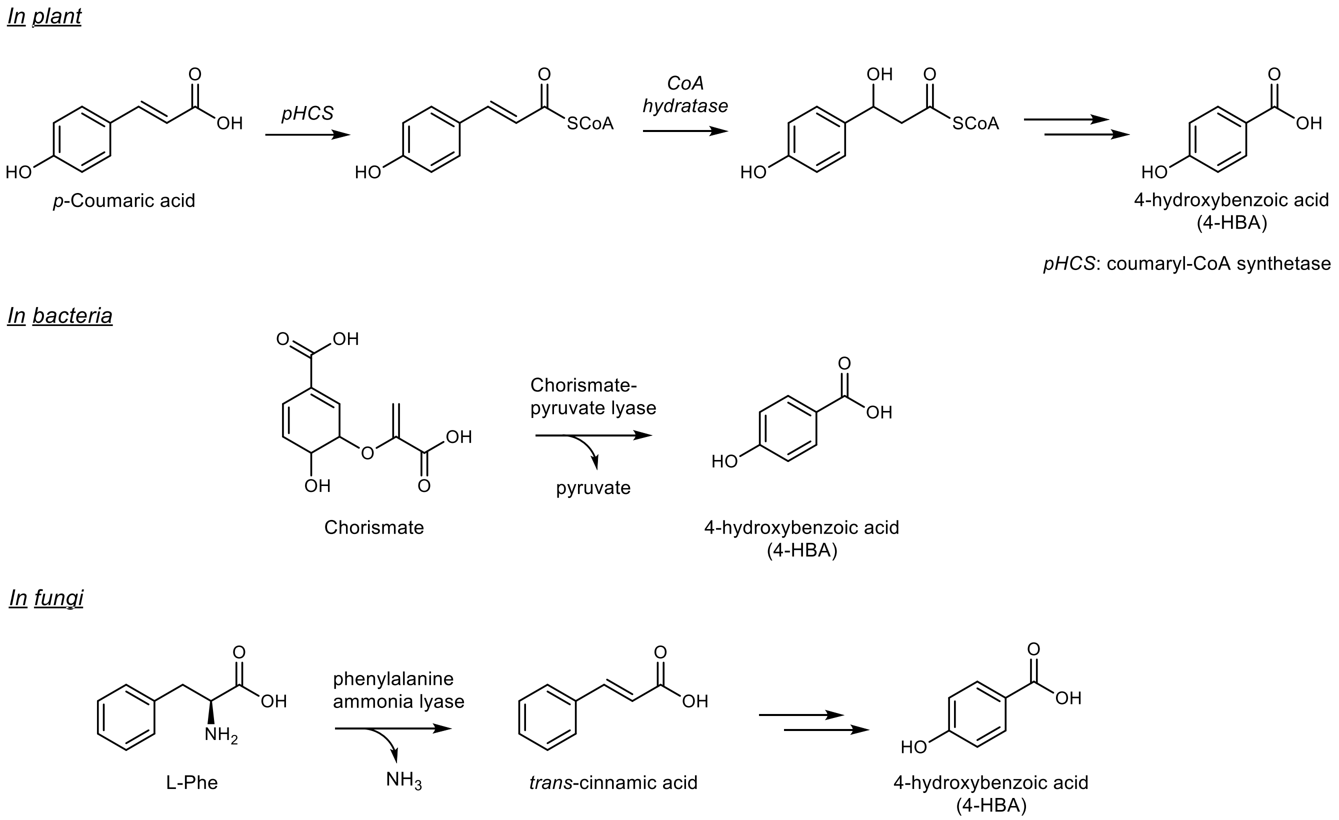


**Supplementary Fig. 15 | Pathways for 4-hydroxybenzoic acid (4-HBA) formation in plants, bacteria, and fungi.** In plants, 4-HBA formation has been proposed to proceed via a CoA-dependent route, in which coumaric acid is first activated to its CoA thioester by coumaroyl-CoA synthase, followed by hydration by a CoA hydratase and subsequent C–C bond cleavage and oxidation to yield 4-HBA[25]. In bacteria, 4-HBA is generated from chorismate through the action of chorismate pyruvate lyase, which cleaves pyruvate from chorismate[26]. In fungi, phenylalanine ammonia-lyase (PAL) catalyzes the deamination of L-Phe to form cinnamic acid, which is further converted to 4-HBA through downstream oxidative steps[22]. However, the 4-hydroxybenzoic acid moiety in the penipratynolene (**5**) is biosynthesized from a pathway starting from L-Tyr, which is different from the plant, bacterial, and canonical fungal catabolic pathways described above.

a

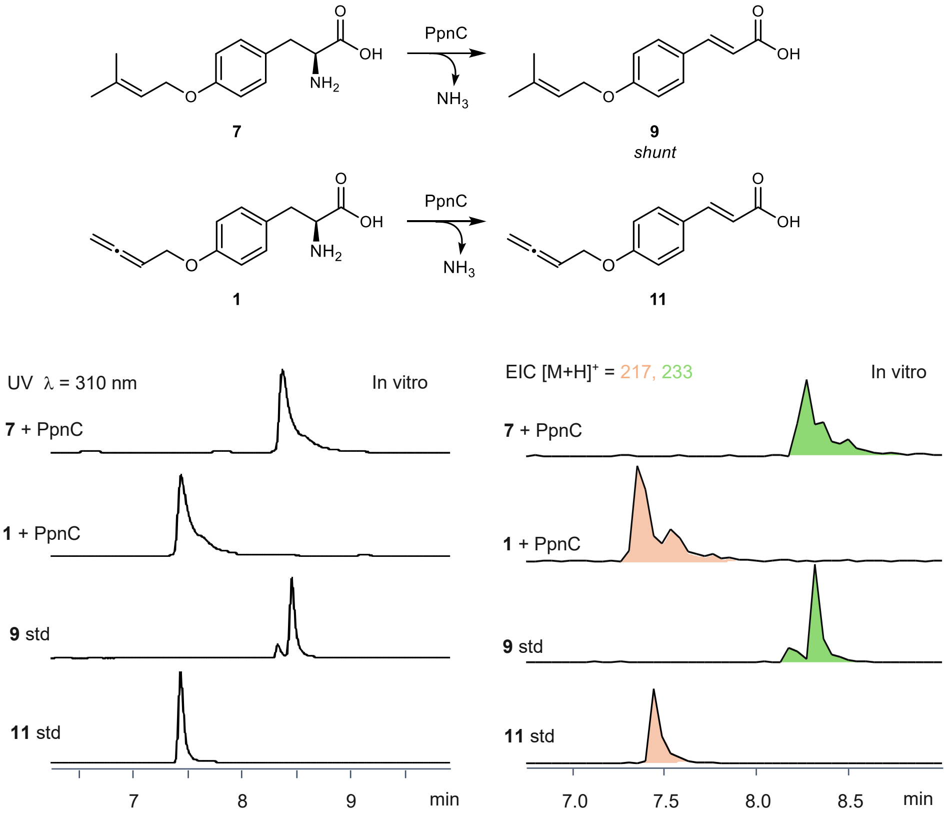


b

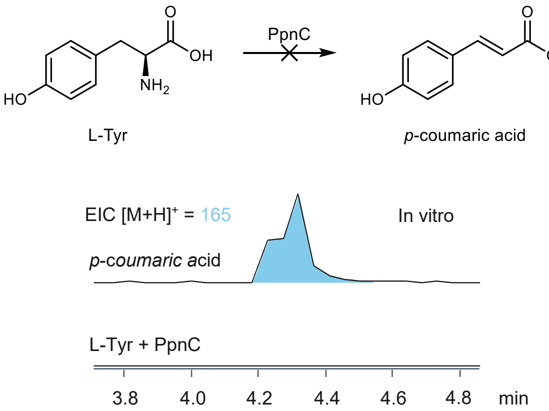


**Supplementary Fig. 16 | In vitro reaction of PpnC. a. Left panel**, LC-MS traces of in vitro reaction of PpnC using substrates *O*-homoallenyl L-Tyr (**1**)**,** *O*-prenyl L-Tyr (**7**), and *O*-but-2-ynyl L-Tyr (**6**). PpnC efficiently convert **1** and **7** to form **11** and **9**, respectively. Trace amount deaminated **6** is also observed. **Right panel**, EIC of in vitro reaction in **a**, showing the desired product, $[M + H]^+ = 233$ for **9** and $[M + H]^+ = 217$ for **11**. **b.** LC-MS traces of in vitro reaction of PpnC with L-Tyr as the substrate, no deaminated product *p*-coumaric acid (*p*-CA) was observed.

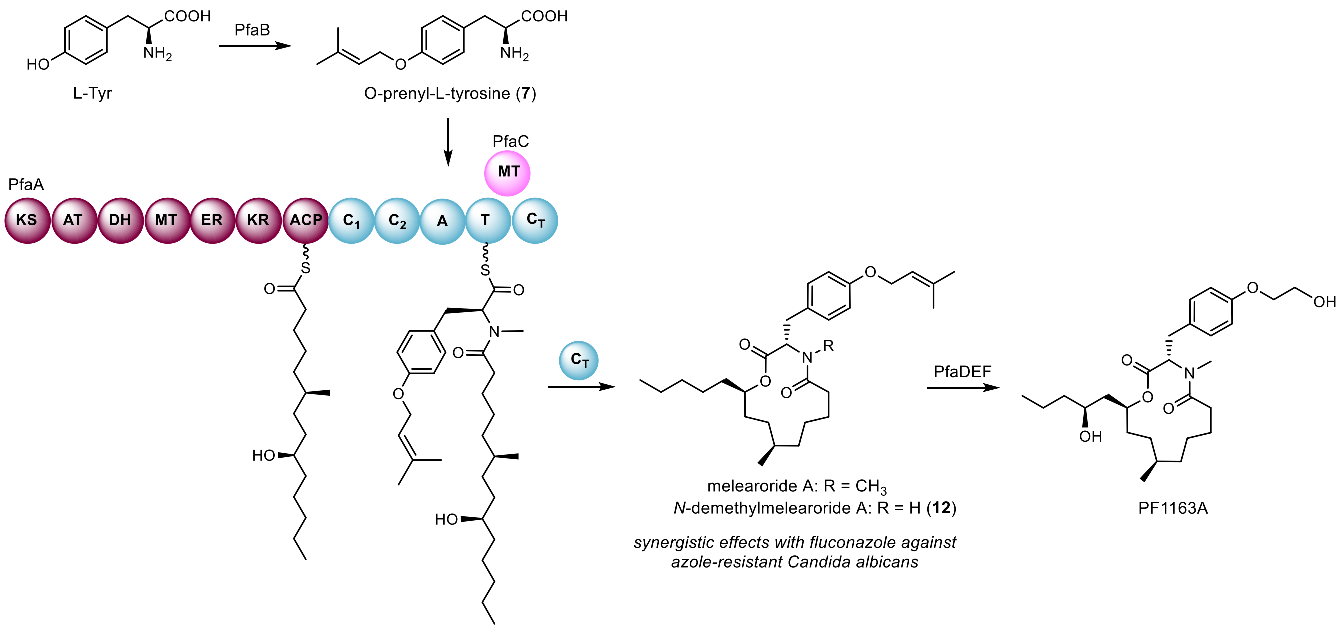


**Supplementary Fig. 17 | Reported native pathway for melearolide A and PF1163A[27].** Compound **7** is incorporated by PfaA to form *N*-demethylmelearolide A or melearolide A. Downstream tailoring enzymes further modify the side chain and prenyl group to yield PF1163A. The pathway enzymes include: PfaA (PKS–NRPS hybrid; domain architecture: KS–AT–DH–MT–ER–KR–ACP–C1–C2–A–T–CT), PfaB (prenyltransferase), PfaC (methyltransferase), PfaD (short-chain reductase), PfaE (α-ketoglutarate–dependent dioxygenase), and PfaF (globin-like ER-bound oxygenase).

**a**

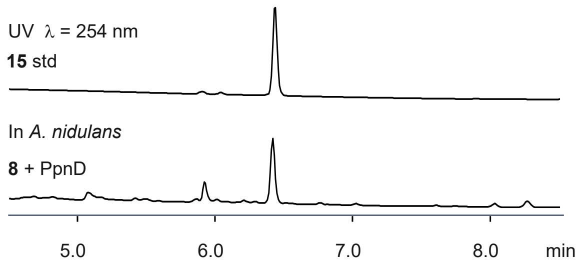


**b**

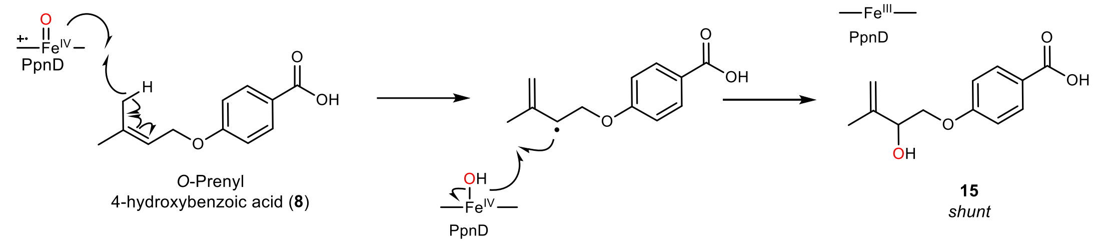


**Supplementary Fig. 18 | PpnD catalyzes oxidative modification of *O*-prenyl-4-hydroxybenzoic acid (8). a.** HPLC analysis showing that direct feeding of *O*-prenyl-4-hydroxybenzoic acid (**8**) to *A. nidulans* expressing PpnD resulted in the formation of shunt **15**, indicating that PpnD catalyzes an oxidative modification in the later stage of the pathway. **b.** Proposed mechanism of the PpnD-catalyzed reaction.

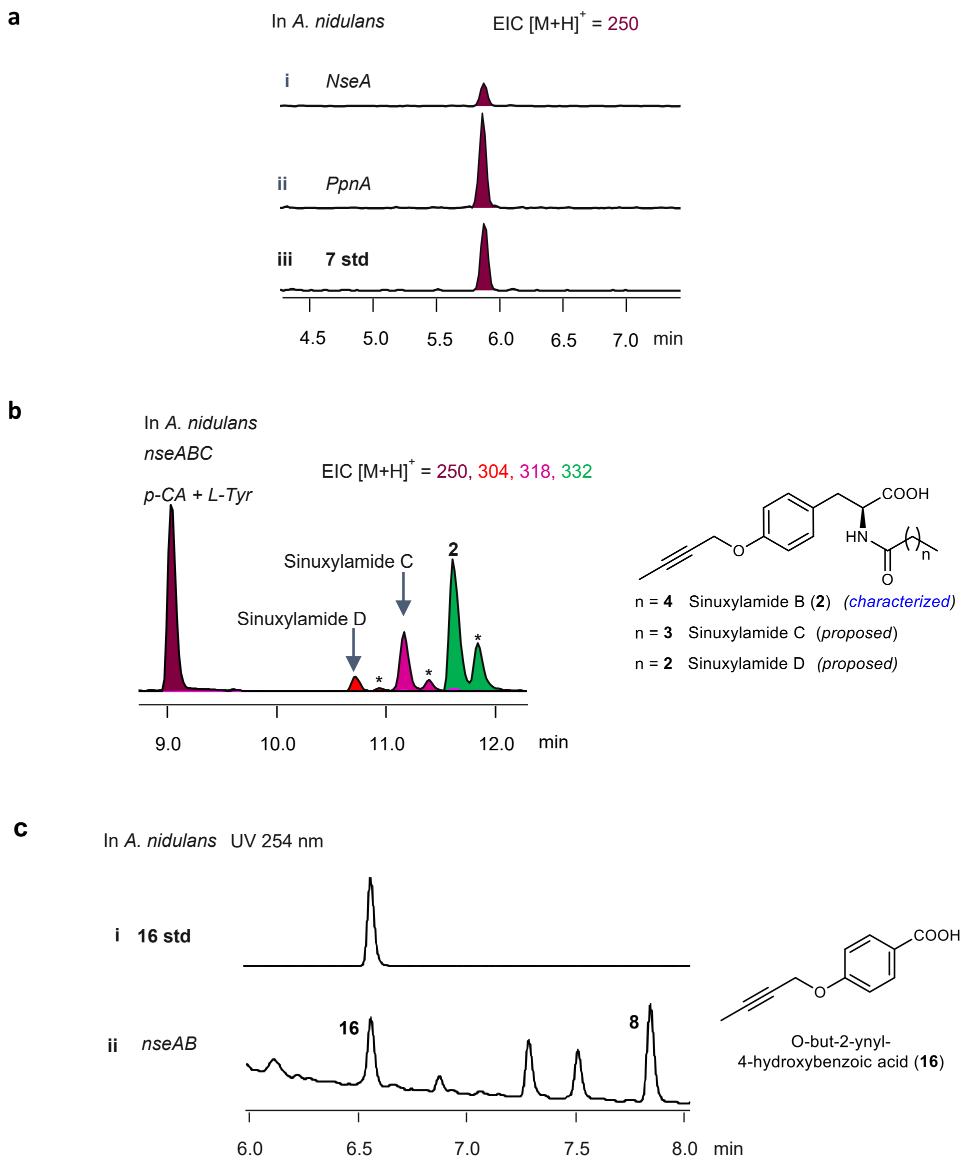


**Supplementary Fig. 19 | Overexpression of NseABC leads to production of internal alkyne-containing natural products, sinuxylamides. a**. Compound **7** was also observed when *nesA* was expressed in *A. nidulans*. **b.** Overexpression of NseABC in *A. nidulans* led to the formation of Sinuxylamide B, C, and D upon feeding of L-Tyr and *p*-coumaric acid (*p*-CA). * Proposed allene analogs of sinuxylamides. **c**. Overexpression of NseAB in *A. nidulans* without feeding of L-Tyr and *p*-CA resulted in the detection of compound **16**, suggesting that *O*-but-2-ynyl L-Tyr (**6**) was metabolized to **16** through an endogenous pathway.

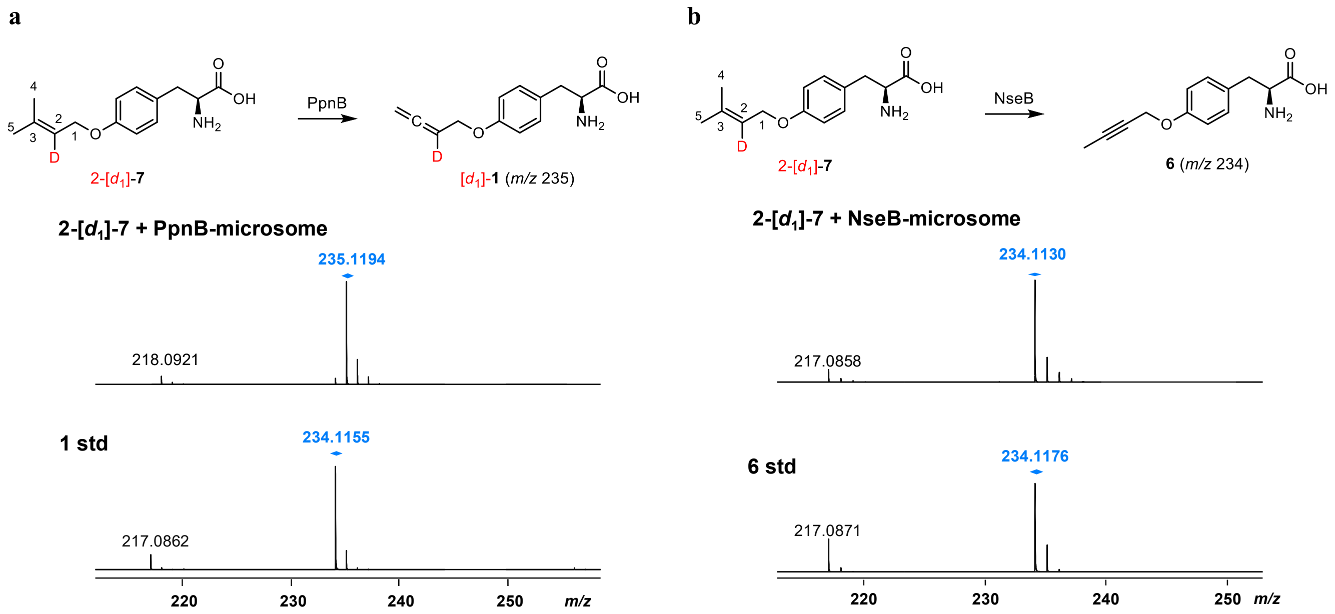


**Supplementary Fig. 20 | LC–QTOF analysis of *A. nidulans* microsomes containing PpnB or NseB incubated with 2-[$d_1$]-7.** +1 Da mass shift was observed for product **1** when 2-[$d_1$]-**7** was supplied in the microsomal assays containing PpnB. The deuterium was lost for product **6** when 2-[$d_1$]-**7** was supplied in the microsomal assays containing NseB.

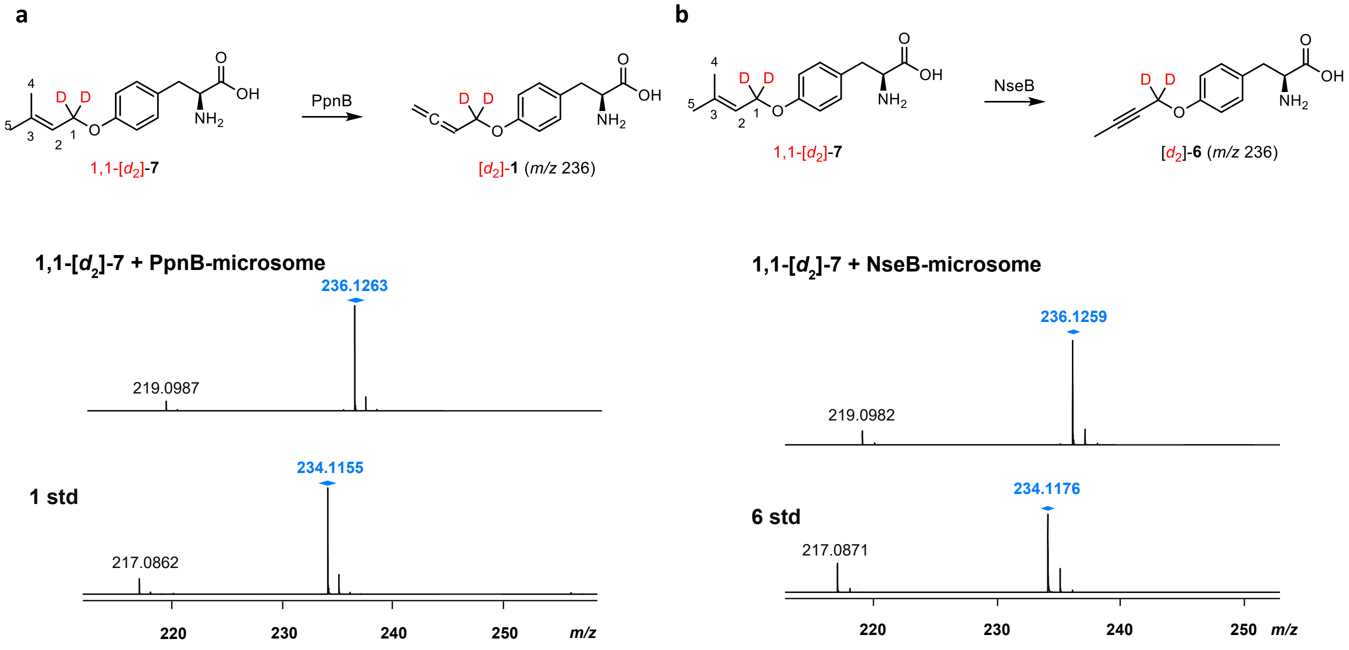


**Supplementary Fig. 21 | LC–QTOF analysis of *A. nidulans* microsomes containing PpnB or NseB incubated with 1,1-[$d_2$]-7.** +2 Da mass shift was observed for products **1** and **6** when 1,1-[$d_2$]-**7** was supplied in the microsomal assays, supporting retention of the hydrogen atoms on the prenyl methylene group during PpnB- and NseB-catalyzed transformations.

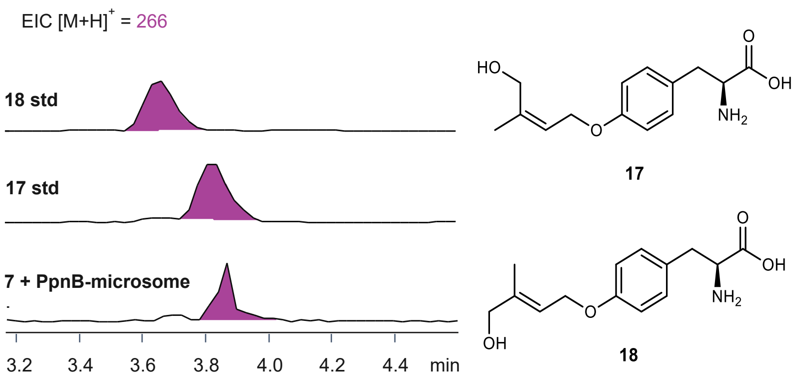


**Supplementary Fig. 22 | Identification of intermediate 17 by incubating *O*-prenyl L-Tyr (7) with *ppnB*.** Compound **17** was detected based on the retention time match with the synthetic standard. This further confirms that the **17** is a bona fide intermediate in the pathway.

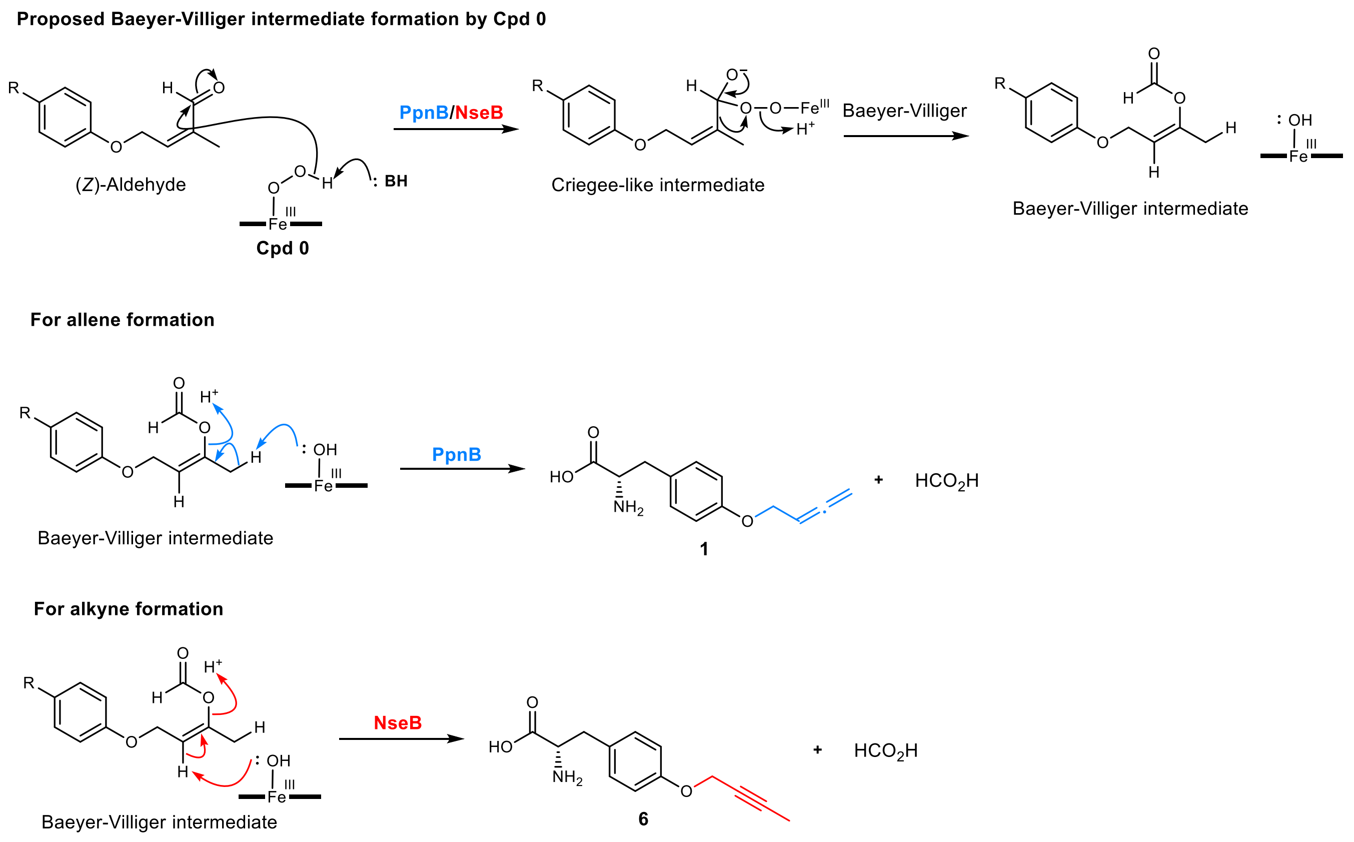


**Supplementary Fig. 23 | Alternative proposed mechanism of allene and alkyne formation via Baeyer-Villiger (BV) intermediate.** Based on the proposed mechanism of the CYP51-catalyzed reaction involving a BV intermediate, analogous reaction mechanism for PpnB- and NseB-catalyzed allene and alkyne formation can also be proposed.

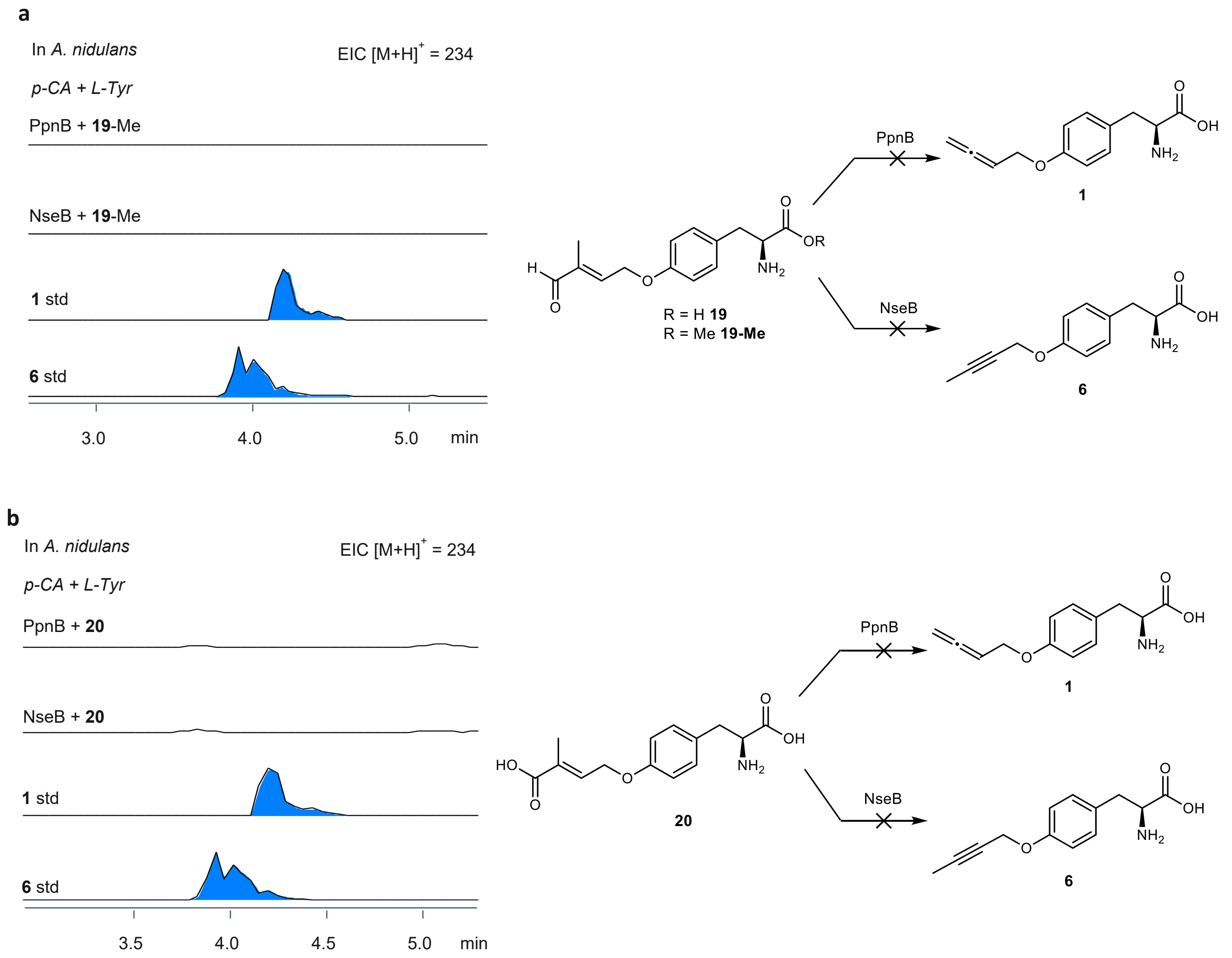


**Supplementary Fig. 24 | Compounds 15 and 16 are not the intermediate toward 1 or 6.** Feeding compounds **19** (**a**) or **20** (**b**) to *A. nidulans* expressing PpnB or NseB did not result in production of **1** or **6**, indicating that neither **19** nor **20** functions as a biosynthetic intermediate. These results indirectly suggest that the pathway leading to **1** and **6** must proceed through the *Z*-aldehyde or *Z*-carboxylic acid intermediates; however, chemical synthesis of these intermediates resulted in isomerization to *E* configuration, preventing direct verification.

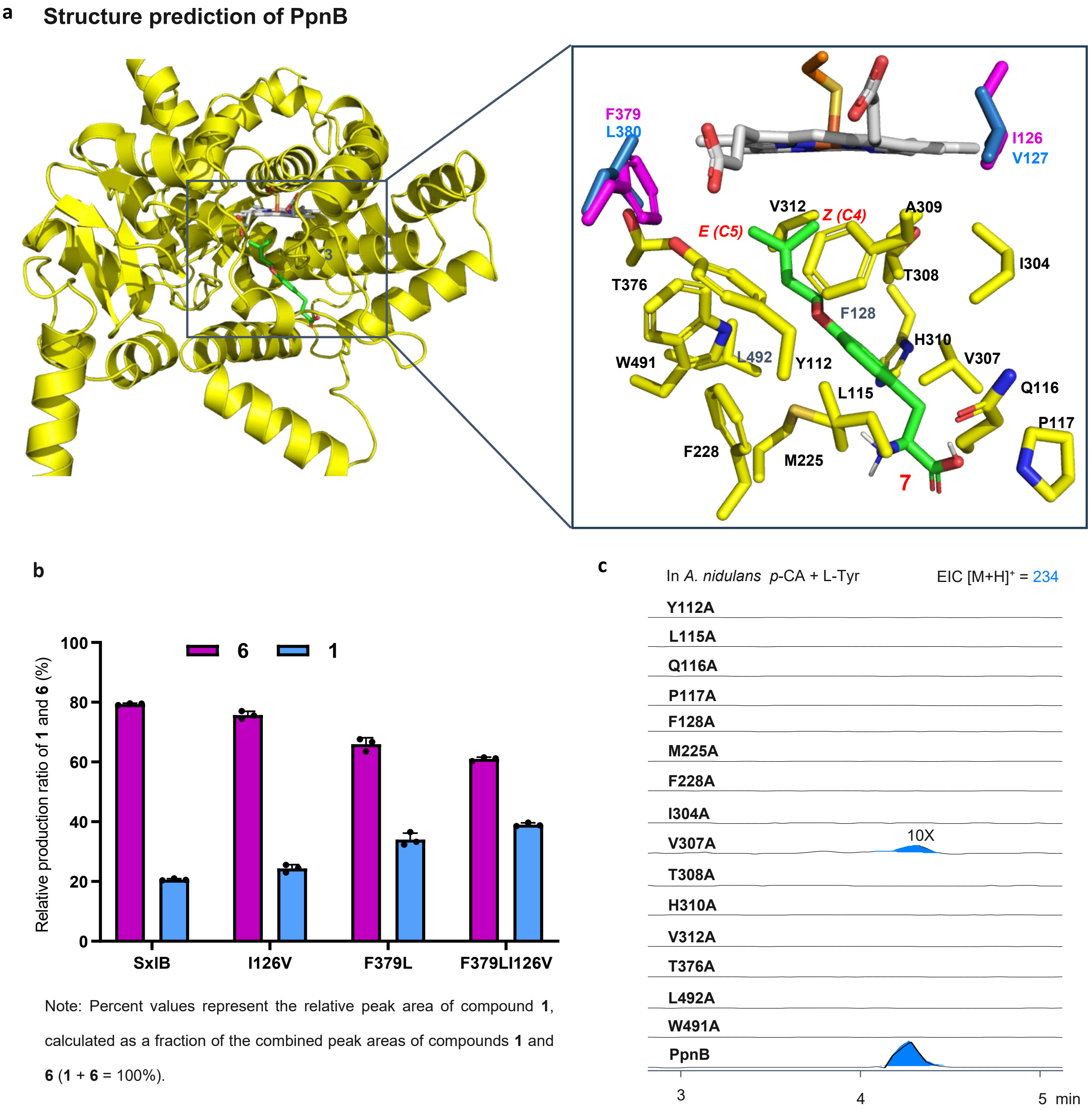


**Supplementary Fig. 25 | Comparison of residues forming substrate pocket between PpnB and NseB. a.** Docking simulation of **7** in the AlphaFold 3[28] predicted structure of PpnB containing heme cofactor. The active site was performed using Autodock Vina[29]. One representative binding mode is shown, with **7** rendered in green and the heme cofactor in grey. PpnB and NseB share high overall sequence identity and display nearly identical substrate-binding pockets, consistent with their conserved active-site architectures. The only differences within the pocket occur at two positions: NseB contains **I126** and **F379** (magenta), whereas PpnB carries the corresponding **V127** and **L380** residues (blue). **b**. Extracted ion chromatograms (EICs) showing the production of **1** by wild-type NseB and the I126V, F379L, and I126V/F379L mutants. Values and error bars represent the average and s.d. of three independent replicates (black filled circles), respectively ($n = 3$). The relative peak area of **1** increase from 19% in wild-type NseB to 24% in I126V, 37% in F379L, and 38% in the I126V/F379L double mutant. Differences in endogenous metabolism of **1** and **6** were not considered. Although mutation at these two positions enhanced allene formation, the double mutation did not completely shift the product profile to match PpnB. This suggests that additional distal residues also contribute to shaping the active-site pocket and controlling product selectivity. **c**. Alanine-scanning mutagenesis of the PpnB substrate-binding pocket. The mutants were coexpressed with ppnA in *A. nidulans* supplemented with *p*-CA and L-Tyr. Most alanine substitutions abolished the production of **1**, indicating that residues surrounding the substrate-binding pocket are important for substrate binding and product formation.

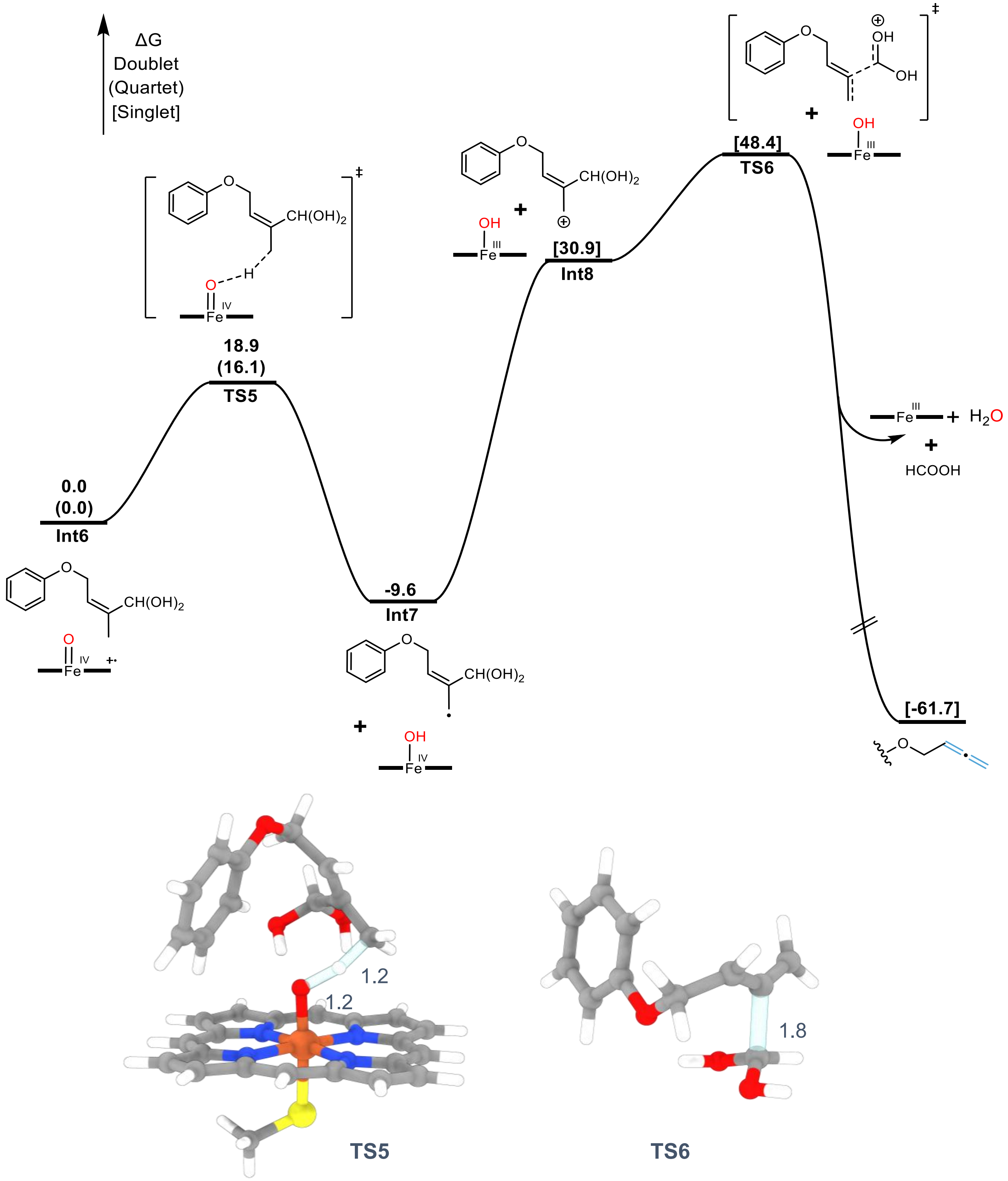


**Supplementary Fig. 26 | Computed energy profile for the potential cationic allene formation mechanism.** The energies were calculated by B3LYP-D3(BJ)/def2-TZVP/SMD($Et_2O$)//B3LYP-D3(BJ)/def2-SVP/IEFPCM($Et_2O$) at 298

K, 1 atm and are given in kcal/mol. Bond distances are labeled in angstrom. For all iron-containing species, possible spin states with free energy are reported. As shown in the Figure, another potential Cpd I mechanism starts from the H-abstraction from the alylic C($sp^3$)-H by Cpd I *via* **TS5** to yield an alylic radical **Int7** and Cpd II. **Int7** is proposed to have a subsequently single electron transfer (SET) that oxidizing itself to alylic cation **Int8** and concurrently reducing Cpd II to Fe(III)-OH. **Int8** then fragments through **TS6** to give the allene product and protonated formic acid, which can react with Fe(III)-OH to restore the Fe(III) state. However, this mechanism is precluded by the highly unfavorable SET process, which is endergonic by 40.5 kcal/mol.

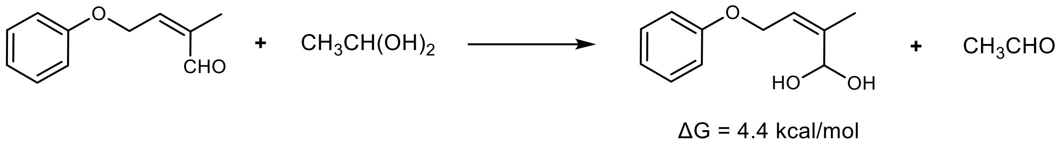


**Supplementary Fig. 27 | Calculated free-energy difference between the gem-diol and aldehyde plus water.** The aldehyde substrate hydration free energy is calculated using the isodesmic reaction developed by Julio Casado *et al*[30]. The energies are calculated by B3LYP-D3(BJ)/def2-TZVP/SMD($H_2O$)//B3LYP-D3(BJ)/def2-SVP/IEFPCM($H_2O$) level of theory. The absolute hydration free energy of the aldehyde substrate is deduced from the experimental hydration equilibrium constant for acetaldehyde, which is measured to be 1.07 at room temperature[31,32]:

$$\Delta G_{rxn} = \Delta G_{substate} - \Delta G_{CH3CHO}$$

$$\Delta G_{CH3CHO} = -RTlnK_{hyd} = -0.03\ kcal/mol$$

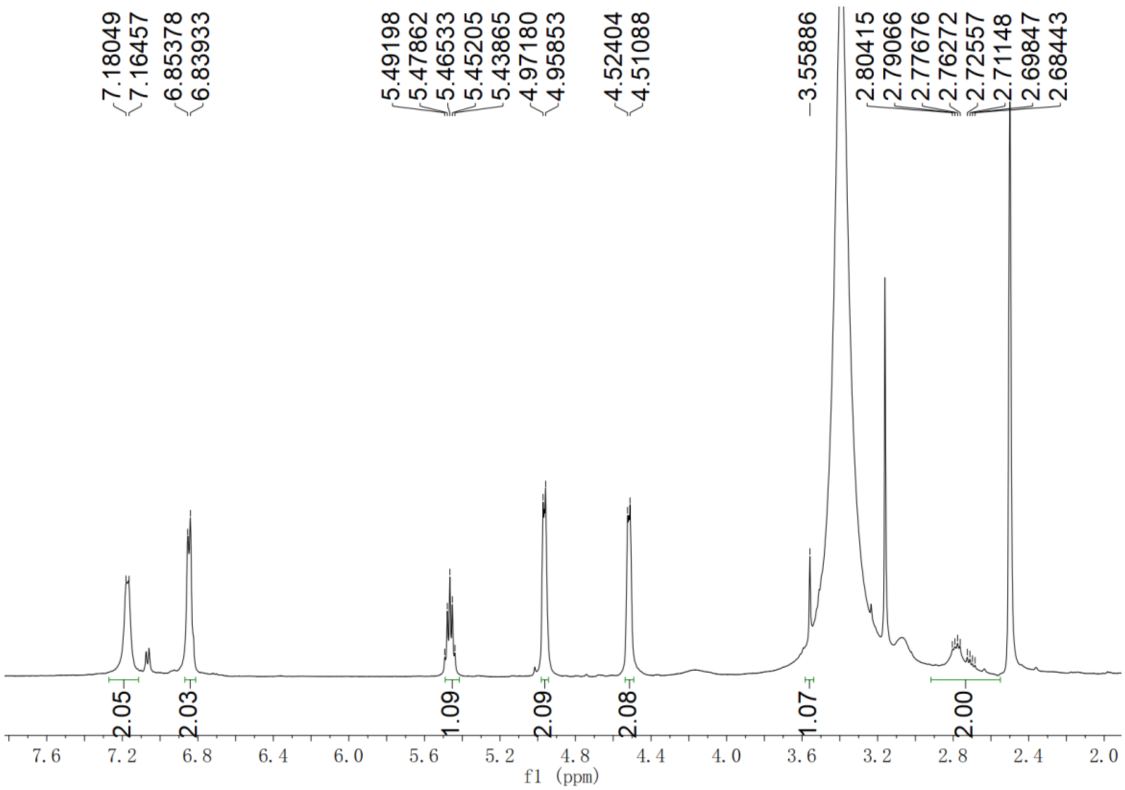


**Supplementary Fig. 28.** $^{1}$H NMR spectrum of *O*-homoallenyl-L-Tyr (**1**) in DMSO-$d_6$

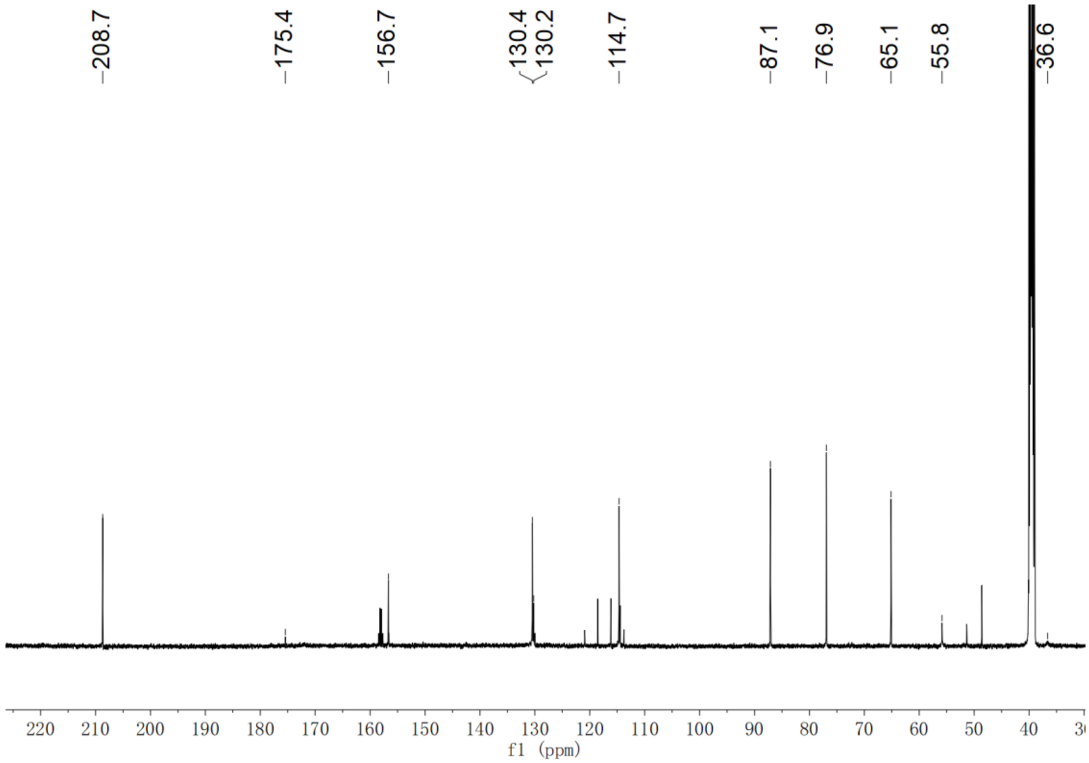


**Supplementary Fig. 29.** $^{13}$C NMR spectrum of *O*-homoallenyl-L-Tyr (**1**) in DMSO-$d_6$

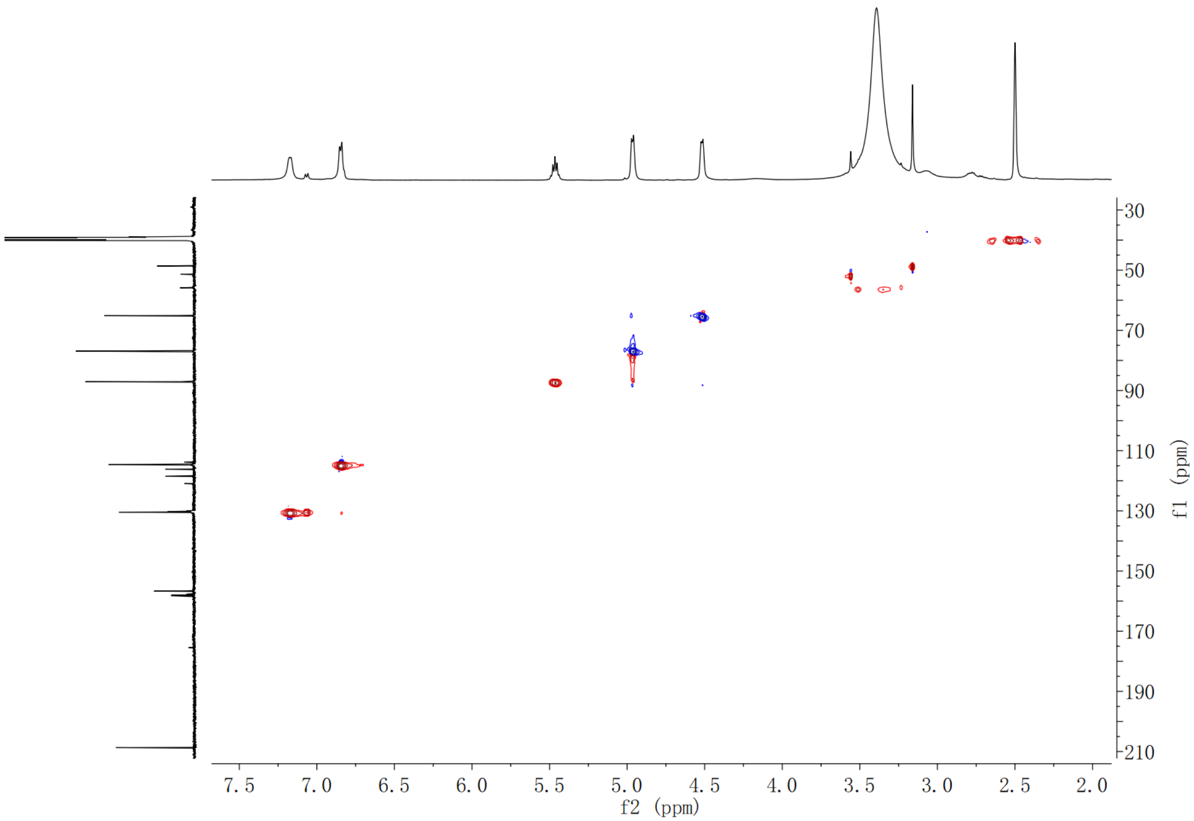


**Supplementary Fig. 30.** HSQC spectrum of *O*-homoallenyl- L-Tyr (**1**) in DMSO-$d_6$

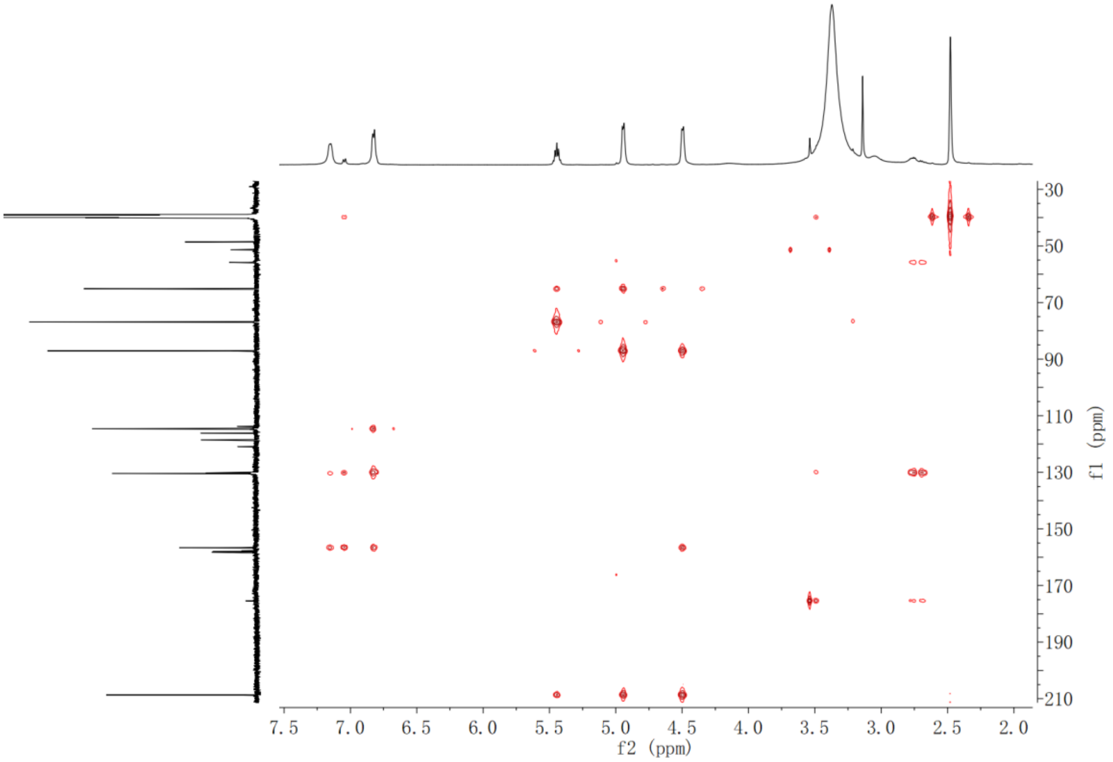


**Supplementary Fig. 31.** HMBC spectrum of *O*-homoallenyl- L-Tyr (**1**) in DMSO-$d_6$

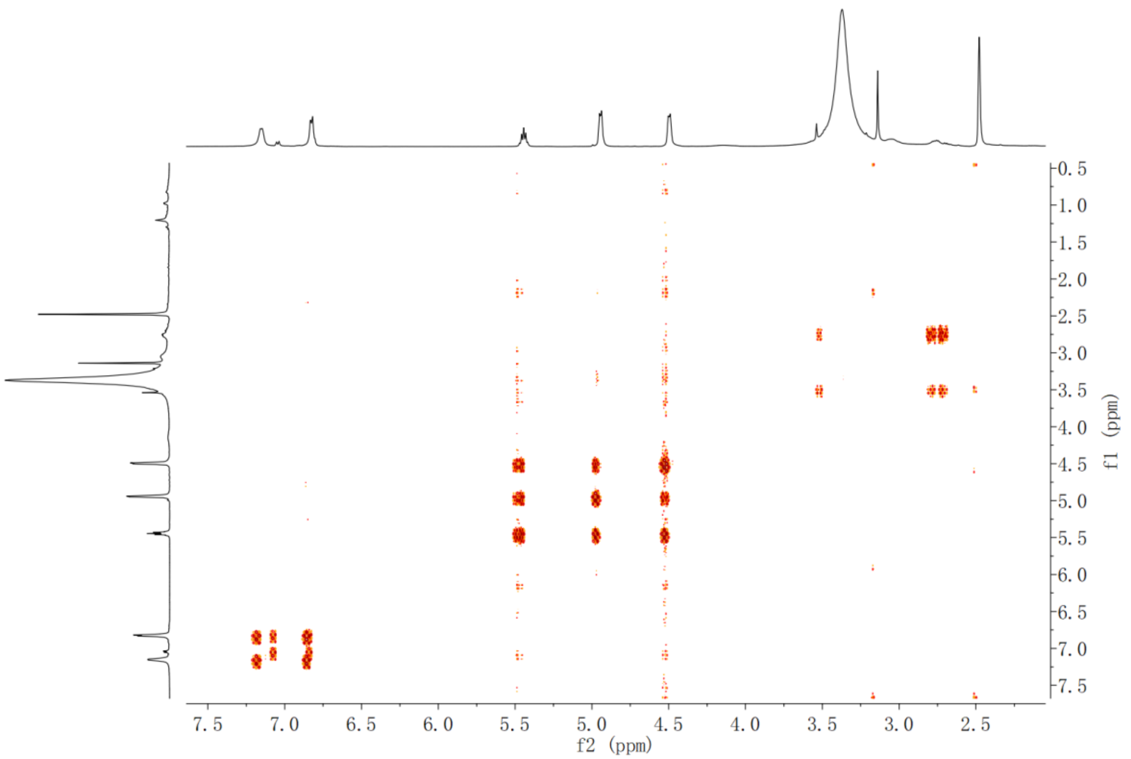


**Supplementary Fig. 32.** $^{1}H$-$^{1}H$ COSY spectrum of *O*-homoallenyl- L -Tyr (**1**) in DMSO-d6

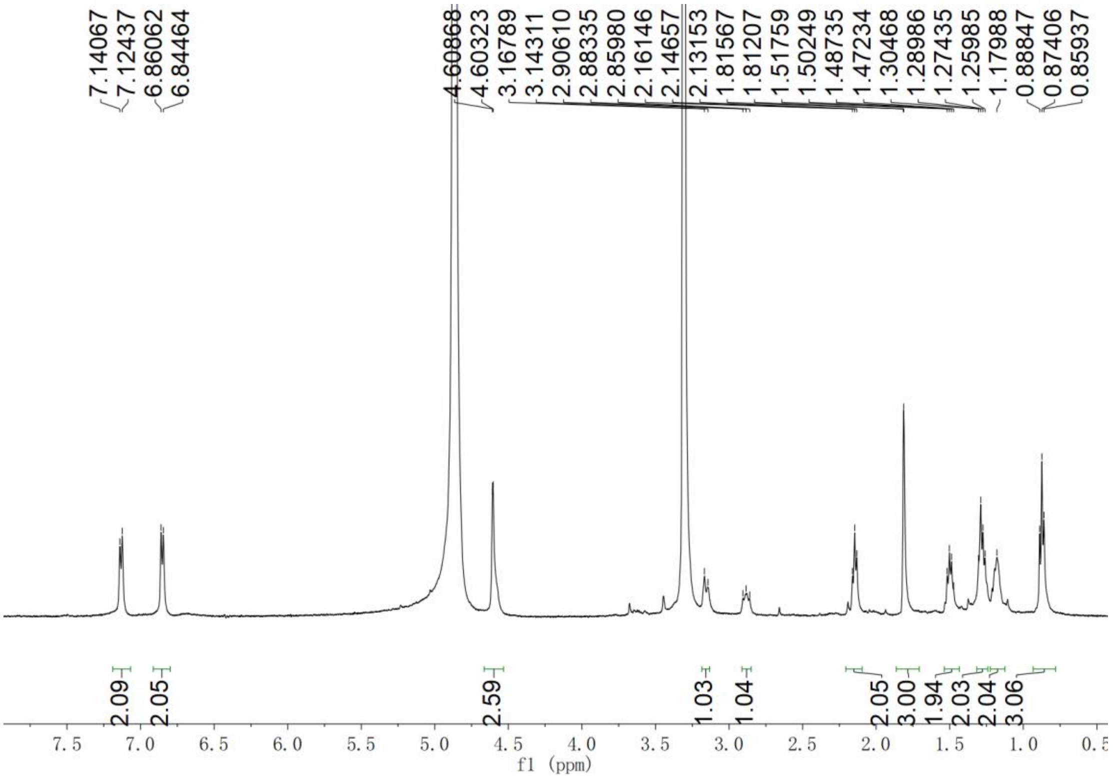


**Supplementary Fig. 33.** $^{1}H$ NMR spectrum of Sinuxylamide B **(2)** in $CD_3OD$

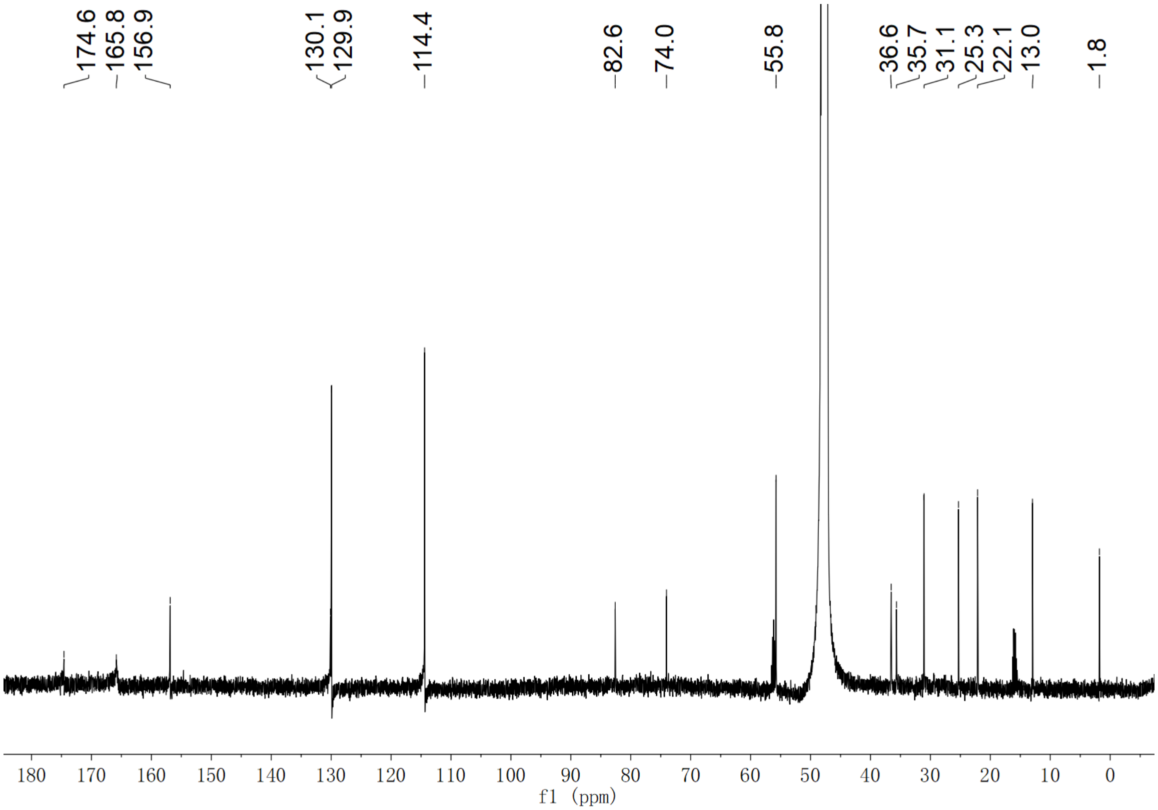


**Supplementary Fig. 34.** $^{13}$C NMR spectrum of Sinuxylamide B **(2)** in $CD_3OD$

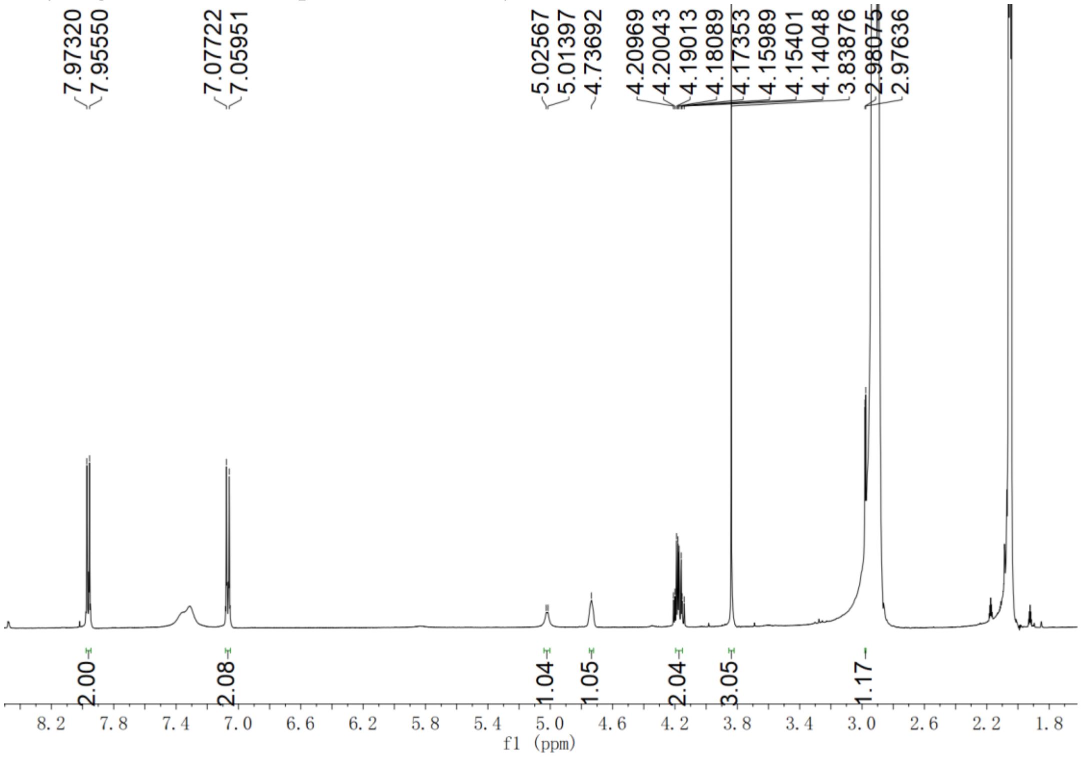


**Supplementary Fig. 35.** $^{1}$H NMR spectrum of penipratynolene (**5**) in acetone-$d_6$

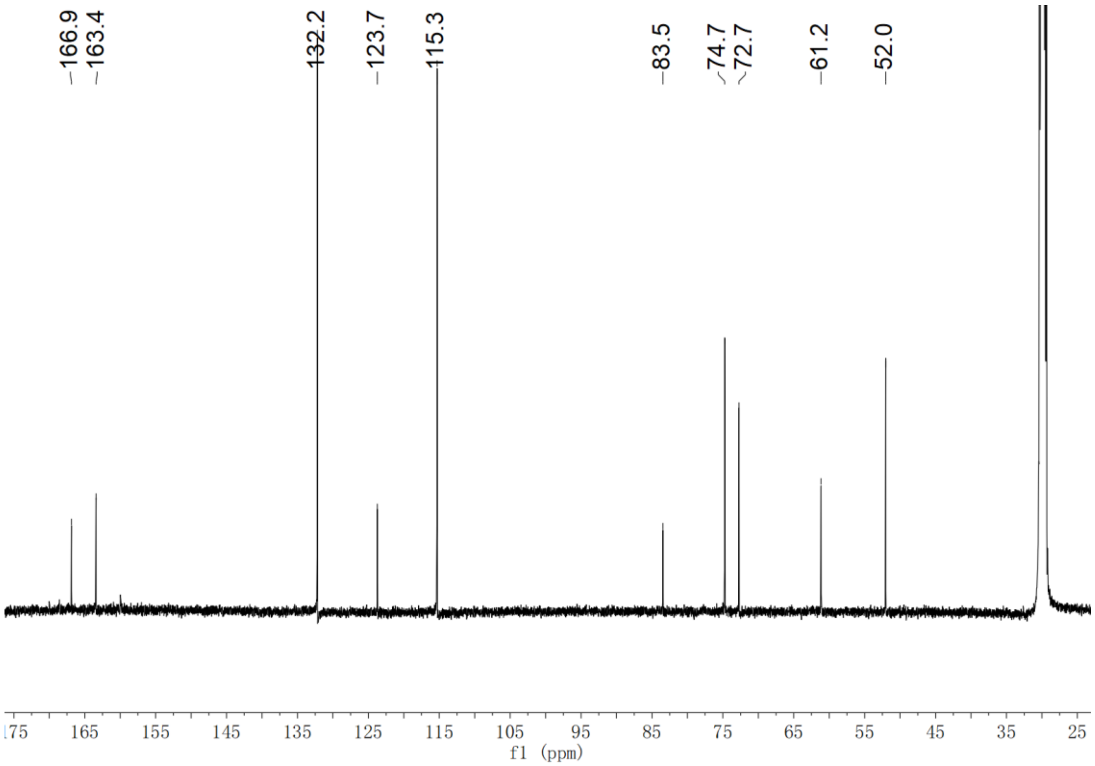


**Supplementary Fig. 36.** $^{13}C$ NMR spectrum of penipratynolene (**5**) in acetone-$d_6$

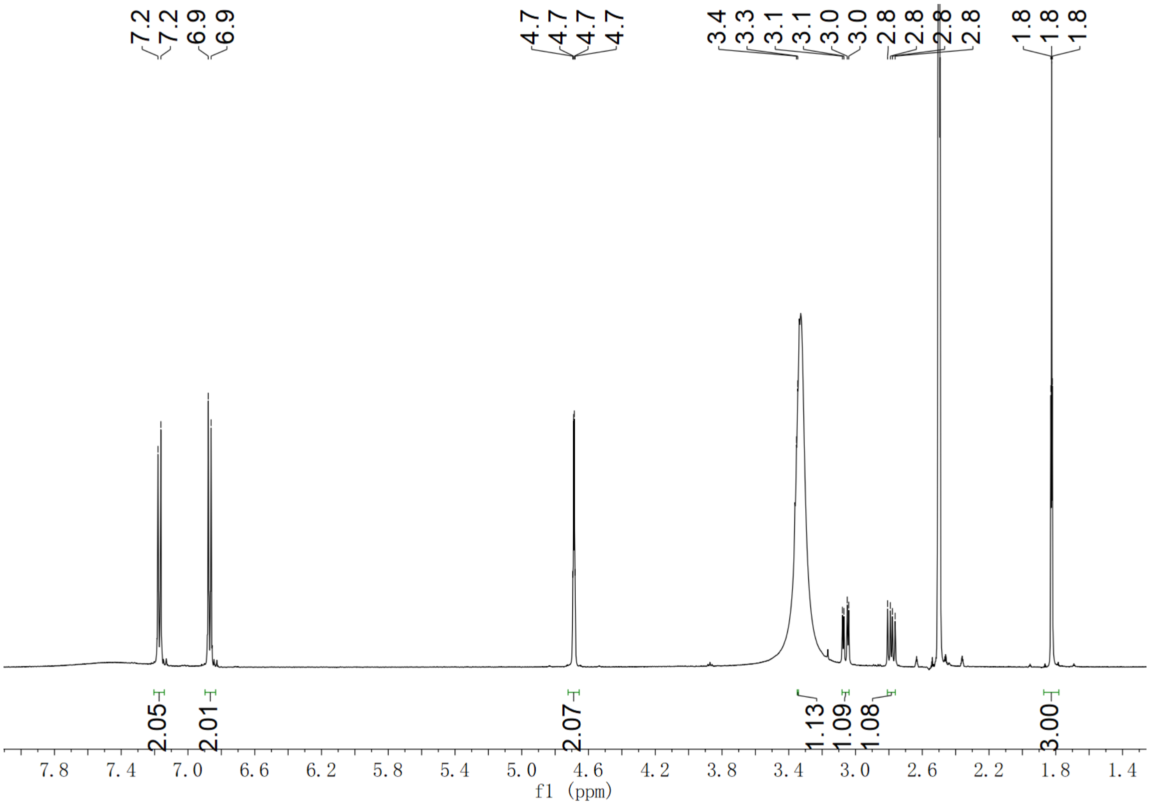


**Supplementary Fig. 37.** $^{1}H$ NMR spectrum of *O*-2-butyn-1-yl L-Tyr (**6**) in DMSO-$d_6$

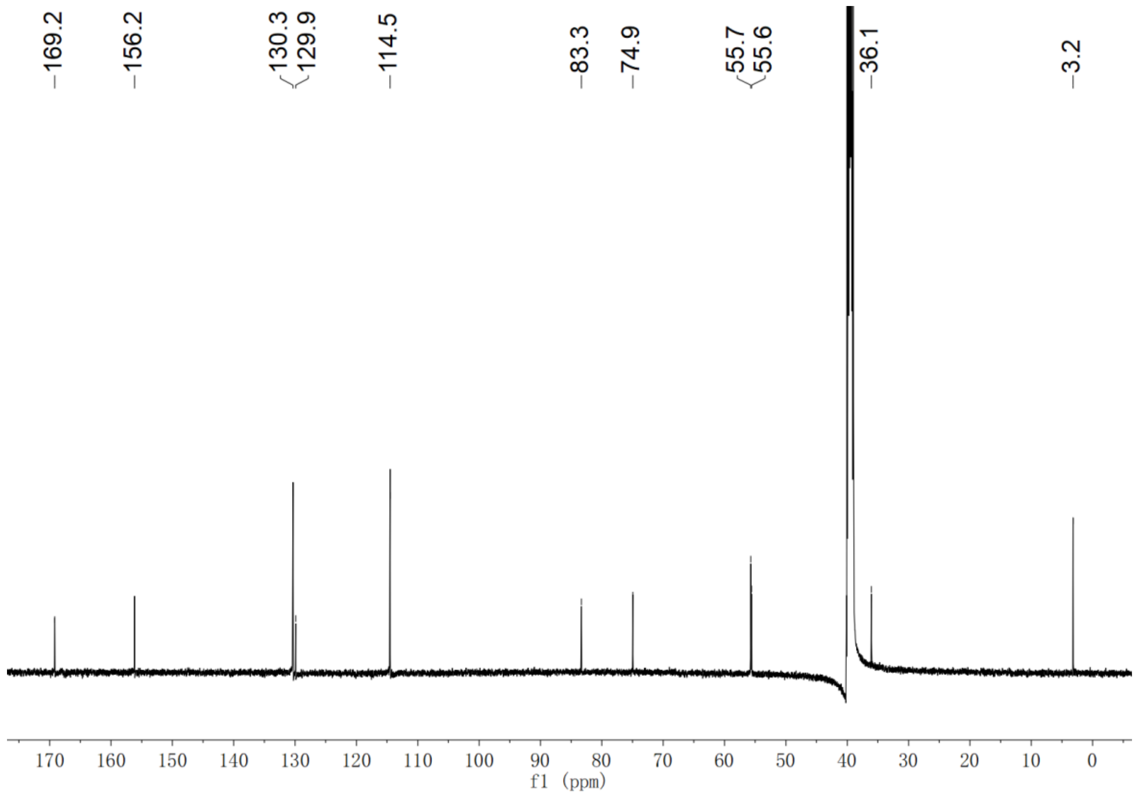


**Supplementary Fig. 38.** [13]C NMR spectrum of *O*-2-butyn-1-yl L-Tyr (**6**) in DMSO-$d_6$

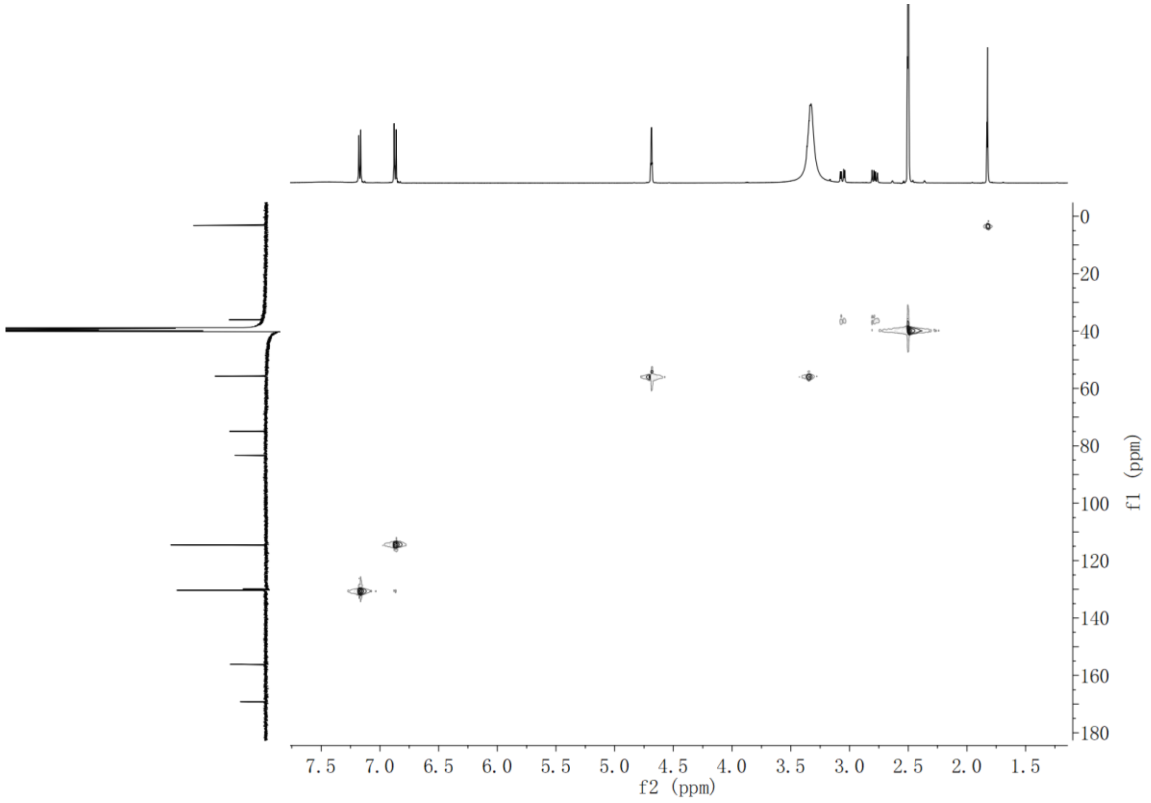


**Supplementary Fig. 39.** HSQC NMR spectrum of *O*-2-butyn-1-yl L-Tyr (**6**) in DMSO-$d_6$

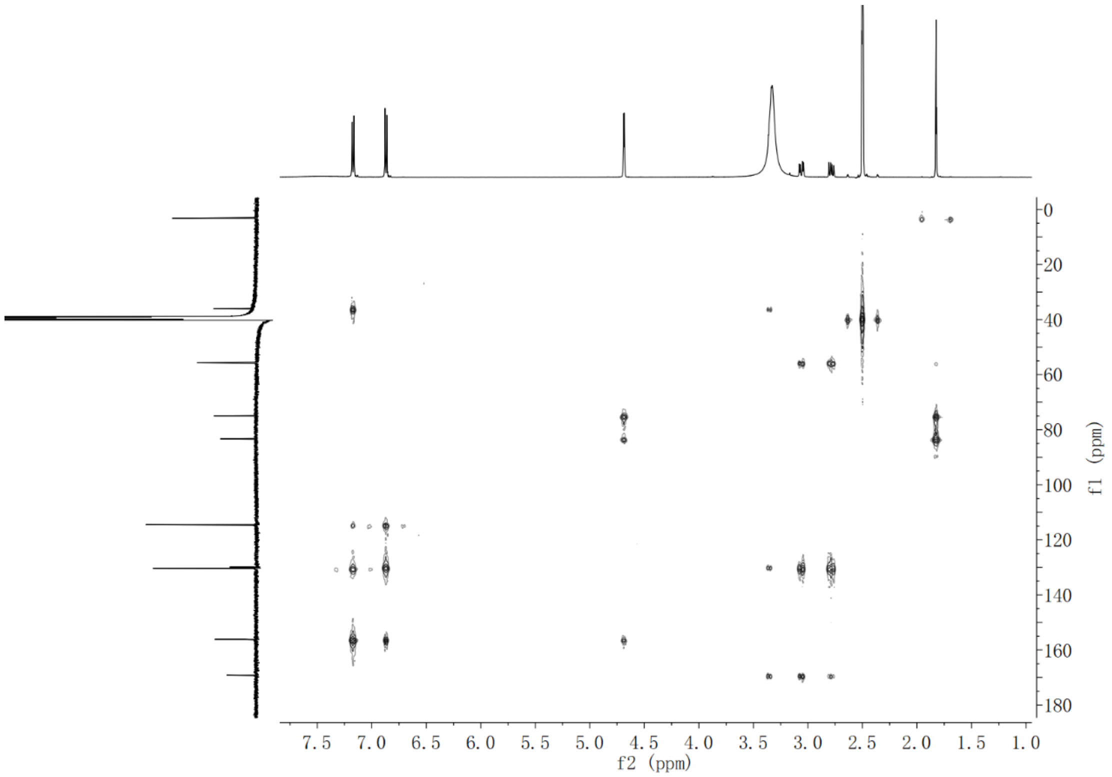


**Supplementary Fig. 40.** HMBC NMR spectrum of *O*-2-butyn-1-yl L-Tyr (**6**) in DMSO-$d_6$

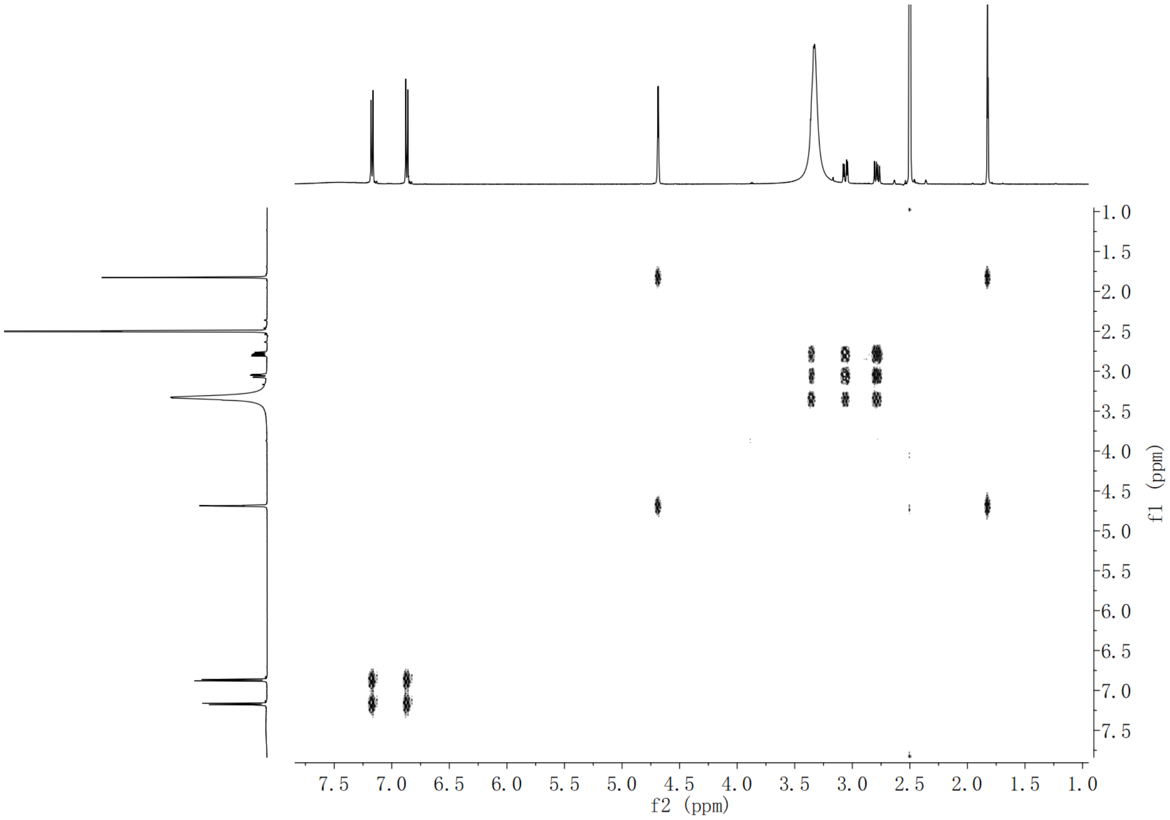


**Supplementary Fig. 41.** $^1$H-$^1$H COSY NMR spectrum of *O*-2-butyn-1-yl L-Tyr (**6**) in DMSO-$d_6$

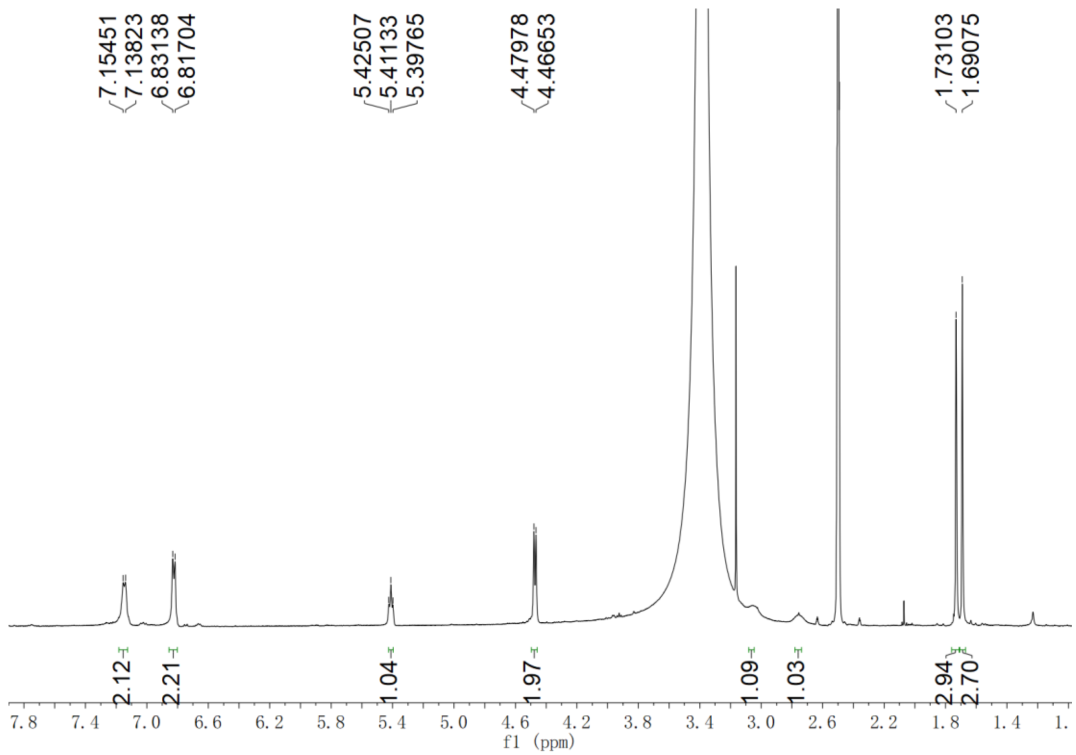


**Supplementary Fig. 42.** $^{1}$H NMR spectrum of *O*-Prenyl L-Tyr (**7**) in DMSO-$d_6$

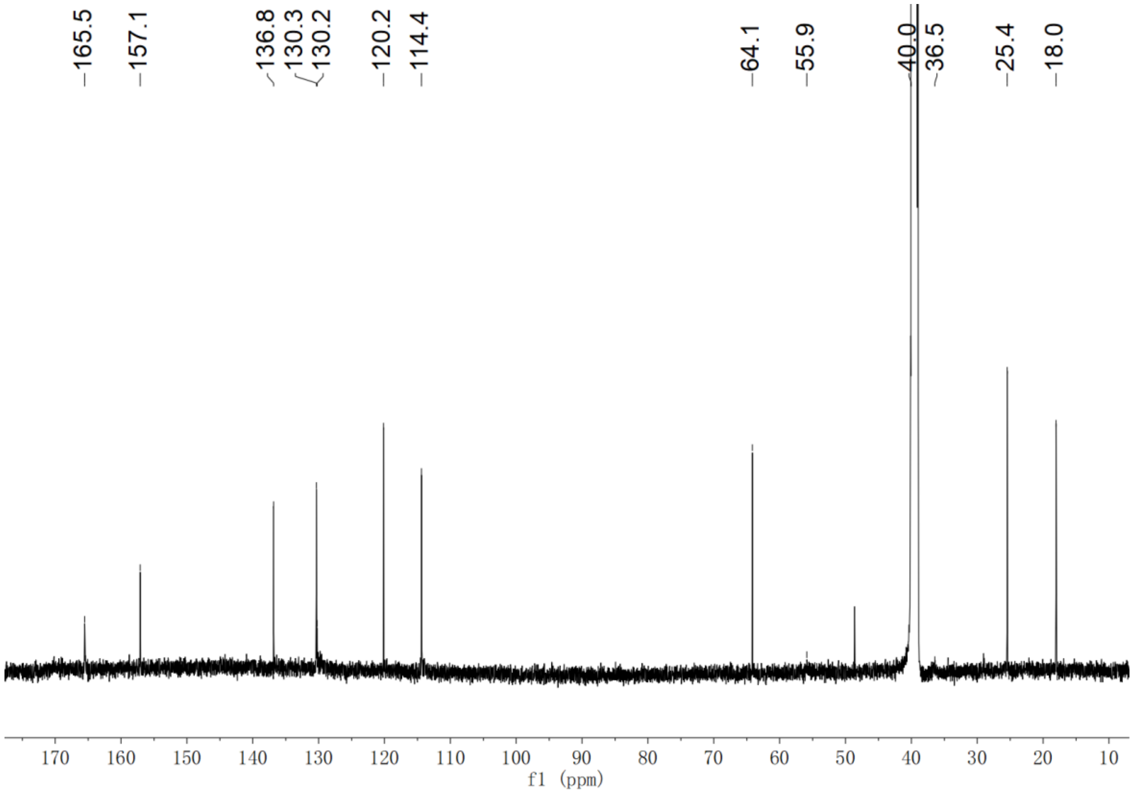


**Supplementary Fig. 43.** $^{13}$C NMR spectrum of *O*-Prenyl L-Tyr (**7**) in DMSO-$d_6$

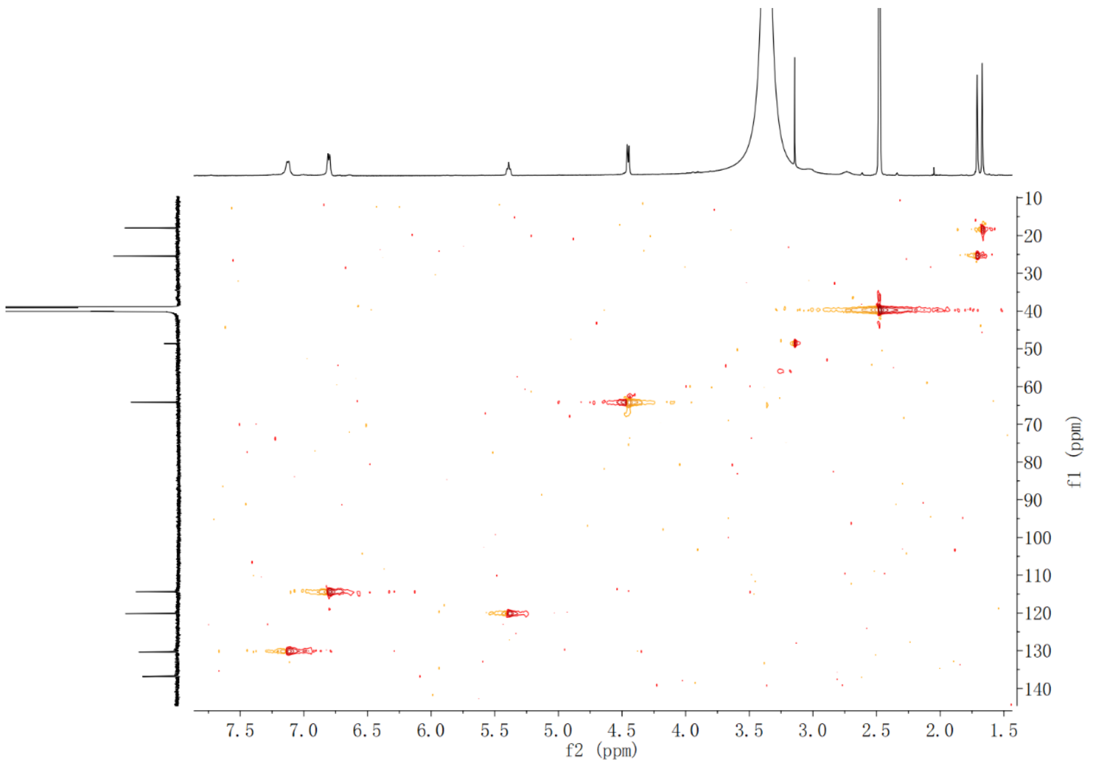


**Supplementary Fig. 44.** HSQC NMR spectrum of *O*-Prenyl L-Tyr (**7**) in DMSO-$d_6$

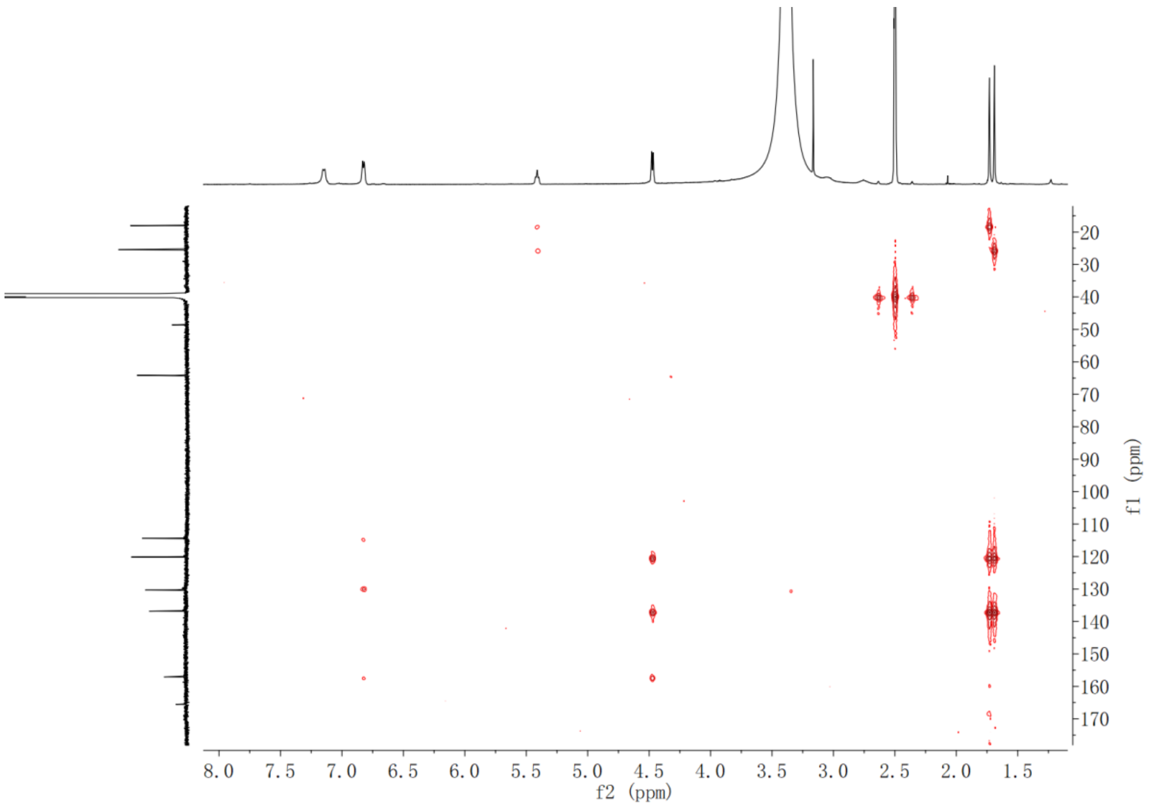


**Supplementary Fig. 45.** HMBC NMR spectrum of *O*-Prenyl L-Tyr (**7**) in DMSO-$d_6$

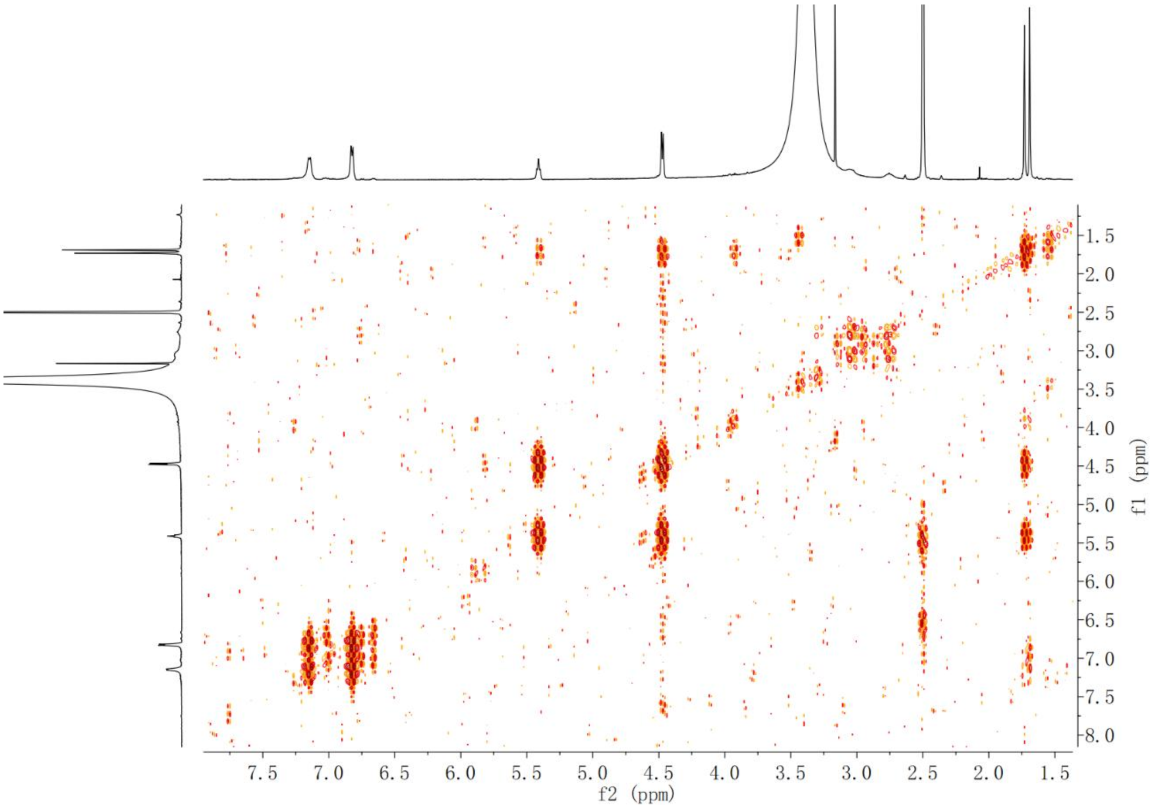


**Supplementary Fig. 46.** $^{1}$H-$^{1}$H COSY NMR spectrum of *O*-Prenyl L-Tyr (**7**) in DMSO-$d_6$

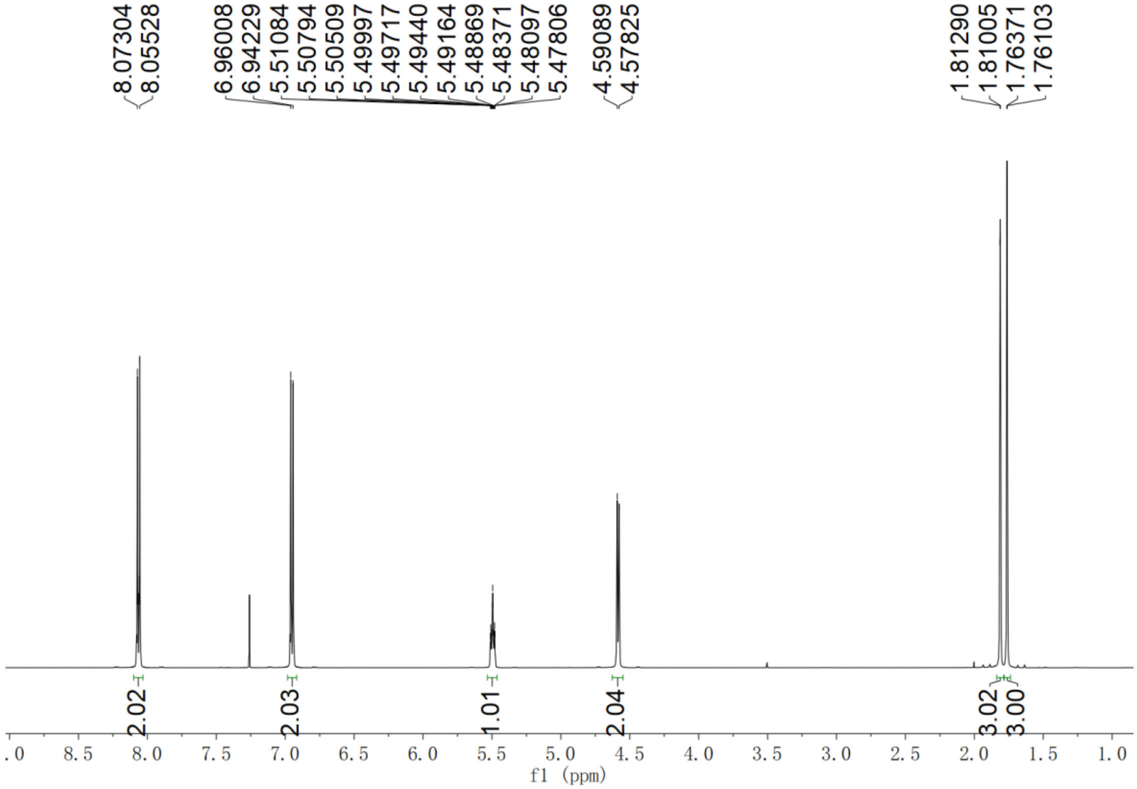


**Supplementary Fig. 47.** $^{1}$H NMR spectrum of *O*-prenyl-4-hydroxybenzoic acid (**8**) in $CDCl_3$

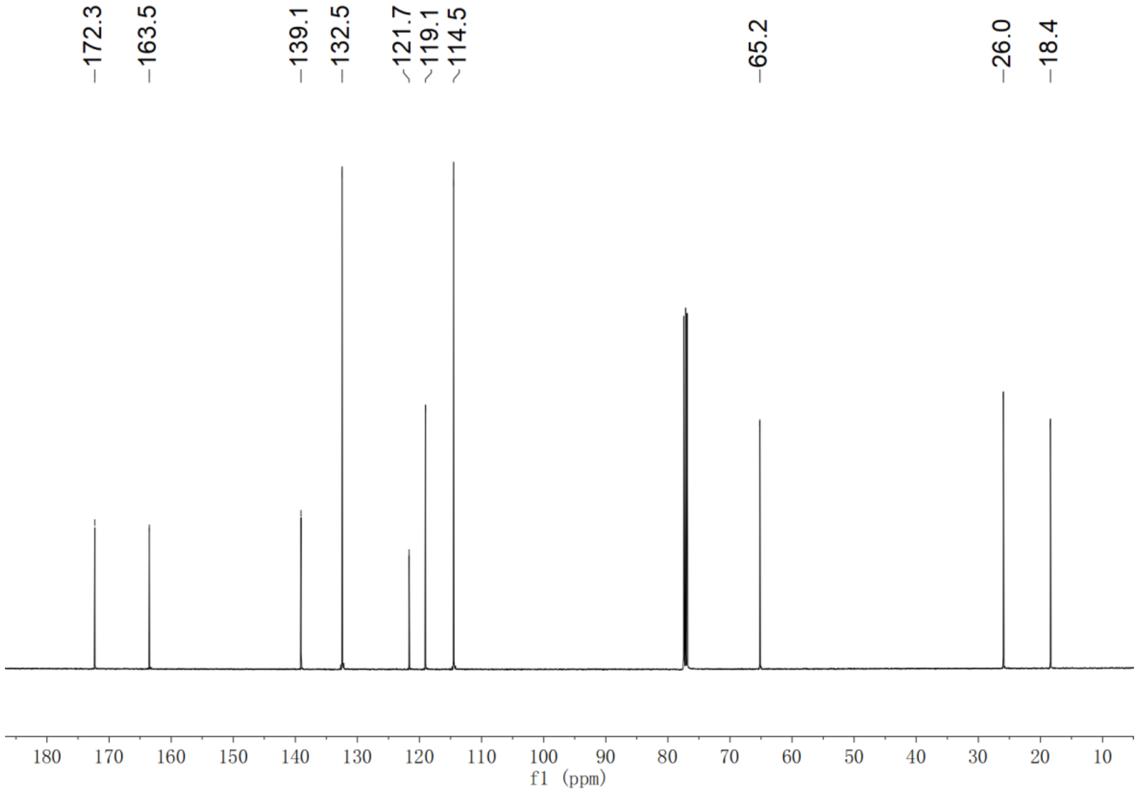


**Supplementary Fig. 48.** $^{13}C$ NMR spectrum of *O*-prenyl-4-hydroxybenzoic acid (**8**) in $CDCl_3$

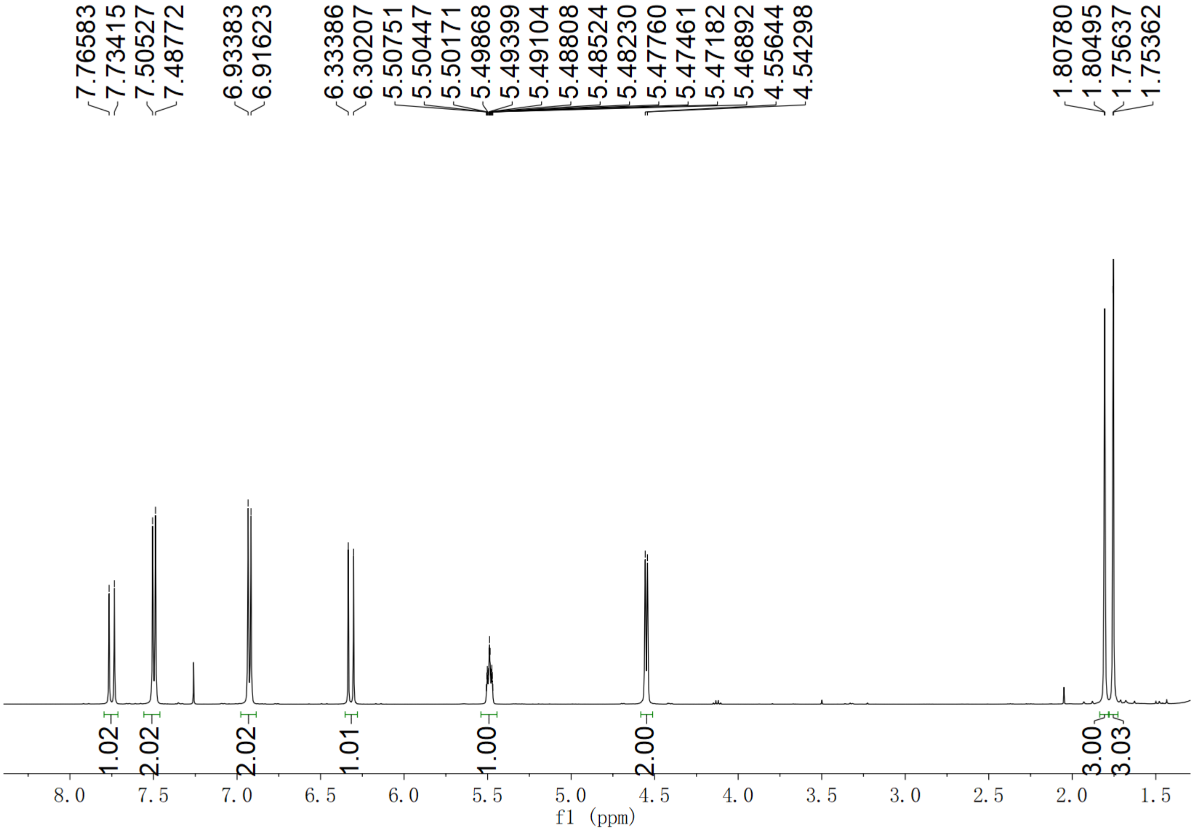


**Supplementary Fig. 49.** $^{1}H$ NMR spectrum of *O*-prenyl *p*-coumaric acid (**9**) in $CDCl_3$

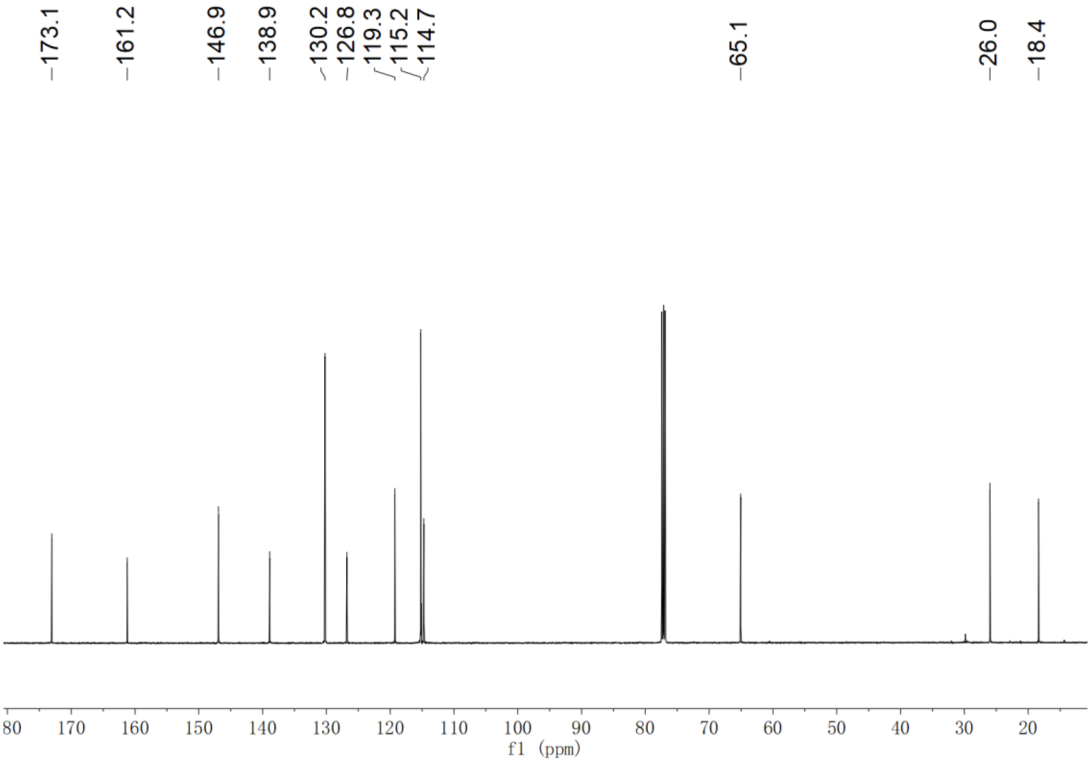


**Supplementary Fig. 50.** $^{13}$C NMR spectrum of *O*-prenyl *p*-coumaric acid (**9**) in $CDCl_3$

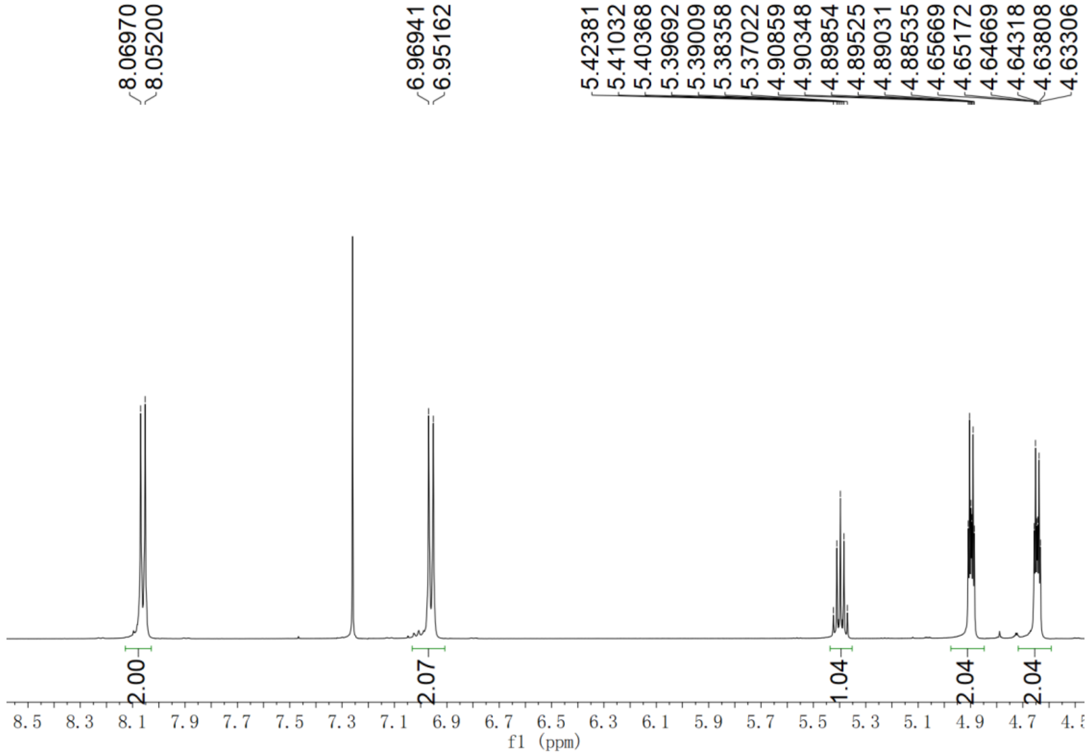


**Supplementary Fig. 51.** $^{1}$H NMR spectrum of *O*-homoallenyl 4-hydroxybenzoic acid (**10**) in $CDCl_3$

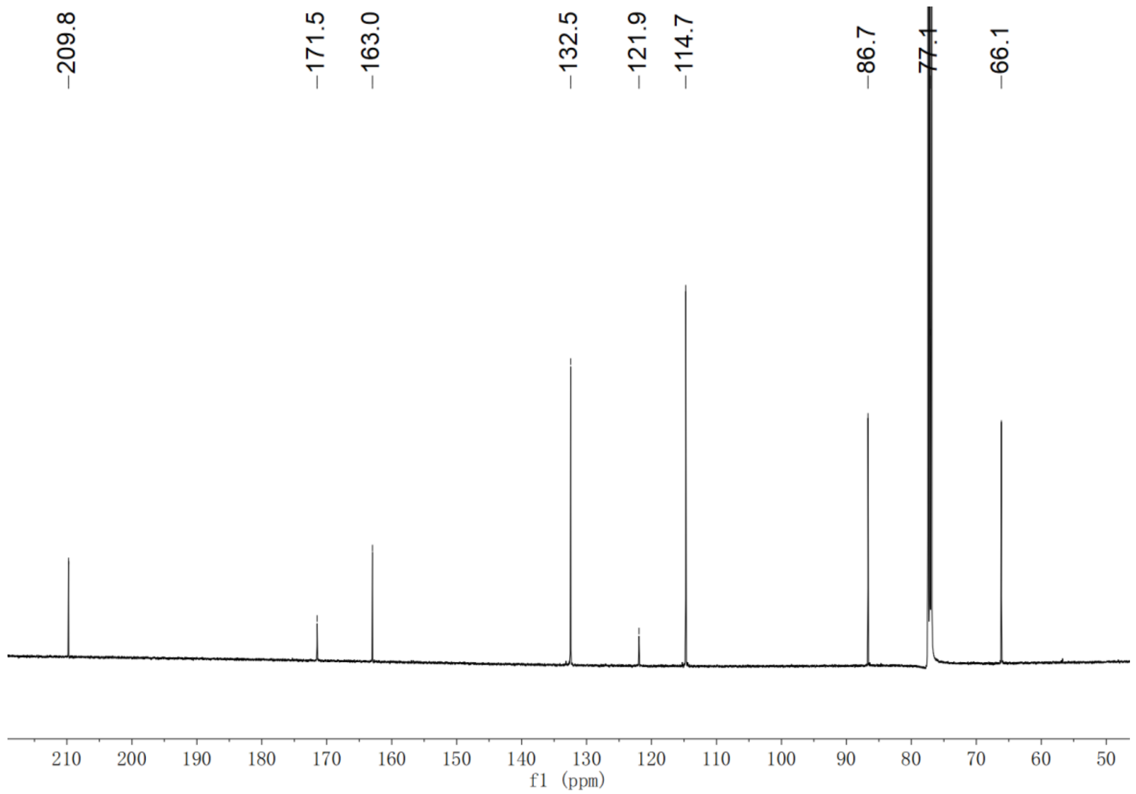


**Supplementary Fig. 52.** $^{13}$C NMR spectrum of *O*-homoallenyl 4-hydroxybenzoic acid (**10**) in $CDCl_3$

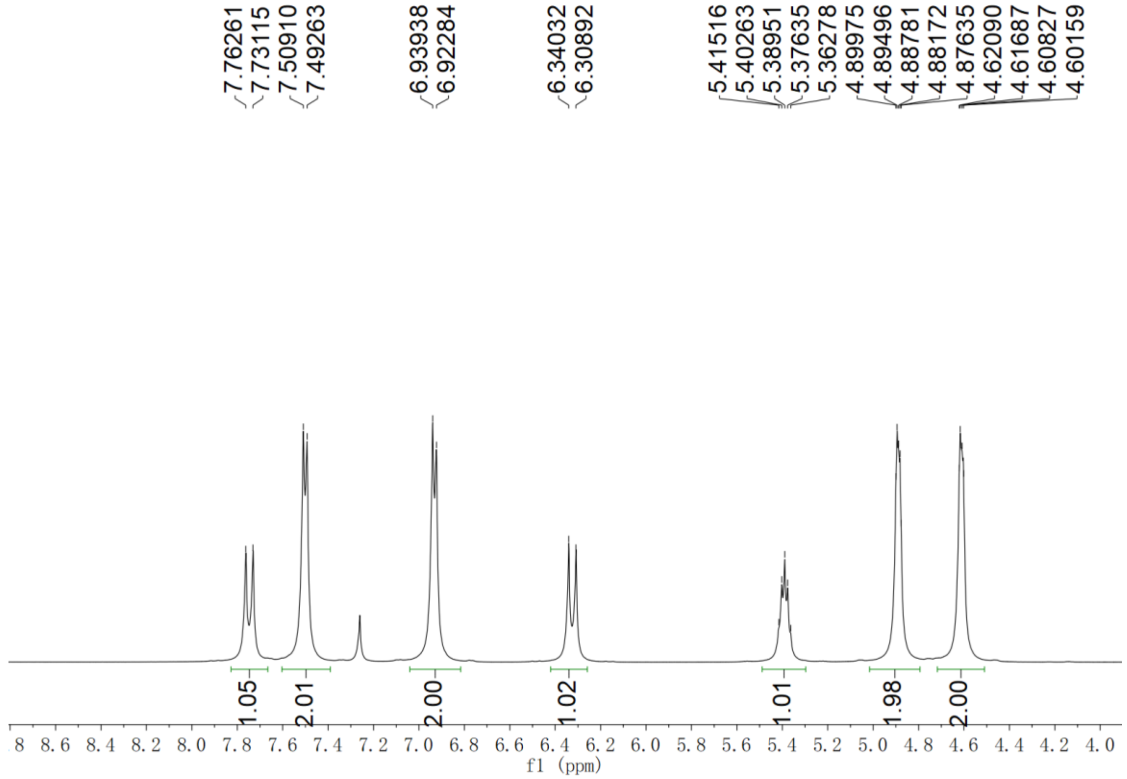


**Supplementary Fig. 53.** $^{1}$H NMR spectrum of *O*-homoallenyl *p*-coumaric acid (**11**) in $CDCl_3$

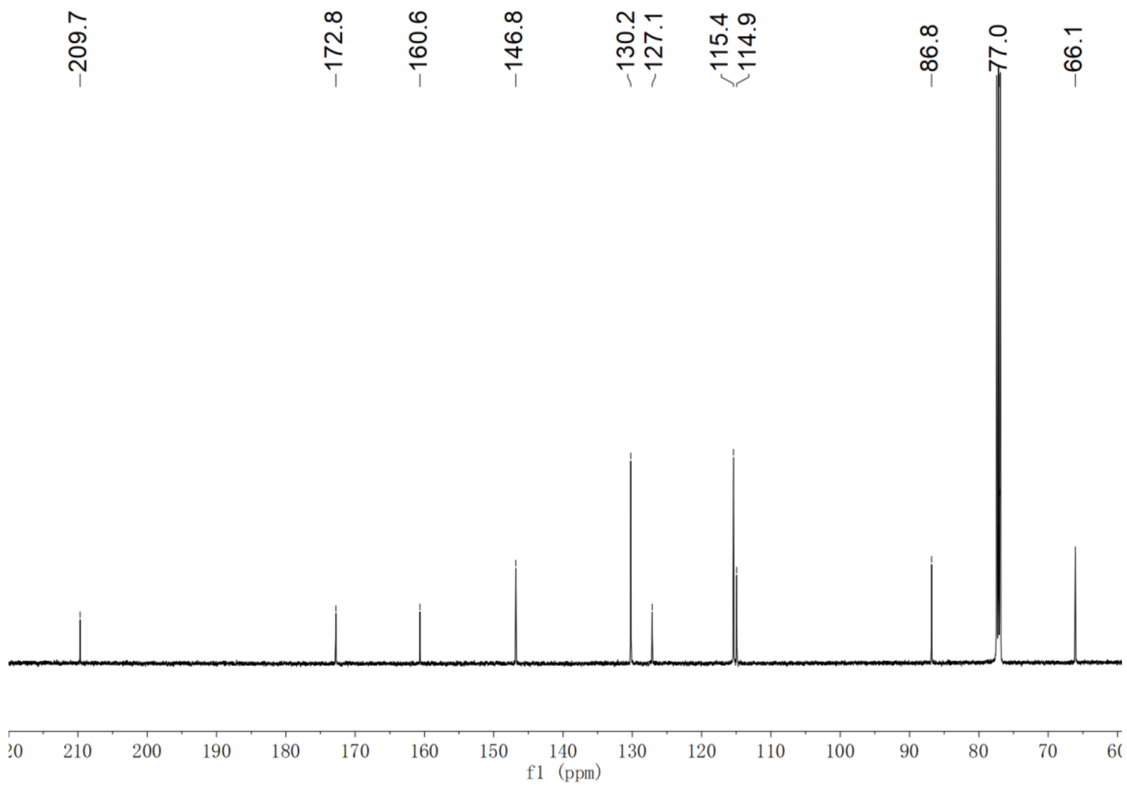

**Supplementary Fig. 54.** $^{13}C$ NMR spectrum of *O*-homoallenyl *p*-coumaric acid (**11**) in $CDCl_3$

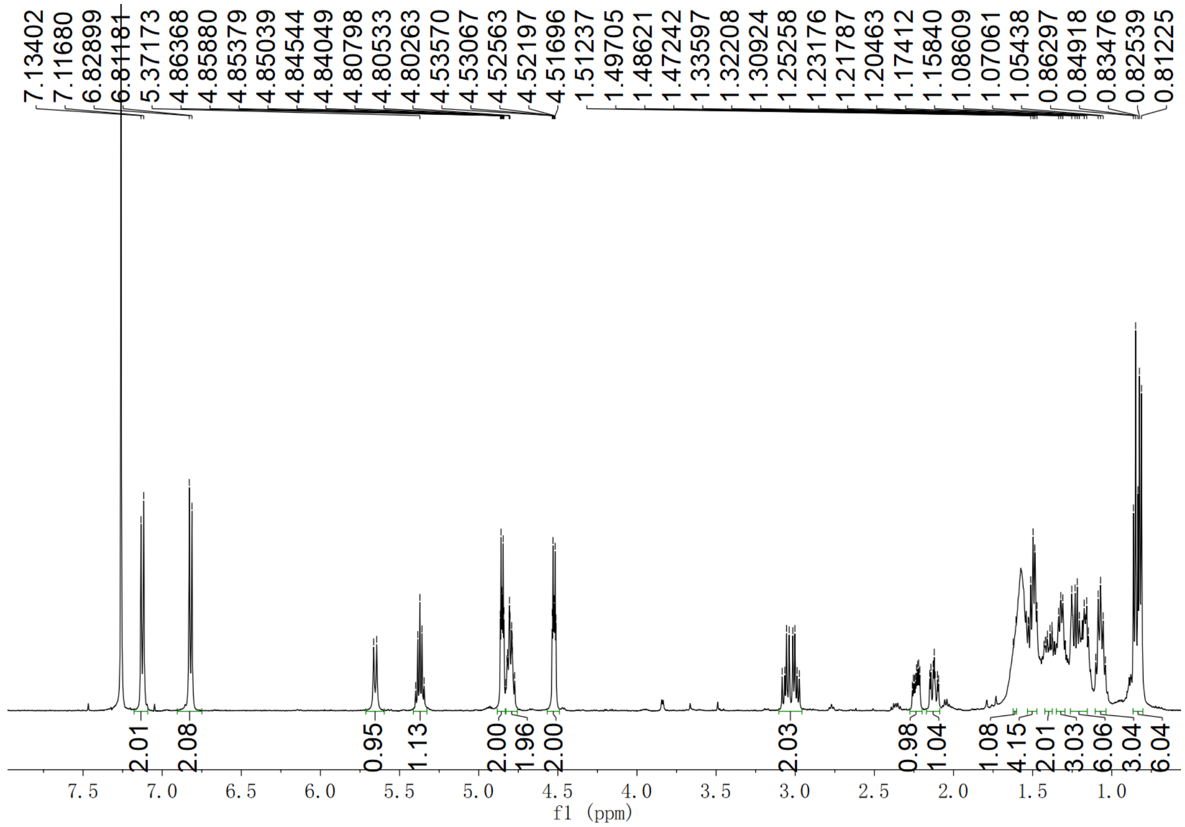

**Supplementary Fig. 55.** $^{1}H$ NMR spectrum of compound **13** in $CDCl_3$

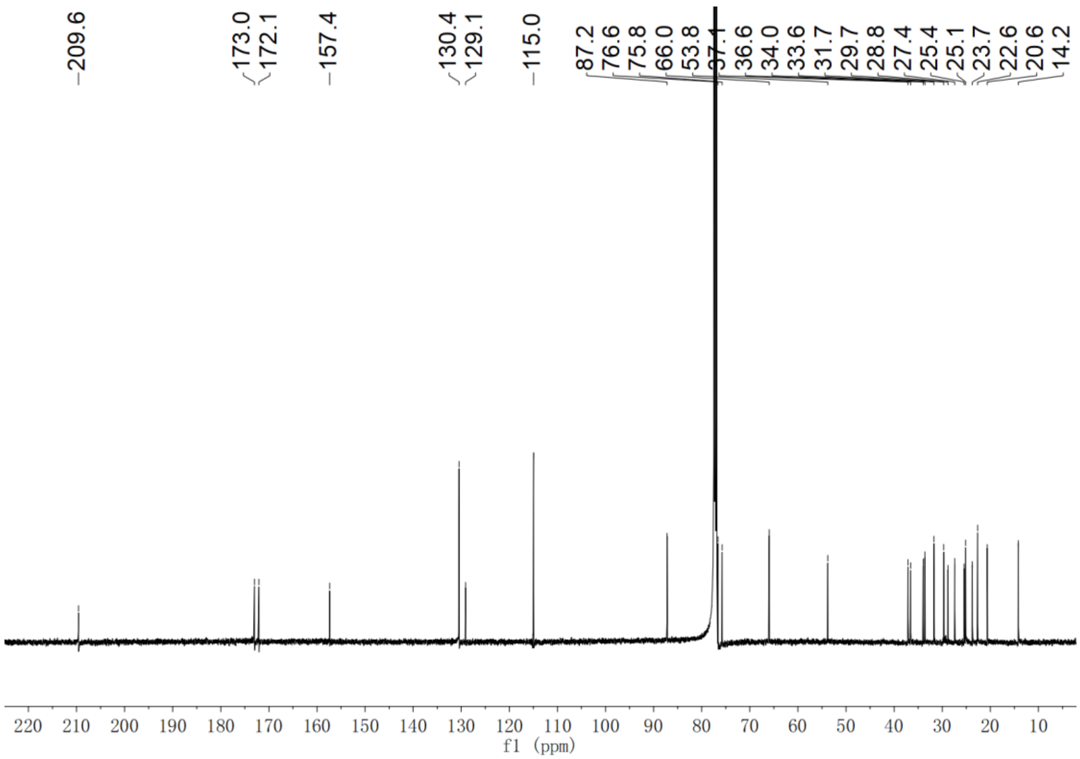


**Supplementary Fig. 56.** $^{13}$C NMR spectrum of compound **13** in $CDCl_3$

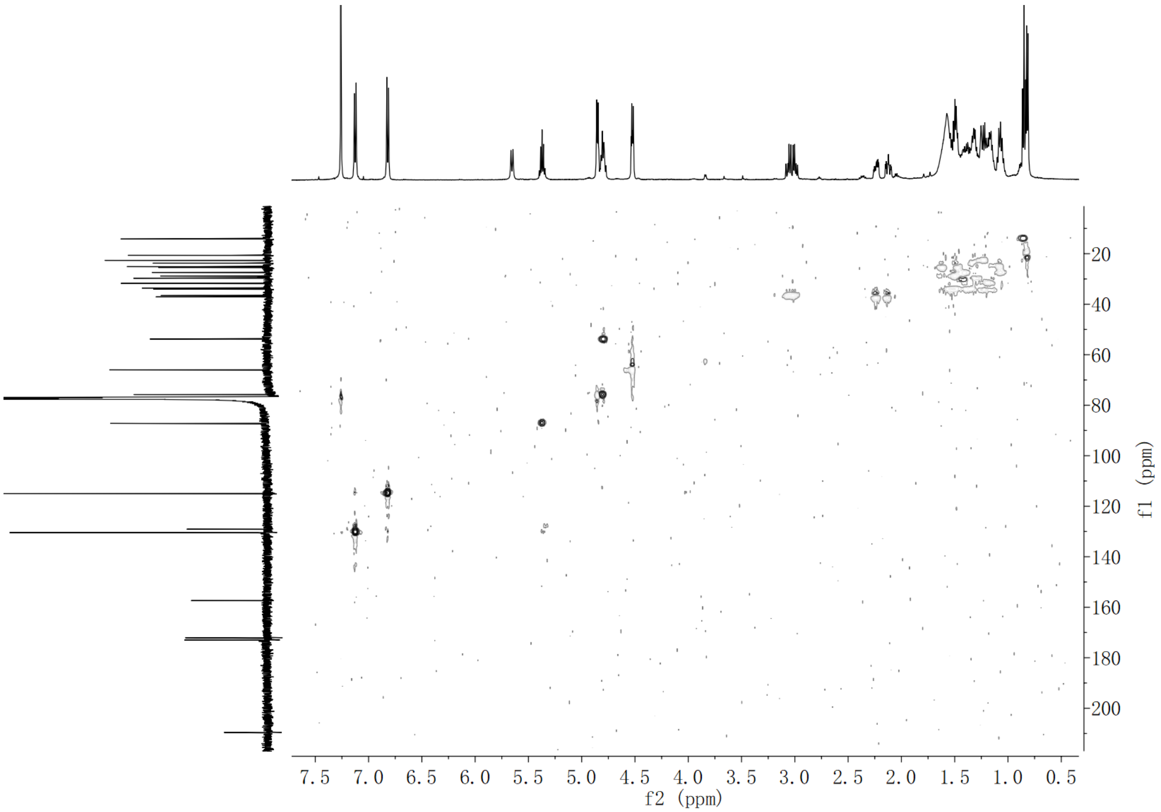


**Supplementary Fig. 57.** HSQC NMR spectrum of compound **13** in $CDCl_3$

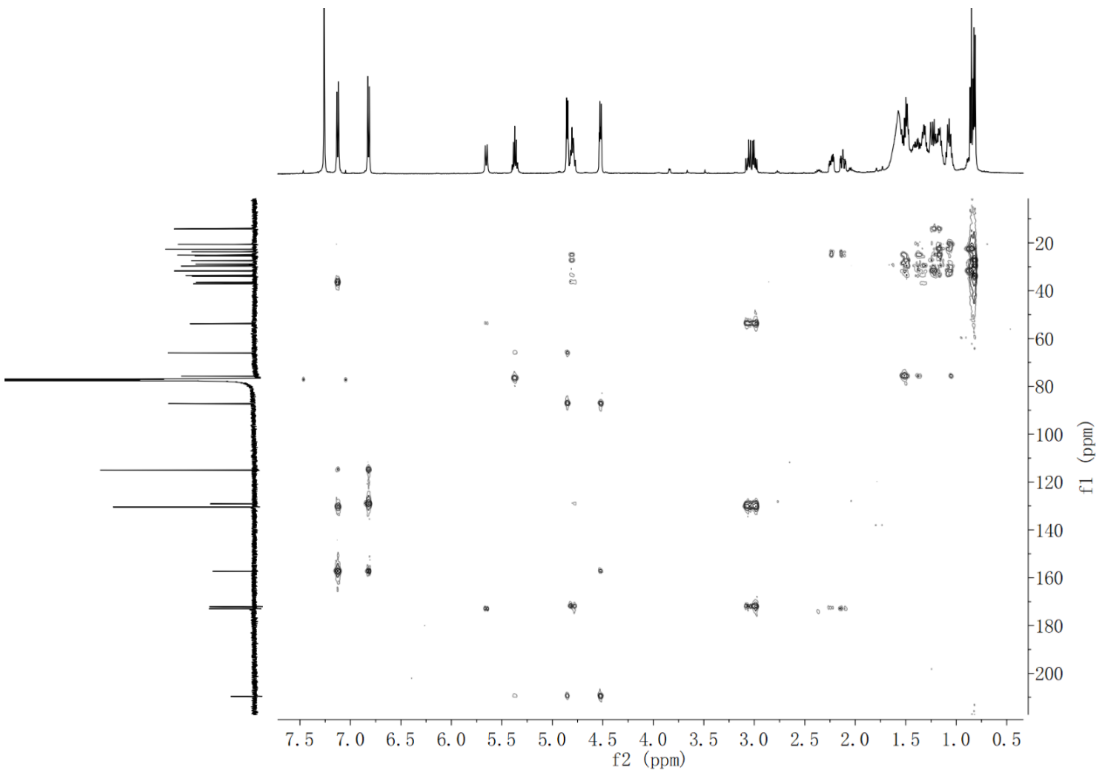


**Supplementary Fig. 58.** HMBC NMR spectrum of compound **13** in $CDCl_3$

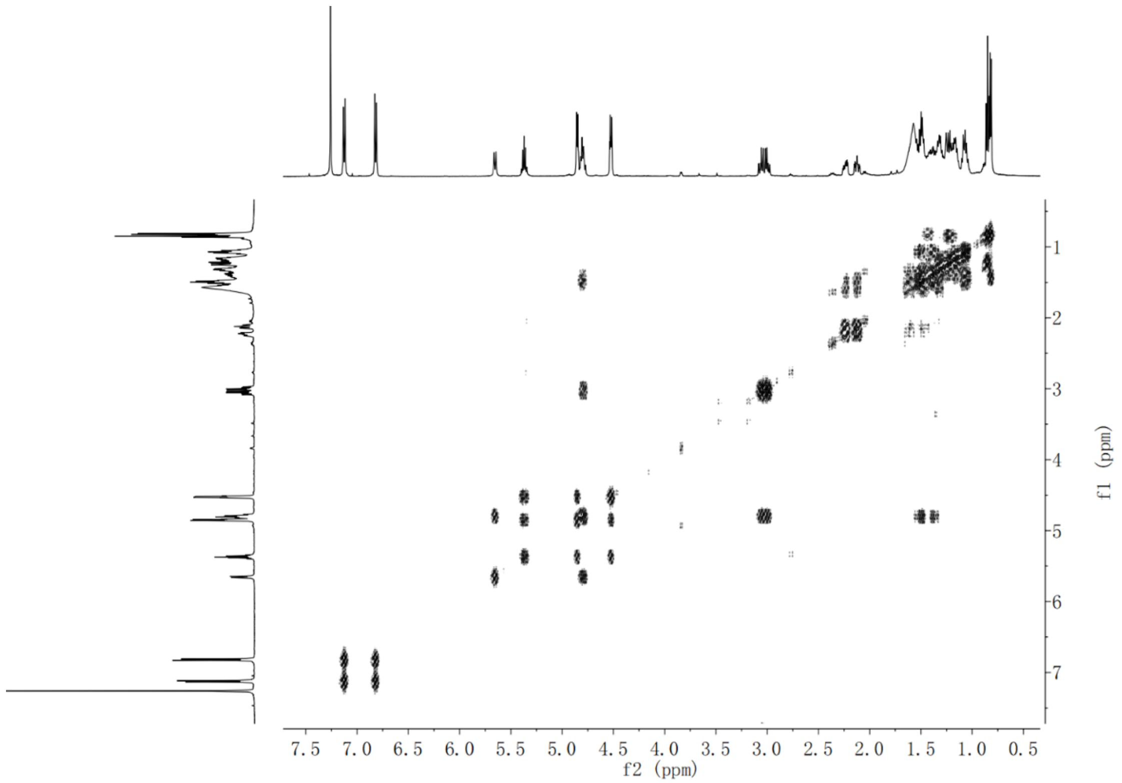


**Supplementary Fig. 59.** $^1H$-$^1H$ COSY NMR spectrum of compound **13** in $CDCl_3$

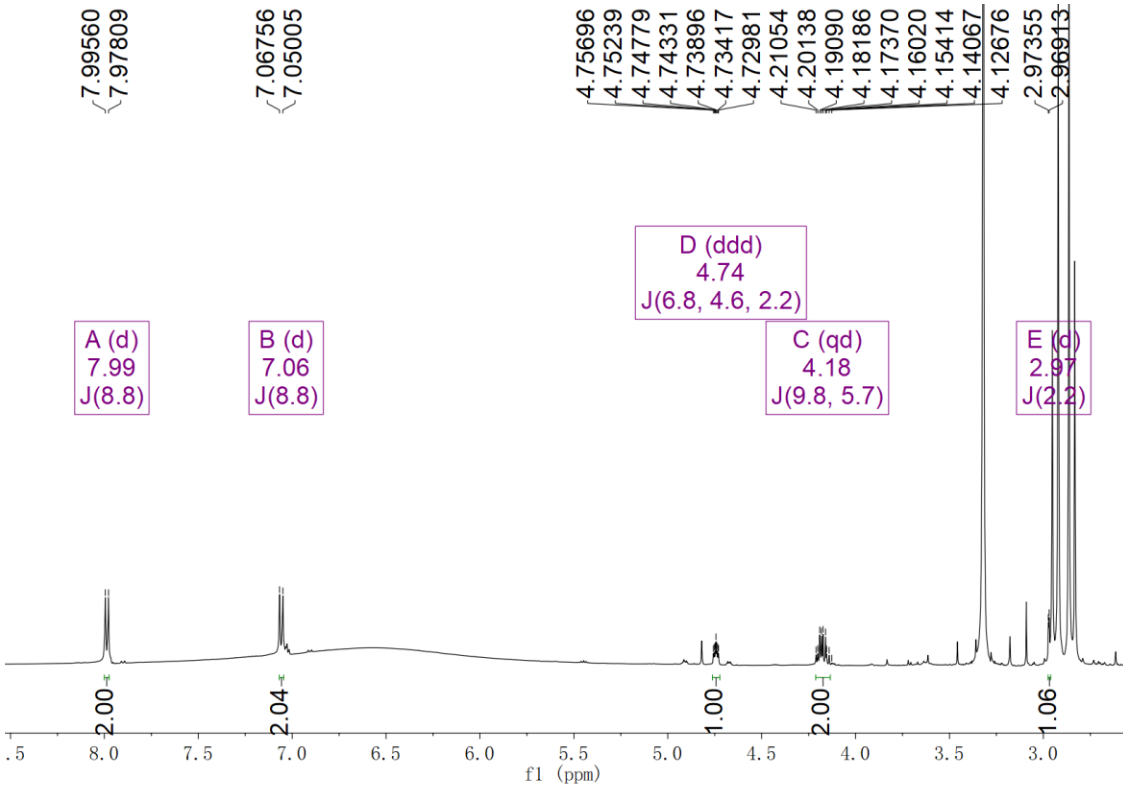


**Supplementary Fig. 60.** $^{1}$H NMR spectrum of demethyl-**5** (**14**) in acetone-$d_6$

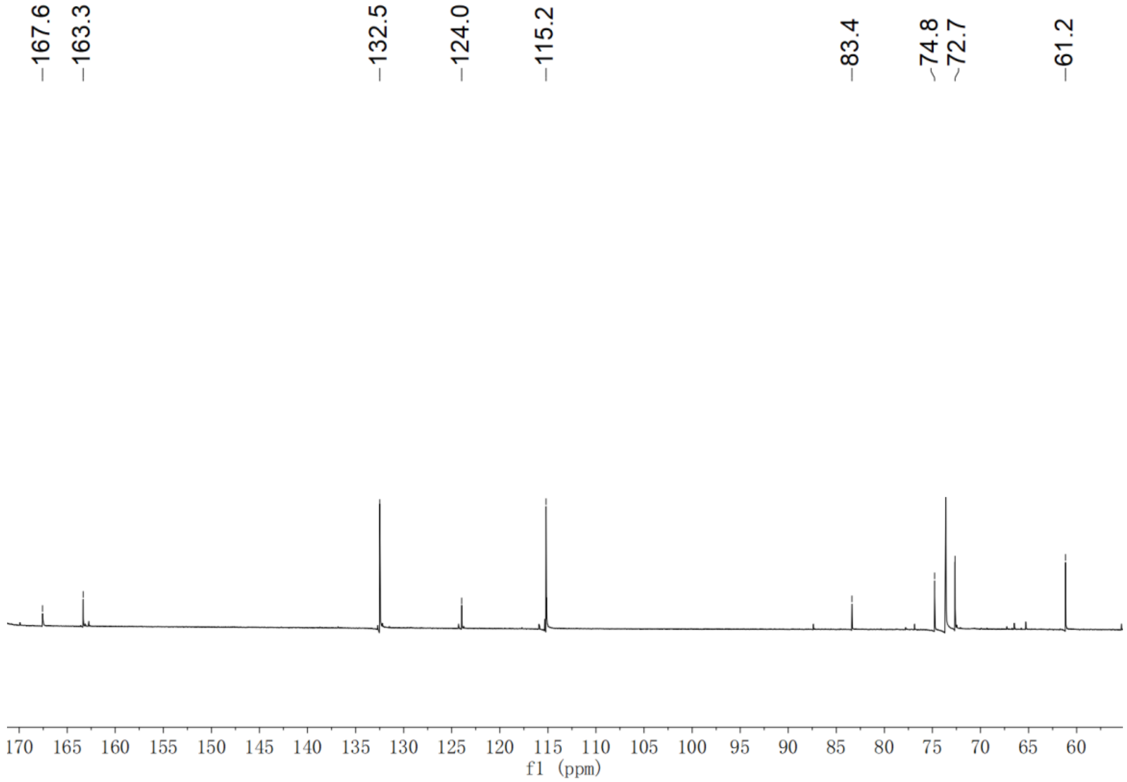


**Supplementary Fig. 61.** $^{13}$C NMR spectrum of demethyl-**5** (**14**) in acetone-$d_6$

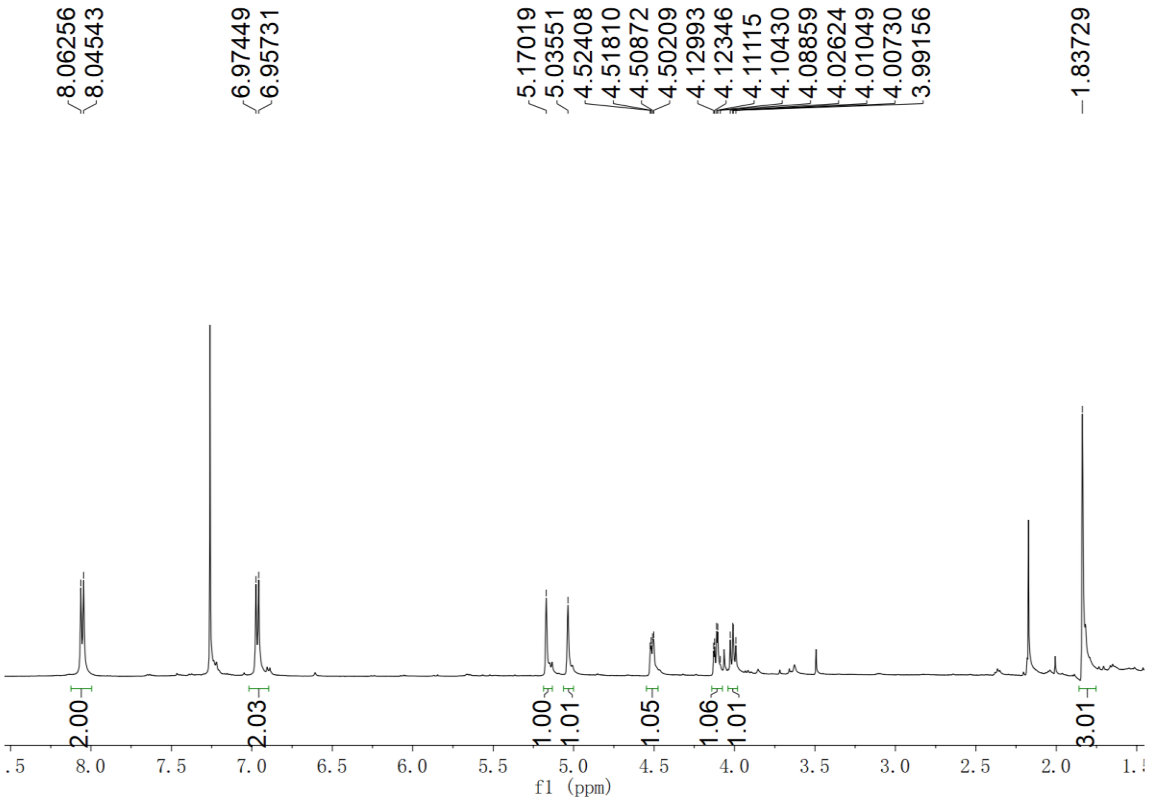


**Supplementary Fig. 62.** $^{1}$H NMR spectrum of **15** in $CDCl_3$

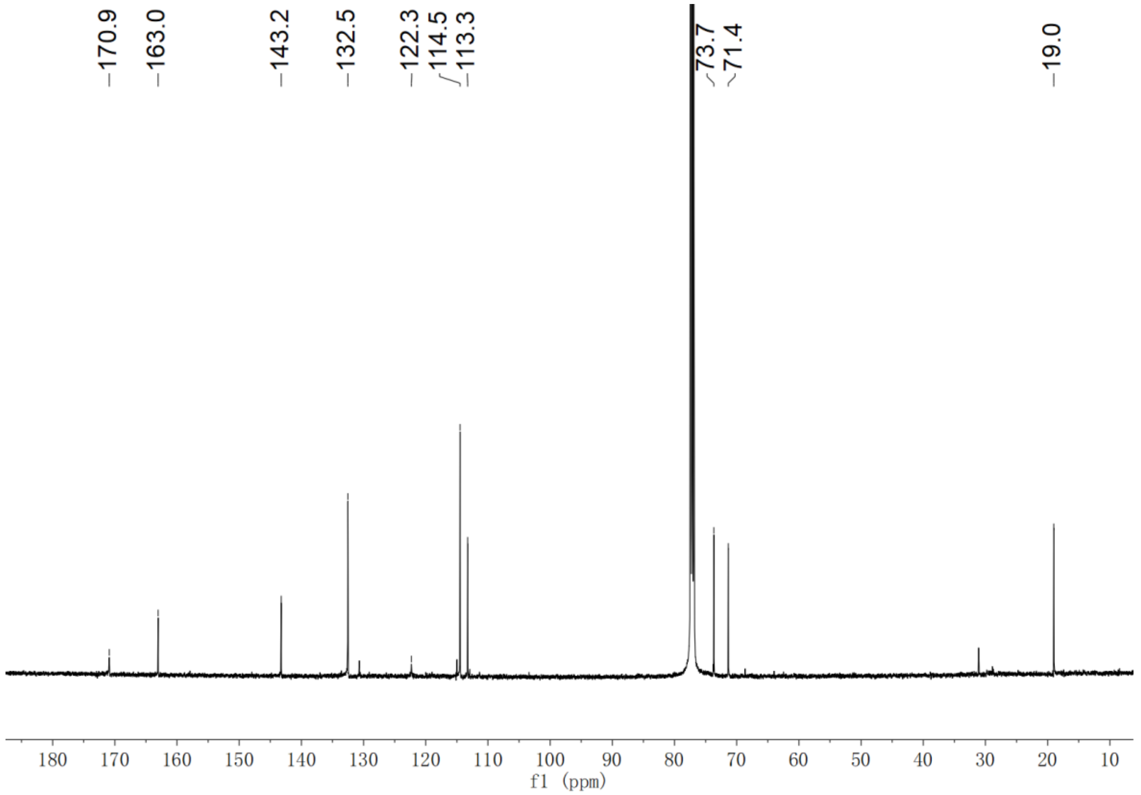


**Supplementary Fig. 63.** $^{13}$C NMR spectrum of **15** in $CDCl_3$

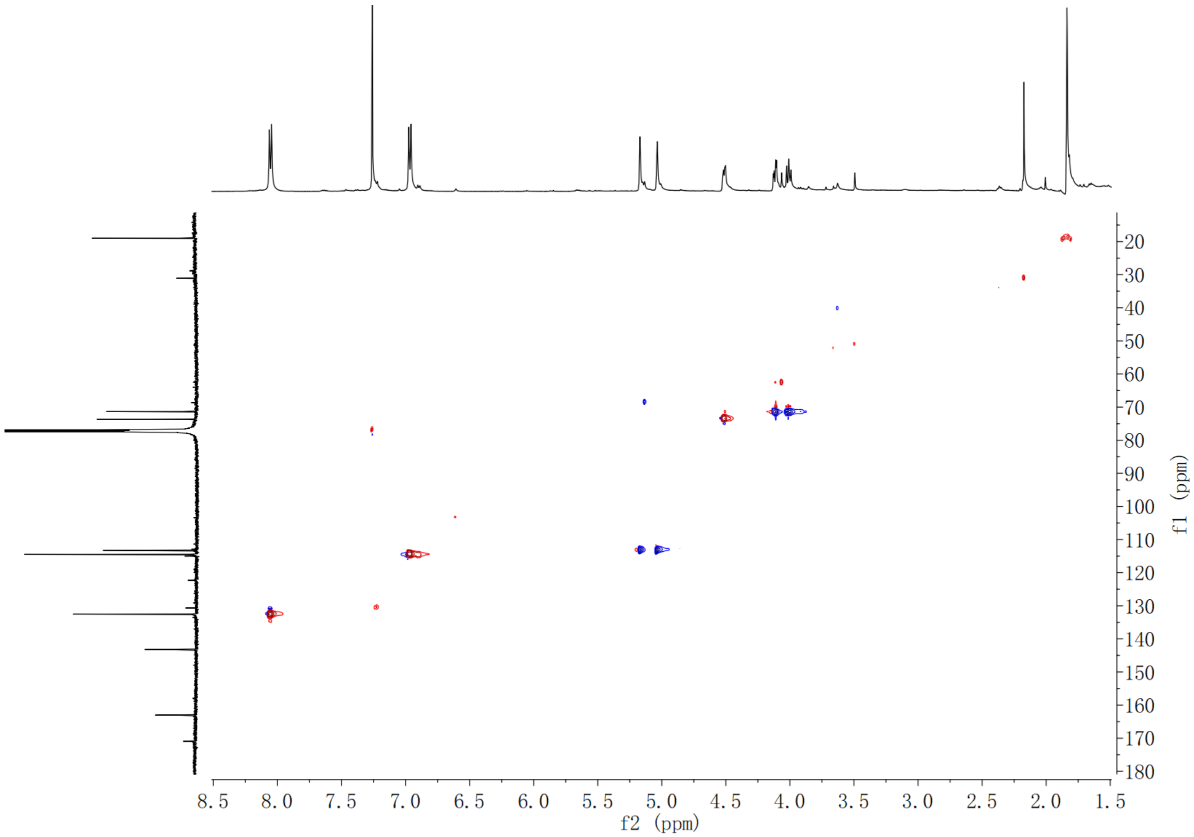


**Supplementary Fig. 64.** HSQC NMR spectrum of **15** in $CDCl_3$

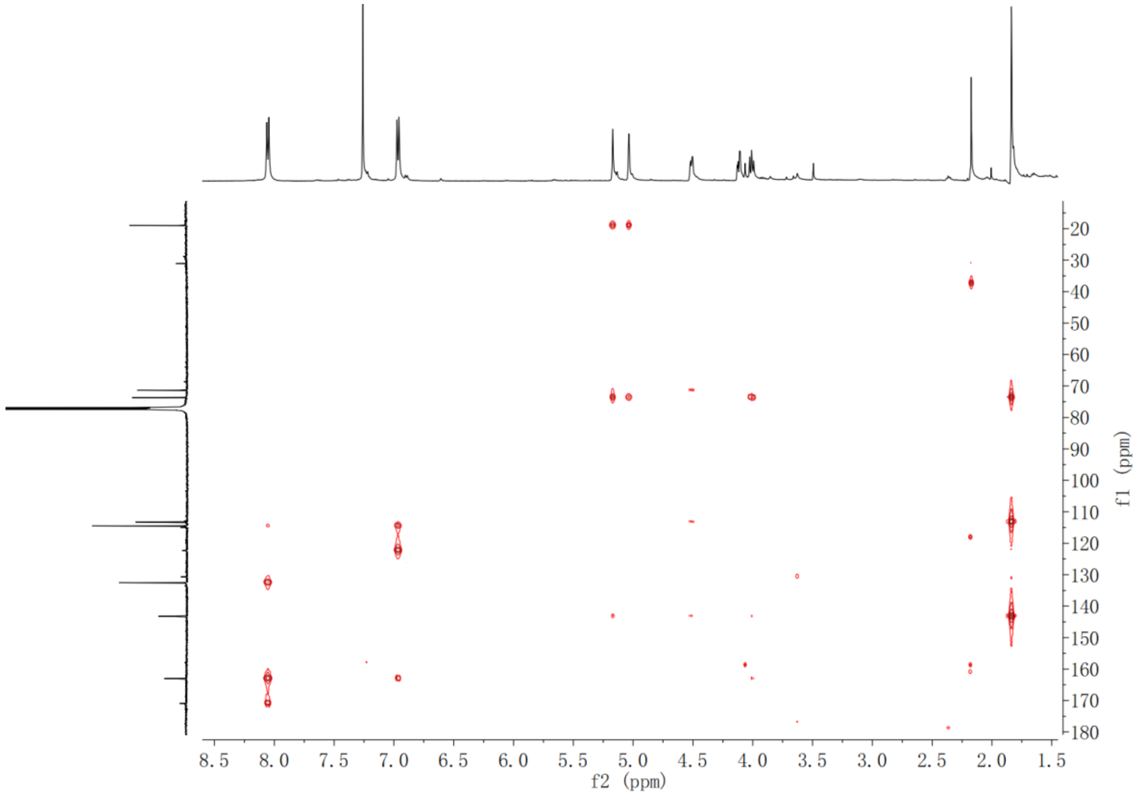


**Supplementary Fig. 65.** HMBC NMR spectrum of **15** in $CDCl_3$

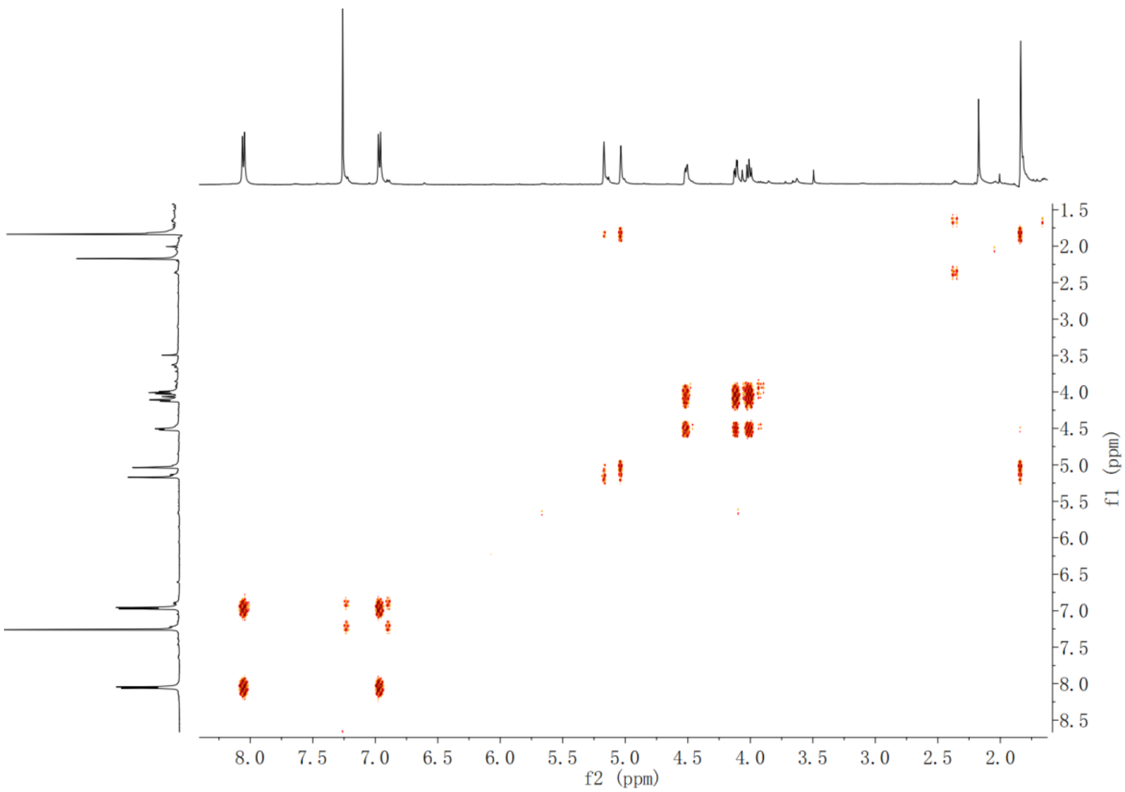


**Supplementary Fig. 66.** $^{1}H$-$^{1}H$ COSY NMR spectrum of **15** in $CDCl_3$

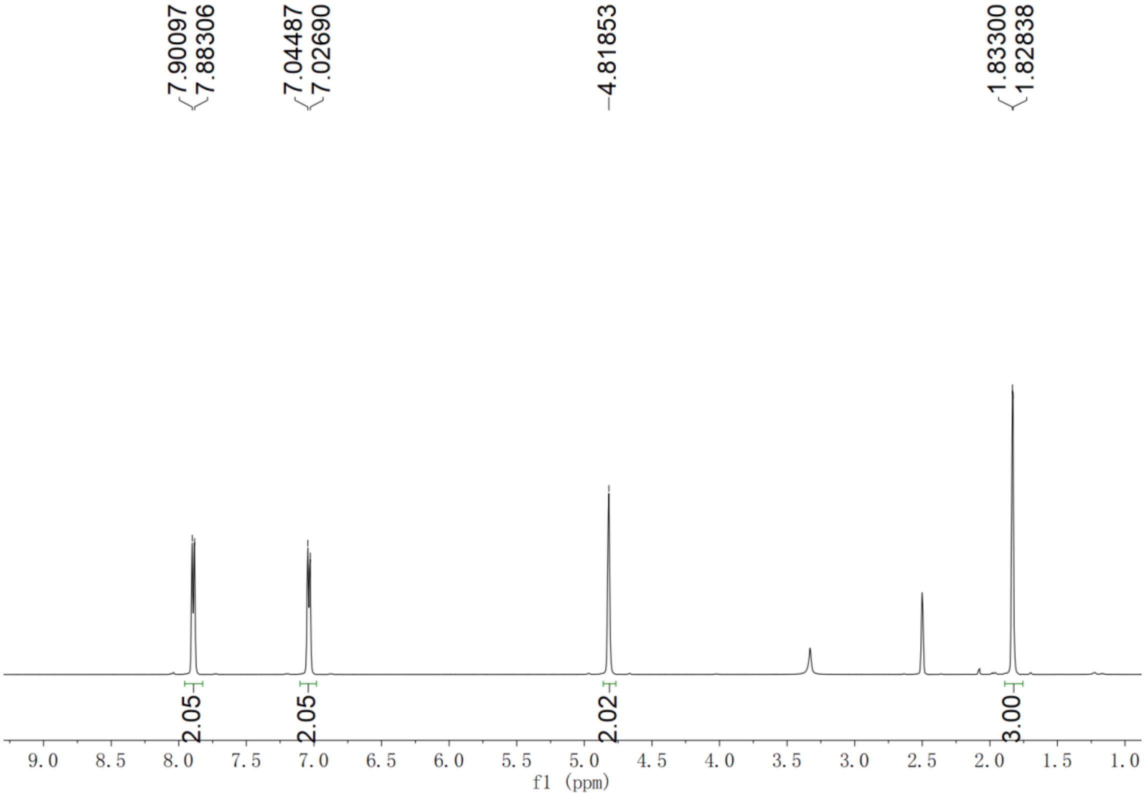


**Supplementary Fig. 67.** $^{1}H$ NMR spectrum of O-but-2-ynyl benzoic acid (**16**) in DMSO-$d_6$

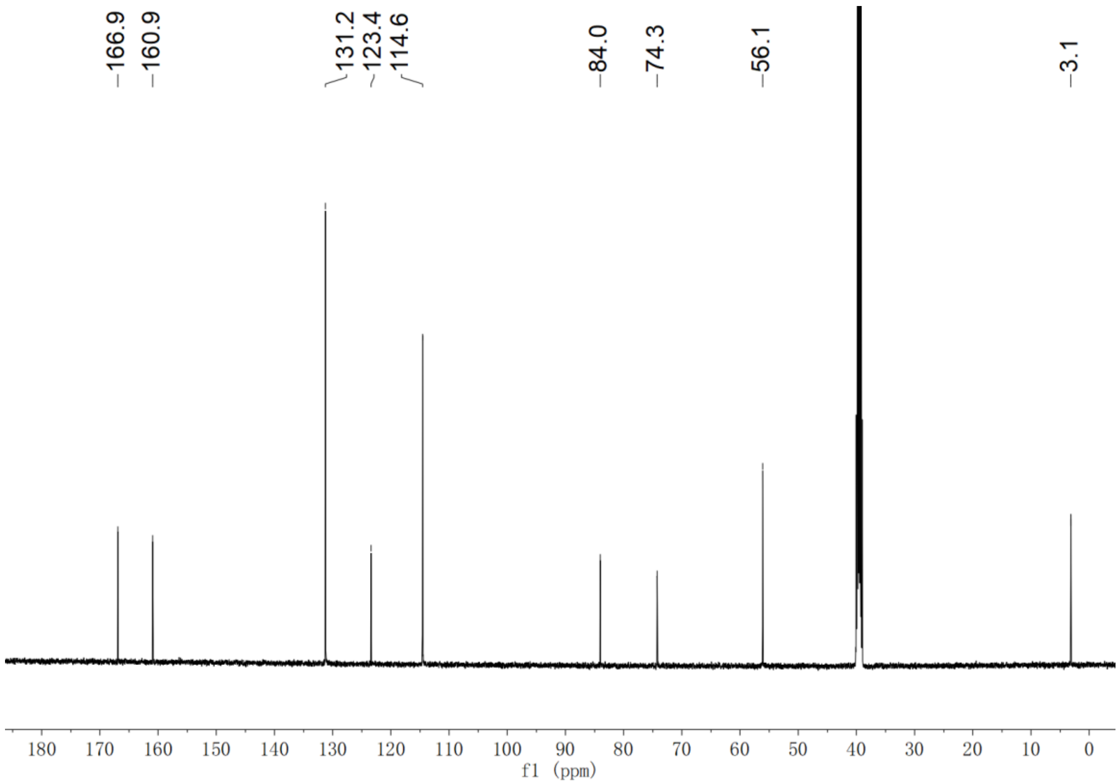


**Supplementary Fig. 68.** $^{13}C$ NMR spectrum of O-but-2-ynyl benzoic acid (**16**) in DMSO-$d_6$

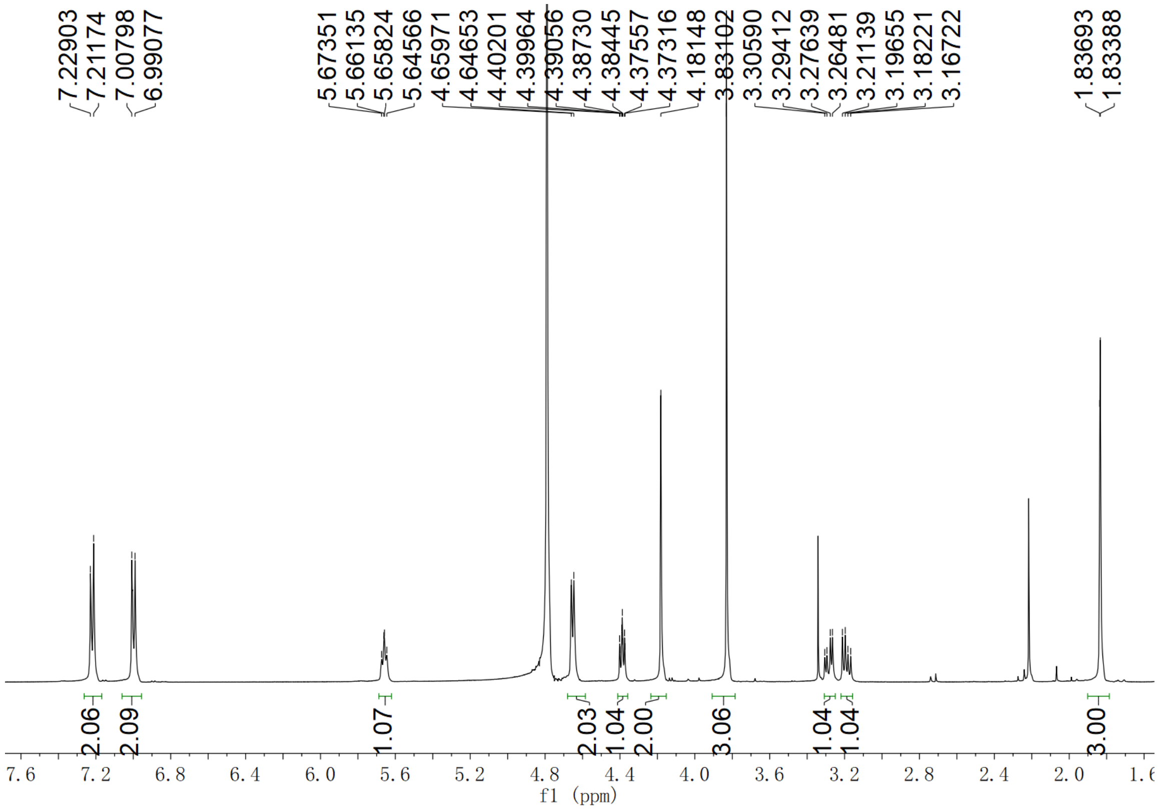


**Supplementary Fig. 69.** $^1H$ NMR spectrum of **17**-Me in $D_2O$

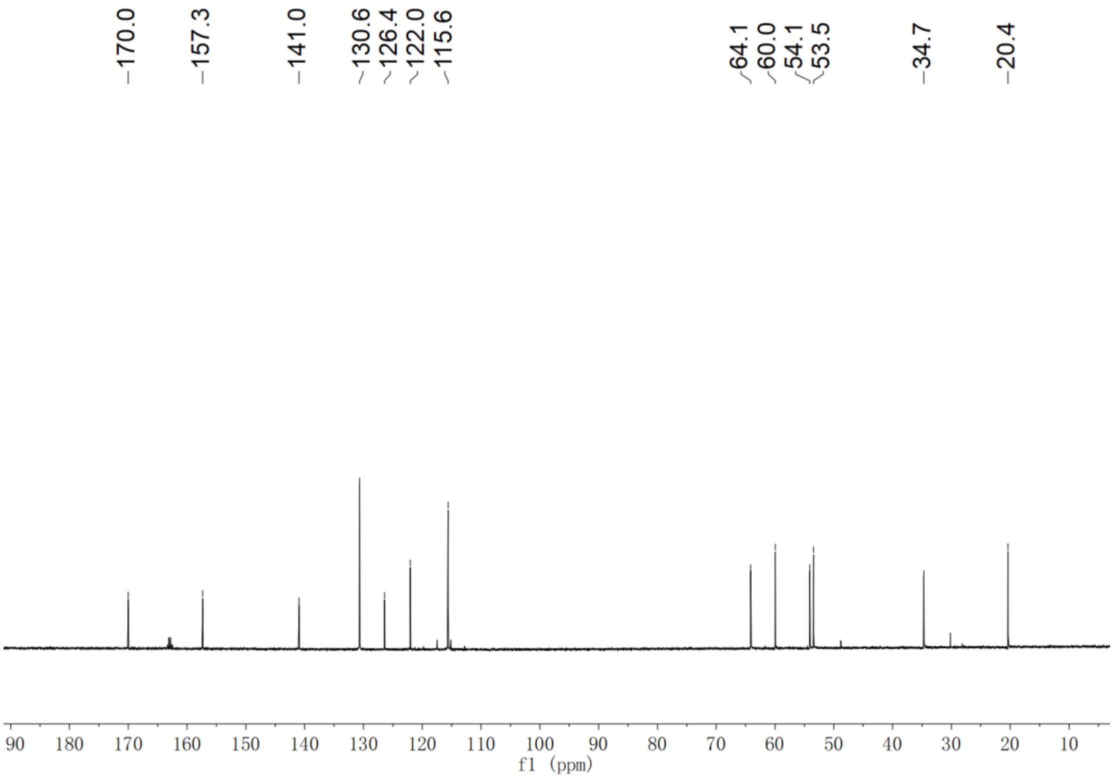


**Supplementary Fig. 70.** $^{13}C$ NMR spectrum of **17**-Me in $D_2O$

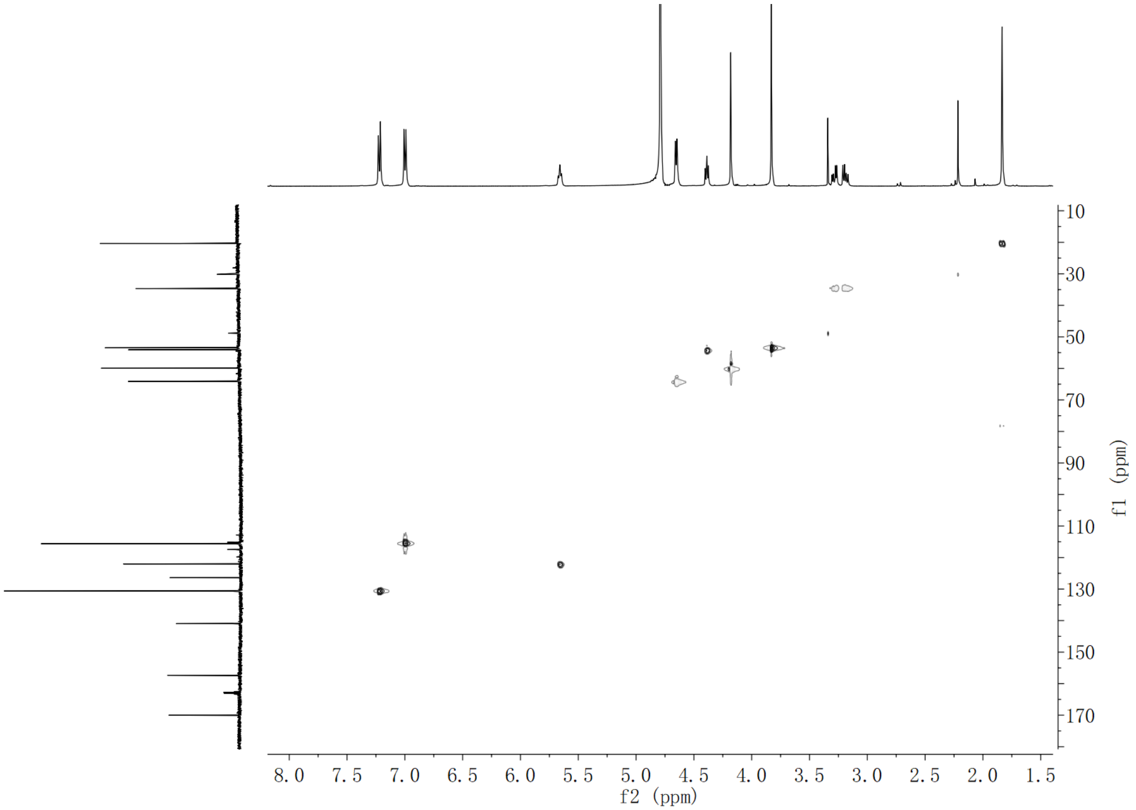


**Supplementary Fig. 71.** HSQC NMR spectrum of **17**-Me in $D_2O$

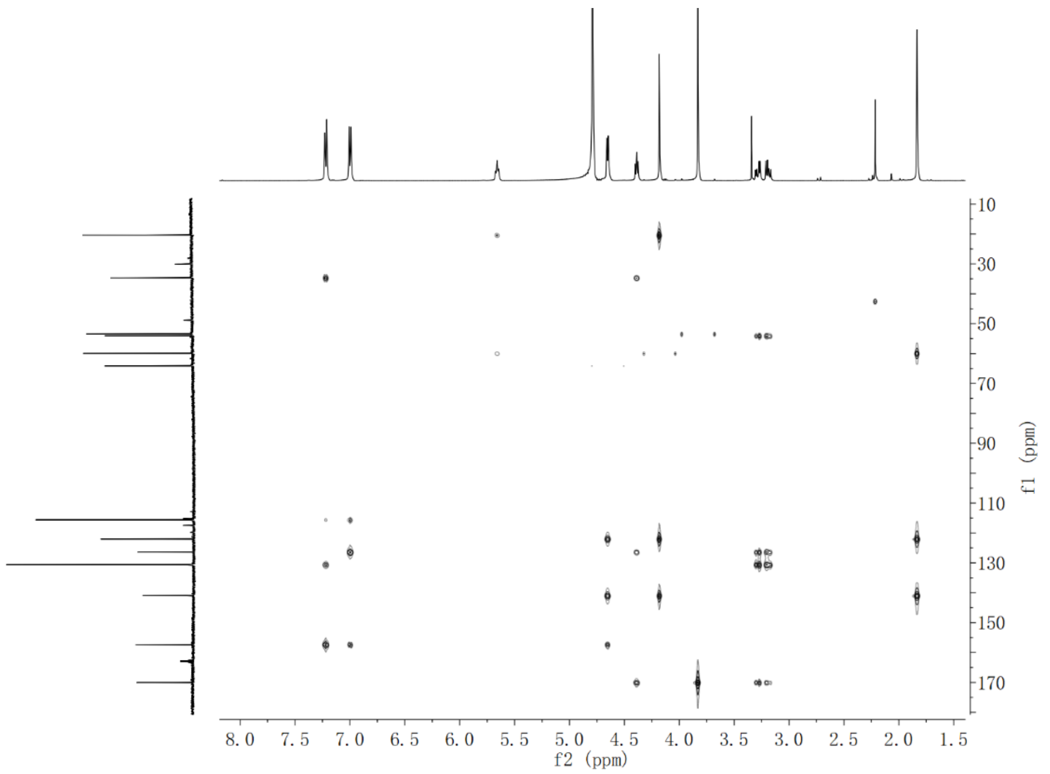


**Supplementary Fig. 72.** HMBC NMR spectrum of **17**-Me in $D_2O$

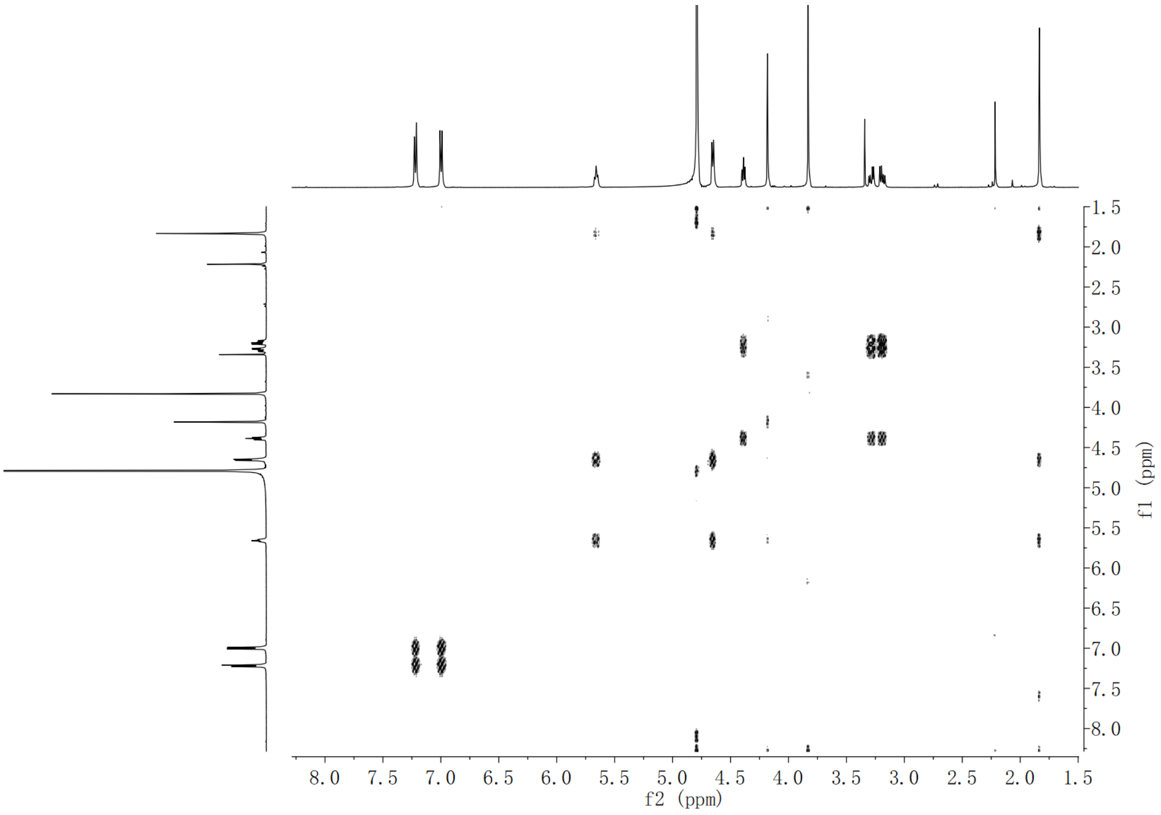


**Supplementary Fig. 73.** $^{1}H$-$^{1}H$ COSY NMR spectrum of **17**-Me in $D_2O$

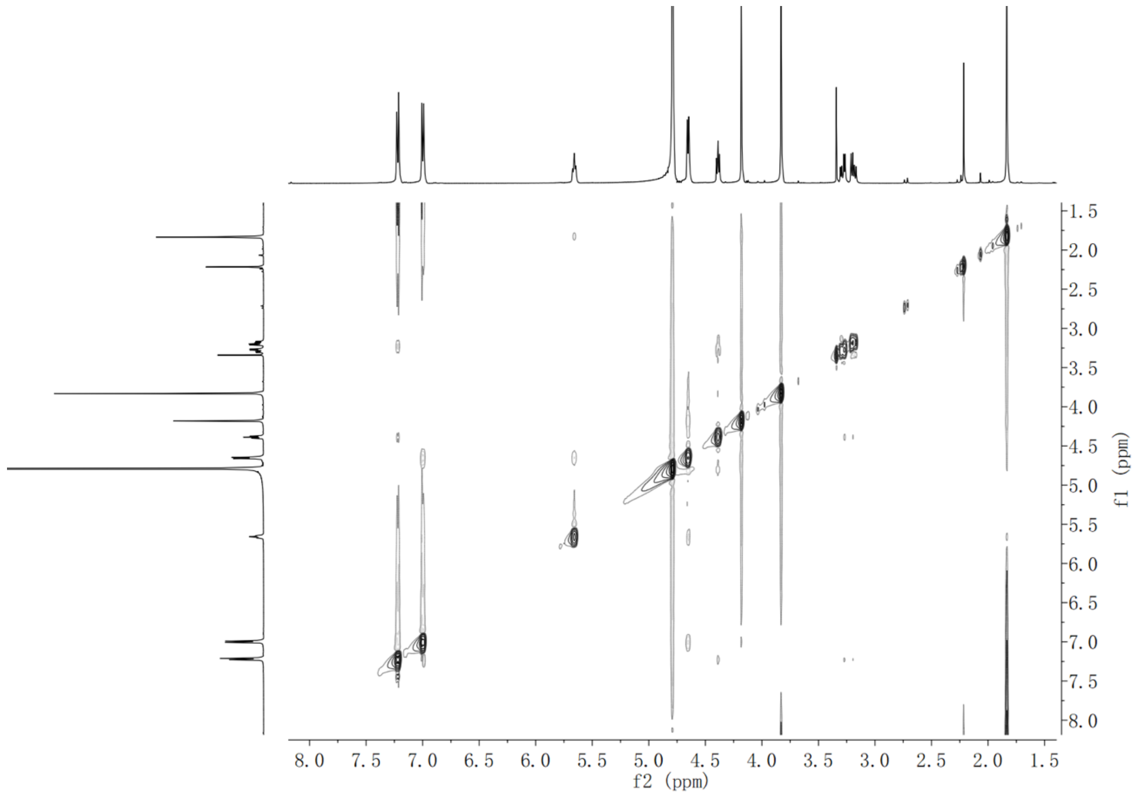


**Supplementary Fig. 74.** NOESY NMR spectrum of **17**-Me in $D_2O$

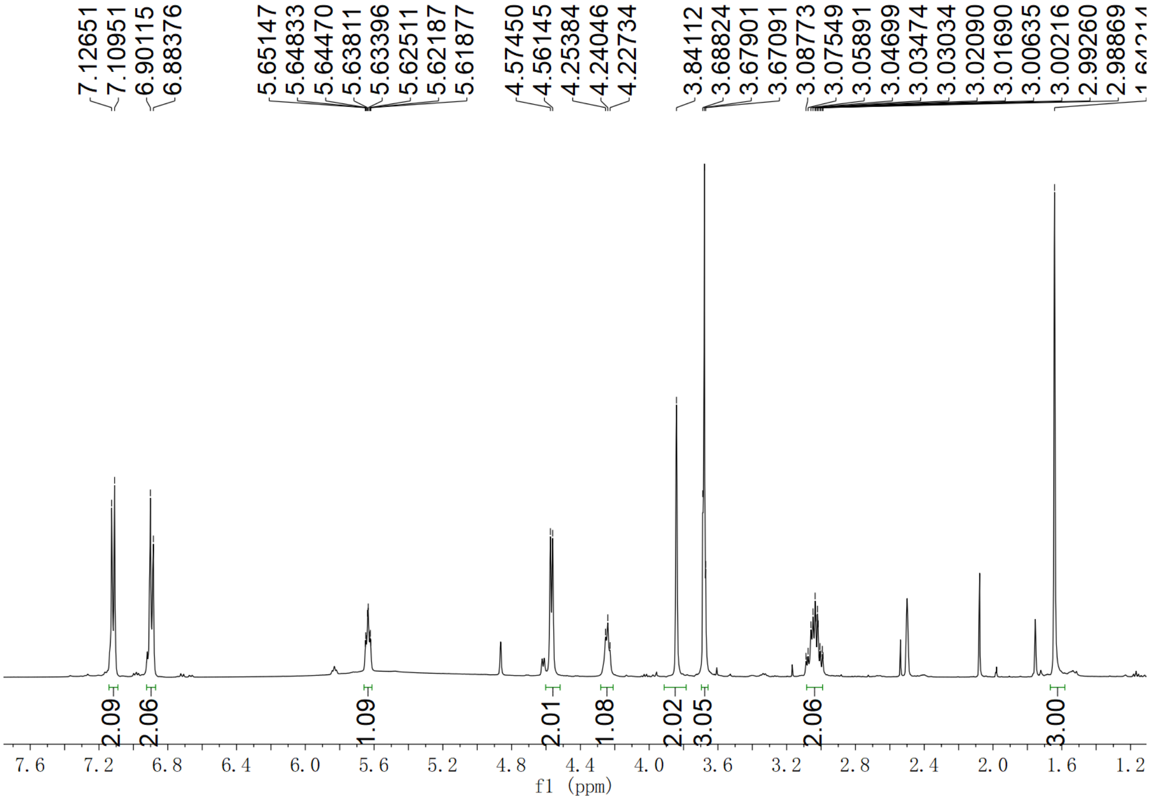


**Supplementary Fig. 75.** [1]H NMR spectrum of **18**-Me in DMSO-*d6*

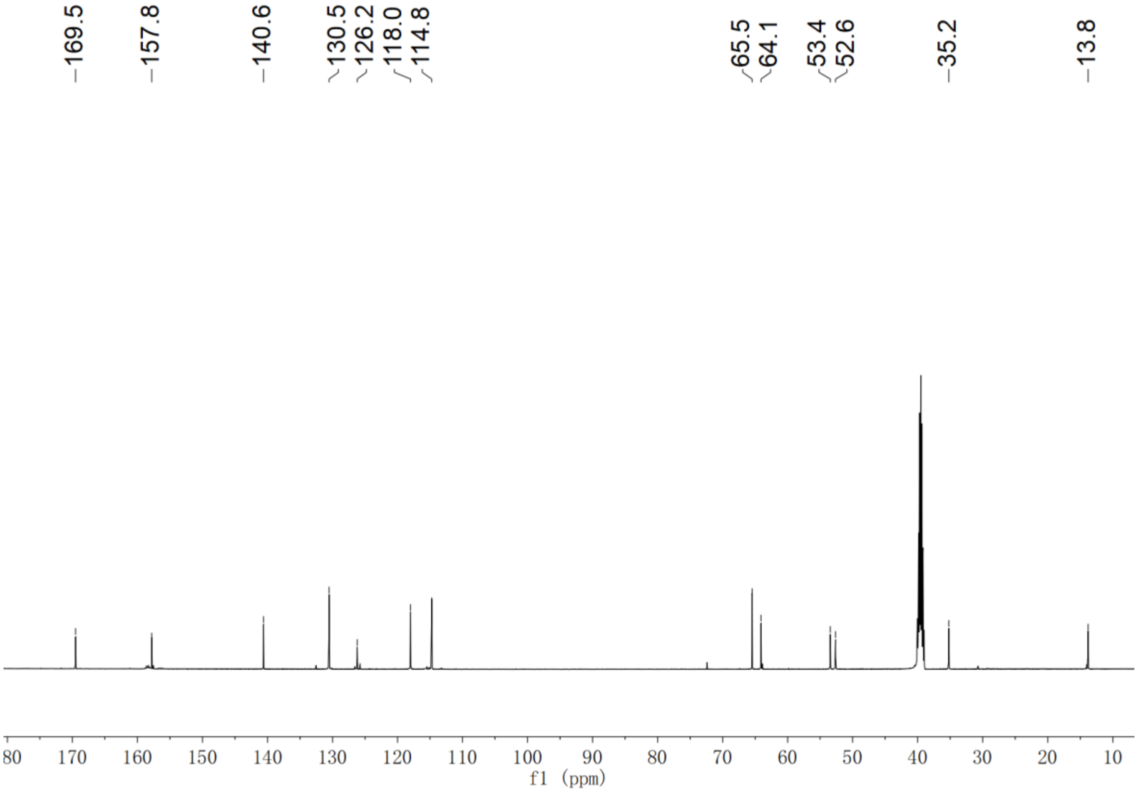


**Supplementary Fig. 76.** $^{13}$C NMR spectrum of **18**-Me in DMSO-*d6*

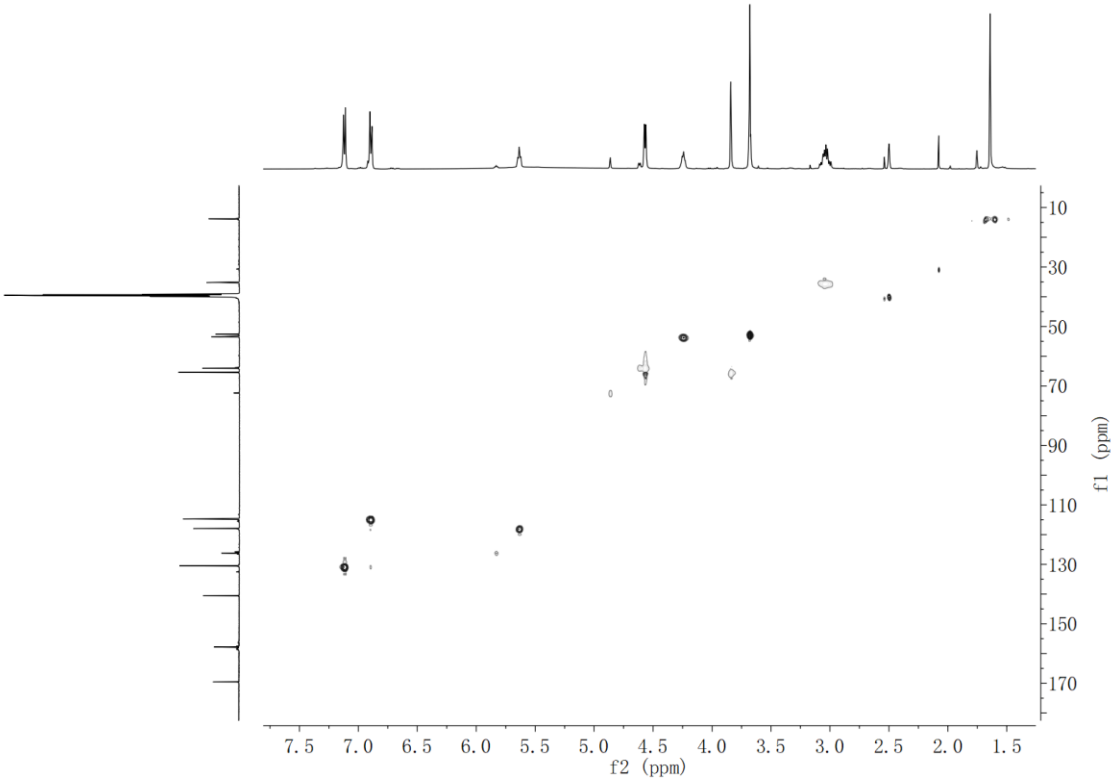


**Supplementary Fig. 77.** HSQC NMR spectrum of **18**-Me in DMSO-*d6*

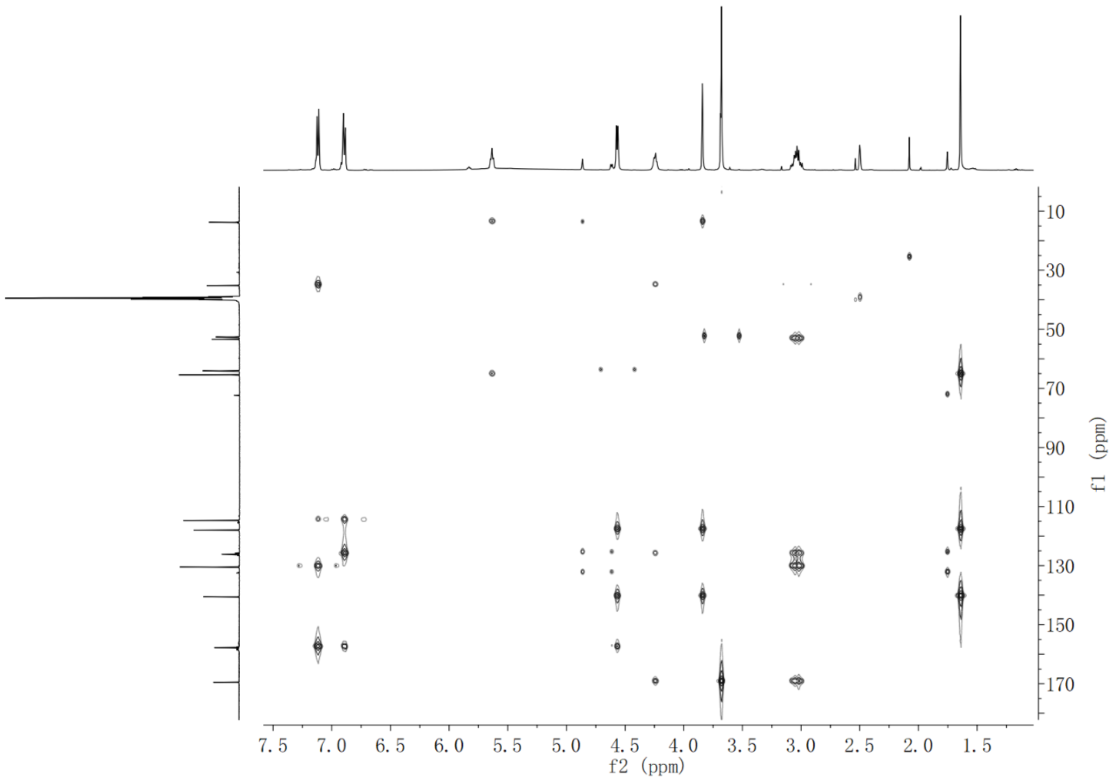


**Supplementary Fig. 78.** HMBC NMR spectrum of **18**-Me in DMSO-*d6*

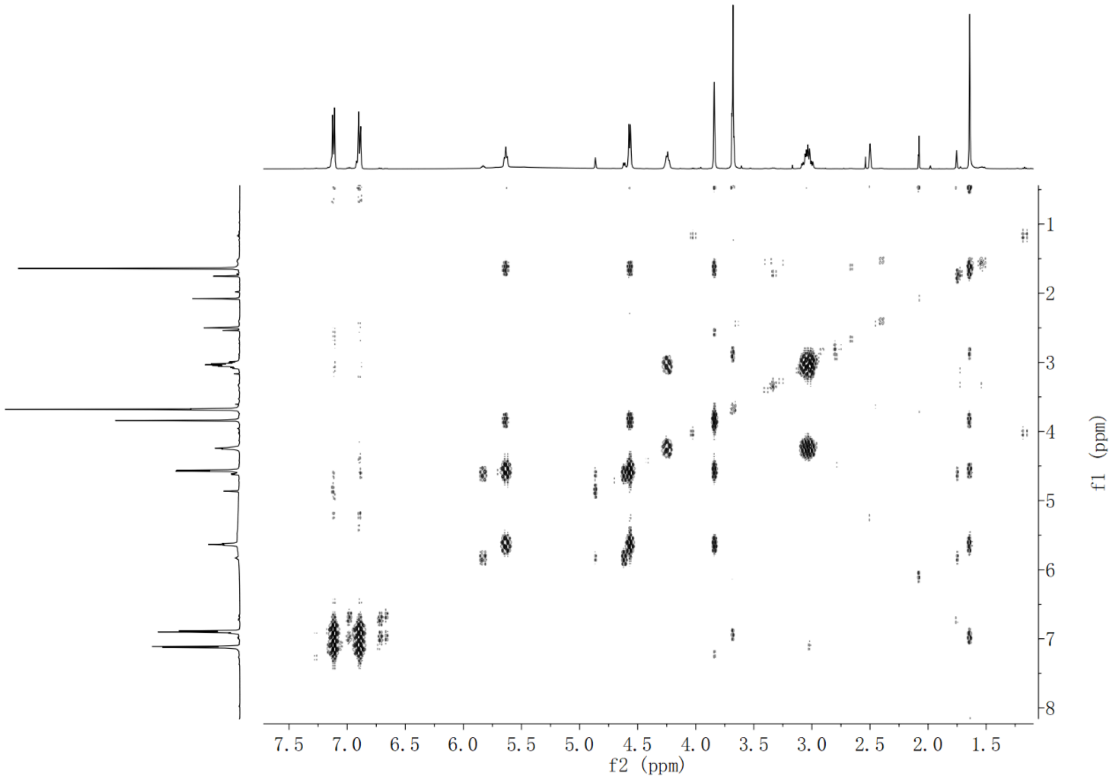


**Supplementary Fig. 79.** ${}^{1}H$-${}^{1}H$ COSY NMR spectrum of **18**-Me in DMSO-*d6*

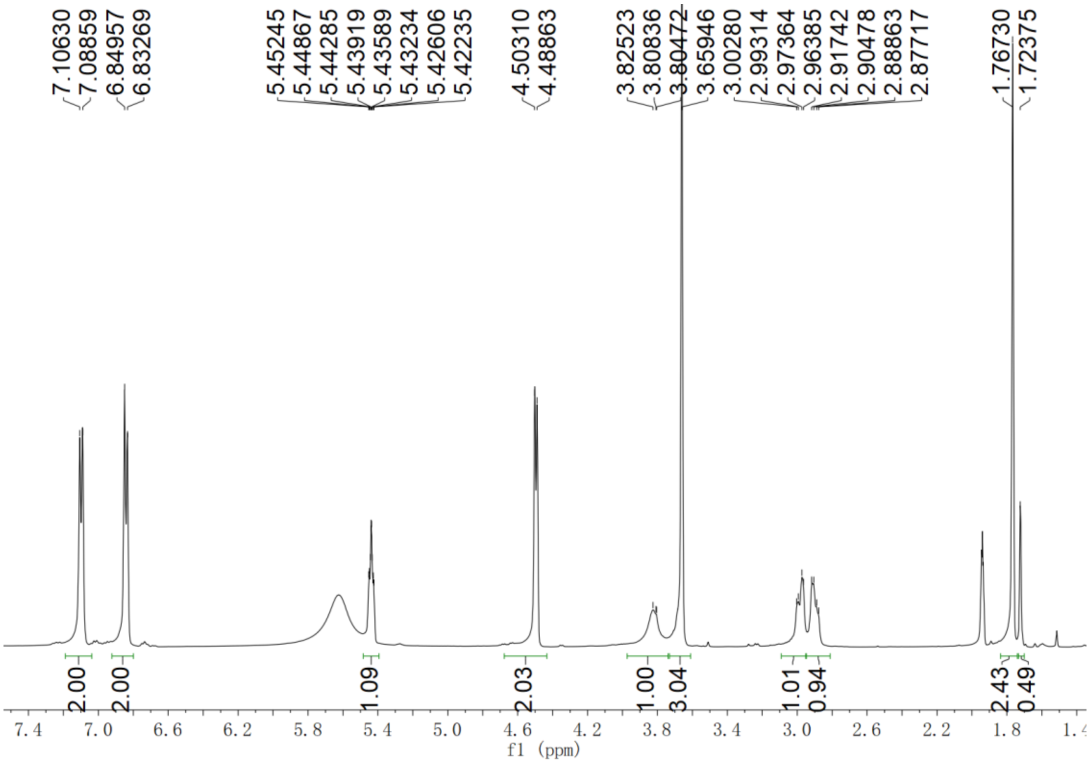


**Supplementary Fig. 80.** $^{1}$H NMR spectrum of 4’,4’,4’-[$^{2}H_3$]-**7-Me** in $CD_3CN$

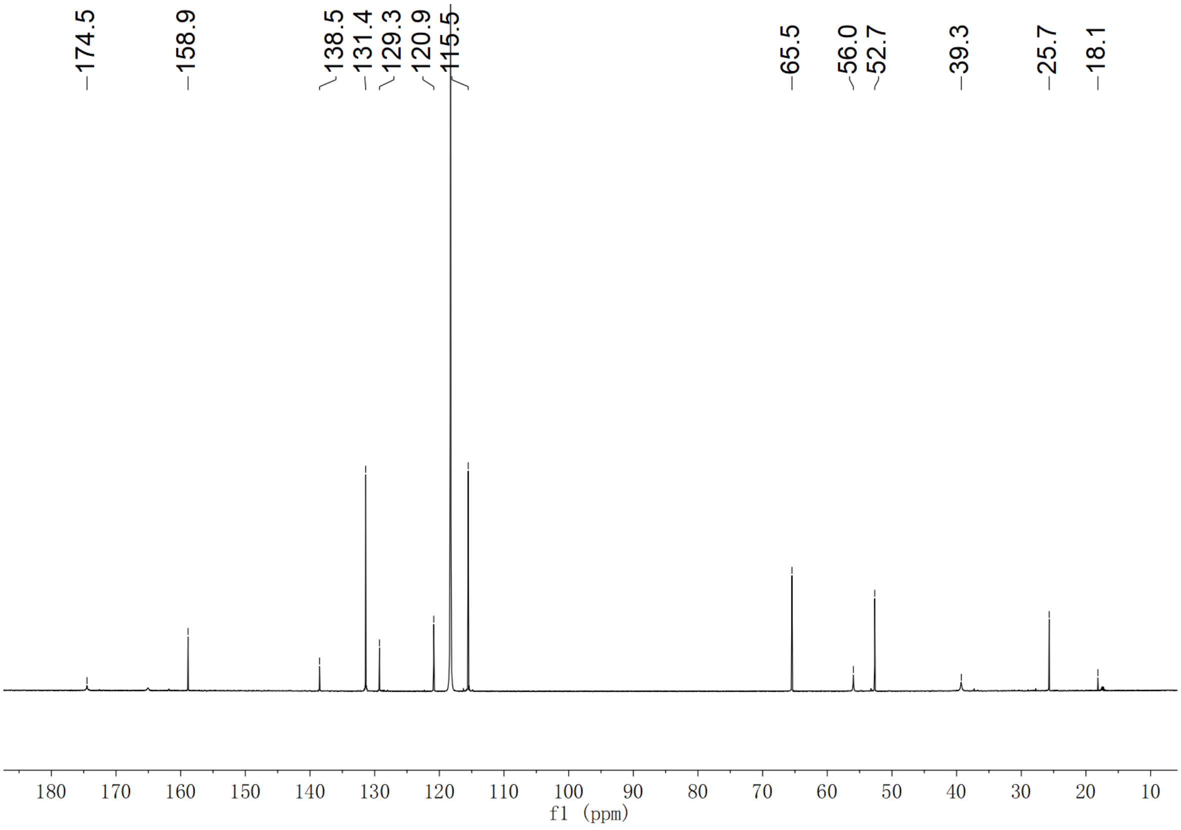


**Supplementary Fig. 81.** $^{13}$C NMR spectrum of 4’,4’,4’-[$^{2}H_3$]-**7-Me** in $CD_3CN$

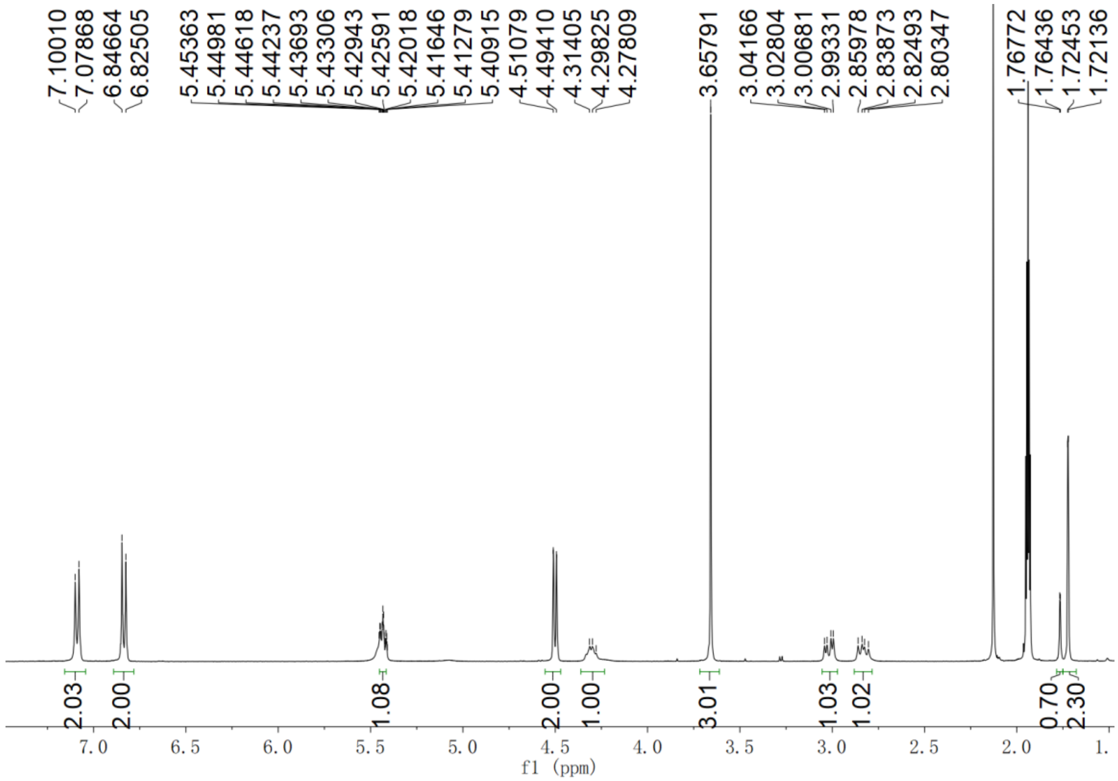


**Supplementary Fig. 82.** $^{1}H$ NMR spectrum of 5',5',5'-[$^{2}H_3$]-**7-Me** in $CD_3CN$

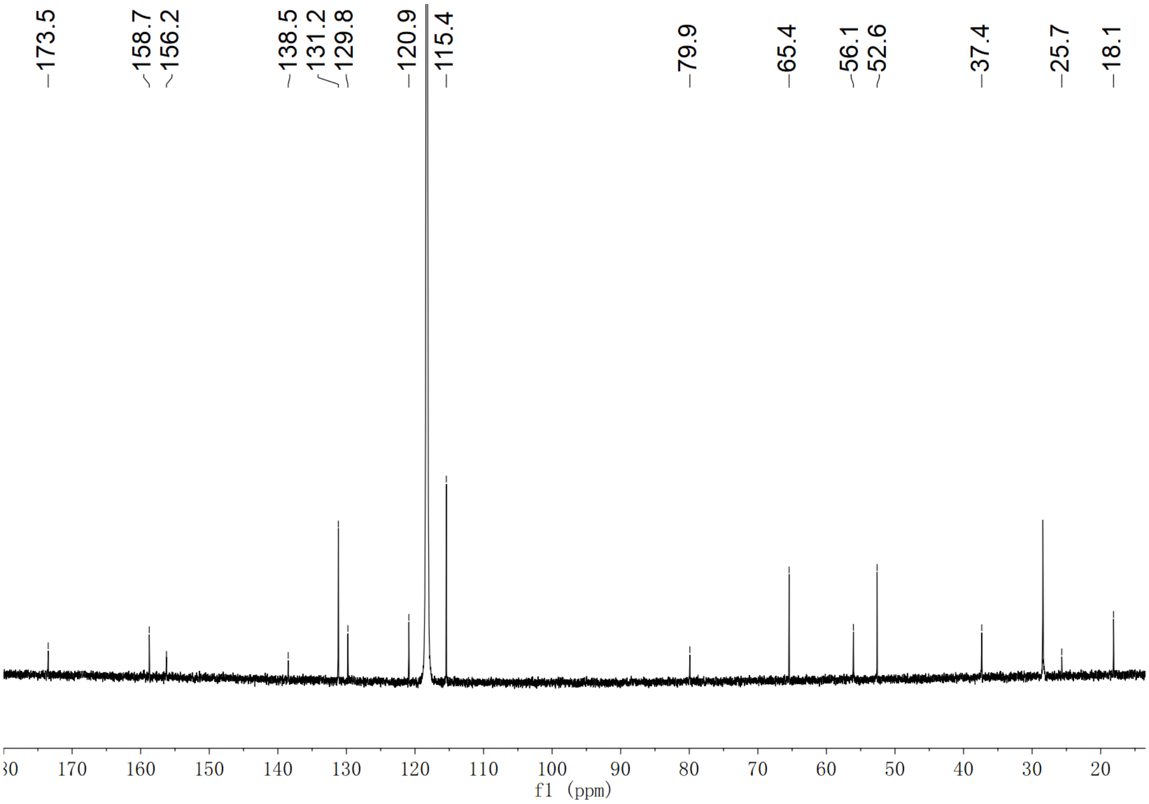


**Supplementary Fig. 83.** $^{13}C$ NMR spectrum of 5',5',5'-[$^{2}H_3$]-**7-Me** in $CD_3CN$

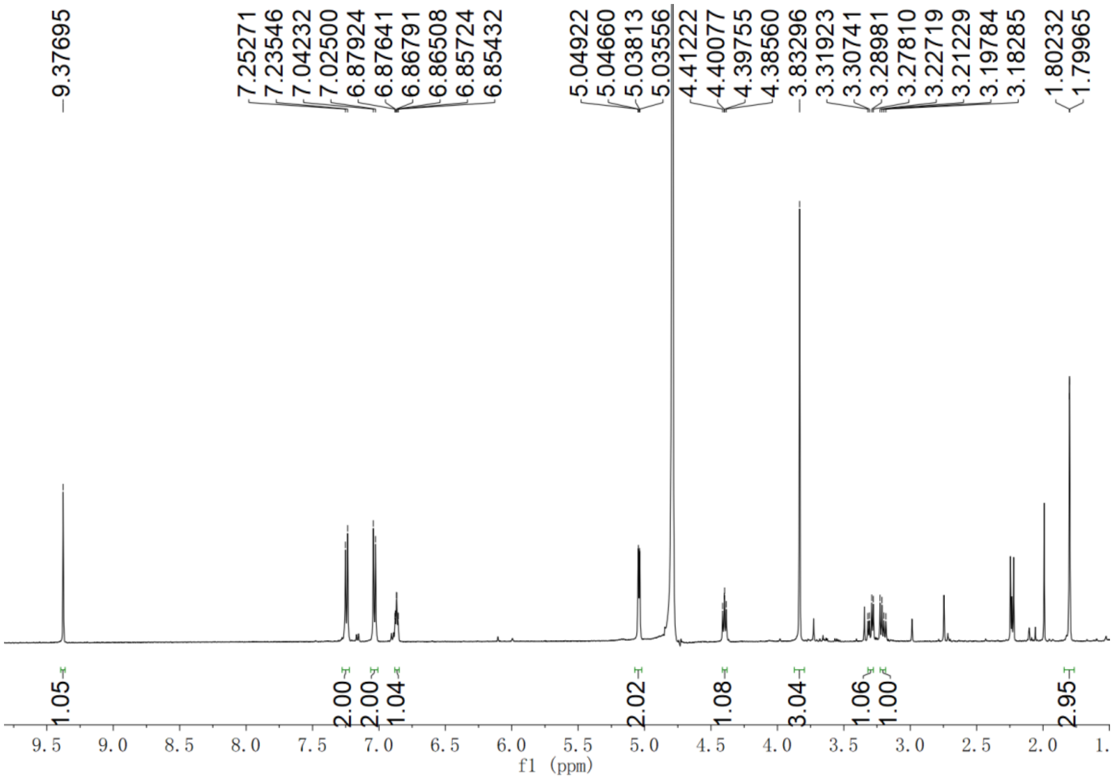


**Supplementary Fig. 84.** $^{1}$H NMR spectrum of compound **19-Me** in $D_2O$

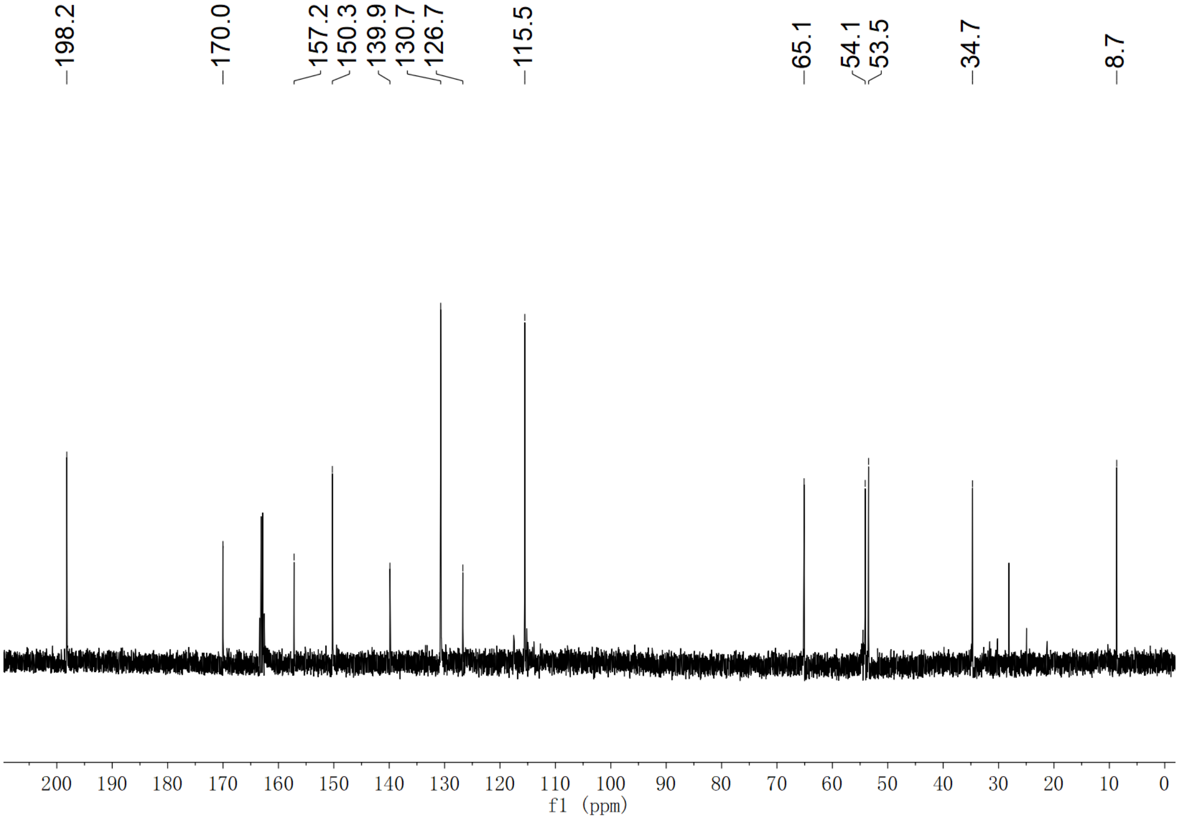


**Supplementary Fig. 85.** $^{13}$C NMR spectrum of compound **19-Me** in $D_2O$

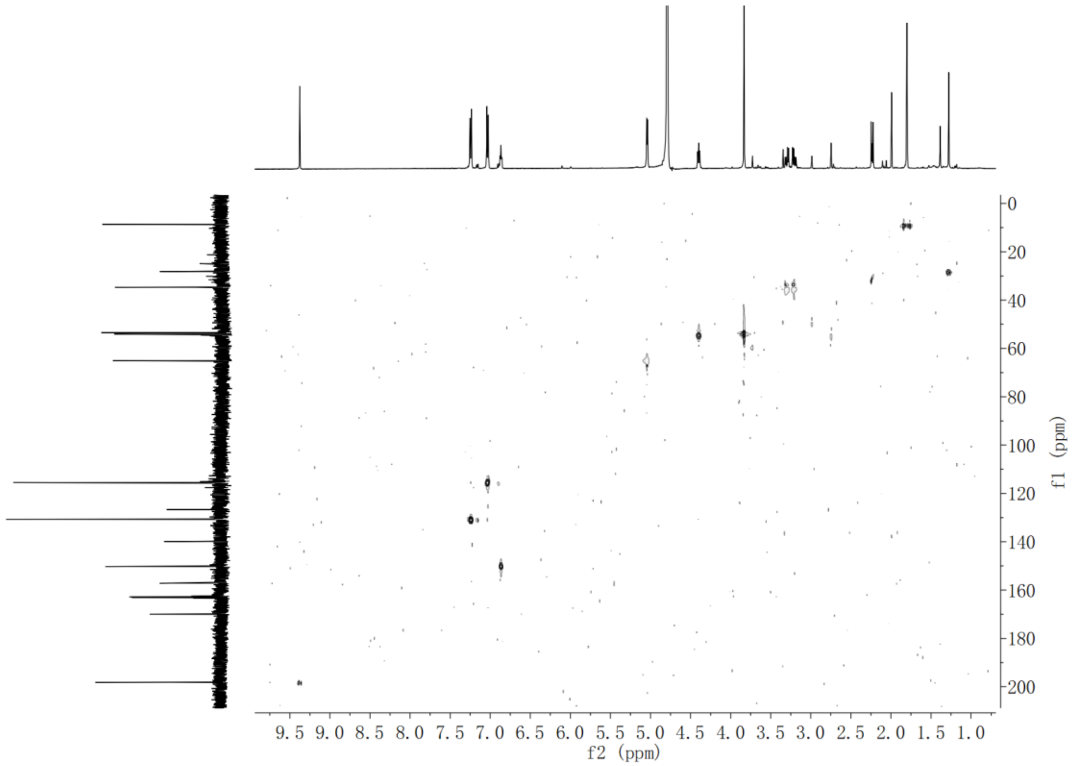


**Supplementary Fig. 86.** HSQC NMR spectrum of compound **19-Me** in $D_2O$

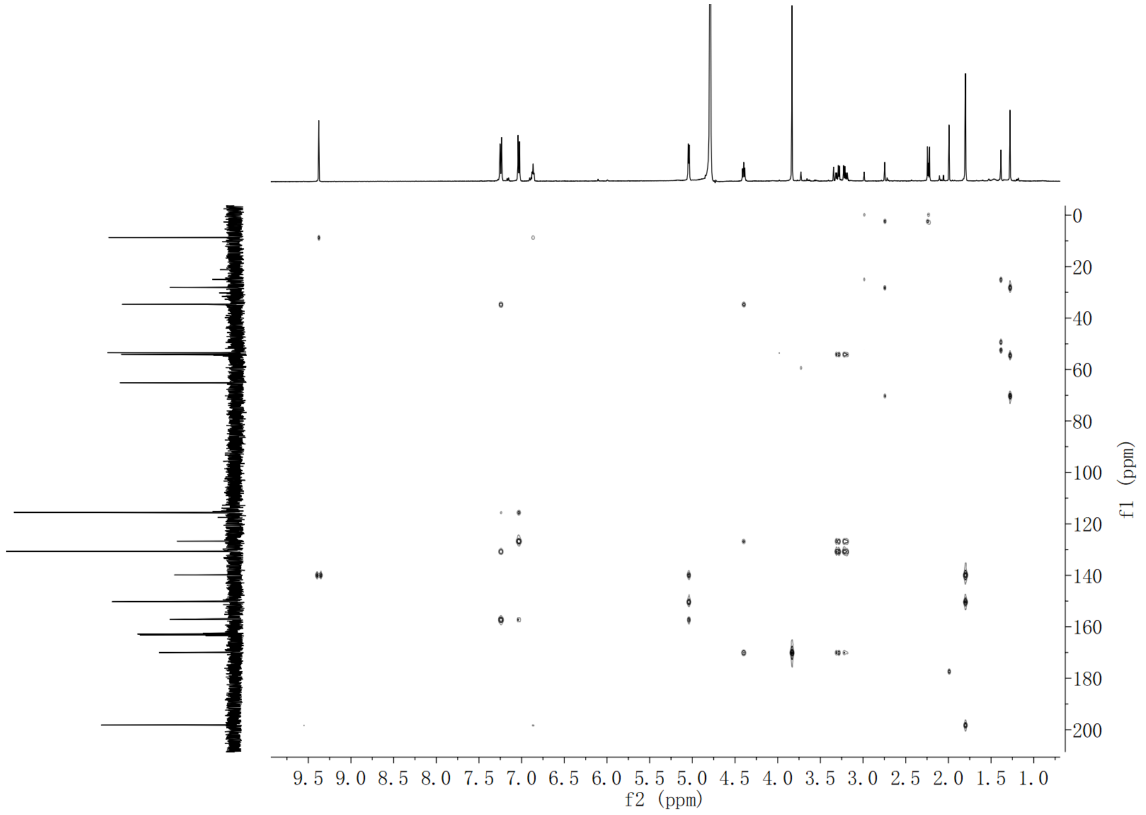


**Supplementary Fig. 87.** HMBC NMR spectrum of compound **19-Me** in $D_2O$

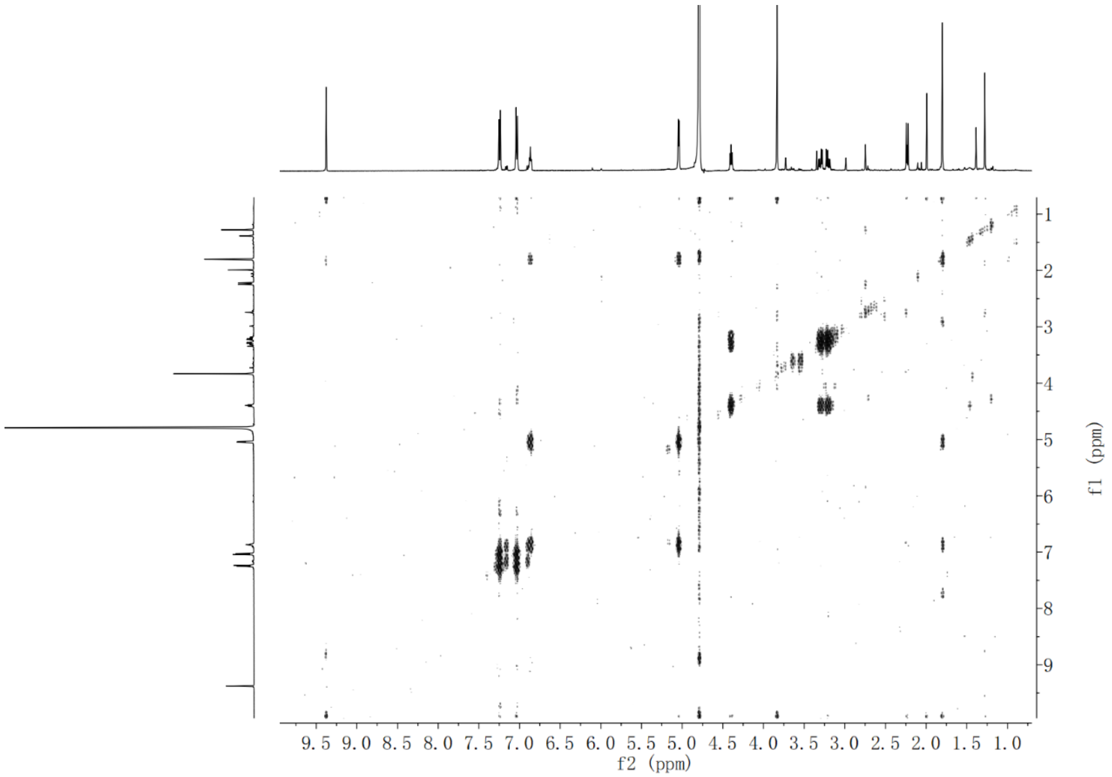


**Supplementary Fig. 88.** $^{1}H$-$^{1}H$ COSY NMR spectrum of compound **19-Me** in $D_2O$

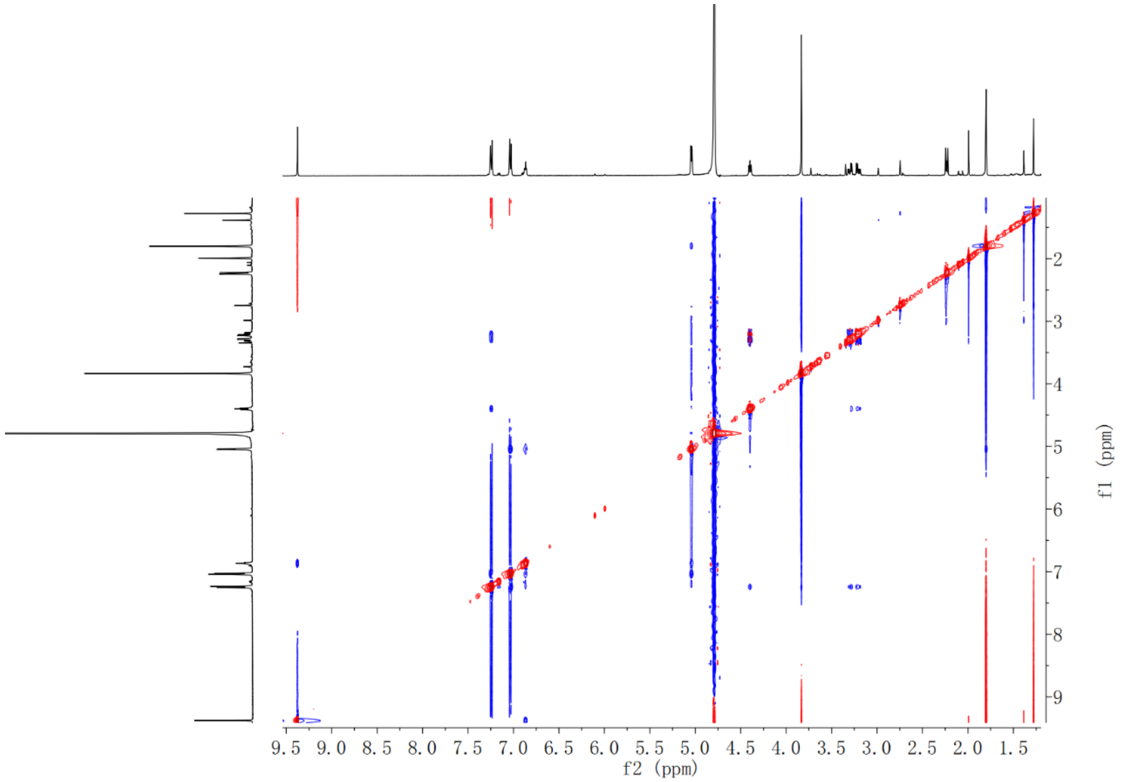


**Supplementary Fig. 89.** NOESY NMR spectrum of compound **19-Me** in $D_2O$

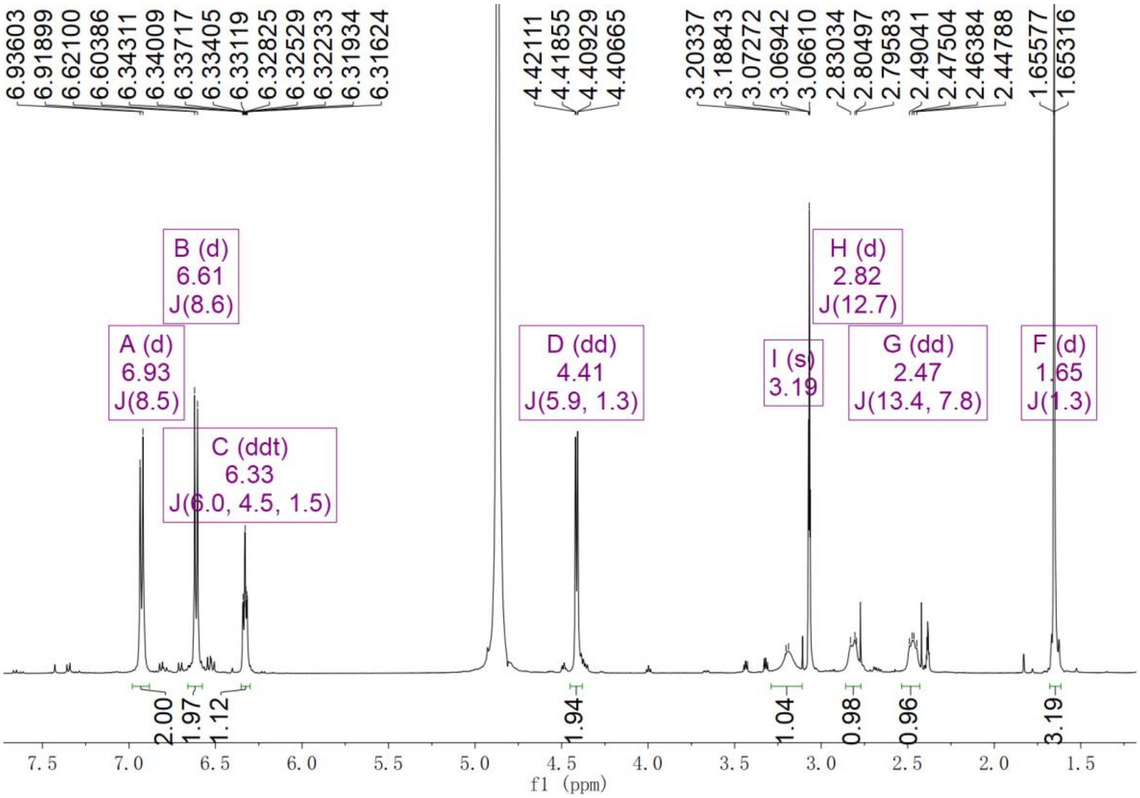


**Supplementary Fig. 90.** $^{1}$H NMR spectrum of compound **20** in $CD_3OD$

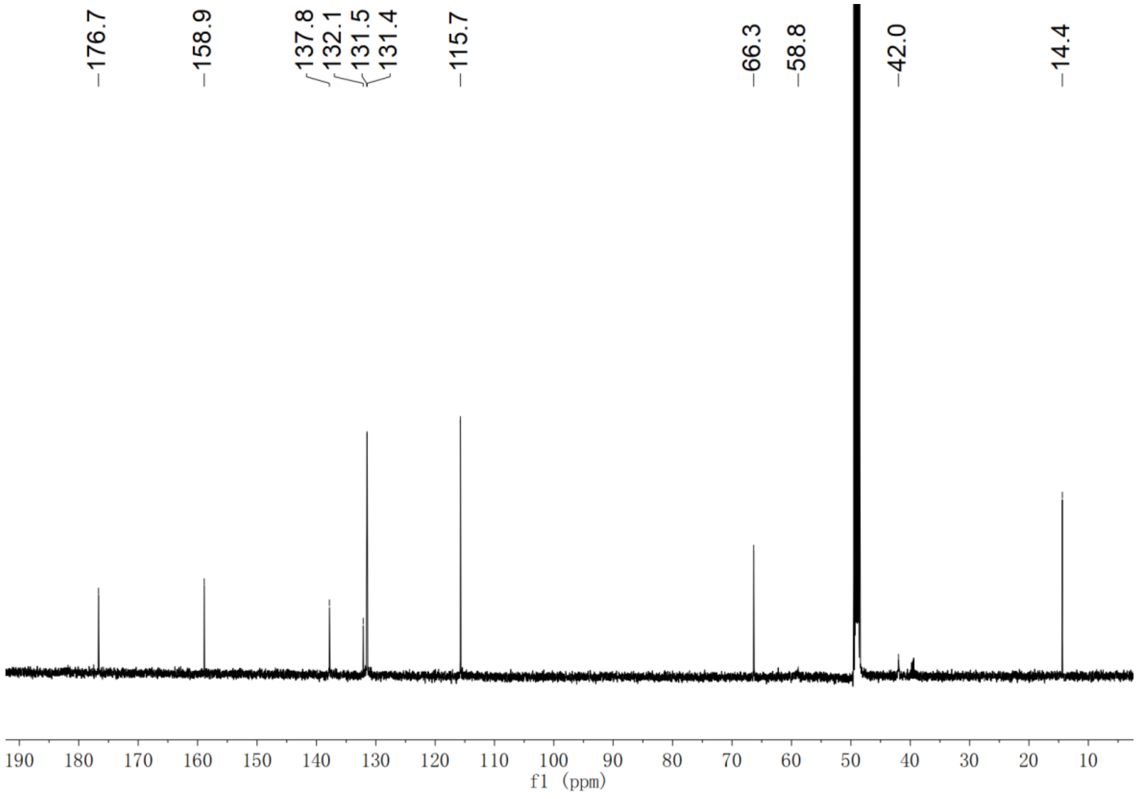


**Supplementary Fig. 91.** $^{13}$C NMR spectrum of compound **20** in $CD_3OD$

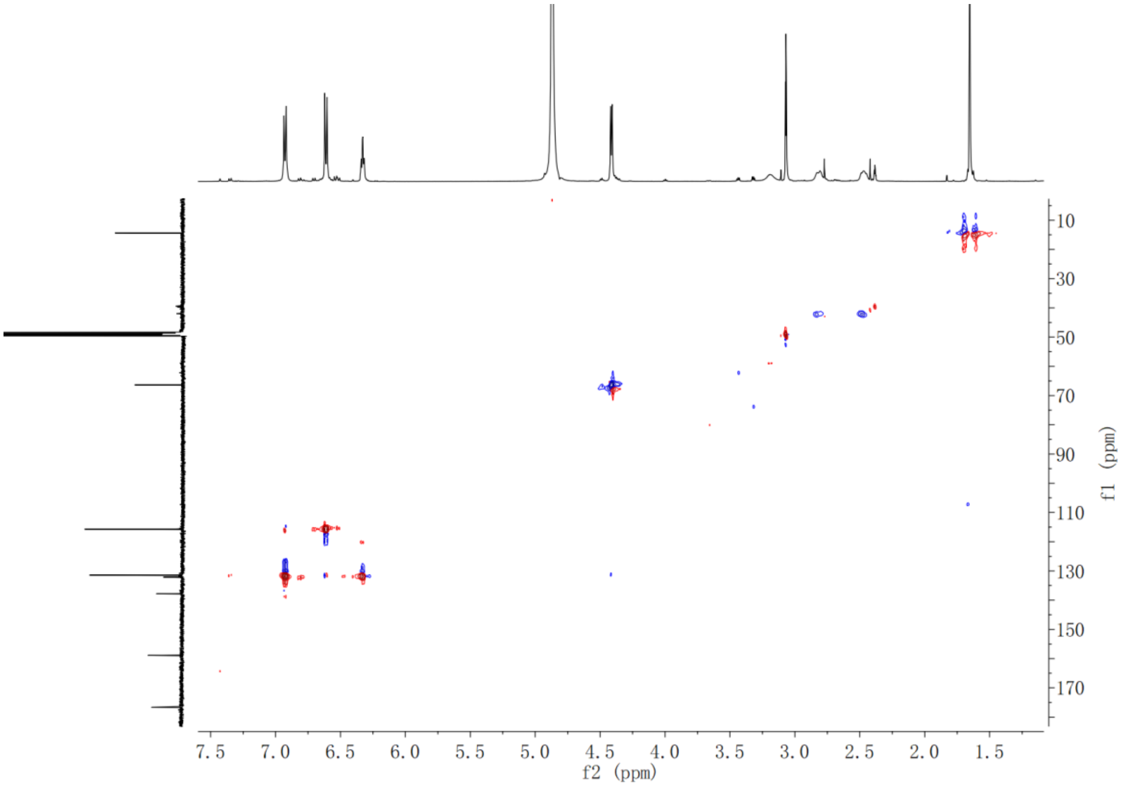


**Supplementary Fig. 92.** HSQC NMR spectrum of compound **20** in $CD_3OD$

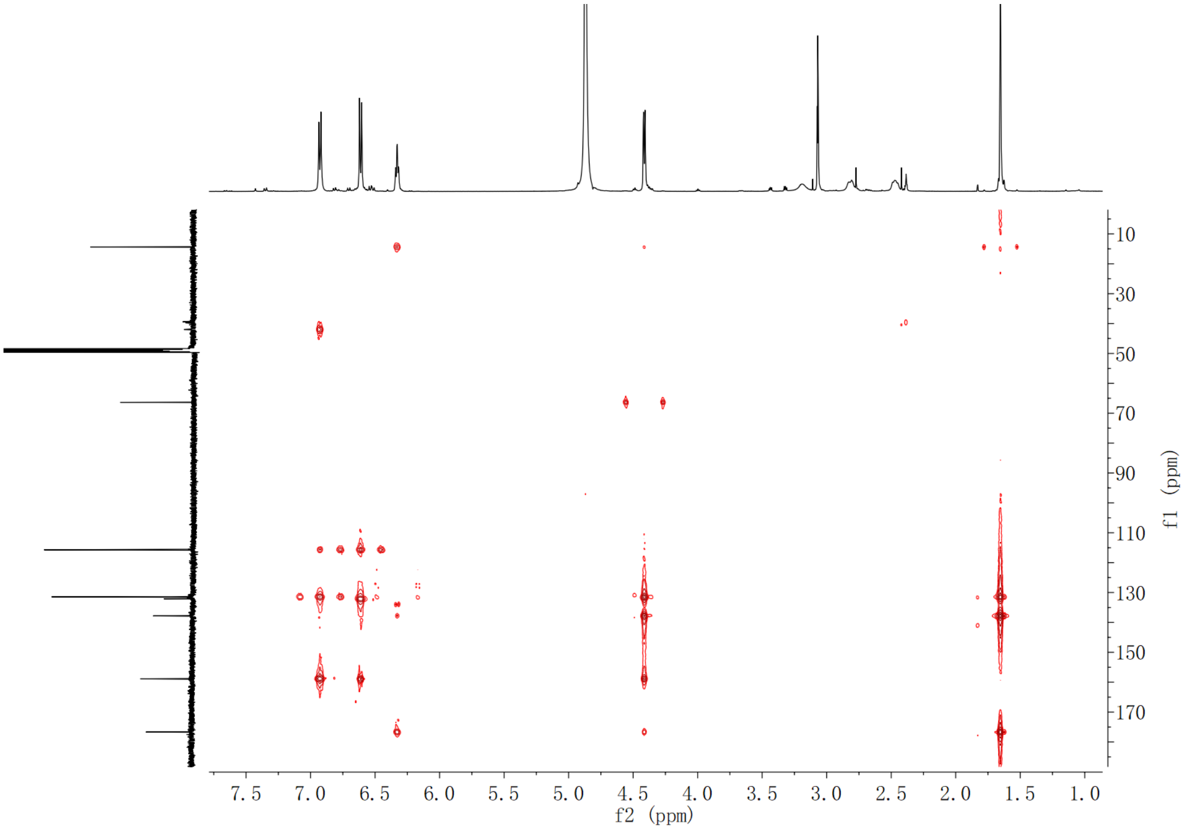


**Supplementary Fig. 93.** HMBC NMR spectrum of compound **20** in $CD_3OD$

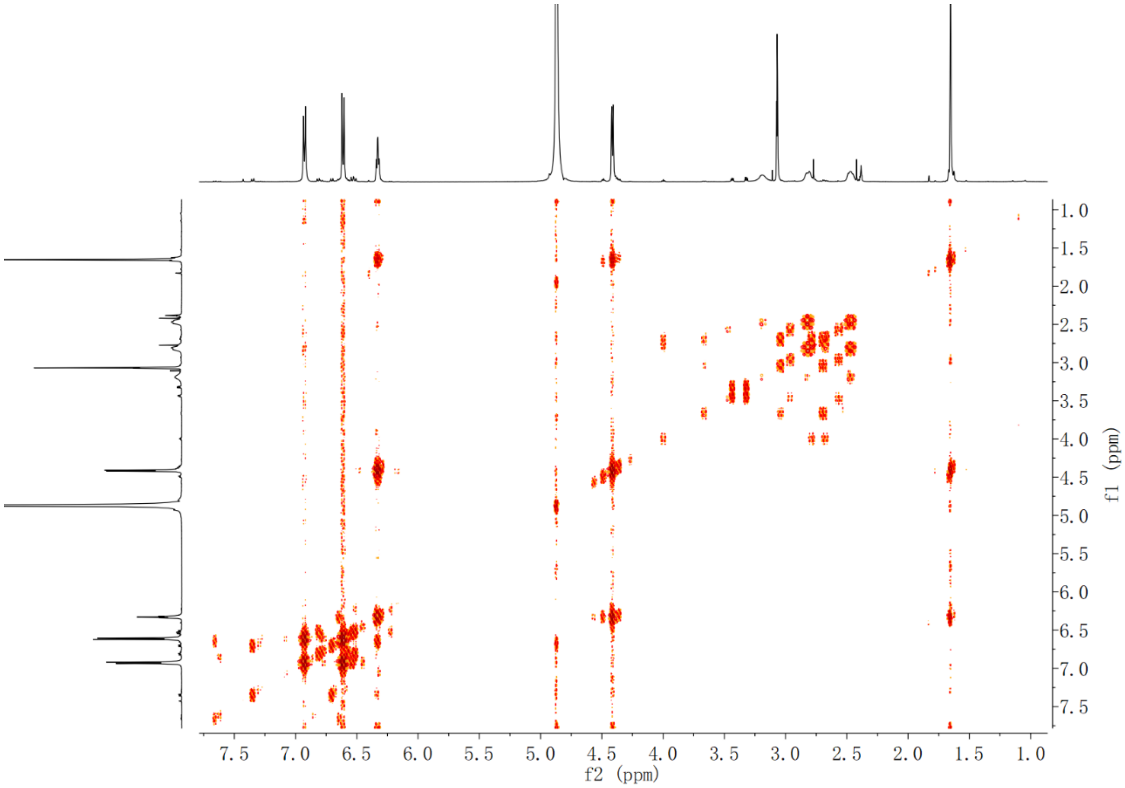


**Supplementary Fig. 94.** $^{1}$H-$^{1}$H COSY NMR spectrum of compound **20** in $CD_3OD$

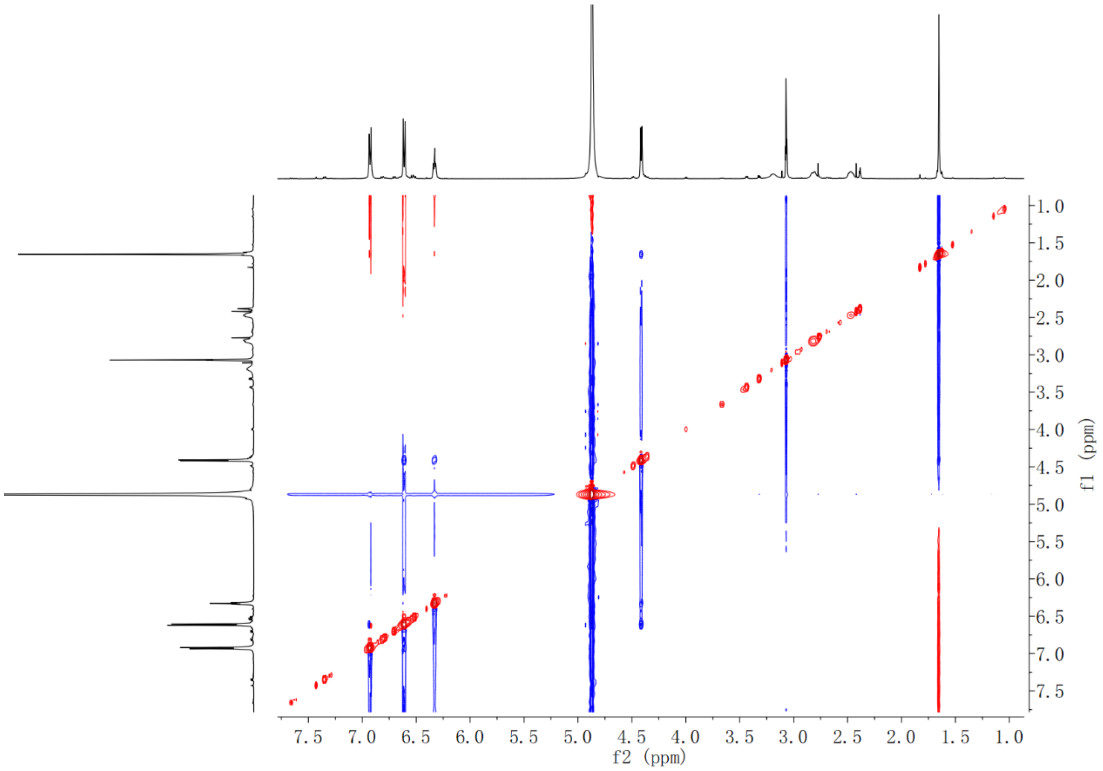

**Supplementary Fig. 95.** NOESY NMR spectrum of compound **20** in $CD_3OD$

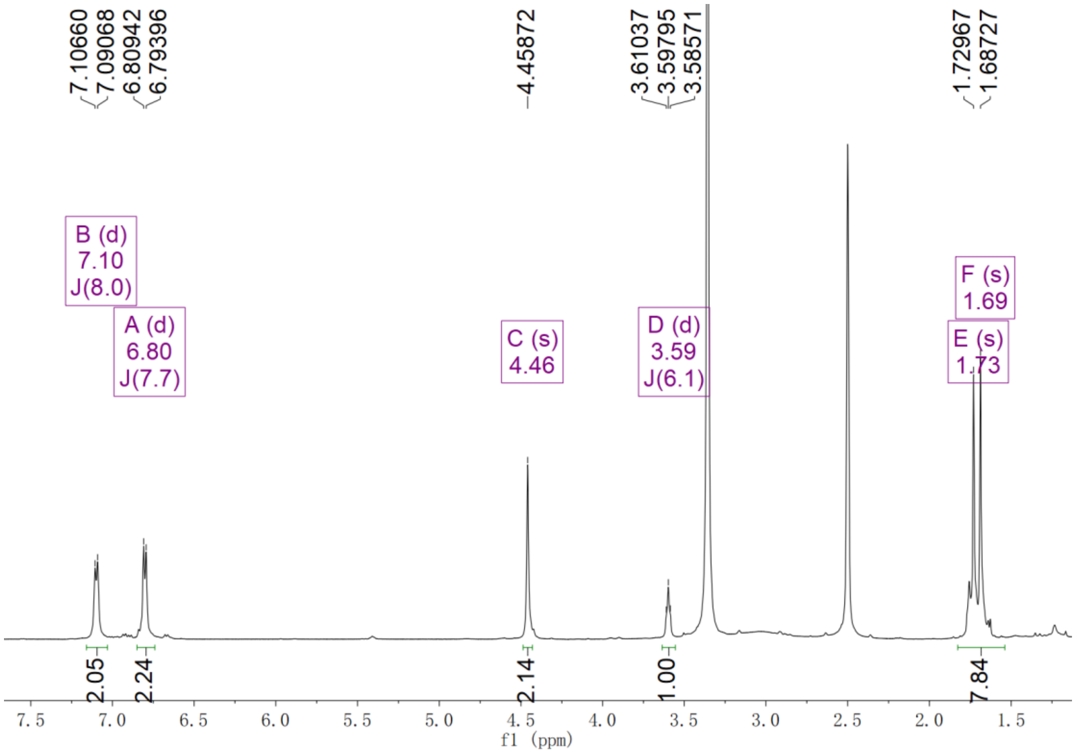


**Supplementary Fig. 96.** $^{1}H$ NMR spectrum of compound **2'-[$d_1$]-7** in DMSO-$d_6$

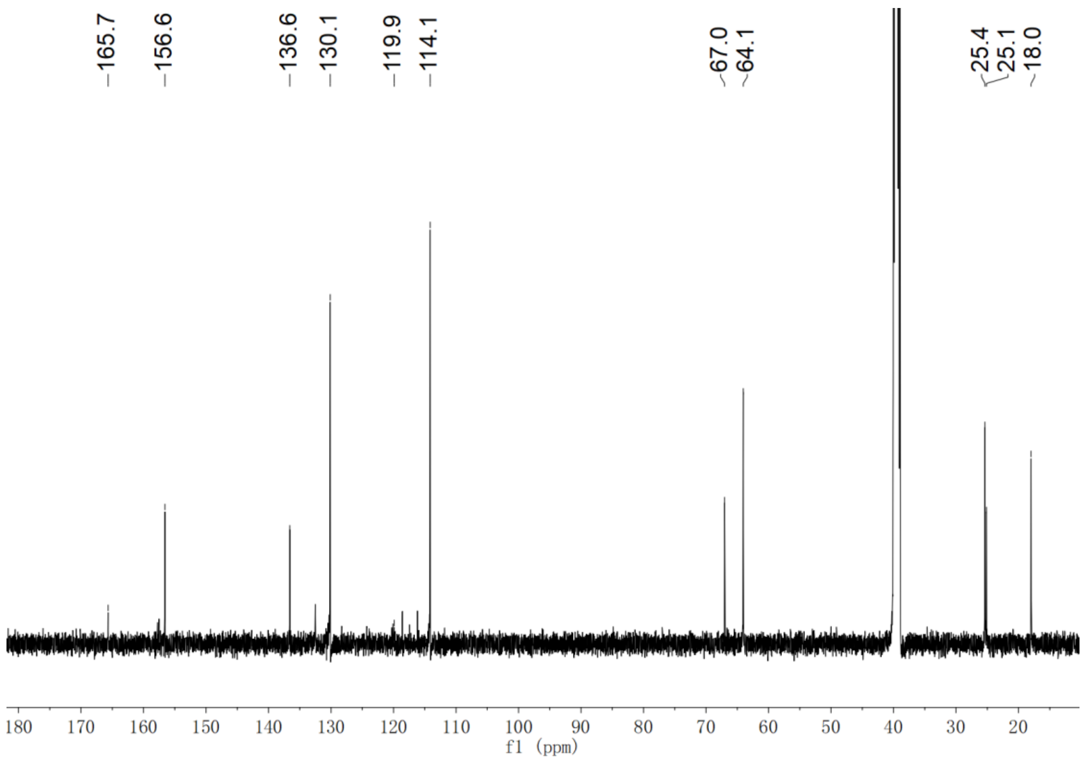


**Supplementary Fig. 97.** $^{13}$C NMR spectrum of compound **2'-[$d_1$]-7** in DMSO-$d_6$

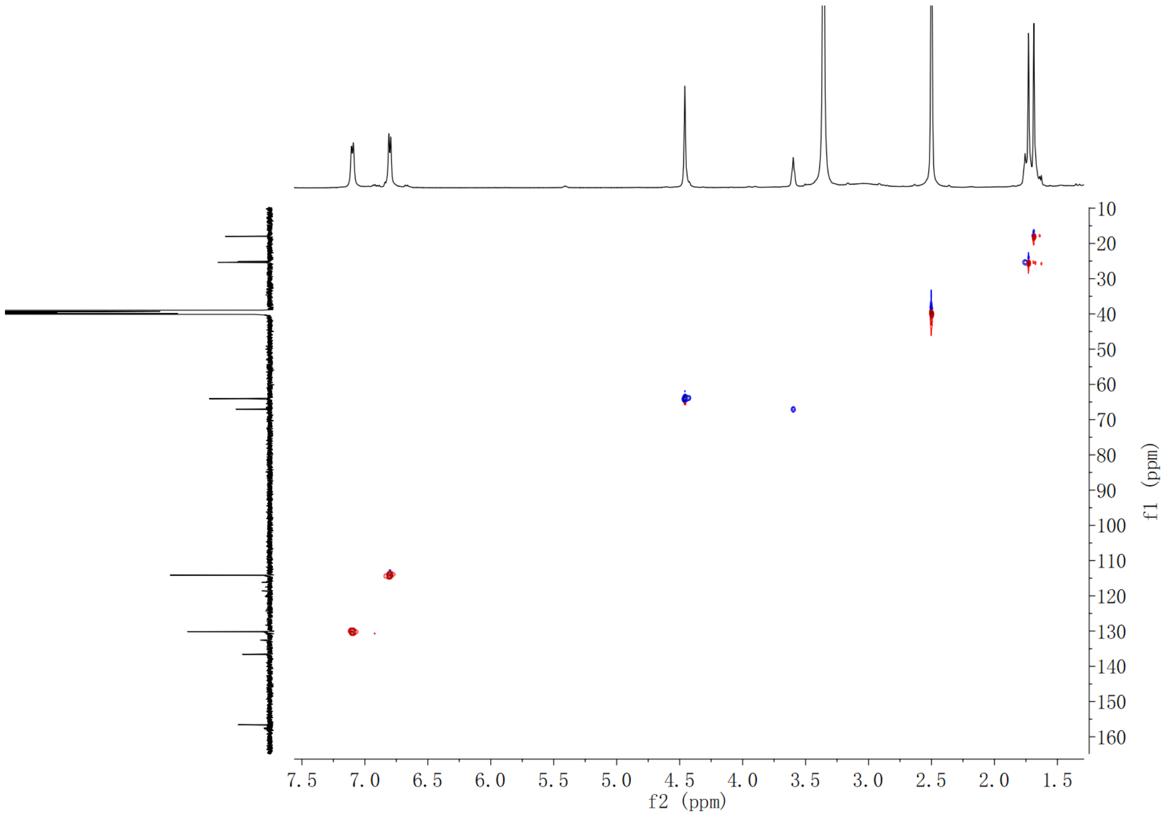


**Supplementary Fig. 98.** HSQC spectrum of compound **2'-[$d_1$]-7** in DMSO-$d_6$

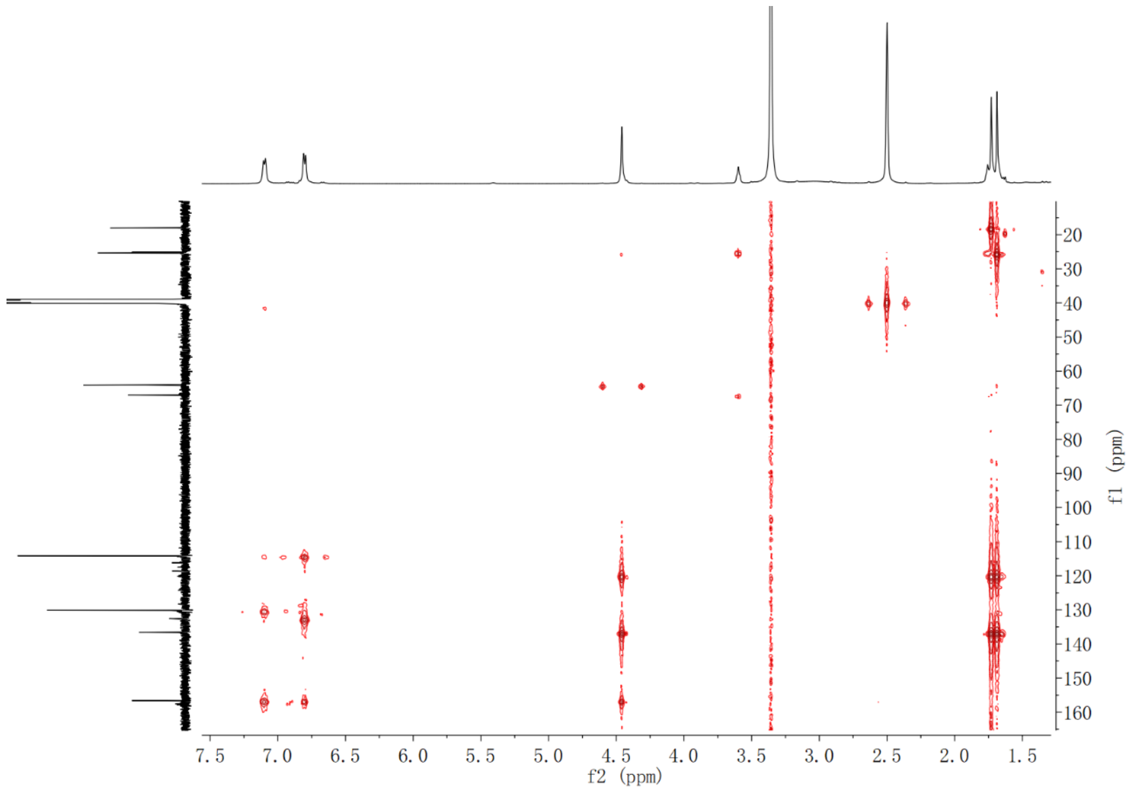


**Supplementary Fig. 99.** HMBC NMR spectrum of compound **2'-[$d_1$]-7** in DMSO-$d_6$

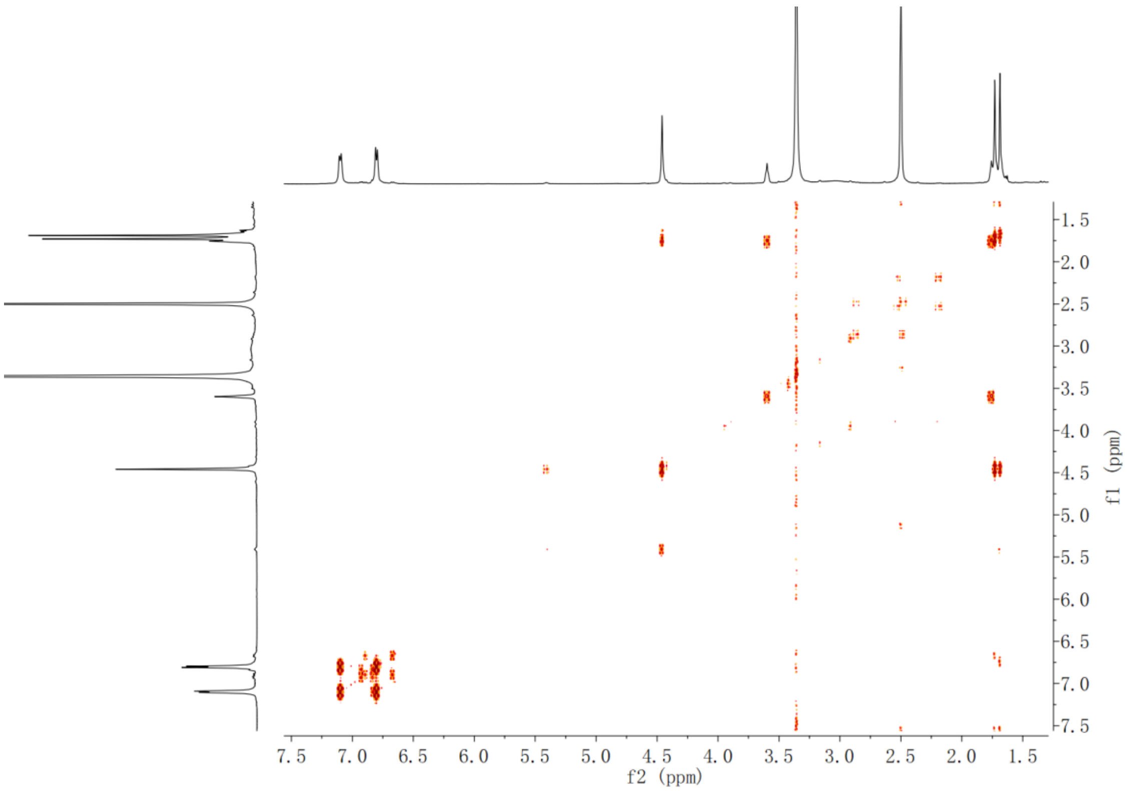


**Supplementary Fig. 100.** $^1$H-$^1$H COSY spectrum of compound **2'-[$d_1$]-7** in DMSO-$d_6$

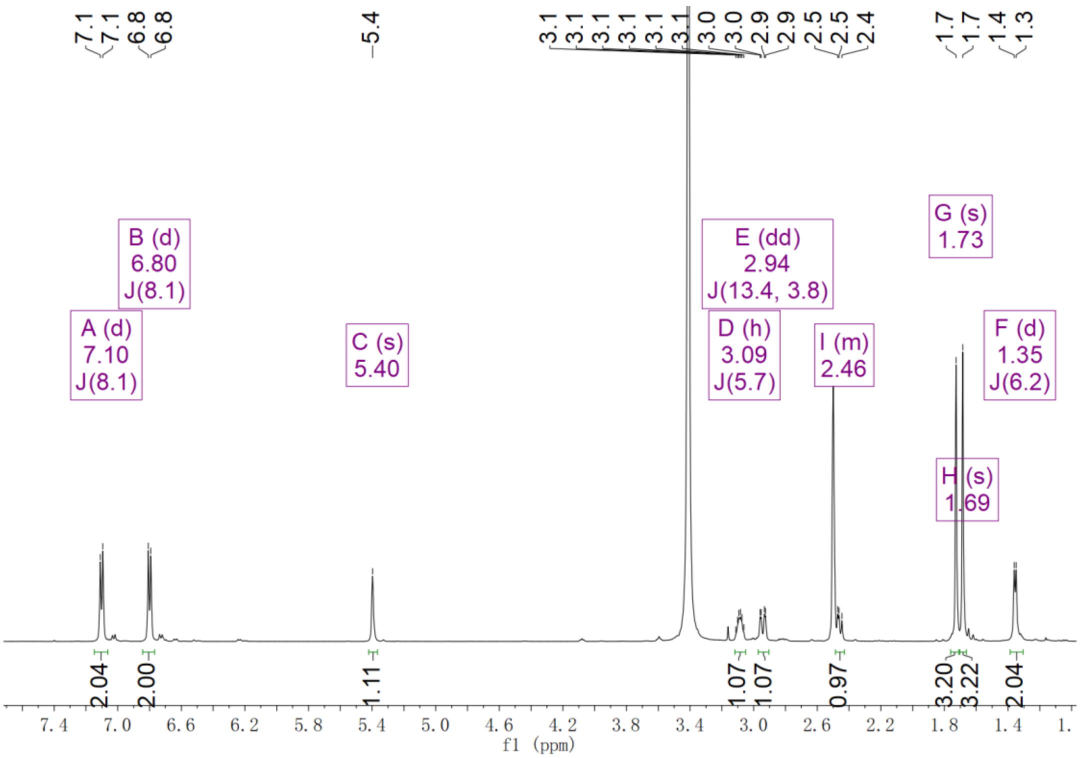

**Supplementary Fig. 101.** $^{1}$H NMR spectrum of compound **1',1'-[$d_2$]-7** in DMSO-$d_6$

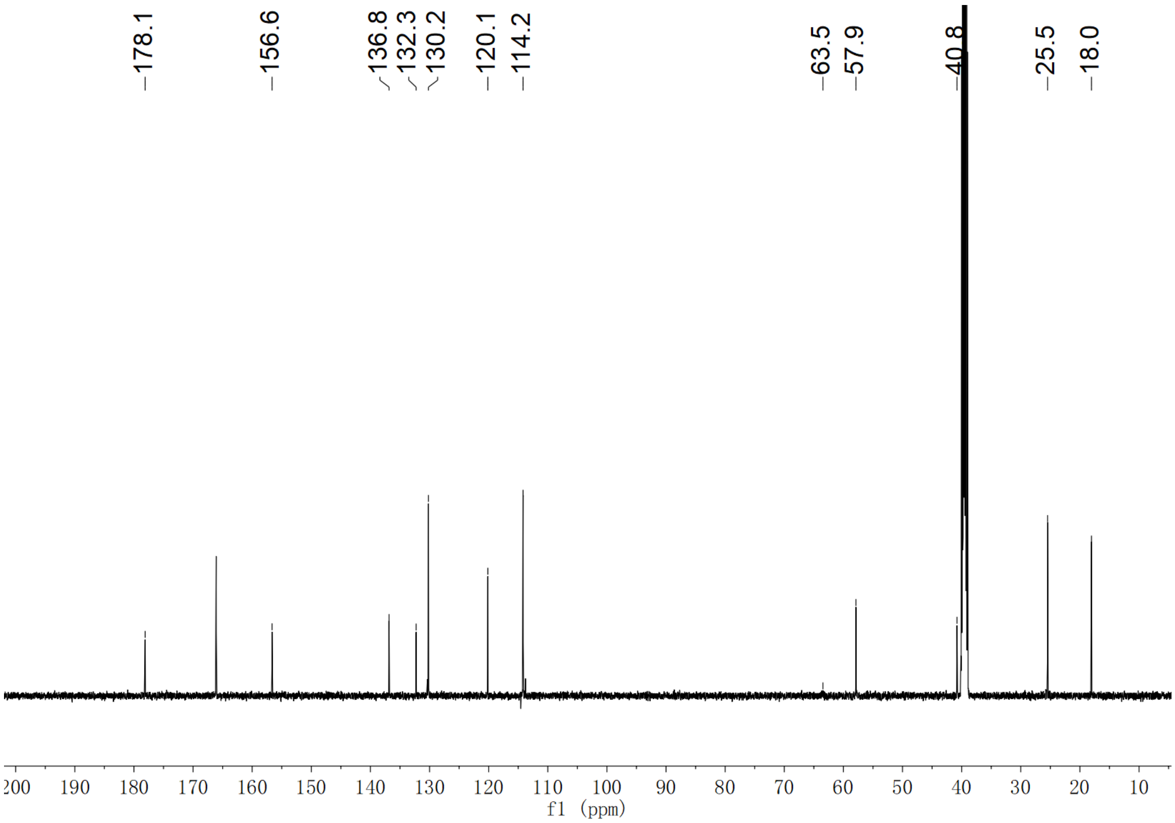

**Supplementary Fig. 102.** $^{13}$C NMR spectrum of compound **1',1'-[$d_2$]-7** in DMSO-$d_6$

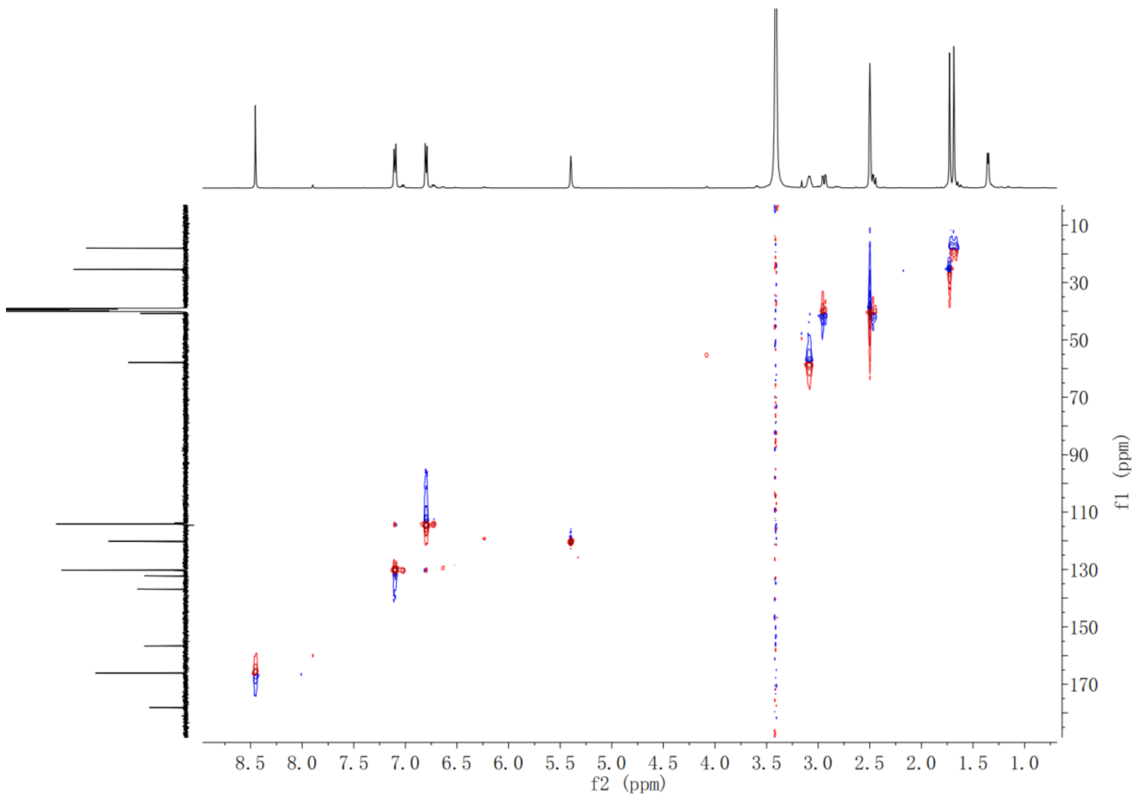


**Supplementary Fig. 103.** HSQC spectrum of compound **1',1'-[*d*$_2$]-7** in DMSO-*d*$_6$

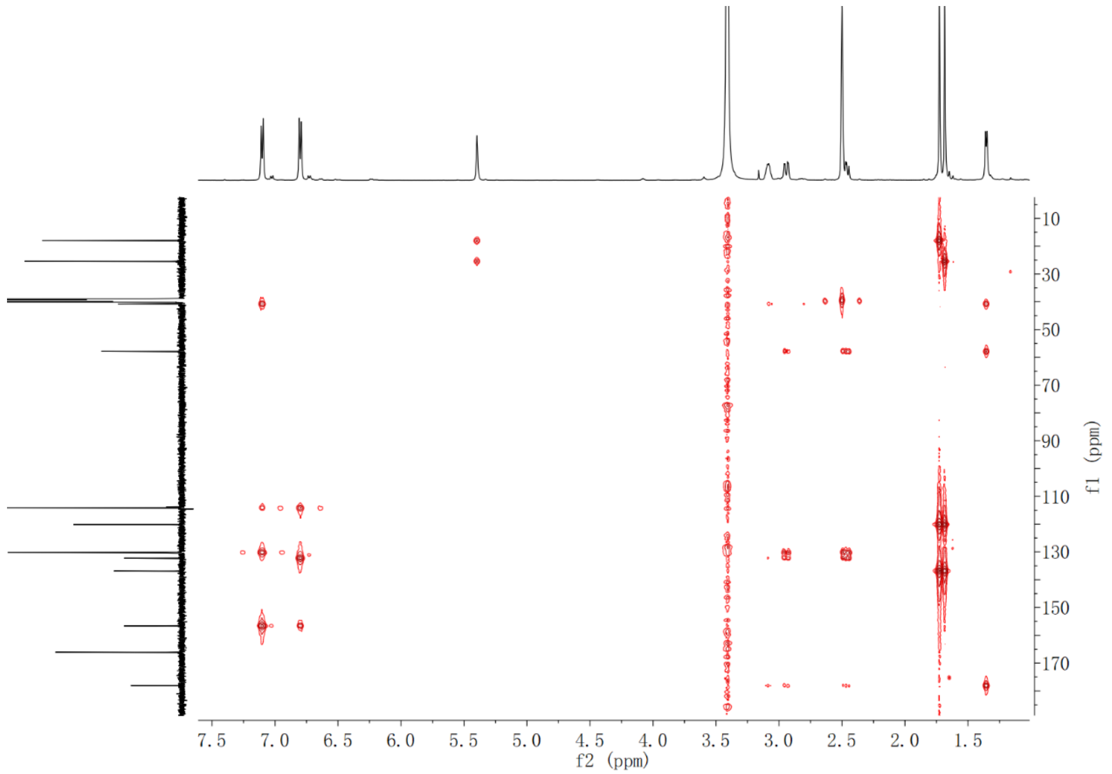


**Supplementary Fig. 104.** HMBC spectrum of compound **1',1'-[*d*$_2$]-7** in DMSO-*d*$_6$

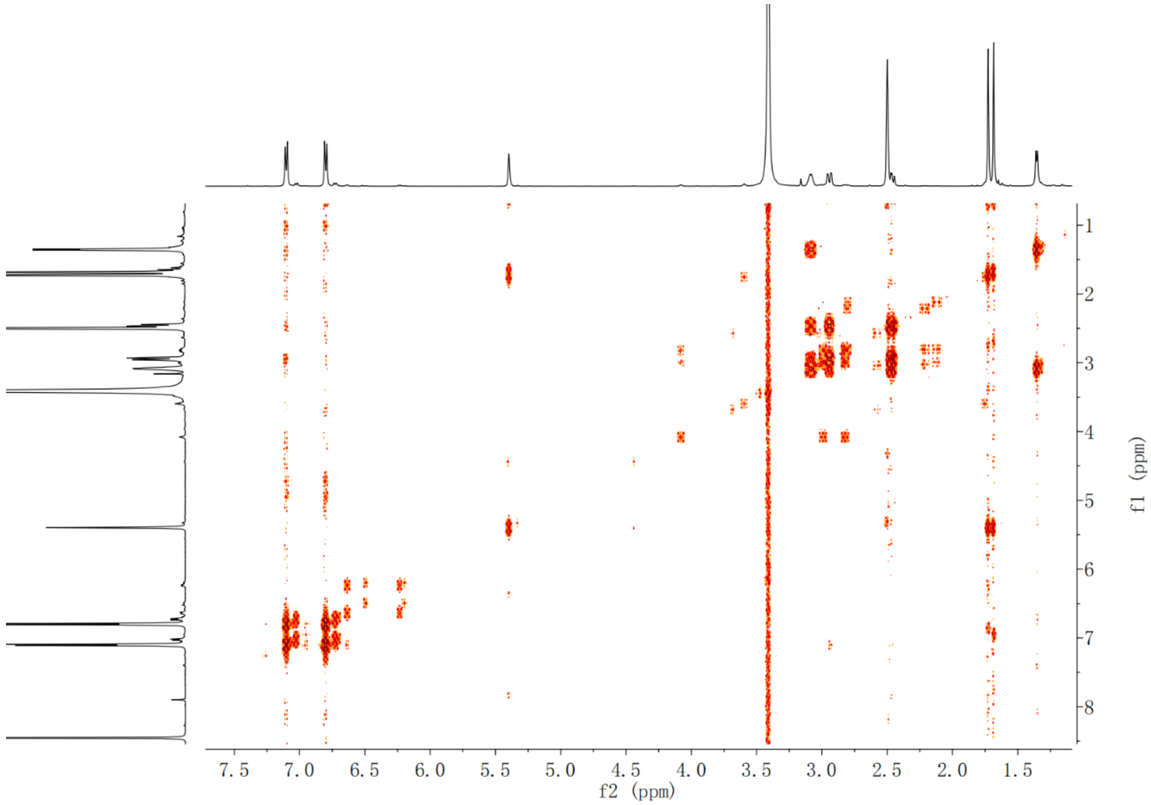


**Supplementary Fig. 105.** $^{1}$H-$^{1}$H COSY spectrum of compound **1',1'-[$d_2$]-7** in DMSO-$d_6$

**Supplementary Computational Data**

Cartesian Coordinates and Energies

For all minimum structures, no imaginary frequency was observed. Energies are reported in this section directly from the output file at the optimization level of theory (B3LYP-D3(BJ) /def2-SVP/IEFPCM(Et2O)). E_SP, H and G are energies combining final single point energy with thermal corrections. (B3LYP-D3(BJ) /def2-TZVP/SMD(Et2O)). All the energies here are in Hartree.

**Int1**

E=-3493.251708

E_SP=-3495.602607

H=-3494.997841

G=-3495.103151

Imag. Freq. 0

Cartesian coordinates

| | | | |
|---|---|---|---|
| Fe | -1.089949 | 0.679829 | 0.465867 |
| N | -1.570598 | -0.975122 | 1.528231 |
| N | -2.970326 | 0.720042 | -0.272246 |
| N | 0.794177 | 0.650287 | 1.219110 |
| N | -0.590646 | 2.322069 | -0.606173 |
| C | -0.745951 | -1.647769 | 2.385326 |
| C | -3.474870 | 1.658250 | -1.129456 |
| C | -2.760665 | -1.648626 | 1.522193 |
| C | -3.979896 | -0.168036 | -0.026022 |
| C | 1.274585 | -0.237923 | 2.144571 |
| C | -1.432090 | 3.026103 | -1.423680 |
| C | 1.826427 | 1.503251 | 0.927886 |
| C | 0.630085 | 2.935561 | -0.681329 |
| C | -1.434416 | -2.797035 | 2.941891 |
| C | -4.861281 | 1.359875 | -1.434202 |
| C | -2.688473 | -2.798697 | 2.402067 |
| C | -5.174342 | 0.221536 | -0.749131 |

| | | | |
|---|---|---|---|
| C | 2.657222 | 0.067694 | 2.456060 |
| C | -0.720699 | 4.130274 | -2.038419 |
| C | 3.001299 | 1.148456 | 1.697966 |
| C | 0.564575 | 4.070475 | -1.581020 |
| H | -0.996787 | -3.499504 | 3.650180 |
| H | -5.493099 | 1.952539 | -2.094634 |
| H | -3.503161 | -3.501155 | 2.573929 |
| H | -6.119215 | -0.319898 | -0.724526 |
| H | 3.276113 | -0.498453 | 3.150584 |
| H | -1.163966 | 4.845181 | -2.730703 |
| H | 3.962962 | 1.653671 | 1.629578 |
| H | 1.402023 | 4.727214 | -1.813733 |
| C | 0.570812 | -1.308701 | 2.685075 |
| C | -2.772138 | 2.734673 | -1.663163 |
| C | 1.767860 | 2.558032 | 0.024919 |
| C | -3.892512 | -1.281321 | 0.803380 |
| H | 1.099390 | -1.942549 | 3.399683 |
| H | -3.313176 | 3.397108 | -2.341645 |
| H | 2.677289 | 3.141061 | -0.127445 |
| H | -4.781081 | -1.907142 | 0.904589 |
| O | -0.518088 | -0.340475 | -1.008886 |
| S | -1.836382 | 1.962892 | 2.226157 |
| C | -0.620540 | 3.295530 | 2.498056 |
| H | 0.370528 | 2.894118 | 2.761920 |
| H | -0.502229 | 3.926394 | 1.603265 |
| H | -0.969699 | 3.931445 | 3.327775 |
| O | -1.381576 | -1.460875 | -1.257800 |
| O | -1.995947 | -1.645853 | -3.979795 |
| H | -1.646027 | -2.544855 | -3.789772 |
| H | -1.239170 | -1.181849 | -4.364923 |
| H | -1.745941 | -1.262961 | -2.147135 |
| C | 4.400041 | -0.778828 | -0.499956 |
| C | 4.065650 | 0.289824 | -1.345389 |
| C | 4.981456 | 1.329041 | -1.540056 |
| C | 6.226658 | 1.320133 | -0.909157 |
| C | 6.557680 | 0.245197 | -0.072136 |
| C | 5.656714 | -0.796962 | 0.130180 |
| H | 3.087653 | 0.342089 | -1.819866 |
| H | 4.702191 | 2.162475 | -2.189881 |
| H | 7.528036 | 0.221290 | 0.430740 |
| H | 5.893772 | -1.635798 | 0.787689 |
| C | 2.269259 | -1.811129 | -0.794466 |
| H | 2.345662 | -1.805342 | -1.893859 |
| H | 1.706914 | -0.909175 | -0.500513 |
| C | 1.543985 | -3.012979 | -0.278754 |
| C | 0.518795 | -3.660740 | -0.868059 |
| H | 1.868076 | -3.332374 | 0.717385 |
| C | -0.223190 | -4.790848 | -0.214483 |
| H | -0.263000 | -5.674598 | -0.870659 |
| H | 0.236594 | -5.071614 | 0.743152 |
| H | -1.264805 | -4.484675 | -0.023069 |
| C | 0.022687 | -3.236135 | -2.189227 |
| O | -0.810739 | -3.865542 | -2.835686 |
| H | 0.502311 | -2.337193 | -2.624150 |
| O | 3.576730 | -1.816738 | -0.225550 |
| H | 6.932988 | 2.138615 | -1.064506 |

**Int2**

E=-3493.262949
E_SP=-3495.610185
H=-3495.004439
G=-3495.107698
Imag. Freq. 0

Cartesian coordinates

| | | | |
|---|---|---|---|
| Fe | -1.023736 | 0.588777 | 0.553122 |
| N | -1.559269 | -1.144522 | 1.443245 |
| N | -2.892089 | 0.741952 | -0.206394 |
| N | 0.856750 | 0.424555 | 1.292141 |
| N | -0.469064 | 2.311780 | -0.358904 |
| C | -0.758612 | -1.924008 | 2.230491 |
| C | -3.356333 | 1.763544 | -0.988623 |
| C | -2.774467 | -1.768499 | 1.385885 |
| C | -3.932789 | -0.131433 | -0.046353 |
| C | 1.314632 | -0.582106 | 2.100078 |
| C | -1.279507 | 3.106220 | -1.123231 |
| C | 1.920557 | 1.260163 | 1.072788 |
| C | 0.768849 | 2.895030 | -0.367000 |
| C | -1.491832 | -3.086429 | 2.692689 |
| C | -4.744106 | 1.530506 | -1.338182 |
| C | -2.747208 | -2.990041 | 2.165907 |

| | | | |
|---|---|---|---|
| C | -5.101922 | 0.350609 | -0.753386 |
| C | 2.716463 | -0.374872 | 2.406565 |
| C | -0.532078 | 4.242448 | -1.626282 |
| C | 3.094056 | 0.768982 | 1.766201 |
| C | 0.744203 | 4.108485 | -1.159388 |
| H | -1.079289 | -3.865872 | 3.331946 |
| H | -5.346779 | 2.192203 | -1.958942 |
| H | -3.588887 | -3.671662 | 2.281400 |
| H | -6.061337 | -0.163752 | -0.788982 |
| H | 3.322437 | -1.041804 | 3.018035 |
| H | -0.946486 | 5.027206 | -2.258088 |
| H | 4.076070 | 1.236845 | 1.728991 |
| H | 1.601490 | 4.760144 | -1.323698 |
| C | 0.573639 | -1.671666 | 2.542732 |
| C | -2.619800 | 2.865435 | -1.412844 |
| C | 1.893412 | 2.410696 | 0.293044 |
| C | -3.888001 | -1.308564 | 0.691582 |
| H | 1.083764 | -2.396571 | 3.179671 |
| H | -3.133902 | 3.597123 | -2.039127 |
| H | 2.822832 | 2.973366 | 0.194459 |
| H | -4.796126 | -1.912527 | 0.735433 |
| O | -0.448774 | -0.282166 | -1.011224 |
| S | -1.744552 | 1.696629 | 2.418585 |
| C | -0.537120 | 3.015203 | 2.780418 |
| H | 0.471684 | 2.604950 | 2.942325 |
| H | -0.479150 | 3.748706 | 1.961349 |
| H | -0.851548 | 3.540092 | 3.696861 |
| O | -1.320866 | -1.367764 | -1.403324 |
| O | -2.690006 | -0.587152 | -3.563675 |
| H | -1.883591 | -2.293794 | -3.489460 |
| H | -2.087355 | 0.100361 | -3.881933 |
| H | -2.427375 | -0.671046 | -2.616564 |
| C | 4.460164 | -0.900738 | -0.753440 |
| C | 4.178664 | 0.315075 | -1.397204 |
| C | 5.138755 | 1.332204 | -1.405451 |
| C | 6.378162 | 1.159620 | -0.786485 |
| C | 6.657947 | -0.060170 | -0.154457 |
| C | 5.712438 | -1.082292 | -0.138887 |
| H | 3.210046 | 0.489705 | -1.861576 |
| H | 4.900469 | 2.277671 | -1.899855 |
| H | 7.623217 | -0.214515 | 0.335044 |
| H | 5.910624 | -2.033462 | 0.359388 |
| C | 2.266785 | -1.742838 | -1.160693 |
| H | 2.298580 | -1.579513 | -2.254014 |
| H | 1.813345 | -0.845494 | -0.713475 |
| C | 1.453118 | -2.950400 | -0.801059 |
| C | 0.213273 | -3.223284 | -1.233738 |
| H | 1.917551 | -3.605933 | -0.056704 |
| C | -0.581315 | -4.393750 | -0.728359 |
| H | -0.904335 | -5.042668 | -1.557729 |
| H | -0.001278 | -4.988291 | -0.007498 |
| H | -1.494858 | -4.038083 | -0.224744 |
| C | -0.519114 | -2.280568 | -2.163318 |
| O | -1.386199 | -2.979751 | -2.993120 |
| H | 0.188062 | -1.652641 | -2.738175 |
| O | 3.592484 | -1.933005 | -0.667244 |
| H | 7.119208 | 1.961990 | -0.794786 |

**Int3**

E=-652.151577

E_SP=-652.893914

H=-652.660641

G=-652.714526

Imag. Freq. 0

Cartesian coordinates

| | | | |
|---|---|---|---|
| O | 1.081239 | -0.706577 | -1.581499 |
| H | 1.100078 | -1.807114 | 0.564355 |
| C | -1.239335 | -0.227217 | 0.600086 |
| C | -2.071563 | -1.213339 | 0.052196 |
| C | -3.216054 | -0.844017 | -0.650539 |
| C | -3.543389 | 0.507951 | -0.815212 |
| C | -2.707492 | 1.482862 | -0.269613 |
| C | -1.552452 | 1.126877 | 0.437199 |
| H | -1.800961 | -2.262296 | 0.186029 |
| H | -3.858153 | -1.619467 | -1.075182 |
| H | -2.945789 | 2.541824 | -0.395610 |
| H | -0.907668 | 1.907741 | 0.838107 |
| C | 0.759767 | 0.241300 | 1.881597 |
| H | 1.387170 | -0.376136 | 2.544266 |
| H | 0.201265 | 0.942409 | 2.521866 |

C 1.633724 1.010939 0.925437
C 2.247366 0.590360 -0.194999
H 1.841434 2.041132 1.237772
C 3.066171 1.497511 -1.064446
H 2.657392 1.507668 -2.087601
H 3.075812 2.527571 -0.681654
H 4.106000 1.135412 -1.136636
C 2.120037 -0.866162 -0.749365
O 2.004879 -1.872397 0.202359
H 3.065517 -1.067165 -1.301744
O -0.143958 -0.684729 1.284141
H -4.440056 0.795673 -1.368017

**Int4**

E=-462.513116
E_SP=-463.031332
H=-462.839305
G=-462.88697
Imag. Freq. 0

Cartesian coordinates
C 0.407319 -0.462516 0.451659
C 0.662485 -0.606869 -0.919714
C 1.910992 -0.232752 -1.430611
C 2.903926 0.285539 -0.597496
C 2.639244 0.430567 0.770420
C 1.402295 0.060865 1.293858
H -0.103494 -0.985365 -1.594930
H 2.099756 -0.346204 -2.501123
H 3.404714 0.835345 1.437184
H 1.180045 0.164928 2.357732
C -1.811050 -1.372257 0.299731
H -2.498860 -1.810198 1.041483
H -1.427454 -2.209151 -0.313213
C -2.573883 -0.398017 -0.583742
C -2.477680 0.915256 -0.572241
H -3.274865 -0.890202 -1.275130
C -1.784857 2.040156 0.067551
H -1.342073 1.734006 1.033125
H -0.970357 2.418753 -0.572913
H -2.472605 2.880411 0.261834
O -0.767601 -0.791136 1.057099
H 3.873255 0.576933 -1.007333

**Int5**

E=-3417.374288
E_SP=-3419.626076
H=-3419.035263
G=-3419.136162
Imag. Freq. 0

Cartesian coordinates
Fe 1.172576 -0.408330 0.104061
N 1.430085 0.123025 -1.838717
N 0.483561 -2.205095 -0.487578
N 2.132558 1.289361 0.659557
N 1.214427 -1.052960 2.005735
C 1.880406 1.339188 -2.312256
C 0.059591 -3.227292 0.327334
C 1.007701 -0.582699 -2.942107
C 0.191127 -2.591878 -1.769872
C 2.480428 2.343053 -0.149049
C 0.696423 -2.227815 2.483479
C 2.432991 1.675223 1.942730
C 1.642555 -0.350075 3.102434
C 1.762893 1.383819 -3.748742
C -0.500783 -4.296414 -0.466332
C 1.225617 0.192477 -4.140601
C -0.417838 -3.901877 -1.770611
C 3.010937 3.426536 0.646159
C 0.805583 -2.274928 3.923459
C 2.987139 3.007879 1.946164
C 1.397920 -1.107394 4.308608
H 2.052736 2.234009 -4.363219
H -0.903739 -5.221709 -0.058767
H 0.979705 -0.145098 -5.145714
H -0.740278 -4.432052 -2.664724
H 3.359515 4.375325 0.242268
H 0.467339 -3.104542 4.541503
H 3.309471 3.541226 2.838504

H 1.650885 -0.769532 5.311925
C 2.363366 2.375824 -1.534857
C 0.154388 -3.247683 1.710371
C 2.220523 0.913047 3.083306
C 0.439132 -1.847422 -2.918121
H 2.668676 3.289431 -2.045286
H -0.223798 -4.128808 2.230104
H 2.511229 1.347779 4.040424
H 0.146463 -2.285165 -3.873149
O -0.308358 0.264854 0.199774
S 3.513593 -1.120595 -0.377122
C 3.458282 -2.658199 -1.318280
H 2.862818 -3.421475 -0.798523
H 2.989373 -2.462732 -2.297465
H 4.485336 -3.011589 -1.489060
O -1.224015 2.125880 -1.487544
H 0.412935 3.644713 -0.303614
C -4.764166 0.153706 0.022520
C -5.634502 -0.847826 0.487139
C -5.245655 -2.186203 0.447736
C -3.985941 -2.546064 -0.049675
C -3.128269 -1.546399 -0.511619
C -3.506485 -0.200441 -0.485772
H -6.609236 -0.549852 0.879279
H -5.931687 -2.954575 0.814171
H -2.138920 -1.798917 -0.890883
H -2.812095 0.553233 -0.846216
C -4.340413 2.511635 -0.202285
H -4.996008 3.397830 -0.210469
H -3.921642 2.394728 -1.214563
C -3.268487 2.655441 0.843730
C -1.986242 2.997215 0.648444
H -3.610709 2.436673 1.862052
C -0.999267 3.040733 1.781738
H -0.515908 4.027154 1.863457
H -1.484750 2.802723 2.738533
H -0.207890 2.291938 1.615530
C -1.447692 3.298677 -0.756212
O -0.310837 4.133032 -0.722177
H -2.189941 3.883805 -1.320802
O -5.226587 1.432381 0.095836
H -0.853397 1.431761 -0.887635
H -3.674558 -3.592748 -0.073132

**Int6**

E=-652.17547
E_SP=-652.917876
H=-652.683883
G=-652.736901
Imag. Freq. 0

Cartesian coordinates
O -3.275146 -0.933761 0.662912
H -0.930000 -1.377176 -0.283972
C 1.395961 -0.300706 0.457092
C 2.134619 -1.322974 -0.159314
C 3.385708 -1.046591 -0.706741
C 3.917215 0.247727 -0.647243
C 3.178069 1.259326 -0.032622
C 1.918372 0.997627 0.519512
H 1.711246 -2.328783 -0.195156
H 3.950895 -1.850528 -1.184502
H 3.577817 2.274981 0.019798
H 1.361468 1.807834 0.987460
C -0.678541 0.323432 1.574731
H -0.116314 0.874279 2.342925
H -1.455874 -0.280218 2.059193
C -1.290918 1.252133 0.572463
C -2.189984 0.867583 -0.437904
H -0.997261 2.305475 0.601688
C -2.658096 1.786370 -1.358826
H -2.331095 2.828961 -1.329683
H -3.362867 1.497726 -2.141731
C -2.671200 -0.579328 -0.563848
O -1.662145 -1.473008 -0.925036
H -3.411521 -0.632671 -1.382042
O 0.184143 -0.670837 0.972838
H -3.515684 -1.869752 0.594578
H 4.897155 0.463128 -1.078081

**Int7**

E=-651.962553
E_SP=-652.711429
H=-652.476895
G=-652.530202
Imag. Freq. 0

Cartesian coordinates

| | | | |
|---|---|---|---|
| O | -4.264262 | -1.186360 | -0.388668 |
| H | -5.437111 | 1.087205 | 0.094903 |
| C | 2.018018 | 0.035718 | -0.010660 |
| C | 2.663746 | 1.275469 | -0.080841 |
| C | 4.056736 | 1.323220 | -0.100151 |
| C | 4.806449 | 0.142256 | -0.050871 |
| C | 4.149095 | -1.087122 | 0.016683 |
| C | 2.750558 | -1.153754 | 0.037516 |
| H | 2.065571 | 2.187513 | -0.119044 |
| H | 4.559405 | 2.291283 | -0.153470 |
| H | 4.724047 | -2.014799 | 0.054722 |
| H | 2.266642 | -2.128745 | 0.091133 |
| C | -0.092888 | -1.081315 | 0.082376 |
| H | 0.173499 | -1.706886 | 0.967877 |
| H | 0.096308 | -1.767821 | -0.777023 |
| C | -1.542792 | -0.937677 | 0.151462 |
| C | -2.362261 | 0.196757 | 0.184024 |
| H | -2.065378 | -1.901500 | 0.182774 |
| C | -1.846256 | 1.470559 | 0.094892 |
| H | -0.775155 | 1.636383 | -0.015879 |
| H | -2.530602 | 2.321878 | 0.124700 |
| C | -3.876097 | -0.036676 | 0.305484 |
| O | -4.498853 | 1.120571 | -0.143178 |
| H | -4.112668 | -0.248601 | 1.364192 |
| O | 0.637005 | 0.088219 | 0.008221 |
| H | -4.198720 | -1.001818 | -1.340315 |
| H | 5.897192 | 0.181872 | -0.065051 |

**Int8**

E=-3417.380613
E_SP=-3419.630456
H=-3419.040106
G=-3419.140618
Imag. Freq. 0

Cartesian coordinates

| | | | |
|---|---|---|---|
| Fe | -0.912490 | -0.656262 | 0.187877 |
| N | -1.741135 | 0.903847 | 1.154778 |
| N | 0.628415 | -0.640108 | 1.516302 |
| N | -2.589353 | -0.873694 | -0.896608 |
| N | -0.199964 | -2.399566 | -0.556021 |
| C | -2.921335 | 1.528813 | 0.831153 |
| C | 1.725033 | -1.480338 | 1.525062 |
| C | -1.174179 | 1.646168 | 2.157145 |
| C | 0.875547 | 0.316374 | 2.476266 |
| C | -3.653584 | -0.011081 | -0.942027 |
| C | 0.996441 | -2.999658 | -0.261681 |
| C | -2.829188 | -1.828277 | -1.849424 |
| C | -0.766094 | -3.145720 | -1.561622 |
| C | -3.109293 | 2.693554 | 1.663051 |
| C | 2.675228 | -1.042332 | 2.517960 |
| C | -2.024223 | 2.762562 | 2.493050 |
| C | 2.149077 | 0.068592 | 3.109072 |
| C | -4.601032 | -0.438542 | -1.944339 |
| C | 1.196225 | -4.155765 | -1.103834 |
| C | -4.088987 | -1.571865 | -2.507931 |
| C | 0.099352 | -4.248698 | -1.910936 |
| H | -3.971016 | 3.356851 | 1.613104 |
| H | 3.628347 | -1.529572 | 2.710859 |
| H | -1.800757 | 3.500106 | 3.261549 |
| H | 2.577403 | 0.688774 | 3.894016 |
| H | -5.532524 | 0.075920 | -2.173453 |
| H | 2.071485 | -4.801657 | -1.065523 |
| H | -4.509296 | -2.188791 | -3.300160 |
| H | -0.122190 | -4.989213 | -2.677432 |
| C | -3.816976 | 1.111972 | -0.141565 |
| C | 1.902521 | -2.577122 | 0.703885 |
| C | -1.991233 | -2.894125 | -2.160138 |
| C | 0.038015 | 1.376662 | 2.783780 |
| H | -4.714973 | 1.712314 | -0.291442 |
| H | 2.827023 | -3.143513 | 0.811309 |
| H | -2.317302 | -3.576849 | -2.945915 |
| H | 0.363240 | 2.064794 | 3.564654 |

| | | | |
|---|---|---|---|
| O | -0.179615 | 0.278692 | -0.933670 |
| S | -1.696397 | -2.198144 | 1.993992 |
| C | -2.247838 | -1.203882 | 3.394676 |
| H | -1.385173 | -0.654862 | 3.808583 |
| H | -3.006881 | -0.470265 | 3.089414 |
| H | -2.641959 | -1.871202 | 4.174771 |
| O | -0.383324 | 3.794807 | -0.059230 |
| H | -0.825245 | 1.741997 | -1.545037 |
| C | 3.711332 | 0.796454 | -0.190361 |
| C | 2.951809 | 0.100079 | -1.142404 |
| C | 3.454128 | -1.083476 | -1.690290 |
| C | 4.696830 | -1.590652 | -1.304447 |
| C | 5.446580 | -0.894909 | -0.346831 |
| C | 4.962355 | 0.289638 | 0.204818 |
| H | 1.955490 | 0.438412 | -1.420633 |
| H | 2.844873 | -1.623940 | -2.418812 |
| H | 6.419699 | -1.277446 | -0.027538 |
| H | 5.532601 | 0.843602 | 0.953262 |
| C | 2.050582 | 2.519476 | 0.077014 |
| H | 1.892362 | 3.310161 | 0.821578 |
| H | 1.253523 | 1.777050 | 0.219089 |
| C | 2.027552 | 3.070457 | -1.325863 |
| C | 0.979339 | 3.557383 | -2.007819 |
| H | 2.995772 | 3.035744 | -1.838572 |
| C | 1.101379 | 3.994962 | -3.442526 |
| H | 0.855910 | 5.065463 | -3.559825 |
| H | 2.112700 | 3.824839 | -3.838508 |
| H | 0.383576 | 3.437503 | -4.068388 |
| C | -0.435854 | 3.654915 | -1.459212 |
| O | -1.239792 | 2.583754 | -1.857456 |
| H | -0.917619 | 4.548288 | -1.905701 |
| O | 3.321184 | 1.951102 | 0.404788 |
| H | -1.289123 | 3.727118 | 0.277653 |
| H | 5.076737 | -2.518414 | -1.738271 |

**Product_allene**

E=-461.940864
E_SP=-462.460254
H=-462.279083
G=-462.325092
Imag. Freq. 0

Cartesian coordinates

| | | | |
|---|---|---|---|
| C | 0.759758 | -0.664673 | 0.126684 |
| C | 0.527261 | 0.670883 | 0.489651 |
| C | 1.565650 | 1.604130 | 0.384844 |
| C | 2.827823 | 1.229533 | -0.078291 |
| C | 3.051611 | -0.104807 | -0.441566 |
| C | 2.028939 | -1.045248 | -0.342097 |
| H | -0.448897 | 0.996724 | 0.844612 |
| H | 1.373641 | 2.641666 | 0.669955 |
| H | 4.033479 | -0.416586 | -0.806546 |
| H | 2.186483 | -2.089376 | -0.619726 |
| C | -1.497949 | -1.368070 | 0.596565 |
| H | -1.521593 | -0.769193 | 1.521863 |
| H | -1.941063 | -2.350572 | 0.818387 |
| C | -2.282747 | -0.694023 | -0.504252 |
| C | -2.884423 | 0.463202 | -0.373219 |
| H | -2.334907 | -1.232048 | -1.459546 |
| C | -3.468563 | 1.624545 | -0.218892 |
| H | -4.498352 | 1.704543 | 0.147197 |
| H | -2.943898 | 2.557013 | -0.457266 |
| O | -0.162312 | -1.662724 | 0.207831 |
| H | 3.629445 | 1.966803 | -0.158134 |

**Product_yne**

E=-461.94046
E_SP=-462.460505
H=-462.27848
G=-462.326017
Imag. Freq. 0

Cartesian coordinates

| | | | |
|---|---|---|---|
| C | 0.831341 | 0.654841 | -0.003775 |
| C | 1.982381 | 0.866349 | -0.781377 |
| C | 3.016148 | -0.067082 | -0.769453 |
| C | 2.919864 | -1.224573 | 0.013994 |
| C | 1.772795 | -1.432293 | 0.781249 |
| C | 0.724577 | -0.503976 | 0.779380 |
| H | 2.041305 | 1.774501 | -1.384495 |

| | | | |
|---|---|---|---|
| H | 3.906543 | 0.110929 | -1.377654 |
| H | 1.679924 | -2.332892 | 1.393461 |
| H | -0.166732 | -0.702067 | 1.373265 |
| C | -1.300208 | 1.514162 | 0.706283 |
| H | -1.772628 | 2.506723 | 0.654280 |
| H | -1.045299 | 1.329796 | 1.766057 |
| C | -2.229614 | 0.491132 | 0.219586 |
| C | -2.987906 | -0.364585 | -0.184773 |
| C | -3.898762 | -1.393904 | -0.672242 |
| H | -4.477050 | -1.040332 | -1.541046 |
| H | -3.345730 | -2.296599 | -0.978582 |
| H | -4.614615 | -1.690159 | 0.111601 |
| O | -0.115000 | 1.631975 | -0.071487 |
| H | 3.730583 | -1.956138 | 0.021774 |

**TS1**

E=-3493.230588
E_SP=-3495.576583
H=-3494.978651
G=-3495.079623
Imag. Freq. -1213.7

Cartesian coordinates

| | | | |
|---|---|---|---|
| Fe | 1.021472 | -0.634093 | 0.514743 |
| N | 1.474006 | 1.091109 | 1.467163 |
| N | 2.916810 | -0.702751 | -0.186408 |
| N | -0.880737 | -0.552611 | 1.203117 |
| N | 0.549080 | -2.339737 | -0.464745 |
| C | 0.633721 | 1.805622 | 2.275330 |
| C | 3.442661 | -1.696556 | -0.965947 |
| C | 2.667460 | 1.760728 | 1.459455 |
| C | 3.921299 | 0.202178 | 0.022983 |
| C | -1.383153 | 0.391966 | 2.059064 |
| C | 1.412590 | -3.093673 | -1.212396 |
| C | -1.911958 | -1.413061 | 0.926961 |
| C | -0.671124 | -2.955515 | -0.538279 |
| C | 1.315126 | 2.974304 | 2.796178 |
| C | 4.832880 | -1.412027 | -1.262598 |
| C | 2.580619 | 2.947697 | 2.286023 |
| C | 5.128939 | -0.228439 | -0.650646 |
| C | -2.778561 | 0.117924 | 2.337678 |
| C | 0.717700 | -4.235386 | -1.773830 |
| C | -3.108944 | -1.000949 | 1.630843 |
| C | -0.580542 | -4.145811 | -1.359576 |
| H | 0.863800 | 3.709724 | 3.460806 |
| H | 5.479365 | -2.044384 | -1.869694 |
| H | 3.393808 | 3.655212 | 2.442385 |
| H | 6.070477 | 0.318967 | -0.645878 |
| H | -3.414920 | 0.730020 | 2.974864 |
| H | 1.179284 | -4.992836 | -2.406198 |
| H | -4.074033 | -1.498348 | 1.555571 |
| H | -1.412306 | -4.814999 | -1.576033 |
| C | -0.689294 | 1.485901 | 2.562610 |
| C | 2.756467 | -2.810534 | -1.439295 |
| C | -1.830703 | -2.527006 | 0.099920 |
| C | 3.813764 | 1.361739 | 0.782043 |
| H | -1.233693 | 2.159179 | 3.227135 |
| H | 3.315964 | -3.514153 | -2.058679 |
| H | -2.738704 | -3.114281 | -0.044884 |
| H | 4.697971 | 1.995882 | 0.865640 |
| O | 0.475140 | 0.263841 | -1.079775 |
| S | 1.713382 | -1.767141 | 2.359948 |
| C | 0.556384 | -3.149733 | 2.632581 |
| H | -0.476449 | -2.791937 | 2.763047 |
| H | 0.568714 | -3.862164 | 1.793453 |
| H | 0.859561 | -3.683095 | 3.547716 |
| O | 1.319071 | 1.388911 | -1.385343 |
| O | 2.615523 | 1.175567 | -3.391916 |
| H | 2.061593 | 2.203012 | -3.388921 |
| H | 2.131002 | 0.596643 | -3.998164 |
| H | 2.010516 | 1.074496 | -2.226619 |
| C | -4.435144 | 0.922873 | -0.676840 |
| C | -4.177295 | -0.232023 | -1.432658 |
| C | -5.146915 | -1.237058 | -1.512483 |
| C | -6.371863 | -1.111697 | -0.854612 |
| C | -6.627655 | 0.048243 | -0.109756 |
| C | -5.672745 | 1.057737 | -0.022337 |
| H | -3.219690 | -0.370795 | -1.931086 |
| H | -4.927634 | -2.136058 | -2.094628 |
| H | -7.581271 | 0.164734 | 0.412004 |
| H | -5.851256 | 1.961425 | 0.563948 |

| | | | |
|---|---|---|---|
| C | -2.242606 | 1.783642 | -1.051558 |
| H | -2.299010 | 1.697672 | -2.152022 |
| H | -1.778688 | 0.858742 | -0.677220 |
| C | -1.424981 | 2.966905 | -0.628670 |
| C | -0.213307 | 3.302589 | -1.097718 |
| H | -1.862988 | 3.549645 | 0.188496 |
| C | 0.579964 | 4.454301 | -0.552837 |
| H | 0.833859 | 5.165486 | -1.354249 |
| H | 0.031920 | 4.978785 | 0.243331 |
| H | 1.532707 | 4.092142 | -0.134233 |
| C | 0.465707 | 2.501636 | -2.177676 |
| O | 1.329542 | 3.143679 | -2.934458 |
| H | -0.238148 | 1.817257 | -2.696926 |
| O | -3.556626 | 1.936504 | -0.514948 |
| H | -7.120394 | -1.904479 | -0.919552 |

**TS2**

E=-3493.235287
E_SP=-3495.583654
H=-3494.980608
G=-3495.083209
Imag. Freq. -40.57

Cartesian coordinates

| | | | |
|---|---|---|---|
| Fe | 1.484478 | -0.265890 | 0.280995 |
| N | -0.227115 | -0.517074 | 1.321579 |
| N | 1.697754 | 1.570112 | 1.106981 |
| N | 1.337988 | -2.155042 | -0.436674 |
| N | 3.278526 | -0.070398 | -0.645037 |
| C | -1.053724 | -1.607445 | 1.267357 |
| C | 2.725306 | 2.445657 | 0.879809 |
| C | -0.850477 | 0.405450 | 2.114784 |
| C | 0.814166 | 2.206053 | 1.932723 |
| C | 0.307359 | -3.026347 | -0.218932 |
| C | 4.088073 | 1.031865 | -0.615888 |
| C | 2.188320 | -2.765123 | -1.320632 |
| C | 3.865939 | -0.968070 | -1.495331 |
| C | -2.249830 | -1.363936 | 2.047367 |
| C | 2.483623 | 3.685256 | 1.588310 |
| C | -2.119753 | -0.114633 | 2.577571 |
| C | 1.296185 | 3.535110 | 2.243992 |
| C | 0.519092 | -4.243047 | -0.972870 |
| C | 5.243299 | 0.819372 | -1.462212 |
| C | 1.687684 | -4.080460 | -1.659602 |
| C | 5.103904 | -0.423514 | -2.010829 |
| H | -3.080224 | -2.060598 | 2.138720 |
| H | 3.149185 | 4.547047 | 1.571505 |
| H | -2.818224 | 0.431508 | 3.209431 |
| H | 0.776053 | 4.246860 | 2.883448 |
| H | -0.156199 | -5.097378 | -0.969976 |
| H | 6.043613 | 1.543163 | -1.609523 |
| H | 2.180327 | -4.773154 | -2.340433 |
| H | 5.765815 | -0.940299 | -2.704330 |
| C | -0.810054 | -2.783927 | 0.571713 |
| C | 3.842320 | 2.205240 | 0.089646 |
| C | 3.366178 | -2.223692 | -1.820801 |
| C | -0.371060 | 1.671205 | 2.419829 |
| H | -1.568530 | -3.565926 | 0.628293 |
| H | 4.579789 | 3.005064 | 0.005700 |
| H | 3.944270 | -2.831486 | -2.518615 |
| H | -0.980402 | 2.299405 | 3.071165 |
| O | 0.697819 | 0.363062 | -1.065697 |
| S | 2.587097 | -1.102072 | 2.213441 |
| C | 3.884691 | -2.251783 | 1.649404 |
| H | 4.632069 | -1.748525 | 1.016528 |
| H | 4.398235 | -2.660894 | 2.534457 |
| H | 3.461121 | -3.091264 | 1.076439 |
| O | -1.266692 | 1.522371 | -0.516911 |
| O | -1.368765 | -1.001513 | -2.166493 |
| H | 0.782265 | 2.185132 | -1.379148 |
| H | -0.568321 | -0.507423 | -1.857178 |
| H | -1.827938 | -1.127392 | -1.324872 |
| C | -4.864038 | -0.135775 | -0.301532 |
| C | -5.556537 | -0.382614 | 0.899273 |
| C | -5.820886 | -1.687454 | 1.304658 |
| C | -5.398959 | -2.774950 | 0.526977 |
| C | -4.718409 | -2.528171 | -0.666534 |
| C | -4.455673 | -1.221268 | -1.094518 |
| H | -5.865867 | 0.474217 | 1.501124 |
| H | -6.355068 | -1.859645 | 2.242814 |
| H | -4.379846 | -3.362200 | -1.286746 |

H -3.913166 -1.062941 -2.024872
C -3.920030 1.501596 -1.796778
H -4.594568 1.369868 -2.664834
H -3.059740 0.831891 -1.912284
C -3.481505 2.928412 -1.664352
C -2.205135 3.335008 -1.560408
H -4.281126 3.676641 -1.612847
C -1.808897 4.762994 -1.336351
H -1.250252 4.852155 -0.390640
H -1.123792 5.110861 -2.125416
H -2.690082 5.420499 -1.300571
C -1.054430 2.310731 -1.602809
O 0.192979 2.949334 -1.571551
H -1.137529 1.709824 -2.536289
O -4.645595 1.162542 -0.604078
H -5.600334 -3.798728 0.849718

**TS3**

E=-652.134021
E_SP=-652.87809
H=-652.647473
G=-652.701568
Imag. Freq. -527.21

Cartesian coordinates
O 2.886822 1.459910 -1.123276
H 0.981041 1.343983 1.236265
C -1.323131 0.040495 0.637835
C -1.753615 -1.160910 0.063498
C -2.848546 -1.146597 -0.808479
C -3.507532 0.045417 -1.113858
C -3.064062 1.241358 -0.536687
C -1.976713 1.243532 0.335434
H -1.247807 -2.101645 0.275781
H -3.181491 -2.086372 -1.255318
H -3.568241 2.182776 -0.766982
H -1.620454 2.167873 0.794506
C 0.543987 -1.001068 1.785621
H -0.088638 -1.825457 2.149962
H 1.183798 -0.688482 2.624916
C 1.384694 -1.478313 0.627303
C 1.999372 -0.803557 -0.335997
H 1.528087 -2.569938 0.598901
C 2.768991 -1.286595 -1.503881
H 3.798039 -0.898464 -1.479781
H 2.786951 -2.389236 -1.517302
H 2.319399 -0.930986 -2.444819
C 1.913890 1.150185 -0.414818
O 1.849317 1.606929 0.868807
H 0.903141 1.034916 -0.888275
O -0.268579 0.139645 1.511279
H -4.358315 0.045479 -1.798154

**TS4**

E=-3417.346437
E_SP=-3419.590548
H=-3419.007042
G=-3419.106022
Imag. Freq. -1932.15

Cartesian coordinates
Fe 1.259303 0.466898 -0.088276
N 1.394194 0.006872 1.891963
N 0.302265 2.172838 0.401817
N 2.320677 -1.188040 -0.543612
N 1.258405 0.986567 -2.018572
C 1.891427 -1.161799 2.424304
C -0.218589 3.086245 -0.479118
C 0.890390 0.725569 2.948870
C -0.059745 2.598249 1.652020
C 2.683145 -2.189369 0.323671
C 0.615269 2.065386 -2.566430
C 2.756783 -1.579566 -1.784845
C 1.836641 0.314158 -3.064862
C 1.714526 -1.167283 3.858009
C -0.919303 4.129331 0.237675
C 1.102850 0.007364 4.185559
C -0.814479 3.829208 1.565019
C 3.354161 -3.250259 -0.394388
C 0.789683 2.076121 -4.001666

| | | | |
|---|---|---|---|
| C | 3.409731 | -2.865109 | -1.703634 |
| C | 1.557512 | 0.990602 | -4.311038 |
| H | 2.021443 | -1.979603 | 4.514221 |
| H | -1.421656 | 4.972971 | -0.232104 |
| H | 0.798065 | 0.364435 | 5.167573 |
| H | -1.215236 | 4.371004 | 2.419727 |
| H | 3.734840 | -4.161569 | 0.063247 |
| H | 0.374053 | 2.828706 | -4.669401 |
| H | 3.842578 | -3.394981 | -2.550200 |
| H | 1.904866 | 0.657351 | -5.287353 |
| C | 2.474988 | -2.189397 | 1.701099 |
| C | -0.082289 | 3.040151 | -1.860054 |
| C | 2.558228 | -0.869736 | -2.962381 |
| C | 0.230003 | 1.942704 | 2.845662 |
| H | 2.810783 | -3.065686 | 2.256757 |
| H | -0.548037 | 3.839828 | -2.437718 |
| H | 2.963177 | -1.294399 | -3.881950 |
| H | -0.125211 | 2.401716 | 3.769312 |
| O | -0.263527 | -0.377279 | -0.169021 |
| S | 3.405512 | 1.358165 | 0.410596 |
| C | 3.198758 | 2.899299 | 1.326505 |
| H | 2.563493 | 3.605968 | 0.774487 |
| H | 2.717010 | 2.687659 | 2.295743 |
| H | 4.190628 | 3.333915 | 1.516034 |
| O | -1.208112 | -2.196852 | 1.528395 |
| H | 0.637757 | -3.433982 | 0.469735 |
| C | -4.849177 | -0.499606 | -0.120074 |
| C | -5.754651 | 0.465821 | -0.592019 |
| C | -5.484617 | 1.822597 | -0.417684 |
| C | -4.308735 | 2.236457 | 0.222013 |
| C | -3.417998 | 1.271862 | 0.695809 |
| C | -3.680439 | -0.093198 | 0.539926 |
| H | -6.659058 | 0.126398 | -1.101359 |
| H | -6.195579 | 2.563178 | -0.793451 |
| H | -2.491654 | 1.573804 | 1.186364 |
| H | -2.962941 | -0.816526 | 0.919172 |
| C | -4.219159 | -2.822637 | -0.119609 |
| H | -4.761766 | -3.759573 | -0.326839 |
| H | -3.919566 | -2.838274 | 0.941177 |
| C | -3.041737 | -2.660680 | -1.038179 |
| C | -1.733882 | -2.842994 | -0.737179 |
| H | -3.303293 | -2.291643 | -2.036569 |
| C | -0.681810 | -2.387338 | -1.640167 |
| H | 0.256091 | -2.954459 | -1.636365 |
| H | -1.004800 | -2.087521 | -2.644214 |
| H | -0.362292 | -1.277182 | -1.046529 |
| C | -1.292695 | -3.293455 | 0.662251 |
| O | -0.099037 | -4.042448 | 0.628672 |
| H | -2.037987 | -3.985302 | 1.082377 |
| O | -5.189907 | -1.801218 | -0.346322 |
| H | -0.822651 | -1.432164 | 1.023011 |
| H | -4.085965 | 3.298482 | 0.345413 |

**TS5**

E=-651.936084

E_SP=-652.682481

H=-652.450242

G=-652.502402

Imag. Freq. -459.73

Cartesian coordinates

| | | | |
|---|---|---|---|
| O | -1.570725 | -1.364587 | 1.134755 |
| H | -3.936729 | -0.642770 | -0.663631 |
| C | 1.215680 | -0.328362 | -0.499198 |
| C | 1.708429 | 0.979520 | -0.499257 |
| C | 2.976587 | 1.228494 | 0.041275 |
| C | 3.740998 | 0.190848 | 0.574196 |
| C | 3.233259 | -1.114143 | 0.565242 |
| C | 1.975018 | -1.378874 | 0.027266 |
| H | 1.135831 | 1.812126 | -0.907364 |
| H | 3.361479 | 2.250531 | 0.040518 |
| H | 3.823437 | -1.935705 | 0.976770 |
| H | 1.572579 | -2.393326 | 0.003793 |
| C | -0.770776 | 0.286140 | -1.698191 |
| H | -1.573651 | -0.265998 | -2.231858 |
| H | -0.170982 | 0.785139 | -2.480212 |
| C | -1.464034 | 1.293486 | -0.863393 |
| C | -1.838550 | 1.116669 | 0.440410 |
| H | -1.868395 | 2.154490 | -1.416019 |
| C | -1.831059 | 1.788842 | 1.596383 |
| H | -1.150901 | 2.632414 | 1.746993 |

| | | | |
|---|---|---|---|
| H | -2.408179 | 1.450407 | 2.461717 |
| C | -2.557284 | -0.619826 | 0.675650 |
| O | -3.091510 | -1.079691 | -0.464447 |
| H | -3.268192 | -0.302483 | 1.454615 |
| O | -0.035387 | -0.677538 | -0.985628 |
| H | -0.993158 | -1.621327 | 0.381864 |
| H | 4.728225 | 0.394250 | 0.993074 |

**TS6**

E=-3417.352916
E_SP=-3419.60473
H=-3419.022206
G=-3419.121998
Imag. Freq. -1134.69

Cartesian coordinates

| | | | |
|---|---|---|---|
| Fe | -0.902964 | -0.701262 | 0.297347 |
| N | -1.628027 | 0.899601 | 1.271060 |
| N | 0.733081 | -0.663668 | 1.479667 |
| N | -2.587600 | -0.807284 | -0.812246 |
| N | -0.237453 | -2.400319 | -0.566090 |
| C | -2.812021 | 1.535859 | 1.018870 |
| C | 1.789193 | -1.540928 | 1.448993 |
| C | -1.001419 | 1.625629 | 2.257102 |
| C | 1.061862 | 0.292583 | 2.406868 |
| C | -3.626483 | 0.082595 | -0.802292 |
| C | 0.934895 | -3.055463 | -0.293846 |
| C | -2.872189 | -1.718784 | -1.793766 |
| C | -0.845071 | -3.104532 | -1.574642 |
| C | -2.958956 | 2.679428 | 1.887194 |
| C | 2.816012 | -1.119498 | 2.371997 |
| C | -1.831769 | 2.732400 | 2.660446 |
| C | 2.364770 | 0.022470 | 2.966180 |
| C | -4.606904 | -0.280514 | -1.799160 |
| C | 1.073683 | -4.208357 | -1.151809 |
| C | -4.138503 | -1.404504 | -2.415029 |
| C | -0.033060 | -4.238020 | -1.950590 |
| H | -3.819744 | 3.345595 | 1.892044 |
| H | 3.761485 | -1.637132 | 2.517796 |
| H | -1.565810 | 3.456455 | 3.428010 |
| H | 2.857218 | 0.644619 | 3.710687 |
| H | -5.528081 | 0.267380 | -1.987882 |
| H | 1.920259 | -4.892113 | -1.133591 |
| H | -4.591027 | -1.980016 | -3.220434 |
| H | -0.294012 | -4.953430 | -2.728174 |
| C | -3.733416 | 1.183161 | 0.036727 |
| C | 1.883778 | -2.662994 | 0.641642 |
| C | -2.069343 | -2.794520 | -2.152211 |
| C | 0.255005 | 1.360478 | 2.779765 |
| H | -4.618830 | 1.811243 | -0.067475 |
| H | 2.785922 | -3.268425 | 0.727901 |
| H | -2.427240 | -3.445578 | -2.950866 |
| H | 0.638128 | 2.040470 | 3.541133 |
| O | -0.162621 | 0.285568 | -0.930540 |
| S | -1.934305 | -2.119024 | 1.836046 |
| C | -1.781737 | -1.375306 | 3.485694 |
| H | -0.729177 | -1.220837 | 3.764028 |
| H | -2.316689 | -0.417136 | 3.551738 |
| H | -2.231884 | -2.084150 | 4.197513 |
| O | -0.541688 | 3.782780 | -0.116478 |
| H | -0.720830 | 1.160724 | -1.243592 |
| C | 3.727283 | 0.951868 | -0.298720 |
| C | 2.970173 | 0.112859 | -1.129870 |
| C | 3.542146 | -1.067747 | -1.615431 |
| C | 4.848767 | -1.432187 | -1.282970 |
| C | 5.594632 | -0.593340 | -0.444360 |
| C | 5.043081 | 0.590487 | 0.042032 |
| H | 1.928963 | 0.338704 | -1.359200 |
| H | 2.938900 | -1.717012 | -2.254765 |
| H | 6.618081 | -0.861647 | -0.169015 |
| H | 5.610778 | 1.255820 | 0.695877 |
| C | 1.948477 | 2.559904 | -0.069647 |
| H | 1.754688 | 3.381653 | 0.632359 |
| H | 1.222119 | 1.766567 | 0.151110 |
| C | 1.810336 | 3.018898 | -1.497998 |
| C | 0.692725 | 3.396318 | -2.139775 |
| H | 2.748678 | 3.024584 | -2.065463 |
| C | 0.681648 | 3.735172 | -3.603093 |
| H | 0.345305 | 4.773597 | -3.772281 |
| H | 1.672697 | 3.608169 | -4.061854 |
| H | -0.033607 | 3.082120 | -4.131596 |

| | | | |
|---|---|---|---|
| C | -0.695013 | 3.433095 | -1.483791 |
| O | -1.354733 | 2.268056 | -1.688134 |
| H | -1.270704 | 4.250225 | -1.986225 |
| O | 3.274900 | 2.118398 | 0.227005 |
| H | -1.394771 | 3.628586 | 0.314343 |
| H | 5.282775 | -2.357611 | -1.668476 |

**Int1_Q**

E=-3493.210914
E_SP=-3495.585909
H=-3494.983491
G=-3495.091943
Imag. Freq. 0

Cartesian coordinates

| | | | |
|---|---|---|---|
| Fe | 0.609950 | -0.927258 | 0.412557 |
| N | 0.967391 | 0.980608 | 1.266049 |
| N | 2.592904 | -0.983419 | -0.203319 |
| N | -1.345723 | -0.845774 | 1.019801 |
| N | 0.286113 | -2.825428 | -0.434961 |
| C | 0.061378 | 1.689704 | 1.994167 |
| C | 3.197043 | -2.052597 | -0.816099 |
| C | 2.171123 | 1.609451 | 1.348198 |
| C | 3.562033 | -0.058699 | 0.089745 |
| C | -1.924100 | 0.168634 | 1.727901 |
| C | 1.213677 | -3.579489 | -1.076946 |
| C | -2.295500 | -1.796660 | 0.781295 |
| C | -0.907646 | -3.473995 | -0.478086 |
| C | 0.717606 | 2.858796 | 2.557874 |
| C | 4.613499 | -1.771382 | -0.956416 |
| C | 2.025573 | 2.808876 | 2.158611 |
| C | 4.836471 | -0.541863 | -0.402681 |
| C | -3.324375 | -0.152233 | 1.951190 |
| C | 0.580405 | -4.798855 | -1.560005 |
| C | -3.552541 | -1.366831 | 1.369084 |
| C | -0.735786 | -4.732702 | -1.190468 |
| H | 0.232242 | 3.613789 | 3.175413 |
| H | 5.336239 | -2.437427 | -1.426399 |
| H | 2.826118 | 3.512445 | 2.383955 |
| H | 5.775406 | 0.005208 | -0.334324 |
| H | -4.036112 | 0.488984 | 2.468406 |
| H | 1.083358 | -5.592022 | -2.112783 |
| H | -4.489400 | -1.917656 | 1.309715 |
| H | -1.523380 | -5.462654 | -1.376133 |
| C | -1.280320 | 1.329534 | 2.178463 |
| C | 2.561679 | -3.225962 | -1.241349 |
| C | -2.099210 | -2.998752 | 0.087773 |
| C | 3.364163 | 1.142606 | 0.779282 |
| H | -1.896637 | 2.027271 | 2.749385 |
| H | 3.196163 | -3.955034 | -1.751137 |
| H | -2.978853 | -3.637512 | -0.020654 |
| H | 4.245285 | 1.773371 | 0.911563 |
| O | -0.009709 | -0.160020 | -1.220561 |
| S | 1.340897 | -1.909017 | 2.343184 |
| C | -0.082806 | -2.783691 | 3.072100 |
| H | -0.890894 | -2.095558 | 3.361802 |
| H | -0.492554 | -3.531412 | 2.376172 |
| H | 0.272874 | -3.307601 | 3.974650 |
| O | 1.005096 | 0.338102 | -2.123177 |
| O | 3.482534 | 1.462123 | -2.891228 |
| H | 3.382640 | 2.250468 | -2.323331 |
| H | 2.569683 | 1.282614 | -3.157775 |
| H | 1.826815 | -0.015745 | -1.729142 |
| C | -3.564992 | 1.510254 | -0.950512 |
| C | -3.087083 | 0.354820 | -1.585736 |
| C | -3.991572 | -0.659276 | -1.913115 |
| C | -5.353538 | -0.542556 | -1.627566 |
| C | -5.818918 | 0.615688 | -0.993033 |
| C | -4.934237 | 1.637437 | -0.657933 |
| H | -2.017644 | 0.199953 | -1.752323 |
| H | -3.604004 | -1.568609 | -2.379007 |
| H | -6.881176 | 0.723690 | -0.755781 |
| H | -5.273916 | 2.547939 | -0.159641 |
| C | -1.427713 | 2.580559 | -1.025471 |
| H | -1.411837 | 2.571461 | -2.133642 |
| H | -0.873666 | 1.677689 | -0.713671 |
| C | -0.829201 | 3.845637 | -0.495377 |
| C | 0.461222 | 4.238735 | -0.556657 |
| H | -1.561332 | 4.515580 | -0.030272 |
| C | 0.945077 | 5.557175 | -0.023450 |
| H | 1.452250 | 6.135548 | -0.812522 |

H 0.117262 6.154879 0.385190
H 1.689475 5.395018 0.772411
C 1.469106 3.365698 -1.177306
O 2.624138 3.728235 -1.375773
H 1.147485 2.345092 -1.472700
O -2.779168 2.550386 -0.576349
H -6.045981 -1.347669 -1.886007

**Int2_Q**

E=-3493.240696
E_SP=-3495.586564
H=-3494.982641
G=-3495.088007
Imag. Freq. 0

Cartesian coordinates
Fe -1.032511 0.567857 0.569507
N -1.554793 -1.220736 1.447212
N -2.974296 0.761001 -0.217669
N 0.926278 0.371431 1.334646
N -0.484076 2.355886 -0.314451
C -0.740360 -2.012839 2.201326
C -3.419162 1.785042 -0.993314
C -2.785210 -1.806660 1.383685
C -3.992286 -0.124080 -0.043105
C 1.379586 -0.667312 2.090641
C -1.297681 3.137472 -1.084303
C 1.962015 1.231946 1.124857
C 0.759479 2.919149 -0.295801
C -1.491836 -3.175797 2.646371
C -4.812882 1.544479 -1.340778
C -2.756705 -3.047978 2.141492
C -5.168017 0.361556 -0.751385
C 2.789810 -0.458997 2.385602
C -0.534460 4.276260 -1.570920
C 3.150752 0.717599 1.788498
C 0.737493 4.140452 -1.084825
H -1.090411 -3.980651 3.261180
H -5.427855 2.197482 -1.959356
H -3.600907 -3.726401 2.260016
H -6.131030 -0.146517 -0.792017
H 3.418075 -1.139469 2.959110
H -0.930468 5.067115 -2.207379
H 4.133066 1.186444 1.764831
H 1.592118 4.797854 -1.241913
C 0.609205 -1.765474 2.493692
C -2.644308 2.884567 -1.390515
C 1.884662 2.409337 0.368271
C -3.904237 -1.309077 0.699766
H 1.116724 -2.518360 3.101085
H -3.140665 3.626426 -2.020722
H 2.806603 2.986989 0.275142
H -4.810277 -1.917229 0.753836
O -0.492672 -0.223504 -1.057370
S -1.728532 1.580210 2.505287
C -0.656662 3.030946 2.776777
H 0.402886 2.739457 2.829645
H -0.770936 3.782106 1.980884
H -0.942075 3.489768 3.737263
O -1.324415 -1.345466 -1.429687
O -2.715581 -0.550494 -3.576966
H -1.868530 -2.240819 -3.533391
H -2.125207 0.158688 -3.870104
H -2.464274 -0.648825 -2.629166
C 4.485854 -0.854897 -0.790058
C 4.202229 0.383928 -1.386658
C 5.163782 1.399497 -1.362544
C 6.406698 1.202127 -0.758090
C 6.688902 -0.041043 -0.174569
C 5.742000 -1.061779 -0.191725
H 3.230659 0.576773 -1.837587
H 4.924013 2.363285 -1.819443
H 7.656837 -0.214812 0.303035
H 5.941226 -2.031012 0.269992
C 2.283436 -1.680367 -1.199250
H 2.295986 -1.486241 -2.287703
H 1.841606 -0.794978 -0.717502
C 1.477085 -2.897846 -0.856798
C 0.239428 -3.176744 -1.291481
H 1.946086 -3.557519 -0.119197
C -0.545448 -4.357903 -0.796128

H -0.867606 -4.999501 -1.631426
H 0.040710 -4.956513 -0.083667
H -1.459205 -4.013046 -0.285380
C -0.504572 -2.230424 -2.206551
O -1.358815 -2.927360 -3.050399
H 0.191695 -1.578303 -2.766678
O 3.616614 -1.887838 -0.733858
H 7.148604 2.003546 -0.740157

**Int5_Q**

E=-3417.373611
E_SP=-3419.625435
H=-3419.034625
G=-3419.136253
Imag. Freq. 0

Cartesian coordinates
Fe 1.169049 -0.399258 0.098665
N 1.432913 0.148143 -1.828528
N 0.480431 -2.192331 -0.513483
N 2.132224 1.287446 0.676892
N 1.232062 -1.078250 1.993951
C 1.902986 1.362864 -2.291214
C 0.072066 -3.230699 0.289424
C 1.004833 -0.542113 -2.942050
C 0.174201 -2.559270 -1.797458
C 2.493675 2.346289 -0.118327
C 0.723518 -2.263102 2.455603
C 2.423002 1.661124 1.966801
C 1.644959 -0.382753 3.100394
C 1.790991 1.421355 -3.726579
C -0.495517 -4.288009 -0.515180
C 1.236811 0.241842 -4.130987
C -0.432655 -3.870167 -1.813290
C 3.019968 3.421298 0.690740
C 0.827984 -2.326711 3.895276
C 2.980006 2.992482 1.986896
C 1.403502 -1.156166 4.296875
H 2.094425 2.273102 -4.332232
H -0.889108 -5.221940 -0.118206
H 0.987937 -0.082142 -5.139825
H -0.764913 -4.385968 -2.712195
H 3.376774 4.371650 0.297877
H 0.495880 -3.167150 4.501918
H 3.295593 3.516896 2.886916
H 1.646605 -0.826255 5.305265
C 2.390260 2.389760 -1.504729
C 0.184742 -3.275444 1.670118
C 2.209155 0.886925 3.098218
C 0.420558 -1.798992 -2.935693
H 2.707618 3.303772 -2.006900
H -0.183023 -4.167430 2.178623
H 2.491089 1.314173 4.061282
H 0.121654 -2.219607 -3.896414
O -0.317551 0.249189 0.251460
S 3.522423 -1.127549 -0.401478
C 3.448578 -2.663245 -1.344905
H 2.852854 -3.423840 -0.821595
H 2.971589 -2.461602 -2.318914
H 4.471216 -3.023725 -1.527181
O -1.207286 2.097118 -1.467156
H 0.419132 3.631408 -0.285754
C -4.779053 0.160948 0.023150
C -5.662832 -0.830976 0.482999
C -5.283188 -2.172317 0.458583
C -4.019326 -2.544721 -0.018688
C -3.148174 -1.554642 -0.475956
C -3.517245 -0.205904 -0.465549
H -6.640678 -0.523152 0.859466
H -5.979753 -2.933092 0.820954
H -2.155531 -1.817074 -0.839500
H -2.812841 0.539955 -0.822801
C -4.337514 2.514029 -0.217974
H -4.987314 3.404203 -0.241605
H -3.909472 2.384330 -1.224806
C -3.274936 2.661929 0.837129
C -1.989124 2.994922 0.650187
H -3.627787 2.454568 1.854192
C -1.011508 3.044618 1.791365
H -0.525941 4.030326 1.869055
H -1.506065 2.816478 2.745917

H -0.221137 2.291862 1.638640
C -1.437138 3.279399 -0.752441
O -0.300660 4.113755 -0.717804
H -2.173507 3.857513 -1.331631
O -5.233327 1.443315 0.081512
H -0.848234 1.408660 -0.854360
H -3.715425 -3.593837 -0.030186

**Int8_Q**

E=-3417.380386
E_SP=-3419.6301
H=-3419.039724
G=-3419.140605
Imag. Freq. 0

Cartesian coordinates
Fe -0.878979 -0.674287 0.190521
N -1.739574 0.860700 1.172647
N 0.659635 -0.636614 1.510410
N -2.560145 -0.921797 -0.889002
N -0.134907 -2.396432 -0.570629
C -2.936502 1.459304 0.860224
C 1.769582 -1.461391 1.517876
C -1.186152 1.608201 2.178095
C 0.887996 0.314924 2.481368
C -3.641095 -0.080452 -0.924396
C 1.070094 -2.979919 -0.279666
C -2.779184 -1.868155 -1.854106
C -0.688661 -3.144728 -1.582312
C -3.147840 2.612905 1.702247
C 2.708878 -1.015646 2.516884
C -2.060130 2.701870 2.526815
C 2.163375 0.082168 3.114933
C -4.579479 -0.514583 -1.932679
C 1.289821 -4.125894 -1.130450
C -4.043716 -1.629526 -2.510617
C 0.195637 -4.230005 -1.939793
H -4.025633 3.255497 1.662210
H 3.668884 -1.489363 2.709292
H -1.850225 3.438702 3.299799
H 2.579497 0.702929 3.905962
H -5.521087 -0.016256 -2.156041
H 2.175292 -4.757919 -1.095923
H -4.450503 -2.244622 -3.311285
H -0.013158 -4.968204 -2.712093
C -3.826849 1.030303 -0.111834
C 1.967520 -2.549388 0.690528
C -1.919037 -2.912604 -2.177175
C 0.033147 1.357879 2.798792
H -4.738510 1.611882 -0.252951
H 2.900506 -3.101724 0.797283
H -2.231548 -3.593676 -2.969847
H 0.347362 2.044323 3.585590
O -0.205271 0.287891 -0.946155
S -1.632607 -2.244834 1.982619
C -2.196087 -1.274653 3.395770
H -1.342830 -0.712380 3.811131
H -2.972376 -0.554342 3.102549
H -2.571768 -1.957954 4.171210
O -0.489986 3.786663 -0.047572
H -0.892872 1.738258 -1.551427
C 3.664226 0.874511 -0.200039
C 2.921493 0.171540 -1.160460
C 3.446904 -0.999068 -1.714281
C 4.696779 -1.486611 -1.326626
C 5.430209 -0.784137 -0.361235
C 4.922749 0.387597 0.196578
H 1.920800 0.494285 -1.441456
H 2.850451 -1.545218 -2.449087
H 6.408659 -1.151573 -0.040522
H 5.479616 0.946141 0.951630
C 1.968771 2.561741 0.077550
H 1.794210 3.343959 0.827350
H 1.187404 1.801941 0.214912
C 1.933209 3.121038 -1.321730
C 0.874400 3.590878 -1.999441
H 2.901246 3.108944 -1.835791
C 0.985527 4.039014 -3.431755
H 0.718169 5.104923 -3.542704
H 1.999496 3.891683 -3.830087
H 0.278173 3.470516 -4.059584

C -0.541778 3.657022 -1.448495
O -1.325868 2.573836 -1.854445
H -1.040977 4.544611 -1.887295
O 3.250802 2.017397 0.401778
H -1.393279 3.695865 0.290534
H 5.094853 -2.404527 -1.765062

**TS1_Q**

E=-3493.2081
E_SP=-3495.55266
H=-3494.956626
G=-3495.06053
Imag. Freq. -1227.7

Cartesian coordinates

Fe -1.033014 0.620358 0.524699
N -1.614069 -1.125598 1.440171
N -2.960203 0.827412 -0.307043
N 0.907188 0.403095 1.341767
N -0.429392 2.358446 -0.395922
C -0.830653 -1.915080 2.231194
C -3.373955 1.854496 -1.095221
C -2.854775 -1.691833 1.364470
C -4.004082 -0.023997 -0.122382
C 1.320030 -0.618261 2.143061
C -1.218114 3.150331 -1.182637
C 1.963939 1.238392 1.135032
C 0.824063 2.901532 -0.361275
C -1.613010 -3.052871 2.685864
C -4.772856 1.648889 -1.445228
C -2.864012 -2.914628 2.150974
C -5.162815 0.483173 -0.843186
C 2.724985 -0.426359 2.470005
C -0.425948 4.267957 -1.669463
C 3.124002 0.724574 1.846806
C 0.836392 4.112426 -1.164722
H -1.238947 -3.850353 3.326830
H -5.368173 2.312777 -2.071474
H -3.722651 -3.575141 2.265907
H -6.139510 0.001795 -0.879969
H 3.324249 -1.098950 3.082512
H -0.798176 5.060000 -2.318500
H 4.114673 1.176041 1.837789
H 1.705560 4.751901 -1.315150
C 0.515917 -1.689277 2.552476
C -2.567928 2.928846 -1.497682
C 1.925708 2.392179 0.340723
C -3.952324 -1.193629 0.647405
H 0.989859 -2.437249 3.192091
H -3.039528 3.677537 -2.138584
H 2.858572 2.952665 0.253022
H -4.870646 -1.782665 0.702134
O -0.487197 -0.242260 -1.099876
S -1.745499 1.702672 2.408764
C -0.708754 3.189731 2.609915
H 0.358586 2.927786 2.657236
H -0.857326 3.908676 1.790470
H -0.993457 3.672469 3.558705
O -1.301043 -1.390413 -1.390418
O -2.627339 -1.205240 -3.375413
H -2.050941 -2.215264 -3.390540
H -2.169941 -0.608902 -3.985948
H -2.016516 -1.092066 -2.228957
C 4.469485 -0.876437 -0.715370
C 4.172210 0.324864 -1.378803
C 5.134251 1.338494 -1.438691
C 6.390972 1.176122 -0.852633
C 6.685722 -0.029134 -0.199935
C 5.738080 -1.047198 -0.132107
H 3.191476 0.490796 -1.820427
H 4.884030 2.272783 -1.948195
H 7.664597 -0.175309 0.264425
H 5.948114 -1.987316 0.381990
C 2.268215 -1.731672 -1.057087
H 2.278731 -1.592658 -2.153070
H 1.819530 -0.826660 -0.620980
C 1.471286 -2.934457 -0.652892
C 0.265208 -3.287559 -1.124018
H 1.917766 -3.515713 0.160671
C -0.507729 -4.454726 -0.582319
H -0.757823 -5.163054 -1.387299

| | | | |
|---|---|---|---|
| H | 0.054391 | -4.977752 | 0.205004 |
| H | -1.462614 | -4.110151 | -0.153527 |
| C | -0.429941 | -2.494130 | -2.197881 |
| O | -1.289784 | -3.144481 | -2.948596 |
| H | 0.257420 | -1.792721 | -2.716298 |
| O | 3.603505 | -1.904947 | -0.581872 |
| H | 7.133823 | 1.975347 | -0.902026 |

**TS4_Q**

E=-3417.344345
E_SP=-3419.593956
H=-3419.010608
G=-3419.110549
Imag. Freq. -2049.02

Cartesian coordinates

| | | | |
|---|---|---|---|
| Fe | 1.232342 | -0.444538 | 0.078464 |
| N | 1.451175 | 0.125744 | -1.839250 |
| N | 0.290181 | -2.121284 | -0.542579 |
| N | 2.285055 | 1.173909 | 0.675009 |
| N | 1.243461 | -1.133747 | 1.970522 |
| C | 1.997047 | 1.312854 | -2.290883 |
| C | -0.215253 | -3.113482 | 0.260101 |
| C | 0.965111 | -0.517558 | -2.958271 |
| C | -0.055484 | -2.454153 | -1.821966 |
| C | 2.688991 | 2.223736 | -0.108571 |
| C | 0.614522 | -2.262101 | 2.423989 |
| C | 2.664976 | 1.488793 | 1.957820 |
| C | 1.768989 | -0.518450 | 3.074774 |
| C | 1.875741 | 1.399187 | -3.724717 |
| C | -0.901730 | -4.104301 | -0.541184 |
| C | 1.242599 | 0.263773 | -4.139689 |
| C | -0.799097 | -3.695664 | -1.839103 |
| C | 3.320547 | 3.242235 | 0.701324 |
| C | 0.758111 | -2.373326 | 3.857346 |
| C | 3.311636 | 2.780765 | 1.986184 |
| C | 1.482616 | -1.288283 | 4.262874 |
| H | 2.226815 | 2.238251 | -4.322404 |
| H | -1.390661 | -4.991579 | -0.142990 |
| H | 0.962153 | -0.026742 | -5.150504 |
| H | -1.187787 | -4.172831 | -2.736794 |
| H | 3.722357 | 4.177692 | 0.316127 |
| H | 0.348158 | -3.182020 | 4.459710 |
| H | 3.703483 | 3.257833 | 2.882657 |
| H | 1.793574 | -1.013963 | 5.269352 |
| C | 2.554779 | 2.296574 | -1.493561 |
| C | -0.068110 | -3.186553 | 1.637397 |
| C | 2.447969 | 0.696951 | 3.076158 |
| C | 0.268723 | -1.716835 | -2.958302 |
| H | 2.924958 | 3.194847 | -1.988797 |
| H | -0.521172 | -4.038456 | 2.146438 |
| H | 2.810218 | 1.069539 | 4.035314 |
| H | -0.069066 | -2.101014 | -3.921673 |
| O | -0.310803 | 0.339322 | 0.283782 |
| S | 3.421032 | -1.353618 | -0.424599 |
| C | 3.208676 | -2.822019 | -1.451915 |
| H | 2.550972 | -3.553883 | -0.962249 |
| H | 2.746410 | -2.533737 | -2.410890 |
| H | 4.195058 | -3.261948 | -1.657038 |
| O | -1.189157 | 2.134653 | -1.500747 |
| H | 0.618130 | 3.456107 | -0.454224 |
| C | -4.848300 | 0.480908 | 0.141264 |
| C | -5.743514 | -0.488931 | 0.623876 |
| C | -5.470521 | -1.844303 | 0.443492 |
| C | -4.302068 | -2.252315 | -0.213462 |
| C | -3.421346 | -1.283353 | -0.696686 |
| C | -3.686935 | 0.080187 | -0.534367 |
| H | -6.642551 | -0.153973 | 1.145539 |
| H | -6.173512 | -2.588379 | 0.827288 |
| H | -2.501738 | -1.580637 | -1.201652 |
| H | -2.975153 | 0.806007 | -0.919247 |
| C | -4.231407 | 2.807882 | 0.128089 |
| H | -4.782169 | 3.741992 | 0.326462 |
| H | -3.937653 | 2.814229 | -0.934323 |
| C | -3.049405 | 2.671162 | 1.044695 |
| C | -1.742233 | 2.851723 | 0.736609 |
| H | -3.306872 | 2.330123 | 2.054072 |
| C | -0.685796 | 2.434584 | 1.651710 |
| H | 0.249976 | 3.004425 | 1.626374 |
| H | -1.004997 | 2.169333 | 2.666622 |
| H | -0.371212 | 1.287613 | 1.094767 |

C -1.303835 3.261336 -0.675387
O -0.127971 4.037268 -0.663395
H -2.060242 3.921037 -1.125870
O -5.192277 1.780880 0.371535
H -0.830610 1.383839 -0.958388
H -4.077689 -3.313307 -0.343305

**TS6_Q**

E=-3417.354806
E_SP=-3419.605707
H=-3419.02266
G=-3419.122066
Imag. Freq. -570.92

Cartesian coordinates

| | | | |
|---|---|---|---|
| Fe | -0.906825 | -0.701705 | 0.298338 |
| N | -1.631422 | 0.933169 | 1.244975 |
| N | 0.720553 | -0.655741 | 1.474953 |
| N | -2.613482 | -0.827222 | -0.772417 |
| N | -0.229369 | -2.389424 | -0.568647 |
| C | -2.828385 | 1.556947 | 1.006653 |
| C | 1.771159 | -1.541025 | 1.464075 |
| C | -0.996043 | 1.670816 | 2.214074 |
| C | 1.055671 | 0.319561 | 2.382424 |
| C | -3.665554 | 0.051998 | -0.752967 |
| C | 0.936958 | -3.049700 | -0.290160 |
| C | -2.897083 | -1.747143 | -1.749592 |
| C | -0.836697 | -3.091067 | -1.576684 |
| C | -2.961243 | 2.717078 | 1.857050 |
| C | 2.795756 | -1.111028 | 2.383759 |
| C | -1.822667 | 2.785718 | 2.610528 |
| C | 2.352895 | 0.049159 | 2.950724 |
| C | -4.654122 | -0.334875 | -1.731442 |
| C | 1.081561 | -4.197772 | -1.155875 |
| C | -4.177511 | -1.454311 | -2.350143 |
| C | -0.020313 | -4.220784 | -1.960869 |
| H | -3.824461 | 3.380123 | 1.866895 |
| H | 3.737190 | -1.632163 | 2.543122 |
| H | -1.547738 | 3.521197 | 3.364033 |
| H | 2.849986 | 0.683682 | 3.681623 |
| H | -5.585550 | 0.198720 | -1.911151 |
| H | 1.927029 | -4.882968 | -1.136208 |
| H | -4.633249 | -2.038447 | -3.147550 |
| H | -0.278564 | -4.931777 | -2.743508 |
| C | -3.771107 | 1.168565 | 0.065009 |
| C | 1.869154 | -2.671059 | 0.666575 |
| C | -2.077252 | -2.801593 | -2.130596 |
| C | 0.260610 | 1.402063 | 2.736534 |
| H | -4.667321 | 1.781904 | -0.033436 |
| H | 2.765806 | -3.282142 | 0.768863 |
| H | -2.437046 | -3.455059 | -2.926424 |
| H | 0.653409 | 2.091516 | 3.484352 |
| O | -0.175966 | 0.262008 | -0.955855 |
| S | -2.119975 | -1.919766 | 1.844801 |
| C | -1.436354 | -1.585162 | 3.491522 |
| H | -0.385066 | -1.897342 | 3.560616 |
| H | -1.505887 | -0.513568 | 3.732128 |
| H | -2.039461 | -2.147648 | 4.219513 |
| O | -0.501584 | 3.834740 | -0.148584 |
| H | -0.717250 | 1.128969 | -1.240022 |
| C | 3.738048 | 0.929948 | -0.297933 |
| C | 2.974164 | 0.085463 | -1.117290 |
| C | 3.541737 | -1.099396 | -1.597665 |
| C | 4.849535 | -1.463342 | -1.270057 |
| C | 5.601466 | -0.619828 | -0.441519 |
| C | 5.054824 | 0.568709 | 0.038699 |
| H | 1.933170 | 0.312557 | -1.345761 |
| H | 2.933775 | -1.752222 | -2.228641 |
| H | 6.625933 | -0.887953 | -0.169774 |
| H | 5.627459 | 1.238505 | 0.683622 |
| C | 1.970962 | 2.552562 | -0.084741 |
| H | 1.787211 | 3.391855 | 0.598659 |
| H | 1.236785 | 1.772146 | 0.155081 |
| C | 1.833229 | 2.979900 | -1.523084 |
| C | 0.718392 | 3.358826 | -2.165692 |
| H | 2.766564 | 2.953111 | -2.097798 |
| C | 0.698340 | 3.663728 | -3.636299 |
| H | 0.384301 | 4.705535 | -3.826608 |
| H | 1.681512 | 3.503432 | -4.101324 |
| H | -0.036587 | 3.015267 | -4.143198 |
| C | -0.663068 | 3.434920 | -1.497528 |

O -1.360684 2.289608 -1.661747
H -1.231693 4.239016 -2.033137
O 3.292306 2.103166 0.219079
H -1.341986 3.667660 0.303565
H 5.279871 -2.392246 -1.651278

**Int10**

E=-3226.485017
E_SP=-3228.516489
H=-3227.993971
G=-3228.089308
Imag. Freq. 0

Cartesian coordinates
Fe 1.050657 0.236712 0.218929
N 2.269196 -1.338276 -0.101523
N -0.142669 -0.929489 1.333100
N 2.297225 1.411508 -0.835075
N -0.136218 1.804806 0.579834
C 3.332863 -1.379710 -0.965714
C -1.377829 -0.592470 1.828201
C 2.201734 -2.567894 0.504955
C 0.114367 -2.213829 1.748727
C 3.354185 0.996296 -1.601640
C -1.361115 1.779310 1.197877
C 2.215319 2.771482 -0.974755
C 0.120179 3.115885 0.264608
C 3.942961 -2.689381 -0.926355
C -1.927880 -1.704159 2.567298
C 3.254666 -3.418491 0.002502
C -0.992667 -2.701462 2.537257
C 3.966093 2.136164 -2.249217
C -1.898987 3.117885 1.274070
C 3.271482 3.239505 -1.845133
C -0.973947 3.948960 0.711022
H 4.804324 -2.987070 -1.522011
H -2.901560 -1.699047 3.053863
H 3.422311 -4.446177 0.320091
H -1.041025 -3.692134 2.985915
H 4.825854 2.079437 -2.914706
H -2.864916 3.367258 1.707982
H 3.432149 4.281678 -2.115749
H -1.017899 5.029200 0.583444
C 3.823755 -0.306878 -1.700724
C -1.974079 0.658429 1.737498
C 1.227558 3.582977 -0.431467
C 1.224712 -2.972741 1.403153
H 4.681626 -0.488107 -2.349238
H -2.963867 0.783453 2.176502
H 1.292131 4.655376 -0.618835
H 1.292008 -3.983975 1.805396
O 0.148653 -0.158061 -1.303830
S 2.315039 0.738965 2.033522
C 1.178810 0.991717 3.429313
H 1.788799 1.271243 4.301080
H 0.465355 1.799902 3.214207
H 0.621688 0.072681 3.659515
C -4.165593 -0.545607 -1.120131
C -5.286997 0.169946 -0.669198
C -5.266201 1.564059 -0.658771
C -4.132348 2.259843 -1.096815
C -3.016266 1.540454 -1.525233
C -3.020337 0.142806 -1.540653
H -6.165544 -0.389712 -0.341717
H -6.146364 2.111110 -0.311016
H -2.104740 2.060253 -1.826706
H -2.103769 -0.370955 -1.822267
C -3.310873 -2.690070 -1.776381
H -3.062848 -2.267871 -2.766842
H -3.803715 -3.663499 -1.953600
C -2.072518 -2.898815 -0.955856
C -0.844676 -3.177974 -1.513306
H -2.181826 -2.881257 0.132917
C 0.258258 -3.389948 -2.017730
H 1.234285 -3.575581 -2.427193
H 0.153420 -1.122992 -1.424709
O -4.286255 -1.905688 -1.114952
H -4.113275 3.351265 -1.083348

**Int10_Q**

E=-3226.484842
E_SP=-3228.516372
H=-3227.993872
G=-3228.089873
Imag. Freq. 0

Cartesian coordinates

| | | | |
|---|---|---|---|
| Fe | 1.055438 | 0.231039 | 0.217294 |
| N | 2.258409 | -1.357199 | -0.103370 |
| N | -0.149500 | -0.923937 | 1.330479 |
| N | 2.314354 | 1.393425 | -0.835704 |
| N | -0.115171 | 1.810320 | 0.578531 |
| C | 3.320669 | -1.409150 | -0.968510 |
| C | -1.381103 | -0.574559 | 1.826045 |
| C | 2.178301 | -2.586236 | 0.502245 |
| C | 0.094995 | -2.210708 | 1.746451 |
| C | 3.366985 | 0.967330 | -1.602267 |
| C | -1.340827 | 1.796845 | 1.195652 |
| C | 2.246547 | 2.754100 | -0.974950 |
| C | 0.154154 | 3.118879 | 0.263132 |
| C | 3.916688 | -2.725490 | -0.930937 |
| C | -1.941782 | -1.680501 | 2.565679 |
| C | 3.221510 | -3.447895 | -0.002036 |
| C | -1.016485 | -2.686992 | 2.535634 |
| C | 3.990933 | 2.100996 | -2.249160 |
| C | -1.865692 | 3.140546 | 1.271096 |
| C | 3.307777 | 3.211426 | -1.844783 |
| C | -0.932229 | 3.962496 | 0.708509 |
| H | 4.774240 | -3.031972 | -1.527641 |
| H | -2.915191 | -1.665634 | 3.052600 |
| H | 3.378407 | -4.477594 | 0.314538 |
| H | -1.074523 | -3.677058 | 2.984502 |
| H | 4.850157 | 2.035568 | -2.914545 |
| H | -2.829396 | 3.399412 | 1.704401 |
| H | 3.479396 | 4.251962 | -2.114996 |
| H | -0.965614 | 5.043092 | 0.580716 |
| C | 3.822613 | -0.340749 | -1.702509 |
| C | -1.964905 | 0.682123 | 1.735296 |
| C | 1.266550 | 3.575252 | -0.431955 |
| C | 1.197487 | -2.980956 | 1.400873 |
| H | 4.678237 | -0.530385 | -2.351597 |
| H | -2.953498 | 0.816959 | 2.174083 |
| H | 1.341745 | 4.647003 | -0.619036 |
| H | 1.254315 | -3.992837 | 1.803086 |
| O | 0.154241 | -0.154097 | -1.308622 |
| S | 2.326586 | 0.719485 | 2.030801 |
| C | 1.193432 | 0.978445 | 3.427950 |
| H | 1.806063 | 1.253581 | 4.299261 |
| H | 0.484821 | 1.791191 | 3.214024 |
| H | 0.630802 | 0.062738 | 3.658102 |
| C | -4.178906 | -0.514731 | -1.115169 |
| C | -5.285445 | 0.224598 | -0.665599 |
| C | -5.237549 | 1.618004 | -0.661140 |
| C | -4.091056 | 2.289797 | -1.103786 |
| C | -2.989747 | 1.547200 | -1.530587 |
| C | -3.021118 | 0.149774 | -1.540081 |
| H | -6.174198 | -0.316483 | -0.334468 |
| H | -6.106352 | 2.183536 | -0.314376 |
| H | -2.068874 | 2.047796 | -1.835868 |
| H | -2.115869 | -0.382873 | -1.822923 |
| C | -3.365182 | -2.678301 | -1.760549 |
| H | -3.113810 | -2.269985 | -2.755931 |
| H | -3.874085 | -3.645470 | -1.925639 |
| C | -2.127631 | -2.896733 | -0.941412 |
| C | -0.900292 | -3.176507 | -1.499580 |
| H | -2.235714 | -2.880143 | 0.147539 |
| C | 0.201243 | -3.393211 | -2.005189 |
| H | 1.176109 | -3.582288 | -2.415902 |
| H | 0.139512 | -1.119686 | -1.423283 |
| O | -4.326604 | -1.871987 | -1.104351 |
| H | -4.050803 | 3.380662 | -1.095066 |

**Int11**

E=-537.105015
E_SP=-537.714449
H=-537.526936
G=-537.575148
Imag. Freq. 0

Cartesian coordinates

| | | | |
|---|---|---|---|
| C | 0.968317 | -0.511890 | 0.254218 |

| | | | |
|---|---|---|---|
| C | 1.975180 | -1.190426 | -0.446534 |
| C | 3.154473 | -0.529568 | -0.792475 |
| C | 3.336387 | 0.815636 | -0.453457 |
| C | 2.329197 | 1.487624 | 0.244440 |
| C | 1.148213 | 0.834038 | 0.609981 |
| H | 1.811208 | -2.237226 | -0.709530 |
| H | 3.933553 | -1.068739 | -1.337021 |
| H | 2.462128 | 2.536739 | 0.520395 |
| H | 0.371496 | 1.351602 | 1.169831 |
| C | -1.443583 | -0.696182 | 0.581533 |
| H | -2.086913 | -1.579157 | 0.731803 |
| C | -1.770497 | -0.048679 | -0.770528 |
| C | -3.171649 | 0.340370 | -0.892887 |
| H | -1.123636 | 0.831837 | -0.912410 |
| C | -4.338610 | 0.650408 | -0.992373 |
| H | -5.371765 | 0.931495 | -1.081706 |
| O | -0.136949 | -1.253603 | 0.591751 |
| H | 4.257119 | 1.334419 | -0.728939 |
| H | -1.507617 | -0.770187 | -1.560819 |
| O | -1.648052 | 0.248395 | 1.585036 |
| H | -1.590134 | -0.197113 | 2.442591 |

**Int12**

E=-3226.497534
E_SP=-3228.521208
H=-3227.996429
G=-3228.089754
Imag. Freq. 0

Cartesian coordinates

| | | | |
|---|---|---|---|
| Fe | -0.685114 | 0.110908 | -0.449527 |
| N | -1.888836 | -1.458532 | -0.089499 |
| N | 0.773868 | -1.140227 | -1.006024 |
| N | -2.186482 | 1.372362 | 0.008753 |
| N | 0.497341 | 1.680183 | -0.856025 |
| C | -3.230916 | -1.411406 | 0.206469 |
| C | 1.980737 | -0.794804 | -1.561917 |
| C | -1.538910 | -2.786233 | -0.113616 |
| C | 0.780656 | -2.507402 | -0.870457 |
| C | -3.483678 | 1.030372 | 0.302679 |
| C | 1.730705 | 1.642073 | -1.455647 |
| C | -2.093367 | 2.731459 | 0.159928 |
| C | 0.225567 | 3.000400 | -0.600426 |
| C | -3.740190 | -2.749952 | 0.379150 |
| C | 2.770540 | -1.980576 | -1.793983 |
| C | -2.687825 | -3.602517 | 0.195422 |
| C | 2.034704 | -3.040463 | -1.348774 |
| C | -4.239412 | 2.214922 | 0.632106 |
| C | 2.255762 | 2.981067 | -1.586814 |
| C | -3.372563 | 3.268781 | 0.561676 |
| C | 1.332916 | 3.822571 | -1.036495 |
| H | -4.774318 | -2.988819 | 0.621217 |
| H | 3.771825 | -1.977113 | -2.219690 |
| H | -2.675195 | -4.689755 | 0.247262 |
| H | 2.296231 | -4.097077 | -1.343512 |
| H | -5.294065 | 2.218558 | 0.901508 |
| H | 3.221879 | 3.222661 | -2.025906 |
| H | -3.566231 | 4.323340 | 0.750362 |
| H | 1.370279 | 4.906469 | -0.941544 |
| C | -3.988781 | -0.260425 | 0.364237 |
| C | 2.419302 | 0.495516 | -1.818552 |
| C | -0.962430 | 3.500054 | -0.087953 |
| C | -0.281162 | -3.284784 | -0.432557 |
| H | -5.044486 | -0.380002 | 0.609645 |
| H | 3.406049 | 0.615374 | -2.264456 |
| H | -1.036204 | 4.576598 | 0.069157 |
| H | -0.138233 | -4.364920 | -0.392998 |
| O | -0.077263 | 0.142434 | 1.289621 |
| S | -1.509716 | 0.189035 | -2.561411 |
| C | -1.239074 | -1.435349 | -3.328649 |
| H | -1.681845 | -1.385411 | -4.334983 |
| H | -0.169538 | -1.669965 | -3.415708 |
| H | -1.741718 | -2.228474 | -2.757242 |
| C | 3.038865 | -0.571674 | 1.700567 |
| C | 4.196354 | -1.168527 | 1.174807 |
| C | 5.194492 | -0.382074 | 0.603386 |
| C | 5.060755 | 1.010895 | 0.558303 |
| C | 3.905087 | 1.598086 | 1.077848 |
| C | 2.887606 | 0.820999 | 1.637621 |
| H | 4.285977 | -2.254963 | 1.223630 |
| H | 6.088302 | -0.861714 | 0.195844 |

H 3.770540 2.680590 1.024077
H 1.961626 1.287929 1.965051
C 1.094116 -0.942012 3.082728
H 1.371245 0.012371 3.556958
H 0.951981 -1.692237 3.876323
C -0.228689 -0.790584 2.313355
C -1.314102 -0.390068 3.249658
H -0.466121 -1.791370 1.909628
C -2.118950 0.634405 3.429905
H -2.849541 0.672031 4.247025
H -2.092083 1.497527 2.748089
O 2.134411 -1.425131 2.253002
H 5.844948 1.628782 0.115098

**Int13**

E=-3226.044992
E_SP=-3228.069268
H=-3227.557203
G=-3227.650216
Imag. Freq. 0

Cartesian coordinates
Fe -0.737755 0.220197 -0.419690
N -2.242470 -1.001647 0.000275
N 0.375333 -1.439098 -1.171782
N -1.857392 1.882787 0.309540
N 0.737622 1.457109 -0.926953
C -3.451538 -0.632695 0.541046
C 1.621766 -1.374843 -1.704613
C -2.272798 -2.364287 -0.189024
C -0.009754 -2.740979 -1.168176
C -3.107645 1.823036 0.832511
C 1.913487 1.098595 -1.539404
C -1.450305 3.175078 0.347648
C 0.764980 2.817079 -0.742898
C -4.282747 -1.806539 0.690441
C 2.069823 -2.713349 -2.059270
C -3.554708 -2.874881 0.244283
C 1.052288 -3.566001 -1.724989
C -3.530077 3.156162 1.240700
C 2.712271 2.283623 -1.766593
C -2.495638 3.999475 0.939376
C 2.005530 3.344069 -1.274021
H -5.293193 -1.799626 1.096992
H 3.034220 -2.956402 -2.504177
H -3.845817 -3.923805 0.206267
H 1.014888 -4.648766 -1.841325
H -4.490641 3.402175 1.692578
H 3.698044 2.282608 -2.228481
H -2.437774 5.076565 1.094738
H 2.286857 4.396337 -1.257554
C -3.856177 0.648257 0.931170
C 2.335322 -0.187916 -1.885111
C -0.221017 3.614828 -0.150266
C -1.255724 -3.174361 -0.707293
H -4.858504 0.727988 1.357306
H 3.328347 -0.265752 -2.331686
H -0.004894 4.683330 -0.085149
H -1.469232 -4.243738 -0.762272
O 0.062382 -0.011506 1.348405
S -1.687368 0.645777 -2.612349
C -2.206021 -0.973200 -3.274947
H -2.599267 -0.834940 -4.295583
H -1.362316 -1.679220 -3.322051
H -2.996403 -1.431377 -2.659433
C 3.296150 -0.780032 1.553838
C 4.524978 -1.241025 1.052226
C 5.488467 -0.332477 0.614511
C 5.246422 1.044642 0.683252
C 4.016360 1.494232 1.171939
C 3.034338 0.597466 1.597450
H 4.698774 -2.318528 1.014924
H 6.439994 -0.704425 0.224440
H 3.795506 2.563549 1.191666
H 2.040015 0.941242 1.878935
C 1.349988 -1.418299 2.840570
H 1.597126 -0.544713 3.466422
H 1.229345 -2.294796 3.497644
C 0.029356 -1.168951 2.083545
C -1.075387 -1.171664 3.079394
H -0.111495 -2.072118 1.452081

C -1.978559 -1.137857 3.889796
H -2.791964 -1.102437 4.589590
O 2.425645 -1.736642 1.967941
H 6.001464 1.757283 0.342764

**Int9**

E=-3226.485276
E_SP=-3228.514773
H=-3227.990341
G=-3228.084978
Imag. Freq. 0

Cartesian coordinates
Fe 1.007851 0.198301 0.090081
N 1.527683 -1.756330 0.279598
N -0.630631 -0.147590 1.213265
N 2.798287 0.558174 -0.786342
N 0.679638 2.178703 0.234183
C 2.612663 -2.379721 -0.303385
C -1.571778 0.779286 1.595773
C 0.785772 -2.753398 0.869699
C -1.102282 -1.362798 1.630117
C 3.693113 -0.374666 -1.239045
C -0.409292 2.793141 0.790435
C 3.277387 1.778696 -1.191715
C 1.457391 3.178692 -0.291196
C 2.562542 -3.799812 -0.045544
C -2.677696 0.123527 2.254909
C 1.436301 -4.030037 0.689557
C -2.384293 -1.208713 2.279184
C 4.773408 0.271961 -1.948867
C -0.321454 4.223520 0.616900
C 4.517752 1.612567 -1.914234
C 0.842134 4.464599 -0.057523
H 3.306046 -4.515430 -0.392377
H -3.556901 0.636194 2.638857
H 1.053046 -4.975598 1.069045
H -2.971026 -2.027906 2.690536
H 5.615130 -0.249593 -2.401207
H -1.064847 4.935759 0.970441
H 5.102461 2.428804 -2.334958
H 1.261511 5.417805 -0.374935
C 3.609474 -1.747795 -1.026595
C -1.468449 2.148171 1.421300
C 2.667783 3.002416 -0.952393
C -0.434181 -2.576891 1.502398
H 4.401836 -2.370817 -1.443488
H -2.284703 2.766224 1.793762
H 3.171798 3.896189 -1.323008
H -0.922127 -3.464115 1.907836
O 0.256830 0.004156 -1.336279
S 2.453764 0.161202 2.154432
C 1.460090 -0.465173 3.524463
H 2.058638 -0.441575 4.446890
H 0.534996 0.115549 3.645346
H 1.181805 -1.512510 3.317170
C -4.130988 -0.046797 -0.907223
C -4.973763 0.939375 -0.366758
C -4.622505 2.285350 -0.454360
C -3.426786 2.666294 -1.075630
C -2.582342 1.680565 -1.588598
C -2.922160 0.327310 -1.511376
H -5.905671 0.623121 0.106308
H -5.290945 3.043052 -0.036932
H -1.619051 1.947340 -2.026701
H -2.214352 -0.412097 -1.877175
C -4.006670 -2.344149 -1.623914
H -3.817180 -1.955935 -2.638103
H -4.791484 -3.114072 -1.691967
C -2.752021 -2.949996 -1.044827
C -1.601674 -2.983804 -1.673200
H -2.839290 -3.375269 -0.038132
C -0.455797 -2.953527 -2.302947
H -0.143197 -3.759925 -2.975380
H 0.212742 -2.096964 -2.149979
O -4.573645 -1.330723 -0.798416
H -3.144000 3.719398 -1.134669

**Int9_Q**

E=-3226.485008

E_SP=-3228.514448
H=-3227.990003
G=-3228.08511
Imag. Freq. 0

Cartesian coordinates

| | | | |
|---|---|---|---|
| Fe | 1.008866 | 0.196049 | 0.084189 |
| N | 1.526815 | -1.752041 | 0.282297 |
| N | -0.631869 | -0.144240 | 1.209047 |
| N | 2.799313 | 0.553727 | -0.790019 |
| N | 0.686144 | 2.180595 | 0.233566 |
| C | 2.614897 | -2.378514 | -0.293491 |
| C | -1.572611 | 0.784975 | 1.587476 |
| C | 0.786614 | -2.747025 | 0.881118 |
| C | -1.105698 | -1.356980 | 1.628848 |
| C | 3.694915 | -0.379519 | -1.239670 |
| C | -0.405179 | 2.796338 | 0.782870 |
| C | 3.281688 | 1.774196 | -1.193113 |
| C | 1.464634 | 3.178341 | -0.293741 |
| C | 2.566997 | -3.796021 | -0.024273 |
| C | -2.681131 | 0.132198 | 2.245351 |
| C | 1.440625 | -4.022770 | 0.711719 |
| C | -2.389275 | -1.200299 | 2.274031 |
| C | 4.777330 | 0.265450 | -1.947541 |
| C | -0.317466 | 4.226203 | 0.604185 |
| C | 4.523332 | 1.606337 | -1.913043 |
| C | 0.848495 | 4.465147 | -0.066931 |
| H | 3.312225 | -4.512784 | -0.364913 |
| H | -3.560553 | 0.646946 | 2.625932 |
| H | 1.059071 | -4.966159 | 1.098212 |
| H | -2.977639 | -2.017807 | 2.686384 |
| H | 5.619616 | -0.257158 | -2.397577 |
| H | -1.062670 | 4.939344 | 0.952060 |
| H | 5.110338 | 2.421977 | -2.331697 |
| H | 1.268443 | 5.417148 | -0.387171 |
| C | 3.610894 | -1.751721 | -1.021656 |
| C | -1.466896 | 2.153317 | 1.411769 |
| C | 2.675644 | 2.999272 | -0.953258 |
| C | -0.435203 | -2.570727 | 1.509053 |
| H | 4.403545 | -2.377208 | -1.434213 |
| H | -2.283699 | 2.772716 | 1.780732 |
| H | 3.182224 | 3.891760 | -1.323429 |
| H | -0.921561 | -3.456937 | 1.918548 |
| O | 0.242719 | 0.037356 | -1.339307 |
| S | 2.452317 | 0.162748 | 2.157926 |
| C | 1.450203 | -0.446930 | 3.529569 |
| H | 2.046871 | -0.421305 | 4.453233 |
| H | 0.529001 | 0.141001 | 3.644938 |
| H | 1.165554 | -1.493917 | 3.329337 |
| C | -4.139565 | -0.068435 | -0.908241 |
| C | -4.994469 | 0.903565 | -0.361023 |
| C | -4.659412 | 2.254281 | -0.438325 |
| C | -3.468002 | 2.654191 | -1.055967 |
| C | -2.611651 | 1.682485 | -1.576057 |
| C | -2.935376 | 0.324695 | -1.509343 |
| H | -5.922524 | 0.572545 | 0.109494 |
| H | -5.337069 | 3.000723 | -0.015531 |
| H | -1.651511 | 1.963960 | -2.011885 |
| H | -2.218456 | -0.403248 | -1.880254 |
| C | -3.983635 | -2.360357 | -1.636635 |
| H | -3.796499 | -1.965118 | -2.648530 |
| H | -4.757873 | -3.140379 | -1.710368 |
| C | -2.722670 | -2.951854 | -1.056312 |
| C | -1.570316 | -2.968317 | -1.681716 |
| H | -2.807082 | -3.381894 | -0.051452 |
| C | -0.423688 | -2.920011 | -2.308863 |
| H | -0.098080 | -3.719530 | -2.983305 |
| H | 0.231891 | -2.054063 | -2.152123 |
| O | -4.566495 | -1.358392 | -0.808210 |
| H | -3.197909 | 3.711062 | -1.107001 |

**TS2_Q**

E=-3226.481971
E_SP=-3228.514434
H=-3227.992809
G=-3228.087879
Imag. Freq. -49.93

Cartesian coordinates

| | | | |
|---|---|---|---|
| Fe | 1.065142 | -0.155883 | 0.216963 |
| N | 2.553950 | -1.085514 | -0.749711 |

| | | | |
|---|---|---|---|
| N | -0.002181 | -1.867044 | 0.231206 |
| N | 2.134619 | 1.546956 | 0.237858 |
| N | -0.388956 | 0.751612 | 1.288542 |
| C | 3.781744 | -0.562682 | -1.064852 |
| C | -1.173713 | -2.101976 | 0.907573 |
| C | 2.514142 | -2.347906 | -1.288448 |
| C | 0.314771 | -3.028839 | -0.425082 |
| C | 3.418587 | 1.713244 | -0.213194 |
| C | -1.524881 | 0.164357 | 1.790865 |
| C | 1.722940 | 2.768410 | 0.705409 |
| C | -0.445572 | 2.077745 | 1.632443 |
| C | 4.554893 | -1.533977 | -1.804432 |
| C | -1.606984 | -3.460025 | 0.677042 |
| C | 3.762393 | -2.635494 | -1.956743 |
| C | -0.693884 | -4.030627 | -0.162992 |
| C | 3.831189 | 3.087764 | -0.033419 |
| C | -2.328793 | 1.154570 | 2.468191 |
| C | 2.773553 | 3.745638 | 0.523076 |
| C | -1.662859 | 2.342988 | 2.364666 |
| H | 5.567088 | -1.367882 | -2.169186 |
| H | -2.510148 | -3.896466 | 1.098384 |
| H | 3.987652 | -3.570724 | -2.466509 |
| H | -0.679222 | -5.041125 | -0.567770 |
| H | 4.807617 | 3.480030 | -0.312511 |
| H | -3.285714 | 0.948278 | 2.943608 |
| H | 2.695539 | 4.793901 | 0.806267 |
| H | -1.952872 | 3.321527 | 2.743201 |
| C | 4.208967 | 0.730404 | -0.791694 |
| C | -1.884922 | -1.167567 | 1.646539 |
| C | 0.517339 | 3.034306 | 1.337968 |
| C | 1.468373 | -3.251878 | -1.166870 |
| H | 5.216455 | 1.007735 | -1.103743 |
| H | -2.825682 | -1.487594 | 2.094064 |
| H | 0.329998 | 4.057732 | 1.663568 |
| H | 1.584140 | -4.226939 | -1.641292 |
| O | 0.301211 | 0.313475 | -1.345481 |
| S | 1.939569 | -0.712545 | 2.236977 |
| C | 2.643669 | -2.381340 | 2.089232 |
| H | 3.030611 | -2.662564 | 3.079943 |
| H | 1.873092 | -3.105376 | 1.787806 |
| H | 3.464410 | -2.405888 | 1.358828 |
| C | -3.521108 | -0.160939 | -1.293815 |
| C | -3.667244 | -1.552894 | -1.220953 |
| C | -4.688078 | -2.105352 | -0.451737 |
| C | -5.573641 | -1.280117 | 0.252326 |
| C | -5.426595 | 0.105827 | 0.168909 |
| C | -4.407131 | 0.675563 | -0.602256 |
| H | -2.958788 | -2.181695 | -1.761822 |
| H | -4.790185 | -3.191819 | -0.398480 |
| H | -6.109197 | 0.763133 | 0.712987 |
| H | -4.300808 | 1.759167 | -0.636811 |
| C | -2.429543 | 1.651197 | -2.454385 |
| H | -3.434376 | 1.964533 | -2.799990 |
| H | -1.763141 | 1.678454 | -3.330636 |
| C | -1.909646 | 2.558371 | -1.378604 |
| C | -1.464210 | 3.835506 | -1.631665 |
| H | -1.876119 | 2.176244 | -0.357682 |
| C | -1.070177 | 4.978075 | -1.866149 |
| H | -0.711343 | 5.970342 | -2.067765 |
| H | -0.621955 | 0.007622 | -1.414933 |
| O | -2.473075 | 0.289275 | -2.047741 |
| H | -6.372444 | -1.715140 | 0.856609 |

**TS7**

E=-3226.456336

E_SP=-3228.486879

H=-3227.969759

G=-3228.062594

Imag. Freq. -2001.78

Cartesian coordinates

| | | | |
|---|---|---|---|
| Fe | 1.101121 | 0.150248 | 0.138616 |
| N | 0.881911 | -1.767693 | 0.628499 |
| N | -0.599364 | 0.593647 | 1.116295 |
| N | 2.893866 | -0.260226 | -0.696429 |
| N | 1.403084 | 2.115867 | -0.217529 |
| C | 1.797636 | -2.780952 | 0.438747 |
| C | -1.100251 | 1.845025 | 1.377697 |
| C | -0.224030 | -2.348253 | 1.193425 |
| C | -1.528209 | -0.298561 | 1.577749 |
| C | 3.529602 | -1.471292 | -0.719296 |

| | | | |
|---|---|---|---|
| C | 0.620880 | 3.156985 | 0.208503 |
| C | 3.690578 | 0.609517 | -1.393661 |
| C | 2.395824 | 2.665252 | -0.988794 |
| C | 1.250736 | -4.030429 | 0.906906 |
| C | -2.379540 | 1.735310 | 2.043823 |
| C | -0.009849 | -3.764662 | 1.361361 |
| C | -2.656548 | 0.405497 | 2.147710 |
| C | 4.772174 | -1.368676 | -1.449662 |
| C | 1.134029 | 4.405924 | -0.307050 |
| C | 4.865618 | -0.076507 | -1.881261 |
| C | 2.228986 | 4.099033 | -1.062331 |
| H | 1.774849 | -4.983848 | 0.872930 |
| H | -2.989000 | 2.582683 | 2.349397 |
| H | -0.738403 | -4.452754 | 1.786568 |
| H | -3.536482 | -0.076573 | 2.568247 |
| H | 5.465717 | -2.192364 | -1.609460 |
| H | 0.693173 | 5.381929 | -0.111858 |
| H | 5.656335 | 0.390962 | -2.465433 |
| H | 2.884630 | 4.769774 | -1.614870 |
| C | 3.038047 | -2.648177 | -0.164151 |
| C | -0.525381 | 3.043116 | 0.986459 |
| C | 3.452484 | 1.968690 | -1.559481 |
| C | -1.376469 | -1.678555 | 1.588203 |
| H | 3.654817 | -3.543755 | -0.249534 |
| H | -1.046261 | 3.962947 | 1.254687 |
| H | 4.176862 | 2.539824 | -2.141765 |
| H | -2.188455 | -2.271866 | 2.008547 |
| O | 0.286157 | 0.019059 | -1.365434 |
| S | 2.250378 | 0.560980 | 2.158842 |
| C | 1.426074 | -0.425867 | 3.444579 |
| H | 1.929014 | -0.198958 | 4.397563 |
| H | 0.363093 | -0.160606 | 3.541107 |
| H | 1.509230 | -1.504553 | 3.247675 |
| C | -4.290170 | -0.206578 | -0.900921 |
| C | -5.279130 | 0.654175 | -0.397438 |
| C | -5.054347 | 2.029536 | -0.357306 |
| C | -3.844954 | 2.564431 | -0.816217 |
| C | -2.860030 | 1.702523 | -1.301301 |
| C | -3.070205 | 0.322311 | -1.346550 |
| H | -6.220716 | 0.220760 | -0.054054 |
| H | -5.833885 | 2.689234 | 0.032638 |
| H | -1.892541 | 2.087317 | -1.628892 |
| H | -2.259813 | -0.314643 | -1.689017 |
| C | -3.894743 | -2.410867 | -1.789782 |
| H | -3.693716 | -1.915028 | -2.753309 |
| H | -4.587817 | -3.248406 | -1.968400 |
| C | -2.614040 | -2.930539 | -1.187464 |
| C | -1.419086 | -2.710675 | -1.731979 |
| H | -2.699791 | -3.521587 | -0.269772 |
| C | -0.298571 | -2.308639 | -2.206395 |
| H | 0.392211 | -2.756802 | -2.923983 |
| H | 0.062118 | -1.141756 | -1.774998 |
| O | -4.606500 | -1.534712 | -0.918473 |
| H | -3.666803 | 3.641413 | -0.778857 |

**TS7_Q**

E=-3226.459867
E_SP=-3228.489148
H=-3227.972012
G=-3228.064805
Imag. Freq. -1876.87

Cartesian coordinates

| | | | |
|---|---|---|---|
| Fe | 1.108201 | 0.183648 | 0.145681 |
| N | 0.882124 | -1.772231 | 0.643317 |
| N | -0.582449 | 0.623842 | 1.127850 |
| N | 2.907175 | -0.249409 | -0.666527 |
| N | 1.392070 | 2.128968 | -0.246574 |
| C | 1.766587 | -2.798484 | 0.397364 |
| C | -1.101954 | 1.872028 | 1.357490 |
| C | -0.207720 | -2.337545 | 1.244467 |
| C | -1.500259 | -0.271373 | 1.616539 |
| C | 3.504031 | -1.482563 | -0.748779 |
| C | 0.606381 | 3.175715 | 0.160744 |
| C | 3.721869 | 0.621185 | -1.344004 |
| C | 2.410462 | 2.676430 | -0.984048 |
| C | 1.216517 | -4.046638 | 0.872524 |
| C | -2.376662 | 1.763991 | 2.029610 |
| C | -0.012778 | -3.762238 | 1.392803 |
| C | -2.633857 | 0.432640 | 2.171581 |
| C | 4.744086 | -1.384155 | -1.482079 |

| | | | |
|---|---|---|---|
| C | 1.132140 | 4.423000 | -0.349699 |
| C | 4.876651 | -0.077638 | -1.857606 |
| C | 2.247218 | 4.111444 | -1.070976 |
| H | 1.719292 | -5.009469 | 0.800858 |
| H | -2.998097 | 2.610546 | 2.312562 |
| H | -0.735156 | -4.441219 | 1.842510 |
| H | -3.506281 | -0.049415 | 2.607795 |
| H | 5.410963 | -2.220939 | -1.682510 |
| H | 0.687913 | 5.400399 | -0.169355 |
| H | 5.677328 | 0.388435 | -2.429263 |
| H | 2.920374 | 4.778466 | -1.606780 |
| C | 2.987937 | -2.671030 | -0.243825 |
| C | -0.537576 | 3.069433 | 0.940872 |
| C | 3.491067 | 1.981726 | -1.510135 |
| C | -1.341455 | -1.649028 | 1.664086 |
| H | 3.580556 | -3.576017 | -0.383863 |
| H | -1.064084 | 3.990116 | 1.195075 |
| H | 4.231591 | 2.552237 | -2.072440 |
| H | -2.150396 | -2.231217 | 2.106167 |
| O | 0.290563 | -0.042965 | -1.351386 |
| S | 2.352430 | 0.252348 | 2.128530 |
| C | 1.226079 | -0.036985 | 3.518319 |
| H | 1.820222 | -0.032779 | 4.443936 |
| H | 0.443636 | 0.732648 | 3.569001 |
| H | 0.744424 | -1.022086 | 3.415888 |
| C | -4.293358 | -0.199723 | -0.927159 |
| C | -5.271669 | 0.682111 | -0.438687 |
| C | -5.024796 | 2.053926 | -0.408673 |
| C | -3.803083 | 2.564976 | -0.862054 |
| C | -2.828509 | 1.682638 | -1.330986 |
| C | -3.060980 | 0.305661 | -1.366705 |
| H | -6.223286 | 0.267332 | -0.099865 |
| H | -5.796803 | 2.729654 | -0.031256 |
| H | -1.852308 | 2.049620 | -1.653142 |
| H | -2.259751 | -0.348079 | -1.698556 |
| C | -3.916653 | -2.423066 | -1.781409 |
| H | -3.692115 | -1.946740 | -2.749759 |
| H | -4.620726 | -3.251788 | -1.958244 |
| C | -2.654732 | -2.949087 | -1.147791 |
| C | -1.445746 | -2.756150 | -1.671997 |
| H | -2.764681 | -3.512331 | -0.214646 |
| C | -0.307074 | -2.385901 | -2.126672 |
| H | 0.398397 | -2.852553 | -2.816633 |
| H | 0.060936 | -1.173818 | -1.702573 |
| O | -4.633354 | -1.521098 | -0.938862 |
| H | -3.607138 | 3.639112 | -0.832247 |

**TS8**

E=-3226.482105

E_SP=-3228.5128

H=-3227.99072

G=-3228.08373

Imag. Freq. -126.28

Cartesian coordinates

| | | | |
|---|---|---|---|
| Fe | 0.931741 | -0.208334 | 0.308025 |
| N | 2.415097 | -1.192974 | -0.633279 |
| N | -0.217937 | -1.872640 | 0.222631 |
| N | 2.056916 | 1.449694 | 0.377121 |
| N | -0.514011 | 0.741618 | 1.338654 |
| C | 3.661049 | -0.706333 | -0.923814 |
| C | -1.434847 | -2.057681 | 0.826776 |
| C | 2.347427 | -2.449950 | -1.180652 |
| C | 0.076996 | -3.040459 | -0.435334 |
| C | 3.350647 | 1.577495 | -0.064879 |
| C | -1.695400 | 0.193402 | 1.783010 |
| C | 1.695142 | 2.674337 | 0.876523 |
| C | -0.521173 | 2.055835 | 1.733345 |
| C | 4.422795 | -1.698316 | -1.649138 |
| C | -1.930845 | -3.382619 | 0.536727 |
| C | 3.602382 | -2.776546 | -1.819002 |
| C | -0.999071 | -3.989225 | -0.256990 |
| C | 3.813922 | 2.931453 | 0.146568 |
| C | -2.470060 | 1.194950 | 2.476992 |
| C | 2.782793 | 3.614474 | 0.721412 |
| C | -1.746038 | 2.352139 | 2.440248 |
| H | 5.446554 | -1.560704 | -1.993039 |
| H | -2.880903 | -3.775547 | 0.893717 |
| H | 3.810113 | -3.716988 | -2.326673 |
| H | -1.014824 | -4.991840 | -0.680981 |
| H | 4.805707 | 3.292155 | -0.120941 |

| | | | |
|---|---|---|---|
| H | -3.450884 | 1.018092 | 2.913465 |
| H | 2.744404 | 4.657191 | 1.031789 |
| H | -2.000054 | 3.328104 | 2.850299 |
| C | 4.116613 | 0.576755 | -0.642708 |
| C | -2.122202 | -1.107879 | 1.570670 |
| C | 0.493547 | 2.974804 | 1.500272 |
| C | 1.259897 | -3.310047 | -1.113000 |
| H | 5.135708 | 0.827583 | -0.939612 |
| H | -3.095330 | -1.389282 | 1.971751 |
| H | 0.347954 | 3.994102 | 1.859073 |
| H | 1.355969 | -4.284563 | -1.593289 |
| O | 0.267788 | 0.267838 | -1.313336 |
| S | 1.687541 | -0.875003 | 2.325018 |
| C | 2.400799 | -2.538269 | 2.154753 |
| H | 2.716333 | -2.862932 | 3.157520 |
| H | 1.652523 | -3.245940 | 1.769968 |
| H | 3.271285 | -2.536703 | 1.484111 |
| C | -3.418291 | 0.165202 | -1.475276 |
| C | -3.867493 | -1.156125 | -1.333965 |
| C | -4.942567 | -1.440552 | -0.496872 |
| C | -5.583620 | -0.415090 | 0.210016 |
| C | -5.133345 | 0.897422 | 0.062726 |
| C | -4.054511 | 1.198881 | -0.776323 |
| H | -3.346522 | -1.944844 | -1.878694 |
| H | -5.279898 | -2.474457 | -0.390771 |
| H | -5.617350 | 1.707928 | 0.612780 |
| H | -3.712001 | 2.229641 | -0.855393 |
| C | -1.947545 | 1.650554 | -2.693906 |
| H | -2.847557 | 2.247867 | -2.938802 |
| H | -1.375072 | 1.525476 | -3.623730 |
| C | -1.108200 | 2.342516 | -1.661069 |
| C | -0.156948 | 3.284002 | -2.001944 |
| H | -1.361642 | 2.210622 | -0.609118 |
| C | 0.688192 | 4.118541 | -2.313428 |
| H | 1.442410 | 4.838104 | -2.571534 |
| H | -0.575490 | -0.180140 | -1.503681 |
| O | -2.346910 | 0.339883 | -2.304426 |
| H | -6.425123 | -0.639887 | 0.868637 |

**TS9**

E=-3226.457049

E_SP=-3228.484482

H=-3227.962674

G=-3228.056101

Imag. Freq. -511.79

Cartesian coordinates

| | | | |
|---|---|---|---|
| Fe | -0.620608 | 0.315694 | -0.371496 |
| N | -2.444403 | -0.489822 | 0.025964 |
| N | -0.042377 | -1.471038 | -1.154034 |
| N | -1.252689 | 2.102432 | 0.288496 |
| N | 1.097168 | 1.134500 | -0.963827 |
| C | -3.473305 | 0.133530 | 0.679039 |
| C | 1.206108 | -1.779588 | -1.624177 |
| C | -2.864539 | -1.768977 | -0.222900 |
| C | -0.790693 | -2.621451 | -1.254530 |
| C | -2.434394 | 2.359061 | 0.935041 |
| C | 2.173527 | 0.481779 | -1.516538 |
| C | -0.572660 | 3.288197 | 0.241720 |
| C | 1.447160 | 2.462735 | -0.898622 |
| C | -4.587261 | -0.774233 | 0.835882 |
| C | 1.259872 | -3.169665 | -2.018667 |
| C | -4.210172 | -1.957558 | 0.267735 |
| C | 0.016507 | -3.689504 | -1.801933 |
| C | -2.493275 | 3.751285 | 1.328098 |
| C | 3.225512 | 1.425229 | -1.815944 |
| C | -1.339590 | 4.331648 | 0.888037 |
| C | 2.769174 | 2.656280 | -1.447168 |
| H | -5.528079 | -0.521739 | 1.322037 |
| H | 2.142467 | -3.663488 | -2.421430 |
| H | -4.773061 | -2.886019 | 0.189335 |
| H | -0.339014 | -4.702517 | -1.982435 |
| H | -3.326427 | 4.209291 | 1.858619 |
| H | 4.187417 | 1.156334 | -2.246841 |
| H | -1.018636 | 5.367656 | 0.982447 |
| H | 3.276326 | 3.617401 | -1.509618 |
| C | -3.468486 | 1.451420 | 1.124826 |
| C | 2.250770 | -0.877613 | -1.779590 |
| C | 0.675295 | 3.473337 | -0.342510 |
| C | -2.109309 | -2.761919 | -0.841718 |
| H | -4.362121 | 1.809965 | 1.638149 |

| | | | |
|---|---|---|---|
| H | 3.189634 | -1.256880 | -2.183071 |
| H | 1.091830 | 4.481654 | -0.338130 |
| H | -2.576898 | -3.738274 | -0.976185 |
| O | -0.074511 | -0.069247 | 1.213046 |
| S | -1.273327 | 0.735810 | -2.616069 |
| C | -2.874329 | -0.034411 | -2.965491 |
| H | -3.172901 | 0.257893 | -3.983782 |
| H | -2.774649 | -1.131241 | -2.937538 |
| H | -3.645978 | 0.270070 | -2.245622 |
| C | 3.121489 | -1.083224 | 1.433329 |
| C | 4.341816 | -1.495923 | 0.878466 |
| C | 5.347954 | -0.562312 | 0.628944 |
| C | 5.151474 | 0.787584 | 0.938048 |
| C | 3.928697 | 1.191864 | 1.480455 |
| C | 2.907773 | 0.271191 | 1.724917 |
| H | 4.477847 | -2.555456 | 0.653112 |
| H | 6.295402 | -0.895014 | 0.196989 |
| H | 3.748524 | 2.248488 | 1.692031 |
| H | 1.931358 | 0.602861 | 2.073901 |
| C | 1.206623 | -1.934656 | 2.637606 |
| H | 1.406296 | -1.084142 | 3.305857 |
| H | 1.257883 | -2.856681 | 3.242086 |
| C | -0.182608 | -1.832503 | 2.046716 |
| C | -1.264706 | -1.689997 | 2.844792 |
| H | -0.307522 | -2.379577 | 1.109790 |
| C | -2.294961 | -1.285517 | 3.536355 |
| H | -2.847451 | -1.944562 | 4.218294 |
| H | -2.657658 | -0.251720 | 3.439237 |
| O | 2.196509 | -2.063774 | 1.634332 |
| H | 5.937813 | 1.519824 | 0.742272 |
